## Supplementary material for "Who infects Whom? - Reconstructing infection chains of *Mycobacterium avium* ssp. *paratuberculosis* in an endemically infected dairy herd by use of genomic data": S1 Figure

Supporting information S1 Fig.

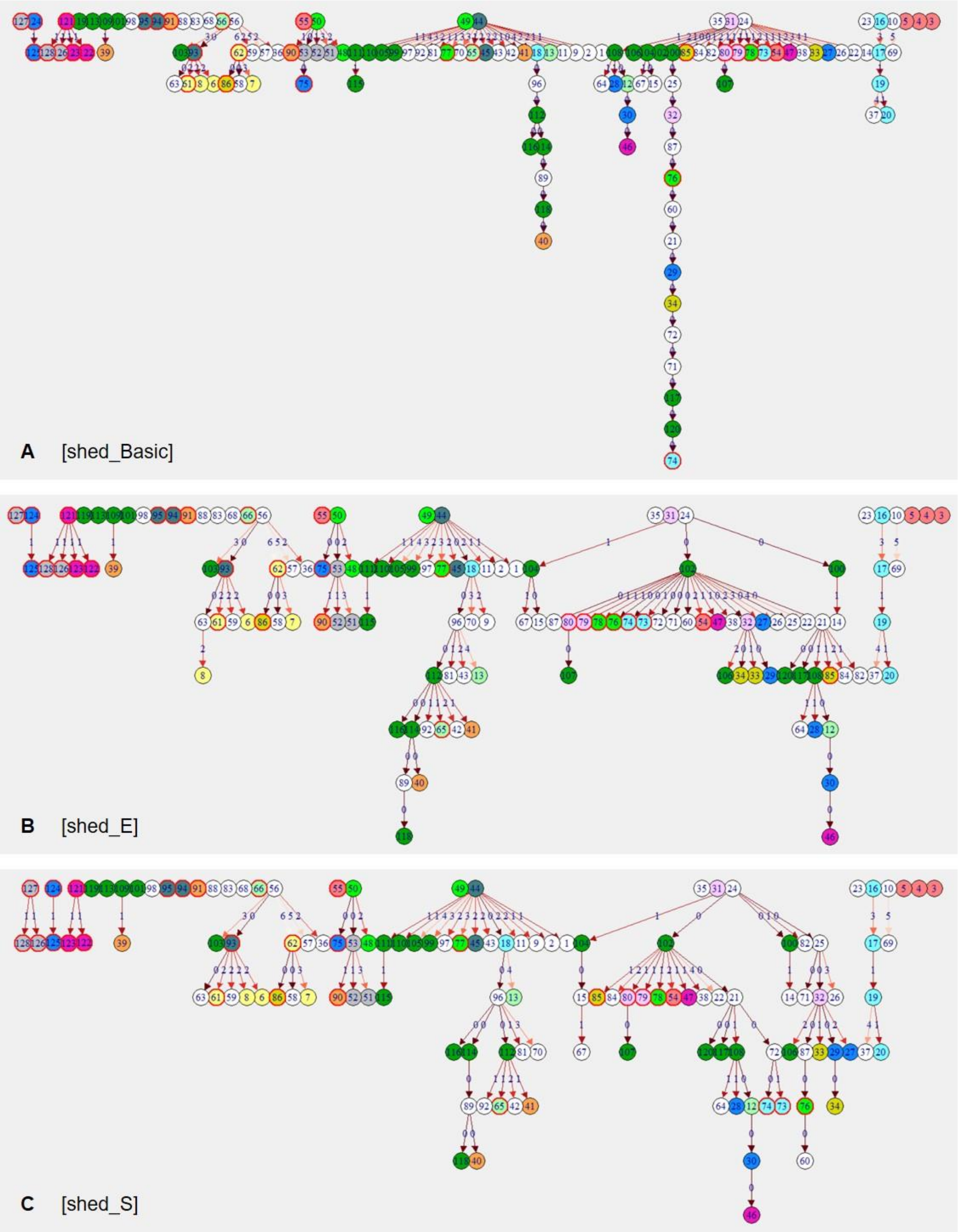


**S1 Fig. Reconstructed transmission tree of three scenarios (n = 128 isolates). (A)** [shed_Basic], **(B)** [shed_E], **(C)** [shed_S]. Isolates sampled from the same cow are labelled with successive numbers and are shown in vertices of same colour and outline. White vertices represent cows with only one isolate. Dark green vertices (labelled 99–120) represent environmental samples. Edge labels and edge colour indicate number of SNPs difference between ancestor and descendant.
