## Supplementary material for "Who infects Whom? - Reconstructing infection chains of *Mycobacterium avium* ssp. *paratuberculosis* in an endemically infected dairy herd by use of genomic data": S2 Figure

Supporting information S2 Fig.

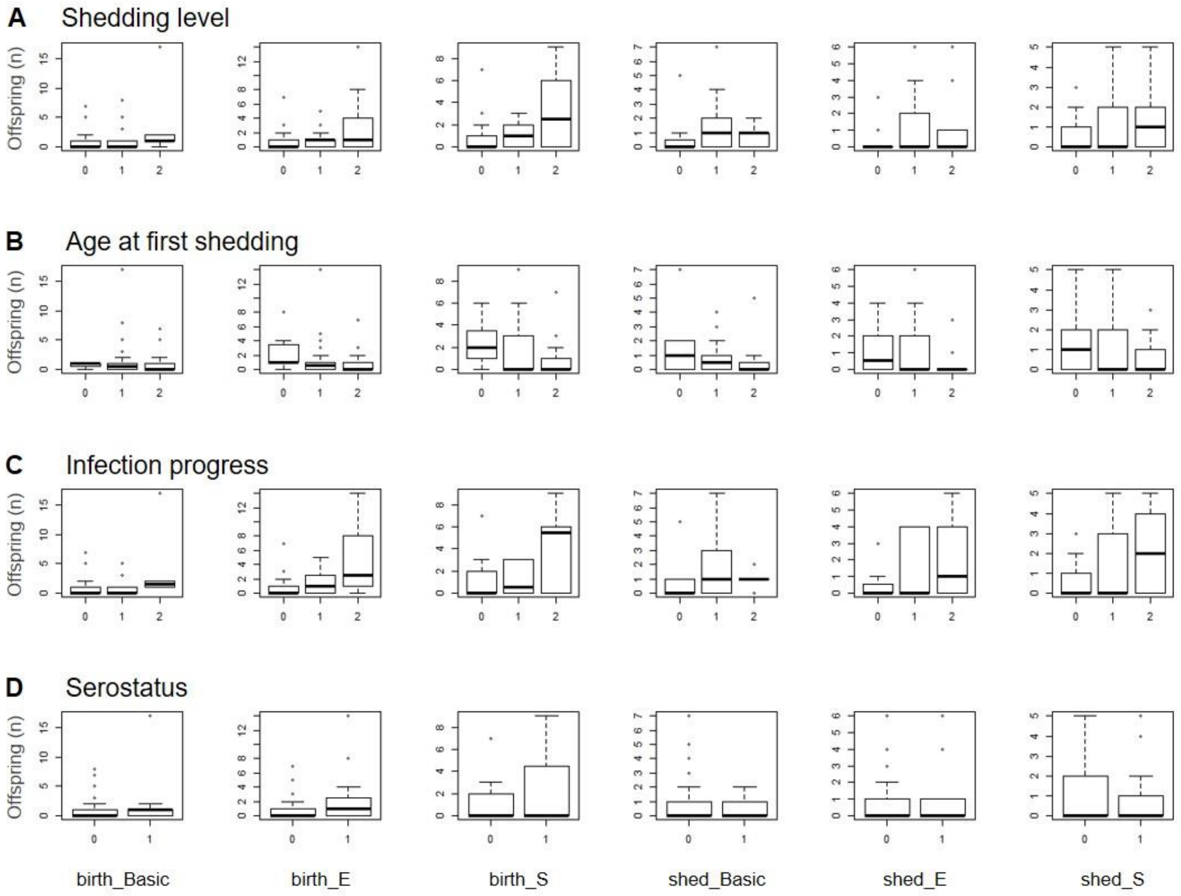


**S2 Fig. Numbers of recipients produced by individual cows.** Boxplots with numbers of recipients produced by individual cows, by disease phenotype and scenario. **(A)** shedding level (0 – always faecal culture negative, 1 - low, 2 – high), **(B)** age at first shedding (0 - ≤3 years, 1 - >3 years, 2 - ante mortem negative), **(C)** infection progress (0 - ante mortem negative, 1 - non-progressor, 2 - progressor), **(D)** serostatus (0 - ELISA-negative, 1 - ELISA-positive). Scenarios (from left-most to right-most column): [birth_Basic], [birth_E], [birth_S], [shed_Basic], [shed_E] and [shed_S].
