## Supplementary material for "Who infects Whom? - Reconstructing infection chains of *Mycobacterium avium* ssp. *paratuberculosis* in an endemically infected dairy herd by use of genomic data": S1 File.fasta

>MAP 009_A_tb1045
CCCCTTAGGCGCCTCGAGCGGGCCCCCGCTTCGCCAGTTCACCCGTCCCAGTAAGCGCCC
GGGGCACCTCGCCTTCGAGGAGAACCCCCAGAACGCCCCACCGTCTCCCAGCTCGCACAC
GCGCACGCGTCGCACTCTCCAGCGCGCGTACTGTTCCAGCCACCGTCCCCATGCTCCCCG
CGGGGCCGTATGACGAGCCACGCCCCGGCCCCGCCGCCGCCGCTCGCCCACCGAGACAAG
CGCAATGCTATTGGGGCCTGACCACGGCTGGGCGGGGCGCTTCCCACGGAGGGCCCAGTC
TTTGGGAGGAGTTGCCCCTCCAGGGATCGGTAGCGCGGTGCGGCTCCCCGCAGCGGCCGC
AGGCGTCGGGGGGATGCTGGGGAGTGGGGGATGCCCCTGCCGGTCGCAGGCGGAACAGGT
TCGCGGTGGTGCACAGGGGCGTGGTGCTGCAGCTCGAGGCCGCGGAGGGGGGCCGCGGTG
GACTTCGGGTCGGAGGGACCGGGATACCAGGGCTCTGGCGCGGGGTAGCCCCAGGACGCT
TTCTCGGGGAGGGGAGGGTACGGGCAGCCAGGTGGGAGTGCCAGAACTGGACGGCGTAAT
CCCTGCTGCCTGCGATGTGGGGTCGGACCCAGCGCACCCTGATGGGGCCTGGCGGCGGGG
GACCCCTGAACGGACGTGGCAGCTACTCTGCCGTTGAGCCGATGCGGACGTGCCCAGTAC
ACGTGGGTGTTGGGGGAACTCCACAGCGGCAGGCGAGGGTTGGGTGTAGCAGGGGGCCCA
CGGGGGCGGCGGGTGGCAGGCGCGAGGGGGGCGCGGGGGAGCGATCCGATGGTCGAGAAC
GATCGGCTAGGAGGCGAGCCCCAGGTGGCACCGGGCGGGTGCGCCCGTGTCGGCGCGGGG
GGGTTGTGGCGACGAGGCCGCCCTTCAGGCGAGGTGGGAGTGGCCCAAGTTGTGGCGCCG
CGGTGGAGGGGGGGCGGTGAGGTGGGGGGCCAGTCGAGGTTGGTGAGGGCGGGACCAGCC
TCGTTTAGTGAAACGCCAACCCGGATAGTAATTCGTGCGGCTCGTGGGAGCGAGTTCTTC
CAATGATGCAGGGGGGGGTGTGGGGCCCGCGCGTCGACCGGTCGCGTCACGACTGCTACG
GGGTCATTACCGCCGGGCTGTCGTAGGCCGGCGCTCGCGGAGGGAGGTTCGAATGCCGGG
GCGGGAGGCCCGGATCCGGCGAAGGGGCGGGGCTCGGGGGACGCGAGGCGGAGGGGCGCG
AGGCGGAAGCCGGTGCAAGCGGCGCGTGGCCCCATGAGGAAGAACTCTCTCGTGCCCCAA
CACCACACATTAGCGCCCCCATCTCGGCCGACTTCGAGCACGCTCAACGAGAACCACCAC
AGGCCTCCAACCCGACGTACCACCCGTCCTCACGAGGCGGGCCGACCCTCGGGTCCAAGC
GAGGGCTCCGCCGCGCACGCCCGGCGTGGTGC
>MAP 055_A_tb1058
CCCCTTAGGCGCCTCGAGCGGGCCCCCGCTTCGCCAGTTCACCCGTCCCAGTAAGCGCCC
GGGGCACCTCGCCTTCGAGGAGAACCCCCAGAACGCCCCACCGTCTCCCAGCTCGCACAC
GCGCACGCGTCGCACTCTCCAGCGCGCGTACTGTTCCAGCCACCGTCCCCATGCTCCCCG
CGGGGCCGTATGACGAGCCACGCCCCGGCCCCGCCGCCGCCGCTCGCCCACCGAGACAAG
CGCAATGCTATTGGGGCCTGACCACGGCTGGGCGGGGCGCTTCCCACGGAGGGCCCAGTC
TTTGGGAGGAGTTGCCCCTCCAGGGATCGGTAGCGCGGTGCGGCTCCCCGCAGCGGCCGC
AGGCGTCGGGGGGATGCTGGGGAGTGGGGGATGCCCCTGCCGGTCGCAGGCGGAACAGGT
TCGCGGTGGTGCACAGGGGCGTGGTGCTGCAGCTCGAGGCCGCGGAGGGGGGCCGCGGTG
GACTTCGGGTCGGAGGGACCGGGATACCAGGGCTCTGGCGCGGGGTAGCCCCAGGGCGCT
TTCTCGGGGAGGGGAGGGTACGGGCAGCCAGGTGGGAGTGCCAGAACTGGACGGCGTAAT
CCCTGCTGCCTGCGATGTGGGGTCGGACCCAGCGCACCCTGATGGGGCCTGGCGGCGGGG
GACCCCTGAACGGACGTGGCAGCTACTCTGCCGTTGAGCCGATGCGGACGTGCCCAGTAC
ACGTGGGTGTTGGGGGAACTCCACAGCGGCAGGCGAGGGTTGGGTGTAGCAGGGGGCCCA
CGGGGGCGGCGGGTGGCAGGCGCGAGGGGGGCGCGGGGGAGCGATCCGATGGTCGAGAAC
GATCGGCTAGGAGGCGAGCCCCAGGTGGCACCGGGCGGGTGCGCCCGTGTCGGCGCGGGG
GGGTTGTGGCGACGAGGCCGCCCTTCAGGCGAGGTGGGAGTGGCCCAAGTTGTGGCGCCG
CGGTGGAGGGGGGGCGGTGAGGTGGGGGGCCAGTCGAGGTTGGTGAGGGCGGGACCAGCC
TCGTTTAGTGAAACGCCAACCCGGATAGTAATTCGTGCGGCTCGTGGGAGCGAGTTCTTC
CAATGATGCAGGGGGGGGTGTGGGGCCCGCGCGTCGACCGGTCGCGTCACGACTGCTACG
GGGTCATTACCGCCGGGCTGTCGTAGGCCGGCGCTCGCGGAGGGAGGTTCGAATGCCGGG
GCGGGAGGCCCGGATCCGGCGAAGGGGCGGGGCTCGGGGGACGCGAGGCGGAGGGGCGCG
AGGCGGTAGCCGGTGCAAGCGGCGCGTGGCCCCATGAGGAAGAACTCTCTCGTGCCCCAA
CACCACACATTAGCGCCCCCATCTCGGCCGACTTCGAGCACGCTCAACGAGAACCACCAC
AGGCCTCCAACCCGACGTACCACCCGTCCTCACGAGGCGGGCCGACCCTCGGGTCCAAGC
GAGGGCTCCGCCGCGCACGCCCGGCGTGGTGC
>MAP 010_A_tb1076
CCCCTTAGGCGCCTCGAGCGGGCCCCCGCTTCGCCAGTTCACCCGTCCCAGTAGGCGCCC
GCGGCCCCTCGCCTTCGCGGAGAACCCCCAGAACGCCCCACCGTCTCCCAGCTCGCACAC
GCGTACGCGTCGCACTCTCCAGCGCGCGGACTGTTCCAGCCACCGACCCCATGCTCCCCG
CGGGGCCGTATTCTGACCCACGCCCTGGCCCCGCCGCCGCCGCTCGCCTACTGAGACAAG
CGCAACGCTATTGCGGCCTGACCACGACTGGTCGGGGCGCCTCCCACGGAGGGCCCAGTC
TTTGGGAGGAGTCGCCCCTCAAGGGATCGGTAGCGCGATGCGGCCCCCCGCAGCGGCCGC
GGATGTCGGGGTGATGTTGGGGAGTAGGAGGTGCCCCTTCCGGCCACAGGCGGAACAGGT
TCGCGGAGGTGCACAGGGGCGTGGTGCTGCAGCTCGAGGCCGCGGAGGGGGGCCGCGGTG
GACTTCGGGTCGGAGGGACCGATATACCAGGGCTCTGGCGTGGGGTAGCCCCAGGACGCT
CTCTCAGGGAGGGGAGGGTACGGGCAGCTAGGTGGGAGTGCAGGAACTGGACGACGTAAT
CCCTGGTGACTGCGATATGAGGTAGGACCCAGCGCACCCTGATGGGGCCTGGCGGCGGGG
GACCCCTGGACGGACGTGGCAGCTACCCTGCCGTTGGGCCGATGCGGACTTGCCCAGTAC
ACGTGGGTGTTGGGGAAACTCCACAGCGGCAGGCGAGGGTTGGGTGTGGCCGGGGGACCA
CGGAGGCGGGGGGTGGTGGTCGCGAGGGGGGCGCGGGGGAGCGATCCGATGGTCAAGAAC
GATCGGCTAGGAGGCGAGCCCCAGGTGGCGTAGGGCGGGTGCGCCGGTATCGGCGCGGGG

```
AGGTTGTGGTGACGAGGCCGGTTTTCAGGCGGGGTGGAGTGGCCCAAGTTGTGGCGCCG
CGGTGGAAGGGGGGCGGTGAGGTAGGGGGCCAGTCGGGGTTGATGAGGGCGGGACCAGCC
TCGCTTGGTGCAACGCCGACCCGGATGGTAATTCGCGCGGCTCGTGAGCGCGAGTTCTTC
CGATGATTCAGGAGGAGGTGTGGGGCCCGCGCGACGACCAGCCGCGTCACGACTTCTATG
GGGTCATTACCACCGGGCTGTCGTAGGCCGGCGACCGCGGGAGGTGGTTCGAATGTCGGG
GCGGGAGGCCCGGCTCAGGCGAAGGGGCGGGGCCCAGGGCACGCGAGGCGGGGGGGCGCG
AGGCGGTAGCCGGTGCCAGCGGCGCGTGGCCCCATGAGGTAGAACTCTCTCGTGCCCCAA
CACCACACATTAGCCCCCCCATCTCGGCCGACTTCGAGCGCGCTCGGCGAGAACCACCAC
AGGCCTCCAACCCGACGTACCACCCGTCCTCACGAGGCAGGCTGACCCCCGGGCACAAGC
GAGGGATCCGGCAAGCACGCCCGGCGTGGTGC
>MAP 011_A_tb1076
CCCCTCAGGCGCCCCAAGCGGGCCCCCGCTTCGCCAGCTCACCCGTTCCAGTAAGCGCCC
ACGGCCCCTCGCCTTCGCGGGGGACCCCCAGAACGCCCCACCGTCTCCCAGCTCGCACGC
GCACACTCGTCGCACCCTCCAGCGCGCGGACTGCTCCAGCAGTCATCCCCATGCTCCCCG
CGGGGCTGCACGACGAGCCACGCCTCGGCACCGCCGCTACCGCCCACCCATCGAGACAAG
CGCAACGCTATTGGGGCCTAACCACGGCTGGGTGGGGCGCTTCCCACGGAGGGCTCCGTC
CTCGGGAGGAGCTGCCCCTCAAGGGATCGGTAGCGCGGTGCGGTTCCCCGCAACGGCCGT
GGGCGTCGGGGTGATCCCGAGGAGTGGGGGATGCCCCTTCCGGCAGAGGGCGGAGCGGGT
TCGCAGTGGTGCACAGGGGCGTGGTGCCGCGGCCTGGGGCGGCGGCCGGGGGCCGCGGTG
GACTTCGGGTCGGATGGACCGATATGGCACGGCTCTGGCGCGGCGTGGCCCCAGGACGCT
CTCTCGGGGACGGGAGGGTACGGGCAACCAGGTGGGAGCGCAGGAGCTGGACGGCGTAAT
CCCTGGTGACTGCGACGTGGGGTAGGACCCAGGGCACCCTGGCGATGCCTGGCGGCGGGG
GCCCCCCGGACGGACGCGCCGGCTACCCTGCTGTTGGGACGGTACAGACGTACCCAGTAC
ACGTGGGTATTGGGGGAACTCTGGAGCGGCAGGCGAGGGTTAGGTGTGAGCGGGGGCCCC
CGGGGGCGGCGGGGGGCGGTCGCGAGGAGGGCGCGAGGAAGTGATGCGACGGTCGAGAAC
GATCGGCCAGGAGGCGAGCCCCAGCTCCCGCAGGGCGGGTGCGCCCGTGTCCGCGCGGGG
GGGTTGTGGCGACGAGGCCGCCCCTCAGGCGGGTCGGGAGTGGCCCAAGTTGTGGCGCTG
CGGTGGAGGGGGGGCAGTGAGGCGGAGGGTGAGTCGGGGTTCGTGCAGGCGGGACCAGCC
TCGCTTGGCGCATGGCCAAGCCGGATAGTAATTCGTGCGACTCGCAGGCACGAGTTCTTC
CAATGATTCAGGGGGGGGTGTGAGGCCCCCGCGTCGGCCGGCCGCGTCACGAGTGCCGCA
GGGTCATTACCGCCGGACCACCGTAGACCGGCAATTGCAGGAGGTGGTTCGAATGCCGGG
CCGGGAGGCCCGGATTAGGCGGAGGGGCGGGGCCCGGGGCACGCGAGGCACGGGGGCGCG
AAGCGGTAGCCGGTGCAAGCGGCCCGTGGCCCCATGAGGTAGAACTCTCTCGTGCCCCAA
CACCACACATTAGCGCCCACGCCTCGTCCGACCTCAAGCGTACTCAGCGGGAACCGCCAC
AGGCCTCTAACCCTACGTTCCACCCGTCCTCACGAGGCGGTCCGACCCCCGGGCCCGAGC
GAGGGCTTCGGCGAGCACGCCCGGCATGGCGC
>MAP 017_A_tb1076
CCCCTTAGGCGCCTCGAGCGGGCCCCTGCCTCGCCAGTTCACCCGTCCCAGTAAGCGCCT
GCGGCCCCTCGCCTTCGCGGAGAACCCCCAGAACGCCCCACCGTATCCCTGCTCGCACAC
GCGCACGCGTCGCACTCTCCAGCGTGCGGACTGTTTCAGCCACCGTCCCCATGCTCCCCG
CGGGGCCGTATGACGAGCCACGCCCCGGCCCCGTCGCCGCGGCTCGCCCACCGAGACAAG
CGCAACGCCATTGGGGCCTGACCACGGCTAGGCGGGGCGCTTCCCACGGAGAACCCAGCC
TTTGGGAGGAGTTGCCCCTCAGGGGATCGGTAGCGCGGTACGGCTCCCCGCAGCGGCCGC
GGGCGTCGGGGTGATGCTGGGGAGGGGGGGATGCCCCTTCCGGCCGCAGGCGGAACAGGT
TCGCGGTGGTGCACAGGGGCGTGGGGCTGCAGCTCGAGGCCGCAGAGGGGGGCCGCGGTG
GACTTCGGGTCGGAGGGACCGATATACCAGGGCTCTGGCGCGGGGTAGCCCCAGGACGCT
CTCCCGGGGAGGGGAGGGTACGGGCAGCCAAGTGGGAGTGCAGGAACTGGACGGCGTAAT
CCCCGGTGAATGCGATGTTGGGGAGGACCCAGCGCACCCTGATGGGGCCTGACGGCGGGG
GATCCCTGGACCGACGTGGCAGCTACCCTGCCGTTGGGCCGATGCGGACGTGCCCAGTAC
ACGTGGATGTTGGGGGAACTCCACAGCGGCGGGCGAGTATTGGGTGTGGCCGGGGGCCCA
CAGGGGCGAGGGGTGACGGTCGCGAGGGGGGCGTGGGGGAGCGATCCGATGGTCGAAGAC
GATCGGCTAGGAGGCGAGACCCAGGCGGCGCAGGGCGGGTGCGCCCGTGTCGGCGCGGGG
GGGTTCAGGCGACGAGGCCGCCCTTCAGGCGGGGTGGGAGTGGTCCAAGTTGTGGCGCCG
GGGCGGAGGGGGGGCGGTGAGGTGGGGGGCCAGTCGGGGTTGGTGAGGACGGGACCAGCC
TTGCTTGGTGCAACGCCAGCCCGGATAGTAATTCGTGCGGCTCGTGGGCGCGGGTTCTTC
CAATGATTCAGGGGGGGGTGTGGGGCCCGCGCGTCGACCGGCCGCGTCACCACTGCTACG
GGGTCATTACCGCCGGGCTGTCGTAGGCCGGCGATCGCGGGAGGTGGTTCGAATGCCGGG
GCCGGAGGCCCGAATCAGGCGAAGGGGCGGGGTCCGGGGCACGCGAGGCGGGGGGGCGCG
AGGCGGTAGCCGGTGCAAGCGGCGCCTGGCCCTATGAGGTAGAACTCTCTCGTCCCCCAA
CACCACACACTAGCGCCTCCATCTCGGCCGACTTCGAGCGCGCTCAGGGAGAACCACCTC
AGGCCTCCAACCCGACGTACCGCCCGTCCTCACGATGCGGGCCGACCCCCGGGCCTAAGC
CAGGGCTCCGGCGAGCACGCCCGGCGTGGTAC
>MAP 190_A_fb1077
CCCCTTAGGCGCCTCGAGCGGGCCCCCGCTTCGCCAGTTCACCCGTCCCAGTAAGCGCCC
GGGGCACCTCGCCTTCGAGGAGAACCCCCAGAACGCCCCACCGTCTCCCAGCTCGCACAC
GCGCACGCGTCGCACTCTCCAGCGCGCGTACTGTTCCAGCCACCGTCCCCATGCTCCCCG
CGGGGCCGTATGACGAGCCACGCCCCGGCCCCGCCGCCGTCGCTCGCCCGCCGAGACAAG
CGCAATGCTATTGGGGCCTGATCACGGCTGGGCGGGGCGCTTCCCACGGAGGGCCCAGTC
```

TTTGGGAGGAGTTGCCCCTCCAGGGATCGGTAGCGCGGTGCGGCTCCCCGCAGCGGCCGC
AGGCGTCGGGGGGATGCTGGGGAGTGGGGGATGCCCCTTCCGGTCGCAGGCGGAACAGGT
TCGCGGTGGTGCACAGGGGCGTGGTGCTGCAGCTCGAGGCCGCGGAGGGGGGCCGCGGTG
GACTTCGGGTCGGAGGGACCGATATACCAGGGCTCTGGCGCGGGGTAGCCCCAGGACGCT
TTCTCGGGGAGGGGAGGGTACGGGCAGCCAGGTGGGAGTGCCAGAACTGGACGGCGTAAT
CCCTGCTGCCTGCGATGTGGGGTCGGACCCAGCGCACCCTGATGGGGCCTGGTGGCGGGG
GACCCCTGAACGGACGTGGCAGATACTCTGCCGTTGAGCCGATGCGGACGTGCCCAGTAC
ACGTGGGTGTTGGGGGAACTCCACAGCGGCAGGCGAGGGTTGGGTGTAGCAGGGGGCCCA
CGGGGGCGGCGGGTGGCAGGCGCGAGGGGGGCGCGGGGGAGCGATCCGATGGTCGAGAAC
GATCGGCTAGGAGGCGAGCCCCAGGTGGCACCGGGCGGGTGCGCCCGTGTCGGCGCGGGG
GGGTTGTGGCGACGAGGCCGCCCTTCAGGCGAGGTGGGAGTGGCCCAAGTTGTGGCGCCG
CGGTGGAGGGGGGGCGGTGAGGTGGGGGGCCAGTCGAGGTTGGTGAGGGCGGGACCAGCC
TCGTTTAGTGAAACGCCAACCCGGATAGTAATTCGTGCGGCTCGTGGGAGCGAGTTCTTC
CAATGATGCAGGGGGGGGTGTGGGGCCCGCGCGTCGACCGGTCGCGTCACGACTGCTACG
GGGTCATTACCGCCGGGCTGTCCTAGGCCGGCGCTCGCGGGAGGTGGTTCGAATGCCGGG
GCGGGAGGCCCGGATCCGGCGAAGGGGCGGGGCTCGGGGGACGCGAGGCGGAGGGGCGCG
AGGCGGTAGCCGGTGCAAGCGGCGCGTGGCCCCATGAGGAAGAACTCTCTCGTGCCCCAA
CACCACACATTAGCGCCCCCATCTCGGCCGACTTCGAGCACGCTCAACGAGAACCACCAC
AGGCCTCCAACCCGACGTACCACCCGTCCTCACGAGGCGGGCCGACCCTCGGGTCCAAGC
GAGGGCTCCGCCGCGTATGCCCGGCGTGGTGC
>MAP 202_A_fb1077
CCCCTTAGGCGCCTCGAGCGGGCCCCCGCTTCGCCAGTTCACCCGTCCCAGTAAGCGCCC
GGGGCACCTCGCCTTCGAGGAGAACCCCCAGAACGCCCCACCGTCTCCCAGCTCGCACAC
GCGCACGCGTCGCACTCTCCAGCGCGCGTACTGTTCCAGCCACCGTCCCCATGCTCCCCG
CGGGGCCGTATGACGAGCCACGCCCCGGCCCCGCCGCCGCCGCTCGCCCACCGAGACAAG
CGCAATGCTATTGGGGCCTGACCACGGCTGGGCGGGGCGCTTCCCACGGAGGGCCCAGTC
TTTGGGAGGAGTTGCCCCTCCAGGGATCGGTAGCGCGGTGCGGCTCCCCGCAGCGGCCGC
AGGCGTCGGGGGGATGCTGGGGAGTGGGGGATGCCCCTTCCGGTCGCAGGCGGAACAGGT
TCGCGGTGGTACACAGGGGCGTGGTGCTGCAGCTCGAGGCCGCGGAGGGGGGCCGCGGTG
GACTTCGGGTCGGAGGGACCGATATACCAGGGCTCTGGCGCGGGGTAGCCCCAGGACGCT
TTCTCGGGGAGGGGAGGGTACGGGCAGCCAGGTGGGAGTGCCAGAACTGGACGGCGTAAT
CCCTGCTGCCTGCGATGTGGGGTCGGACCCAGCGCACCCTGATGGGGCCTGGCGGCGAGG
GACCCCTGAACGGACGTGGCAGCTACTCTGCCGTTGAGCCGATGCGGACGTGCCCAGTAC
ACGTGGGTGTTGGGGGAACTCCACAGCGGCAGGCGAGGGTTGGGTGTAGCAGGGGGCCCA
CGGGGGCGGCGGGTGGCAGGCGCGAGGGGGGCGCGGGGGAGCGATCCGATGGTCGAGAAC
GATCGGCTAGGAAGCGAGCCCCAGGTGGCACCGGGCGGGTGCGCCCGTGTCGGCGCGGGG
GGGTTGTGGCGACGAGGCCGCCCTTCAGGCGAAGTGGGAGTGGCCCAAGTTGTGGCGCCG
CGGTGGAGGGGGGGCGGTGAGGTGGGGGGCCAGTCGAGGTTGGTGAGGGCGGGACCAGCC
TCGTTTAGTGAAACGCCAACCCGGATAGTAATTCGTGCGGCTCGTGGGAGCGAGCTCTTC
CAATGATGCAGGGGGGGGTGTGGGGCCCGCGCGTCGACCGGTCGCGTCACGACTGCTACG
GGGTCATTACCGCCGGGCTGTCGTAGGCCGGCGCTCGCGGGAGGTGGTTCGAATGCCGGG
GCGGGAGGCCCGGATCCGGCGAAGGGGCGGGGCTCGGGGGACGCGAGGCGGAGGGGCGCG
AGGCGGTAGCCGGTGCAAGCGGCGCGTGGCCCCATGAGGAAGAACTCTCTCGGGCCCCAA
CACCACACATTAGCGCCCCCATCTCGGCCGACTTCGAGCACGCTCAACGAGAACCACCAC
AGGCCTCCAACCCGACGTACCACCCGTCCTCACGAGGCGGGCCGACCGTCGGGTCCAAGC
GAGGGCTCCGCCGCCCACGCCCGGCGTGGTGC
>MAP 207_A_fb1077
CCCCTTAGGCGCCTCGAGCGGGCCCCCGCTTCGCCAGTTCACCCGTCCCAGTAAGCGCCC
GGGGCACCTCGCCTTCGAGGAGAACCCCCAGAACGCCCCACCGTCTCCCAGCTCGCACAC
GCGCACGCGTCGCACTCTCCAGCGCGCGTACTGTTCCAGCCACCGTCCCCATGCTCCCCG
CGGGGCCGTATGACGAGCCACGCCCCGGCCCCGCCGCCGTCGCTCGCCCGCCGAGACAAG
CGCAATGCTATTGGGGCCTGACCACGGCTGGGCGGGGCGCTTCCCACGGAGGGCCCAGTC
TTTGGGAGGAGTTGCCCCTCCAGGGATCGGTAGCGCGGTGCGGCTCCCCGCAGCGGCCGC
AGGCGTCGGGGGGATGCTGGGGAGTGGGGGATGCCCCTTCCGGTCGCAGGCGGAACAGGT
TCGCGGTGGTGCACAGGGGCGTGGTGCTGCAGCTCGAGGCCGCGGAGGGGGGCCGCGGTG
GACTTCGGGTCGGAGGGACCGATATACCAGGGCTCTGGCGCGGGGTAGCCCCAGGACGCT
TTCTCGGGGAGGGGAGGGTACGGGCAGCCAGGTGGGAGTGCCAGAACTGGACGGCGTAAT
CCCTGCTGCCTGCGATGTGGGGTCGGACCCAGCGCACCCTGATGGGGCCTGGCGGCGGGG
GACCCCTGAACGGACGTGGCAGATACTCTGCCGTTGAGCCGATGCGGACGTGCCCAGTAC
ACGTGGGTGTTGGGGGAACTCCACAGCGGCAGGCGAGGGTTGGGTGTAGCAGGGGGCCCA
CGGGGGCGGCGGGTGGCAGGCGCGAGGGGGGCGCGGGGGAGCGATCCGATGGTCGAGAAC
GATCGGCTAGGAGGCGAGCCCCAGGTGGCACCGGGCGGGTGCGCCCGTGTCGGCGCGGGG
GGGTTGTGGCGACGAGGCCGCCCTTCAGGCGAGGTGGGAGTGGCCCAAGTTGTGGCGCCG
CGGTGGAGGGGGGGCGGTGAGGTGGGGGGCCAGTCGAGGTTGGTGAGGGCGGGACCAGCC
TCGTTTAGTGAAACGCCAACCCGGATAGTAATTCGTGCGGCTCGTGGGAGCGAGTTCTTC
CAATGATGCAGGGGGGGGTGTGGGGCCCGCGCGTCGACCGGTCGCGTCACGACTGCTACG
GGGTCATTACCGCCGGGCTGTCCTAGGCCGGCGCTCGCGGGAGGTGGTTCGAATGCCGGG
GCGGGAGGCCCGGATCCGGCGAAGGGGCGGGGCTCGGGGGACGCGAGGCGGAGGGGCGCG

AGGCGGTAGCCGGTGCAAGCGGCGCGTGGCCCCATGAGGAAGAACTCTCTCGTGCCCCAA
CACCACACATTAGCGCCCCCATCTCGGCCGACTTCGAGCACGCTCAACGAGAACCACCAC
AGGCCTCCAACCCGACGTACCACCCGTCCTCACGAGGCGGGCCAACCCTCGGGTCCAAGC
GAGGGCTCCGCTGCGTATGCCCGGCGTGGTGC
>MAP 092_A_fb1078
CCCCTTAGGCGCCTCGAGCGGGCCCCCGCTTCGCCAGTTCACCCGTCCCAGTAAGCGCCC
GGGGCACCTCGCCTTCGAGGAGAACCCCCAGAACGCCCCACCGTCTCCCAGCTCGCAGAC
GCGCACGCGTCGCACTCTCCAGCGCGCGTACTGTTCCAGCCACCGTCCCCATGCTCCCCG
CGGGGCCGTATGACGAGCCACGCCCCGGCCCCGCCGCCGCCGCTCGCCCACCGAGACAAG
CGCAATGCTATTGGGGCCTGACCACGGCTGGGCGGAGCGCTTCCCACGGAGGGCCCAGTC
TTTGGGAGGAGTTGCCCCTCCAGGGATCGGTAGCGCGGTGCGGCTCCCCGCAGCGGCCGC
AGGCGTCGGGGGGATGCTGGGGAGTGGGGGATGCCCCTGCCGGTCGCAGGCGGAACAGGT
TCGCGGTGGTGCACAGGGGCGTGGTGCTGCAGCTCGAGGCCGCGGAGGGGGGCCGCGGTG
GACTTCGGGTCGGAGGGACCGGGATACCAGGGCTCTGGCGCGGGGTAGCCCCAGGACGCT
TTCTCGGGGAGGGGAGGGTACGGGCAGCCAGGTGGGAGTGCCAGAACTGGACGGCGTAAT
CCCTGCTGCCTGCGATGTGGGGTCGGACCCAGCGCACCCTGATGGGGCCTGGCGGCGGGG
GACCCCTGAACGGACGTGGCAGCTACTCTGCCGTTGAGCCGATGCGGACGTGCCCAGTAC
ACGTGGGTGTTGGGGGAACTCCACAGCGGCAGGCGAGGGTTGGGTGTAGCAGGGGGCCCA
CGGGGGCGGCGGGTGGCAGGCGCGAGGGGGGCGCGGGGGAGCGATCCGATGGTCGAGAAC
GATCGGCTAGGAGGCGAGCCCCAGGTGGCACCGGGCGGGTGCGCCCGTGTCGGCGCGGGG
GGGTTGTGGCGACGAGGCCGCCCTTCAGGCGAGGTGGGAGTGGCCCAAGTTGTGGCGCCG
CGGTGGAGGGGGGGCGGTGAGGTGGGGGGCCAGTCGAGGTTGGTGAGGGCGGGACCAGCC
TCGTTTAGTGAAACGCCAACCCGGATAGTAATTCGTGCGGCTCGTGGGAGCGAGTTCTTC
CAATGATGCAGGGGGGGGTGTGGGGCCCGCGCGTCGACCGGTCGCGTCACGACTGCTACG
GGGTCATTACCGCCGGGCTGTCGTAGGCCGGCGCTCGCGGAGGGAGGTTCGAATGCCGGG
GCGGGAGGCCCGGATCCGGCGAAGGGGCGGGGCTCGGGGGACGCGAGGCGGAGGGGCGCG
AGGCGGTAGCCGGTGCAAGCGGCGCGTGGCCCCATGAGGAAGAACTCTCTCGTGCCCCAA
CACCACACATTAGCGCCCCCATCTCGGCCGACTTCGAGCACGCTCAACGAGAACCACCAC
AGGCCTCCAACCCGACGTACCACCCGTCCTCACGAGGCGGGCCGACCCTCGGGTCCAAGC
GAGGGCTCCGCCGCGCACGCCCGGCGTGGTGC
>MAP 022_A_fb1085
CCCCTTAGGCGCCTCGAGCGGGCCCCCGCCTCGCCAGTTCACCCGTCCCAGTAAGCGCCT
GCGGCCCCTCGCCTTCGCGGAGAACCCCCAGAACGCCCCACCGTATCCCTGCTCGCACAC
GCGCACGCGTCGCACTCTCCAGCGCGCGGACTGTCCCGCCCACCGTCCCCATCCTCCCCG
CGGGGCCGTATGACGAGCCACGCCCCGGCCCCGCCGCCGCCGCTCGCCCACCGAGACAAG
CGCAACGCTATTGGGGCCTGACCGCGGCTGGGCGGGGCGCTTCCCACGGAGAACCCAGCC
TTTGGAAGGAATTGCCCCTCAGGGGGTCGGTAGCGCGGTGCGGCTCCCCGCGGCGGCCGC
GGGCGTCGGGGTGATGCTGGGGAGTGGGGGATGCCCCTTCCGGCCGCAGGCGGAACACGT
TCGCGGTGGTGCACAGGGGCGTGGTGCTACAGCTCGAGGCCGCAGAGGGGGGCCGCGGTG
GACTTCGGGTCCGAGGGACCGATATACCAGGGCTCTGGCGCGGGGTAGCCCCAGGACGCT
CTCCCGGGGAGGGGAGGGTACGGGCAGCCAGGTGGGAGTGCAGGAACTGGACGGCGTAAT
CCCTGGTGACTGCGATGTTGGGGAGGACCCAGCGCACCCTGATGGGGCCTGACGGCGGGG
GATCCCTGGACCGACGTGGCAGCTACCCTACCGTTGGGCCGATGCGGACGCGCCCAGTAC
ACGTGGGTGTTGGGGGAACGCCACAGCGGCGGGCGAGTGTTGGGTGTGGCCGGGGGCCCA
CAGGGGCGAGGGGTGACGGTCGCGAGGGGGGCGTGGGGGAGCGATCCGGTGGTCGAGAAC
GATCGGCTAGGAGGCGAGACCCAGGCGGCGCAGGGCGGGTGCGCCCGTGTCGGCGCGGGG
GGGTTGTGGCGACGAGGCCGCCCTTCAGGCGGGGTGGGAGTGGCCCAAGCTGCGGAGCCG
CGGCGGAGGGGGGGCGGTGAGGTGGGGGGCCAGTCGGGGTTGGTGAGGACGGGACCAGCC
TTGCTCGGTGCAACGCCAGCCCGGATAGTAATTCGTGCGGCTCGTGGGCGCGGGTTCTTC
CAATGATTCAGGGGGGCGTGTGGGGCCCGCGCGTCGACCGGCCGCGTCACGACTGCTACG
GGGTCATTACCGCCGGGCTGTCGTAGGCCGGCGATCGCGGGAGGTGGTTCGAATGCCGGG
GCCGGAGGCCCGAATCAGGCGAAGGGGCGGGGTCCGGGGCACGCGAGGCGGGGGGGCGCG
AGGCGGTGGCCGGTGCAAGCGGCGCCTGGCCCCATGAGGTAGAACTCTCTCGTCCCCCAA
CACCACACACTAGCGCCTCCATCTCGGCCGACTTCGAGCGCGCTCAGGGAGAATCACCAC
AGGCCTCCAACCCGACGTACCACCCGTCCTCACGAGGCGGGCCGACCCCCGGGCCCAAGC
GAGGGCTCCGGCGAGCACGCCCGGCGAGGTAC
>MAP 025_A_fb1099
CCCCTTAGGCGCCTCGAGCGGGCCCCCGCTTCGCCAGTTCACCCGTCCCAGTAAGCGCCC
GGGGCACCTCGCCTTCGAGGAGAACCCCCAGAACGCCCCACCGTCTCCCAGCTCGCACAC
GCGCACGCGTCGCACTCTCCAGCGCGCGTACTGTTCCAGCCACCGTCCCCATGCTCCCCA
CGGGGCCGTATGACGAGCCACGCCCCGGCCCCGCCGCCGCCGCTCGCCCACCGAGACAAG
CGCAATGCTATTGGGGCCTGACCACGGCTGGGCGGGGCGCTTCCCACGGAGGGCCCAGTC
TTTGGGAGGAGTTGCCCCTCCAGGGATCGGTAGCGCGGTGCGGCTCCCCGCAGCGGCCGC
AGGCGTCGGGGGGATGCTGGGGAGTGGGGGATGCCCCTGCCGGTCGCAGGCGGAACAGGT
TCGCGGTGGTGCACAGGGGCGTGGTGCTGCAGCTCGAGGCCGCGGAGGGGGGCCGCGGTG
GACTTCGGGTCGGAGGGACCGGGATACCAGGGCTCTGGCGCGGGGTAGCCCCAGGACGCT
TTCTCGGGGAGGGGAGGGTACGGGCAGCCAGGTGGGAGTGCCAGAACTGGACGGCGTAAT
CCCTGCTGCCTGCGATGTGGGGTCGGACCCAGCGCACCCTGATGGGGCCTGGCGGCGGAG

```
GACCCCTGAACGGACGTGGCAGCTACTCTGCCGTTGAGCCGATGCGGACGTGCCCAGTAC
ACGTGGGTGTTGGGGGAACTCCACAGCGGCAGGCGAGGGTTGGGTGTAGCAGGGGGCCCA
CGGGGGCGGCGGGTGGCAGGCGCGAGGGGGGCGCGGGGGAGCGATCCGATGGTCGAGAAC
GATCGGCTAGGAGGCGAGCCCCAGGTGGCACCGGGCGGGTGCGCCCGTGTCGGCGCGGGG
GGGTTGTGGCGACGAGGCCGCCCTTCAGGCGAGGTGGGAGTGGCCCAAGTTGTGGCGCCG
CGGTGGAGGGGGGGCGGTGAGGTGGGGGGCCAGTCGAGGTTGGTGAGGGCGGGACCAGCC
TCGTTTAGTGAAACGCCAACCCGGATAGTAATTCGTGCGGCTCGTGGGAGCGAGTTCTTC
CAATGATGCAGGGGGGGGTGTGGGGCCCGCGCGTCGACCGGTCGCGTCACGACTGCTACG
GGGTCATTACCGCCGGGCTGTCGTAGGCCGGCGCTCGCGGAGGGAGGTTCGAATGCCGGG
GCGGGAGGCCCGGATCCGGCGAAGGGGCGGGGCTCGGGGGACGCGAGGCGGAGGGGCGCG
AGGCGGTAGCCGGTGCAAGCGGCGCGTGGCCCCATGAGGAAGAACTCTCTCGTGCCCCAA
CACCACACATTAGCGCCCCCATCTCGGCCGACTTCGAGCACGCTCAACGAGAACCACCAC
AGGCCTCCAACCCGACGTACCACCCGTCCTCACGAGGCGGGCCGACCCTCGGGTCCAAGC
GAGGGCTCCGCCGCGCACGCCCGGCGTGGTGC
>MAP 149_A_tb1101
CCCCTTAGGCGCCTCGAGCGGGCCCCCGCTTCGCCAGTTCAGCCGTCCCAGTAAGCGCCC
GGGGCACCTCGCCTTCGAGGAGAACCCCCAGAACGCCCCACCGTCTCCCAGCTCGCACAC
GCGCACGCGTCGCACTCTCCCGCGCGCGTACTGTTCCAGCCACCGTCCCCATGCTCCCCG
CGGGGCCGTATGACGAGCCACGCCCCGGCCCCGCCGCCGCCGCTCGCTCACCGAGACCAG
CGCAATGCTATTGGGGCCTGACCACGGCTGGGCGGGGCGCTTCCCACGGAGGGCCCAGTC
TTTGGGAGGAGTTGCCCCTCCAGGGATCGGCCGCGCGGTGCGGCTCCCCGCAGCGGCCGC
GGGCGTCGGGGGGATGCTAGGGAGTGGGGGATGCCCCTTCCGGTCGCAGGCGGAACAGGT
TCGCGGTGGTGCACAGGGGCGTGGTGCTGCAGCTCGAGGCCGCGGAGGGGGGCCGCGGTG
GACTTCGGGTCGGAGGGACCGATATACCAGGGCTCTGGCGCGGGGTAGCCCCAGGACGCT
TTCTCGGGGAGGGGAGGGTACGGGCAGCCAGGTGGGAGTGCAAGAACTGGACGGCCTAAT
CCCTGCTGCCTGCGATGTGGGGTCGGACCCAGCGCACCCTGATGGGGCCTGGCGGCGGGG
GACCCCTGAACGGACGTGGCAGCTACTCTGCCGTTGAGCCGATGCGGACGTGCCCAGTAC
ACGTGGGTGTTGGGGGAACTCCACAGCGGCAGGCGAGGGTTGGGTGTAGCAGGGGGCCCA
CGGGGGCGGCGGGTGGCAGGCGCGAGGGGGGCGCGGGGGAGCGATCCGATGGTCGAGAAC
GATCGGCTAGGAGGCGAGCCCCAGGTGGCGCCGGGCGGGTGCGCCCGTGTCGGCGCGGGG
GGGTTGTGGCGACGAGGCCGCCCTTCAGGCGAGGTGGGAGTGGCCCACGTTGTGGCGCCG
CGGTGGAGGGGGGGCGGTGAGGTGGGGGGCCAGTCGAGGTTGGTGAGGGCGGGACCAGCC
TCGCTTAGTGCAACGCCAACCCGGATAGTAATTCGTGCGGCTCGTGGGAGCGAGTTCTTC
CAATGATGCAGGGGGGGGTGTGGGGCCCGCGCGTCGACCGGCCGCGTCACGACTGCTACG
GGGTCATTACCGCCGGGCTGTCGTAGGCCGGCGCTCGCGGGAGGTGGTTCGAATGCCGGG
GCGGGAGGCCCGGATCCGGCCAAGGGGCGGGGCTCGGGGGACGCGAGGCGGAGGGGCGCG
AGGCGGTAGCCGGTGCAAGCGGCGCGTGGCCCCATGAGGAAGAACTCTCTCGTGCCCCAA
CACCACACATTAGCGCCCCCATCTCGGCCGACTTCGAGCACGCTCAGCGAGAACCACCAC
AGGCCTCCAACCCGACGTACCACCCGTCCTCACGAGGCGGGCCGACCCTCGGGTCCAAGC
GAGGGCTCCGGCGCGCACGCCCGGCGTGGTGC
>MAP 158_A_tb1101
CCCCTTAGGCGCCTCGAGCGGGCCCCCGCTTCGCCAGTTCACCCGTCCCAGTAAGCGCCC
GGGGCACCTCGCCTTCGAGGAGAACCCCCAGAACGCCCCACCGTCTCCCAGCTCGCACAC
GCGCACGCGTCGCACTCTCCAGCGCGCGTACTGTTCCAGCCACCGTCCCCATGCTCCCCG
CGGGGCCGTATGACGAGCCACGCCCCGGCCCCGCCGCCGCCGCTCGCCCACCGAGACAAG
CGCAATGCTATTGGGGCCTGACCACGGCTGGGCGGGGCGCTTCCCACCGAGGGCCCAGTC
TTTGGGAGGAGTTGCCCCTCCAGGGATCGGTAGCGCGGTGCGGCTCCCCGCAGCGGCCGC
AGGCGTCGGGGGGATGCTGGGGAGTGGGGGATGCCCCTGCCGGTCGCAGGCGGAACAGGT
TCGCGGTGGTGCACAGGGGCGTGGTGCTGCAGCTCGAGGCCGGGGAGGGGGGCCGCGGTG
GACTTCGGGTCGGAGGGACCGGGATACCAGGGCTCTGGCGCGGGGTAGCCCCAGGACGCT
TTCTCGGGGAGGGGAGGGTACGGGCAGCCAGGTGGGAGTGCCAGAACTGGACGGCGTAAT
CCCTGCTGCCTGCGATGTGGGGTCGGACCCAGCGCACCCTGATGGGGCCTGGCGGCGGGG
GACCCCTGAACGGACGTGGCAGCTACTCTGCCGTTGAGCCGATGCGGACGTGCCCAGTAC
ACGTGGGTGTTGGGGGAACTCCACAGCGGCAGGCGAGGGTTGGGTGTAGCAGGGGGCCCA
CGGGGGCGGCGGGTGGCAGGCGCGAGGGGGGCGCGGGGGAGCGATCCGATGGTCGAGAAC
GATCGGCTAGGAGGCGAGCCCCAGGTGGCACCGGGCGGGTGCGCCCGTGTCGGCGCGGGG
GGGTTGTGGCGACGAGGCCGCCCTTCAGGCGAGGTGGGAGTGGCCCAAGTTGTGGCGCCG
CGGTGGAGGGGGGGCGGTGAGGTGGGGGGCCAGTCGAGGTTGGTGAGGGCGGGACCAGCC
TCGTTTAGTGAAACGCCAACCCGGATAGTAATTCGTGCGGCTCGTGGGAGCGAGTTCTTC
CAATGATGCAGGGGGGGGTGTGGGGCTCGCGCGTCGACCGGTCGCGTCACGACTGCTACG
GGGTCATTACCGCCGGGCTGTCGTAGGCCGGCGCTCGCGGAGGGAGGTTCGAATGCCGGG
GCGGGAGGCCCGGATCCGGCGAAGGGGCGGGGCTCGGGGGACGCGAGGCGGAGGGGCGCG
AGGCGGTAGCCGGTGCAAGCGGCGCGTGGCCCCATGAGGAAGAACTCTCTCGTGCCCCAA
CACCACACATTAGCGCCCCCATCTCGGCCGACTACGAGCACGCTCAACGAGAACCACCAC
AGGCCTCCAACCCGACGTACCACCCGTCCTCACGAGGCGGGCCGACCCTCGGGTCCAAGC
GAGGGCTCCGCCGCGCACGCCCGGCGTGGTGC
>MAP 061_A_tb1111
CCCCTTAGGCGCCTCGAGCGGGCCCCCGCTTCGCCAGTTCAGCCGTCCCAGTAAGCGCCC
```

```
GGGGCACCTCGCCTTCGAGGAGAACCCCCAGAACGCCCCACCGTCTCCCAGCTCGCACAC
GCGCACGCGTCGCACTCTCCCGCGCGCGTACTGTTCCAGCCACCGTCCCCATGCTCCCCG
CGGGGCCGTATGACGAGCCACGCCCCGGCCCCGCCGCCGCCGCTCGCTCACCGAGACCAG
CGCAATGCTATTGGGGCCTGACCACGGCTGGGCGGGGCGCTTCCCACGGAGGGCCCAGTC
TTTGGGAGGAGTTGCCCCTCCAGGGATCGGCAGCGCGGTGCGGCTCCCCGCAGCGGCCGC
GGGCGTCGGGGGGATGCTAGGGAGTGGGGGATGCCCCTTCCGGTCGCAGGCGGAACAGGT
TCGCGGTGGTGCACAGGGGCGTGGTGCTGCAGCTCGAGGCCGCGGAGGGGGGCCGCGGTG
GACTTCGGGTCGGAGGGACCGATATACCAGGGCTCTGGCGCGGGGTAGCCCCAGGACGCT
TTCTCGGGGAGGGGAGGGTACGGGCAGCCAGGTGGGAGTGCAAGAACTGGACGGCCTAAT
CCCTGCTGCCTGCGATGTGGGGTCGGACCCAGCGCACCCTGATGGGGCCTGGCGGCGGGG
GACCGCTGAACGGACGTGGCAGCTACTCTGCCGTTGAGCCGATGCGGACGTGCCCAGTAC
ACGTGGGTGTTGGGGGAACTCCACAGCGGCAGGCGAGGGTTGGGTGTAGCAGGGGGGCCCA
CGGGGGCGGCGGGTGGCAGGCGCGAGGGGGGCGCGGGGGAGCGATCCGATGGTCGAGAAC
GATCGGCTAGGAGGCGAGCCCCAGGTGGCGCCGGGCGGGTGCGCCCGTGTCGGCGCGGGG
GGGTTGTGGCGACGAGGCCGCCCTTCAGGCGAGGTGGGAGTGGCCCACGTTGTGGCGCCG
CGGTGGAGGGGGGGCGGTGAGGTGGGGGGCCAGTCGAGGTTGGTGAGGGCGGGACCAGCC
TCGCTTAGTGCAACGCCAACCCGGATAGTAATTCGTGCGGCTCGTGGGAGCGAGTTCTTC
CAATGATGCAGGGGGGGGTGTGGGGCCCGCGCGTCGACCGGCCGCGTCACGACTGCTACG
GGGTCATTACCGCCGGGCTGTCGTAGGCCGGCGCTCGCGGGAGGTGGTTCGAATGCCGGG
GCGGGAGGCCCGGATCCGGCCAAGGGGCGGGGCTCGGGGGACGCGAGGCGGAGGGGCGCG
AGGCGGTAGCCGGTGCAAGCGGCGCGTGGCCCCATGAGGAAGAACTCTCTCGTGCCCCAA
CACCACACATTAGCGCCCCCATCTCGGCCGACTTCGAGCACGCTCAGCGAGAACCACCAC
AGGCCTCCAACCCGACGTACCACCCGTCCTCACGAGGCGGGCCGACCCTCGGGTCCAAGC
GAGGGCTCCGGCGCGCACGCCCGGCGTGGTGC
>MAP 082_A_tb1120
CCCCTTAGGCGCCTCGAGCGGGCCCCCGCTTCGCCAGTTCAGCCGTCCCAGTAAGCGCCC
GGGGCACCTCGCCTTCGAGGAGAACCCCCAGAACGCCCCACCGTCTCCCAGCTCGCACAC
GCGCACGCGTCGCACTCTCCCGCGCGCGTACTGTTCCAGCCACCGTCCCCATGCTCCCCG
CGGGGCCGTATGACGAGCCACGCCCCGGCCCCGCCGCCGCCGCTCGCTCACCGAGACCAG
CGCAATGCTATTGGGGCCTGACCACGGCTGGGCGGGGCGCTTCCCACGGAGGGCCCAGTC
TTTGGGAGGAGTTGCCCCTCCAGGGATCGGCAGCGCGGTGCGGCTCCCCGCAGCGGCCGC
GGGCGTCGGGGGGATGCTAGGGAGTGGGGGATGCCCCTTCCGGTCGCAGGCGGAACAGGT
TCGCGGTGGTGCACAGGGGCGTGGTGCTGCAGCTCGAGGCCGCGGAGAGGGGCCGCGGTG
GACTTCGGGTCGGAGGGACCGATATACCAGGGCTCTGGCGCGGGGTAGCCCCAGGACGCT
TTCTCGGGGAGGGGAGGGTACGGGCAGCCAGGTGGGAGTGCAAGAACTGGACGGCCTAAT
CCCTGCTGCCTGCGATGTGGGGTCGGACCCAGCGCACCCTGATGGGGCCTGGCGGCGGGG
GACCCCTGAACGGACGTGGCAGCTACTCTGCCGTTGAGCCGATGCGGACGTGCCCAGTAC
ACGTGGGTGTTGGGGGAACTCCACAGCGGCAGGCGAGGGTTGGGTGTAGCAGGGGGGCCCA
CGGGGGCGGCGGGTGGCAGGCGCGAGGGGGGCGCGGGGGAGCGATCCGATGGTCGAGAAC
GATCGGCTAGGAGGCGAGCCCCAGGTGGCGCCGGGCGGGTGCGCCCGTGTCGGCGCGGGG
GGGTTGTGGCGACGAGGCCGCCCTTCAGGCGAGGTGGGAGTGGCCCACGTTGTGGCGCCG
CGGTGGAGGGGGGGCGGTGAGGTGGGGGGCCAGTCGAGGTTGGTGAGGGCGGGACCAGCC
TCGCTTAGTGCAACGCCAACCCGGATAGTAATTCGTGCGGCTCGTGGGAGCGAGTTCTTC
CAATGATGCAGGGGGGGGTGTGGGGCCCGCGCGTCGACCGGCCGCGTCACGACTGCTACG
GGGTCATTACCGCCGGGCTGTCGTAGGCCGGCGCTCGCGGGAGGTGGTTCGAATGCCGGG
GCGGGAGGCCCGGATCCGGCCAAGGGGCGGGGCTCGGGGGACGCGAGGCGGAGGGGCGCG
AGGCGGTAGCCGGTGCAAGCGGCGCGTGGCCCCATGAGGAAGAACTCTCTCGTGCCCCAA
CACCACACATTAGCGCCCCCATCTCGGCCGACTTCGAGCACGCTCAGCGAGAACCACCAC
AGGCCTCCAACCCGACGTACCACCCGTCCTCACGAGGCGGGCCGACCCTCGGGTCCAAGC
GAGGGCTCCGGCGCGCACGCCCGGCGTGGTGC
>MAP 003_A_fb1127
CCCCTTAGGCGCCTCGAGCGGGCCCCCGCTTCGCCAGTTCACCCGTCCCAGTAAGCGCCC
GGGGCACCTCGCCTTCGAGGAGAACCCCCAGAACGCCCCACCGTCTCCCAGCTCGCACAT
GCGCACGCGTCGCACTCTCCAGCGCGCGTACTGTTCCAGCCACCGTCCCCATGCTCCCCG
CGGGGCCGTATGACGAGCCACGCCCCGGCCCCGCCGCCGCCGCTCGCCCACCGAGACAAG
CGCAATGCTATTGGGGCCTGACCACGGCTGGGCGGGGCGCTTCCCACGGAGGGCCCAGTC
TTTGGGAGGAGTTGCCCCTCCAGGAATCGGTAGCGCGGTGCGGCTCCCCGCAGCGGCCGC
AGGCGTCGGGGGGATGCTGGGGAGTGGGGGATGCCCCTTCCGGTCGCAGGCGGAACAGGT
TCGCGGTGGTGCACAGGGGCGTGGTGCTGCAGCTCGAGGCCGCGGAGGGGGGCCGCGGTG
GACTTCGGGTCGGAGGGACCGATATACCAGGGCTCTGGCGCGGGGTAGCCCCAGGACGCT
TTCTCGGGGAGGGGAGGGTACGGGCAGCCAGGTGGGAGTGCCAGAACTGGGCGGCGTAAT
CCCTGCTGCCTGCGATGTGGGGTCGGACCCAGCGCACCCTGATGGGGCCTAGCGGCGGGG
GACCCCTGAACGGACATGGCAGCTACTCTGCCGTTGAGCCGATGCGGACGTGCCCAGTAC
ACGTGGGTGTTGGGGGAACTCCACAGCGGCAGGCGAGGGTTGGGTGTAGCAGGGGGGCCCA
CGGGGGCGGCGGGTGGCAGGCGCGAGGGGGGCGCGGGGGAGCGATCCGATGGTCGAGAAC
GACCGGCTAGGAGGCGAGCCCCAGGTGGCACCGGGCGGGTGCGCCCGTGTCGGCGCGGGG
GGGTTGTGGCGACGAGGCCGCCCTTCAGGCGAGGTGGGAGTGGCCCAAGTTGTGGCGCCG
CGGTGGAGGGGGGGTGGTGAGGTGGGGGGCCAGTCGAGGTTGGTGAGGGCGGGACCAGCC
```

TCGTTTAGTGAAACGCCAACCCGGATAGTAATTCGTGCGGCTCGTGGGAGCGAGTTCTTC
CAATGATGCAGGGGGGGGTGTGGGGCCCGCGCGTCGACCGGTCGCGTCACGACTGCTACG
GGGTCATTACCGCCGGGCTGTCGTAGGCCGGCGCTCGCGGGAGGTGGTTCGAATGCCGGG
GCGGGAGGCCCGGATCCGGCGAAGGGGCGGGGCTCGGGGGACGCGAGGCGGAGGGGCGCG
AGGCGGTAGCCGGTGCAAGCGGCGCGTGGCCCCATGAGGAAGAACTCTCTCGTGCCCCAA
CACCACACATTAGCGCCCCCATCTCGGCCGACTTCGAGCACGCTCAACGAGAACCACCAC
AGGCCTCCAACCCGACGTACCACCTGTCCTCACGAGGCGGGCCGACCCTCGGGTCCAAGC
GAGGGCTCCGCCGCGCACGCCCGGCGTGGTGC
>MAP 028_A_fb1127
CCCCTTAGGCGCCTCGAGCGGGCCCCCGCTTCGCCAGTTCACCCGTCCCAGTAAGCGCCC
GGGGTACCTCGCCTTCGAGGAGAACCCCCAGAACGCCCCACCGTCTCCCAGCTCGCACAT
GCGCACGCGTCGCACTCTCCAGCGCGCGTACTGTTCCAGCCACCGTCCCCATGCTCCCCG
CGGGGCCGTATGACGAGCCACGCCCCGGCCCCGCCGCCGCCGCTCGCCCACCGAGACAAG
CGCAATGCTATTGGGGCCTGACCACGGCTGGGCGGGGCGCTTCCCACGGAGGGCCCAGTC
TTTGGGAGGAGTTGCCCCTCCAGGAATCGGTAGCGCGGTGCGGCTCCCCGCAGCGGCCGC
AGGCGTCGGGGGGATGCTGGGGAGTGGGGGATGCCCCTTCCGGTCGCAGGCGGAACAGGT
TCGCGGTGGTGCACAGGGGCGTGGTGCTGCAGCTCGAGGCCGCGGAGGGGGGCCGCGGTG
GACTTCGGGTCGGAGGGACCGATATACCAGGGCTCTGGCGCGGGGTAGCCCCAGGACGCT
TTCTCGGGGAGGGGAGGGTACGGGCAGCCAGGTGGGAGTGCCAGAACTGGACGGCGTAAT
CCCTGCTGCCTGCGATGTGGGGTCGGACCCAGCGCACCCTGATGGGGCCTGGCGGCGGGG
GACCCCTGAACGGACATGGCAGCTACTCTGCCGTTGAGCCGATGCGGACGTGCCCAGTAC
ACGTGGGTGTTGGGGGAACTCCACAGCGGCAGGCGAGGGTTGGGTGTAGCAGGGGGCCCA
CGGGGGCGGCGGGTGGCAGGCGCGAGGGGGGGCGCGGGGGAGCGATCCGATGGTCGAGAAC
GACCGGCTAGGAGGCGAGCCCCAGGTGGCACCGGGCGGGTGCGCCCGTGTCGGCGCGGGG
GGGTTGTGGCGACGAGGCCGCCCTTCAGGCGAGGTGGGAGTGGCCCAAGTTGTGGCGCCG
CGGTGGAGGGGGGGGTGGTGAGGTGGGGGGCCAGTCGAGGTTGGTGAGGGCGGGACCAGCC
TCGTTTAGTGAAACGCCAACCCGGATAGTAATTCGTGCGGCTCGTGGGAGCGAGTTCTTC
CAATGATGCAGGGGGGGGTGTGGGGCCCGCGCGTCGACCGGTCGCGTCACGACTGCTACG
GGGTCATTACCGCCGGGCTGTCGTAGGCCGGCGCTCGCGGGAGGTGGTTCGAATGCCGGG
GCGGGAGGCCCGGATCCGGCGAAGGGGCGGGGCTCGGGGGACGCGAGGCGGAGGGGCGCG
AGGCGGTAGCCGGTGCAAGCGGCGCGTGGCCCCATGAGGAAGAACTCTCTCGTGCCCCAA
CACCACACATTAGCGCCCCCATCTCGGCCGACTTCGAGCACGCTCAACGAGAACCACCAC
AGGCCTCCAACCCGACGTACCACCTGTCCTCACGAGGCGGGCCGACCCTCGGGTCCAAGC
GAGGGCTCCGCCGCGCACGCCCGGCGTGGTGC
>MAP 051_A_tb1127
CCCCTTAGGCGCCTCGAGCGGGCCCCCGCTTCGCCAGTTCACCCGTCCCAGTAAGCGCCC
GGGGCACCTCGCCTTCGAGGAGAACCCCCAGAACGCCCCACCGTCTCCCAGCTCGCACAC
GCGCACGCGTCGCACTCTCCAGCGCGCGTACTGTTCCAGCCACCGTCCCCATGCTCCCCG
CGGGGCCGTATGACGAGCCACGCCCCGGCCCCGCCGCCGCCGCTCGCCCACCGAGACAAG
CGCAATGCTATTGGGGCCTGACCACGGCTGGGCGGGGCGCTTCCCACGGAGGGCCCAGTC
TTTGGGAGGAGTTGCCCCTCCAGGGATCGGTAGCGCGGTGCGGCTCCCCGCAGCGGCCGC
AGGCGTCGGGGGGATGCTGGGGAGTGGGGGATGCCCCTGCCGGTCGCAGGCGGAACAGGT
TCGCGGTGGTGCACAGGGGCGTGGTGCTGCAGCTCGAGGCCGCGGAGGGGGGCCGCGGTG
GACTTCGGGTCGGAGGGACCGGGATACCAGGGCTCTGGCGCGGGGTAGCCCCAGGACGCT
TTCTCGGGGAGGGGAGGGTACGGGCAGCCAGGTGGGAGTGCCAGAACTGGACGGCGTAAT
CCCTGCTGCCTGCGATGTGGGGTCGGACCCAGCGCACCCTGATGGGGCCTGGCGGCGGGG
GACCCCTGAACGGACGTGGCAGCTACTCTGCCGTTGAGCCGATGCGGACGTGCCCAGTAC
ACGTGGGTGTTGGGGGAACTCCACAGCGGCAGGCGAGGGTTGGGTGTAGCAGGGGGCCCA
CGGGGGCGGCGGGTGGCAGGCGCGAGGGGGGGCGCGGGGGAGCGATCCGATGGTCGAGAAC
GATCGGCTAGGAGGCGAGCCCCAGGTGGCACCGGGCGGGTGCGCCCGTGTCGGCGCGGGG
GGGTTGTGGCGACGAGGCCGCCCTTCAGGCGAGGTGGGAGTGGCCCAAGTTGTGGCGCCG
CGGTGGAGGGGGGGCGGTGAGGTGGGGGGCCAGTCGAGGTTGGTGAGGGCGGGACCAGCC
TCGTTTAGTGAAACGCCAACCCGGATAGTAATTCGTGCGGCTCGTGGGAGCGAGTTCTTC
CAATGATGCAGGGGGGGGTGTGGGGCCCGCGCGTCGACCGGTCGCGTCACGACTGCTACG
GGGTCATTACCGCCGGGCTGTCGTAGGCCGGCGCTCGCGGAGGGAGGTTCGAATGCCGGG
GCGGGAGGCCCGGATCCGGCGAAGGGGCGGGGCTCGGGGGACGCGAGGCGGAGGGGCGCG
AGGCGGTAGCCGGTGCAAGCGGCGCGTGGCCCCATGAGGAAGAACTCTCTCGTGCCCCAA
CACCACACATTAGCGCCCCCATCTCGGCCGACTTCGAGCACGCTCAACGAGAACCACCAC
AGGCCTCCAACCCGACGTACCACCCGTCCTCACGAGGCGGGCCGACCCTCGGGTCCAAGC
GAGGGCTCCGCCGCGCACGCCCGGCGTGGTGC
>MAP 054_A_tb1127
CCCCTTAGGCGCCTCGAGCGGGCCCCCGCTTCGCCAGTTCACCCGTCCCAGTAAGCGCCC
GGGGCACCTCGCCTTCGAGGAGAACCCCCAGAACGCCCCACCGTCTCCCAGCTCGCACAT
GCGCACGCGTCGCACTCTCCAGCGCGCGTACTGTTCCAGCCACCGTCCCCATGCTCCCCG
CGGGGCCGTATGACGAGCCACGCCCCGGCCCCGCCGCCGCCGCTCGCCCACCGAGACAAG
CGCAATGCTATTGGGGCCTGACCACGGCTGGGCGGGGCGCTTCCCACGGAGGGCCCAGTC
TTTGGGAGGAGTTGCCCCTCCAGGAATCGGTAGCGCGGTGCGGCTCCCCGCAGCGGCCGC
AGGCGTCGGGGGGATGCTGGGGAGTGGGGGATGCCCCTTCCGGTCGCAGGCGGAACAGGT

TCGCGGTGGTGCACAGGGGCGTGGTGCTGCAGCTCGAGGCCGCGGAGGGGGGCCGCGGTG
GACTTCGGGTCGGAGGGACCGATATACCAGGGCTCTGGCGCGGGGTAGCCCCAGGACGCT
TTCTCGGGGAGGGGAGGGTACGGGCAGCCAGGTGGGAGTGCCAGAACTGGACGGCGTAAT
CCCTGCTGCCTGCGATGTGGGGTCGGACCCAGCGCACCCTGATGGGGCCTGGCGGCGGGG
GACCCCTGAACGGACATGGCAGCTACTCTGCCGTTGAGCCGATGCGGACGTGCCCAGTAC
ACGTGGGTGTTGGGGGAACTCCACAGCGGCAGGCGAGGGTTGGGTGTAGCAGGGGGCCCA
CGGGGGCGGCGGGTGGCAGGCGCGAGGGGGGCGCGGGGGAGCGATCCGATGGTCGAGAAC
GACCGGCTAGGAGGCGAGCCCCAGGTGGCACCGGGCGGGTGCGCCCGTGTCGGCGCGGGG
GGGTTGTGGCGACGAGGCCGCCCTTCAGGCGAGGTGGGAGTGGCCCAAGTTGTGGCGCCG
CGGTGGAGGGGGGGTGGTGAGGTGGGGGGCCAGTCGAGGTTGGTGAGGGCGGGACCAGCC
TCGTTTAGTGAAACGCCAACCCGGATAGTAATTCGTGCGGCTCGTGGGAGCGAGTTCTTC
CAATGATGCAGGGGGGGGGTGTGGGGCCCGCGCGTCGACCGGTCGCGTCACGACTGCTACG
GGGTCATTACCGCCGGGCTGTCGTAGGCCGGCGCTCGCGGGAGGTGGTTCGAATGCCGGG
GCGGGAGGCCCGGATCCGGCGAAGGGGCGGGGCTCGGGGGACGCGAGGCGGAGGGGCGCG
AGGCGGTAGCCGGTGCAAGCGGCGCGTGGCCCCATGAGGAAGAACTCTCTCGTGCCCCAA
CACCACACATTAGCGCCCCCATCTCGGCCGACTTCGAGCACGCTCAACGAGAACCACCAC
AGGCCTCCAACCCGACGTACCACCTGTCCTCACGAGGCGGGCCGACCCTCGGGTCCAAGC
GAGGGCTCCGCCGCGCACGCCCGGCGTGGTGC
>MAP 059_A_tb1127
CCCCTTAGGCGCCTCGAGCGGGCCCCCGCTTCGCCAGTTCACCCGTCCCAGTAAGCGCCC
GGGGCACCTCGCCTTCGAGGAGAACCCCCAGAACGCCCCACCGTCTCCCAGCTCGCACAT
GCGCACGCGTCGCACTCTCCAGCGCGCGTACCGTTCCAGCCACCGTCCCCATGCTCCCCG
CGGGGCCGTATGACGAGCCACGCCCCGGCCCCGCCGCCGCCGCTCGCCCACCGAGACAAG
CGCAATGCTATTGGGGCCTGACCACGGCTGGGCGGGGCGCTTCCCACGGAGGGCCCAGTC
TTTGGGAGGAGTTGCCCCTCCAGGAATCGGTAGCGCGGTGCGGCTCCCCGCAGCGGCCGC
AGGCGTCGGGGGGATGCTGGGGAGTGGGGGATGCCCCTTCCGGTCGCAGGCGGAACAGGT
TCGCGGTGGTGCACAGGGGCGTGGTGCTGCAGCTCGAGGCCGCGGAGGGGGGCCGCGGTG
GACTTCGGGTCGGAGGGACCGATATACCAGGGCTCTGGCGCGGGGTAGCCCCAGGACGCT
TTCTCGGGGAGGGGAGGGTACGGGCAGCCAGGTGGGAGTGCCAGAACTGGACGGCGTAAT
CCCTGCTGCCTGCGATGTGGGGTCGGACCCAGCGCACCCTGATGGGGCCTGGCGGCGGGG
GACCCCTGAACGGACATGGCAGCTACTCTGCCGTTGAGCCGATGCGGACGTGCCCAGTAC
ACGTGGGTGTTGGGGGAACTCCACAGCGGCAGGCGAGGGTTGGGTGTAGCAGGGGGCCCA
CGGGGGCGGCGGGTGGCAGGCGCGAGGGGGGCGCGGGGGAGCGATCCGATGGTCGAGAAC
GACCGGCTAGGAGGCGAGCCCCAGGTGGCACCGGGCGGGTGCGCCCGTGTCGGCGCGGGG
GGGTTGTGGCGACGAGGCCGCCCTTCAGGCGAGGTGGGAGTGGCCCAAGTTGTGGCGCCG
CGGTGGAGGGGGGGTGGTGAGGTGGGGGGCCAGTCGAGGTTGGTGAGGGCGGGACCAGCC
TCGTTTAGTGAAACGCCAACCCGGATAGTAATTCGTGCGGCTCGTGGGAGCGAGTTCTTC
CAATGATGCAGGGGGGGGGTGTGGGGCCCGCGCGTCGACCGGTCGCGTCACGACTGCTACG
GGGTCATTACCGCCGGGCTGTCGTAGGCCGGCGCTCGCGGGAGGTGGTTCGAATGCCGGG
GCGGGAGGCCCGGATCCGGCGAAGGGGCGGGGCTCGGGGGACGCGAGGCGGAGGGGCGCG
AGGCGGTAGCCGGTGCAAGCGGCGCGTGGCCCCATGAGGAAGAACTCTCTCGTGCCCCAA
CACCACACATTAGCGCCCCCATCTCGGCCGACTTCGAGCACGCTCAACGAGAACCACCAC
AGGCCTCCAACCCGACGTACCACCTGTCCTCACGAGGCGGGCCGACCCTCGGGTCCAAGC
GAGGGCTCCGCCGCGCACGCCCGGCGTGGTGC
>MAP 124_A_fb1148
CCCCTTAGGCGCCTCGAGCGGGCCCCCGCTTCGCCAGTTCAGCCGTCCCAGTAAGCGCCC
GGGGCACCTCGCCTTCGAGGAGAACCCCCAGAACGCCCCACCGTCTCCCAGCTCGCACAC
GCGCACGCGTCGCACTCTCCCGCGCGCGTACTGTTCCAGCCACCGTCCCCATGCTCCCCG
CGGGGCCGTATGACGAGCCACGCCCCGGCCCCGCCGCCGCCGCTCGCTCACCGAGACCAG
CGCAATGCTATTGGGGCCTGACCACGGCTGGGCGGGGCGCTTCCCACGGAGGGCCCAGTC
TTTGGGAGGAGTTGCCCCTCCAGGGATCGGCAGCGCGGTGCGGCTCCCCGCAGCGGCCGC
GGGCGTCGGGGGGATGCTAGGGAGTGGGGGATGCCCCTTCCGGTCGCAGGCGGAACAGGT
TCGCGGTGGTGCACAGGGGCGTGGTGCTGCAGCTCGAGGCCGCGGAGGGGGGCCGCGGTG
GACTTCGGGTCGGAGGGACCGATATACCAGGGCTCTGGCGCGGGGTAGCCCCAGGACGCT
TTCTCGGGGAGGGGAGGGTACGGGCAGCCAGGTGGGAGTGCAAGAACTGGACGGCCTAAT
CCCTGCTGCCTGCGATGTGGGGTCGGACCCAGCGCACCCTGATGGGGCCTGGCGGCGGGG
GACCCCTGAACGGACGTGGCAGCTACTCTGCCGTTGAGCCGATGCGGACGTGCCCAGTAC
ACGTGGGTGTTGGGGGAACTCCACAGCGGCAGGCGAGGGTTGGGTGTAGCAGGGGGCCCA
CGGGGGCGGCGGGTGGCAGGCGCGAGGGGGGCGCGGGGGAGCGATCCGATGGTCGAGAAC
GATCGGCTAGGAGGCGAGCCCCAGGTGGCGCCGGGCGGGTGCGCCCGTGTCGGCGCGGGG
GGGTTGTGGCGACGAGGCCGCCCTTCAGGCGAGGTGGGAGTGGCCCACGTTGTGGCGCCG
CGGTGGAGGGGGGGCGGTGAGGTGGGGGGCCAGTCGAGGTTGGTGAGGGCGGGACCAGCC
TCGCTTAGTGCAACGCCAACCCGGATAGTAATTCGTGCGGCTCGTGGGAGCGAGTTCTTC
CAATGATGCAGGGGGGGGGTGTGGGGCCCGCGCGTCGACCGGCCGCGTCACGACTGCTACG
GGGTCATTACCGCCGGGCTGTCGTAGGCCGGCGCTCGCGGGAGGTGGTTCGAATGCCGGG
GCGGGAGGCCCGGATCCGGCCAAGGGGCGGGGCTCGGGGGACGCGAGGCGGAGGGGCGCG
AGGCGGTAGCCGGTGCAAGCGGCGCGTGGCCCCATGAGGAAGAACTCTCTCGTGCCCCAA
CACCACACATTAGCGCCCCCATCTCGGCCGACTTCGAGCACGCTCAGCGAGAACCACCAC

AGGCCTCCAACCCGACGTACCACCCGTCCTCACGAGGCGGGCCGACCCTCGGGTCCAAGC
GAGGGCTCCGGCGCGCACGCCCGGCGTGGTGC
>MAP 029_A_fb1149
CCCCTTAGGCGCCTCGAGCGGGCCCCCGCTTCGCCAGTTCAGCCGTCCCGGTAAGCGCCC
GGGGCACCTCGCCTTCGAGGAGAACCCCCAGAACGCCCTACCGTCTCCCAGCTCGCACAC
GCGCACGCGTCGCACTCTCCCGCGCGCGTACTGTTCCAGCCACCGTCCCCATGCTCCCCG
CGGGGCCGTATGACGAGCCACGCCCCGGCCCCGCCGCCGCCGCTCGCTCACCGAGACCAG
CGCAATGCTATTGGGGCCTGACCACGGCTGGGCGGGGCGCTTCCCACGGAGGGCCCAGTC
TTTGGGAGGAGTTGCCCCTCCAGGGATCGACAGCGCGGTGCGGCTCCCCGCAGCGGCCGC
GGGCGTCGGGGGGATGCTAGGGAGTGGGGGATGCCCCTTCCGGTCGCAGGCGGAACAGGT
TCGCGGTGGTGCACAGGGGCGTGGTGCTGCAGCTCGAGGCCGCGGAGGGGGGCCGCGGTG
GACTTCGGGTCGGAGGGACCGATATACCAGGGCTCTGGCGCGGGGTAGCCCCAGGACGCT
TTCTCGGGGAGGGGAGGGTACGGGCAGCCAGGTGGGAGTGCAAGAACTGGACGGCCTAAT
CCCTGCTGCCTGCGATGTGGGGTCGGACCCAGCGCACCCTGATGGGGCCTGGCGGCGGGG
GACCCCTGAACGGACGTGGCAGCTACTCTGCCGTTGAGCCGATGCGGACGTGCCCAGTAC
ACGTGGGTGTTGGGGGAACTCCACAGCGGCAGGCGAGGGTTGGGTGTAGCAGGGGGCCCA
CGGGGGCGGCGGGTGGCAGGCGCGAGGGGGGCGCGGGGGAGCGATCCGATGGTCGAGAAC
GATCGGCTAGGAGGCGAGCCCCAGGTGGCGCCGGGCGGGTGCGCCCGTGTCGGCGCGGGG
GGGTTGTGGCGACGAGGCCGCCCTTCAGGCGAGGTGGGAGTGGCACACGTTGTGGCGCCG
CGGTGGAGGGGGGGCGGTGAGGTGGGGGGCCAGTCGAGGTTGGTGAGGGCGGGACCAGCC
TCGCTTAGTGCAACGCCAACCCGGATAGTAATTCGTGCGGCTCGTGGGAGCGAGTTCTTC
CAATGATGCAGGGGGGGGTGTGGGGCCCGCGCGTCGACCGGCCGCGTCACGACTGCTACG
GGGTCATTACCGCCGGGCTGTCGTAGGCCGGCGCTCGCGGGAGGTGGTTCGAATGCCGGG
GCGGGAGGCCCGGATCCGGCCAAGGGGCGGGGCTCGGGGGACGCGAGGCGGAGGGGCGCG
AGGCGGTAGCCGGTGCAAGCGGCGCGTGGCCCCATGAGGAAGAACTCTCTCGTGCCCCAA
CACCACACATTAGCGCCCCCATCTCGGCCGACTTCGAGCACGCTCAGCGAGAACCACCAC
AGGCCTCCAACCCGACGTACCACCCGTCCTCACGAGGCGGGCCGACCCTCGGGTCCAAGC
GAGGGCTCCGGCGCGCACGCCCGGCGTGGTGC
>MAP 179_A_tb1158
TCCCTCAGGCGCCTCGAGTGGGCCCCCGCTTCGCCAGTCCACCTGTCCCAGTAAGCGCCC
GCGACCCCTCGCCTTCGCATGGAACCCTTAGAACGCCTCACCGTCTCCCAGCTTGCACGC
GCGCACGCGTCGCACTTTCCAGCGCGCGGACTTCTCCAGCCGCCGTCCCCAGGCTCCCCG
CGGGGCCGCACGACAAGCTACGTCCCGGTCCCGCCGCCGCCGCCCGCCCACCGAGACAAC
TTCAACGCTATTGGGGCTTGACCACGGCTGGGCATGGCACTTCCCACGGAGGGCCCAGTC
TTCCTGAGGAGCTGCCCCTCAAGGGATCGGTAGTGCGGTGCGGTTCCCCGCAGCGGCCGT
GGGCGTCGGGGTGATCCTGGGGAGTGGGGGATGCCCCTTTCTGCCGCGGGTAGCGCGGGT
TCGCGGTGGTGCACAGGCGCGTGGTGCCGCGGTTCGAGGTGGCGGAGGGGGGTTGCGGTG
GACTTCGGGTCGGATGGACCGATAGACCAGGGCTTTGGCGCGGCGCGGCCGCAGGACACT
CTCTCGGGGAGGGGAGGGTACGGGCAGCCAGGTGGGAGTGCAGGCGCTGGACGGCGTACT
CCCTGGTGACTGGGATGTGGGGTAGGATCCAGCGCACTCTGGCGAGGCCTGGCGGCGGGG
GCCCCCCGGACGCACGCGGCGGCTACCTTGGCGTTGGGCCGATGCGGACGTGCCCAGTAC
ACGTGGGTGTTAGGGGAACTCCACAGCGGCAGGCGAGGGTTGGGCGTGGCCGAGGGCCCA
CGGGGGCGGCGGGGGGCGGTCGCGAGGGGGGCGCGGGAGAGCGATCCGATGATCGAGAAC
GATCGGCCAGAAGGCGAGCTCCAGGTGGCGCAGAGCGTGTGCGCCCGCGTCGGCGCGGGG
GGGTTGTGGCGACGCGGCCACCCTTCAGGCGGGGCGGGAGTGGCCCAAGTTGTGGCGCCG
CGGTGGAGGGGGCCCGGTGAGGTGGGGGGCCAGTCGGGGTTGGTGAGGGCGGAACCAGCC
ACGCTTGGCGCATGGCCAAGGCAGATAGTAATTCGTGCAGCTCGCGGGCGCAAGTCCTTC
CAATGATTCAGGGGGGGGTGGGGGGCCCGCGCGTCGGCCGGCCGCGGCACGAGTGCTGCG
GGGCCATTACCGCCGGGCCGCAGTAGGCCGGCGATCGCGGGAGGTGGTTCGTGTGCCGGG
GCGGGAAGGCCGGATCAGGCGAAGGGGCGGGGCCCGGGGCACGCGAGGCGGGGGGACGCG
AGGCGGTAGTCGGCGCAAGCGGCGCGTGGCCTCATGAGGTAGAACTCTCTCATGCCCCAA
CACCATACATTAGCGCCCCCGCTTCGGCCGGGCTTGAGTGCGCTCAGCGAGAACCACCAC
AGGCCTCCAACCCGACGTTCCACTCGTCCTCACGGGGCGGGCCGACCCCGGGGCCCAAGC
GAGGCCTCCGGCGAGCACGCCCGGTGTGGTGC
>MAP 007_A_fb1161
CCCCTTAGGCGCCTCGAGCGGGCCCCCGCTTCGCCAGTTCAGCCGTCCCAGTAAGCGCCC
GGGGCACCTCGCCTTCGAGGAGAACCCCCAGAACGCCCCACCGTCTCCCAGCTCGCACAC
GCGCACGCGTCGCACTCTCCCGCGCGCGTACTGTTCCAGCCACCGTCCCCATGCTCCCCG
CGGGGCCGTATGACGAGCCACGCCCCGGCCCCGCCGCCGCCGCTCGCTCACCGAGACCAG
CGCAATGCTATTGGGGCCTGACCACGGCTGGGCGGGGCGCTTCCCACGGAGGGCCCAGTC
TTTGGGAGGAGTTGCCCCTCCAGGGATCGGCAGCGCGGTGCGGCTCCCCGCAGCGGCCGC
GGGCGTCGGGGGGATGCTAGGGAGTGGGGGATGCCCCTTCCGGTCGCAGGCGGAACAGGT
TCGCGGTGGTGCACAGGGGCGTGGTGCTGCAGCTCGAGGCCGCGGAGGGGGGCCGCGGTG
GACTTCGGGTCGGAGGGACCGATATACCAGGGCTCTGGCGCGGGGTAGCCCCAGGACGCT
TTCTCGGGGAGGGGAGGGTACGGGCAGCCAGGTGGGAGTGCAAGAACTGGACGGCCTAAT
CCCTGCTGCCTGCGATGTGGGGTCGGACCCAGCGCACCCTGATGGGGCCTGGCGGCGGGG
GACCCCTGAACGGACGTGGCAGCTACTCTGCCGTTGAGCCGATGCGGACGTGCCCAGTAC
ACGTGGGTGTTGGGGGAACTCCACAGCGGCAGGCGAGGGTTGGGTGTAGCAGGGGGCCCA

CGGGGGCGGCGGGTGGCAGGCGCGAGGGGGGCGCGGGGGAGCGATCCGATGGTCGAGAAC
GATCGGCTAGGAGGCGAGCCCCAGGTGGCGCCGGGCGGGTGCGCCCGTGTCGGCGCGGGG
GGGTTGTGGCGACGAGGCCGCCCTTCAGGCGAGGTGGGAGTGGCCCACGTTGTGGCGCCG
CGGTGGAGGGGGGGCGGTGAGGTGGGGGGCCAGTCGAGGTTGGTGAGGGCGGGACCAGCC
TCGCTTAGTGCAACGCCAACCCGGATAGTAATTCGTGCGGCTCGTGGGAGCGAGTTCTTC
CAATGATGCAGGGGGGGGGTGTGGGGCCCGCGCGTCGACCGGCCGCGTCACGACTGCTACG
GGGTCATTACCGCCGGGCTGTCGTAGGCCGGCGCTCGCGGGAGGTGGTTCGAATGCCGGG
GCGGGAGGCCCGGATCCGGCCAAGGGGCGGGGCTCGGGGGACGCGAGGCGGAGGGGCGCG
AGGCGGTAGCCGGTGCAAGCGGCGCGTGGCCCCATGAGGAAGAACTCTCTCGTGCCCCAA
CACCACACATTAGCGCCCCCATCTCGGCCGACTTCGAGCACGCTCAGCGAGAACCACCAC
AGGCCTCCAACCCGACGTACCACCCGTCCTCACGAGGCGGGCCGACCCTCGGGTCCAAGC
GAGGGCTCCGGCGCGCACGCCCGGCGTGGTGC
>MAP 020_A_tb1167
CCCCTTAGGCGCCTCGAGCGGGCCCCCGCTTCGCCAGTTCAGCCGTCCCAGTAAGCGCCC
GGGGCACCTCGCCTTCGAGGAGAACCCCCAGAACGCCCCACCGTCTCCCAGCTCGCACAC
GCGCACGCGTCGCACTCTCCCGCGCGCGTACTGTTCCAGCCACCGTCCCCATGCTCCCCG
CGGGGCCGTATGACGAGCCACGCCCCGGCCCCGCCGCCGCCGCTCGCTCACCGAGACCAG
CGCAATGCTATTGGGGCCTGACCACGGCTGGGCGGGGCGCTTCCCACGGAGGGCCCAGTC
TTTGGGAGGAGTTGCCCCTCCAGGGATCGGCAGCGCGGTGCGGCTCCCCGCAGCGGCCGC
GGGCGTCGGGGGGATGCTAGGGAGTGGGGGATGCCCCTTCCGGTCGCAGGCGGAACAGGT
TCGCGGTGGTGCACAGGGGCGTGGTGCTGCAGCTCGAGGCCGCGGAGGGGGGGCCGCGGTG
GACTTCGGGTCGGAGGGACCGATATACCAGGGCTCTGGCGCGGGGTAGCCCCAGGACGCT
TTCTCGGGGAGGGGAGGGTACGGGCAGCCAGGTGGGAGTGCAAGAACTGGACGGCCTAAT
CCCTGCTGCCTGCGATGTGGGGTCGGACCCAGCGCACCCTGATGGGGCCTGGCGGCGGGG
GACCCCTGAACGGACGTGGCAGCTACTCTGCCGTTGAGCCGATGCGGACGTGCCCAGTAC
ACGTGGGTGTTGGGGGAACTCCACAGCGGCAGGCGAGGGTTGGGTGTAGCAGGGGGCCCA
CGGGGGCGGCGGGTGGCAGGCGCGAGGGGGGCGCGGGGGAGCGATCCGATGGTCGAGAAC
GATCGGCTAGGAGGCGAGCCCCAGGTGGCGCCGGGCGGGTGCGCCCGTGTCGGCGCGGGG
GGGTTGTGGCGACGAGGCCGCCCTTCAGGCGAGGTGGGAGTGGCCCACGTTGTGGCGCCG
CGGTGGAGGGGGGGCGGTGAGGTGGGGGGCCAGTCGAGGTTGGTGAGGGCGGGACCAGCC
TCGCTTAGTGCAACGCCAACCCGGATAGTAATTCGTGCGGCTCGTGGGAGCGAGTTCTTC
CAATGATGCAGGGGGGGGGTGTGGGGCCCGCGCGTCGACCGGCCGCGTCACGACTGCTACG
GGGTCATTACCGCCGGGCTGTCGTAGGCCGGCGCTCGCGGGAGGTGGTTCGAATGCCGGG
GCGGGAGGCCCGGATCCGGCCAAGGGGCGGGGCTCGGGGGACGCGAGGCGGAGGGGCGCG
AGGCGGTAGCCGGTGCAAGCGGCGCGTGGCCCCATGAGGAAGAACTCTCTCGTGCCCCAA
CACCACACATTAGCGCCCCCATCTCGGCCGACTTCGAGCACGCTCAGCGAGAACCACCAC
AGGCCTCCAACCCGACGTACCACCCGTCCTCACGAGGCGGGCCGACCCTCGGGTCCAAGC
GAGGGCTCCGGCGCGCACGCCCGGCGTGGTGC
>MAP 087_A_fb1171
CCCCTTAGGCGCCTCGAGCGGGCCCCCGCTTCGCCAGTTCAGCCGTCCCAGTAAGCGCCC
GGGGCACCTCGCCTTCGAGGAGAACCCCCAGAACGCCCCACCGTCTCCCAGCTCGCACAC
GCGCACGCGTCGCACTCTCCCGCGCGCGTACTGTTCCAGCCACCGTCCCCATGCTCCCCG
CGGGGCCGTATGACGAGCCACGCCCCGGCCCCGCCGCCGCCGCTCGCTCACCGAGACCAG
CGCAATGCTATTGGGGCCTGACCACGGCTGGGCGGGGCGCTTCCCACGGAGGGCCCAGTC
TTTGGGAGGAGTTGCCCCTCCAGGGATCGGCAGCGCGGTGCGGCTCCCCGCAGCGGCCGC
GGGCGTCGGGGGGATGCTAGGGAGTGGGGGATGCCCCTTCCGGTCGCAGGCGGAACAGGT
TCACGGTGGTGCACAGGGGCGTGGTGCTGCAGCTCGAGGCCGCGGAGGGGGGGCCGCGGTG
GACTTCGGGTCGGAGGGACCGATATACCAGGGCTCTGGCGCGGGGTAGCCCCAGGACGCT
TTCTCGGGGAGGGGAGGGTACGGGCAGCCAGGTGGGAGTGCAAGAACTGGACGGCCTAAT
CCCTGCTGCCTGCGATGTGGGGTCGGACCCAGCGCACCCTGATGGGGCCTGGCGGCGGGG
GACCCCTGAACGGACGTGGCAGCTACTCTGCCGTTGAGCCGATGCGGACGTGCCCAGTAC
ACGTGGGTGTTGGGGGAACTCCACAGCGGCAGGCGAGGGTTGGGTGTAGCAGGGGGCCCA
CGGGGGCGGCGGGTGGCAGGCGCGAGGGGGGCGCGGGGGAGCGATCCGATGGTCGAGAAC
GATCGGCTAGGAGGCGAGCCCCAGGTGGCGCCGGGCGGGTGCGCCCGTGTCGGCGCGGGG
GGGTTGTGGCGACGAGGCCGCCCTTCAGGCGAGGTGGGAGTGGCCCACGTTGTGGCGCCG
CGGTGGAGGGGGGGCGGTGAGGTGGGGGGCCAGTCGAGGTTGGTGAGGGCGGGACCAGCC
TCGCTTAGTGCAACGCCAACCCGGATAGTAATTCGTGCGGCTCGTGGGAGCGAGTTCTTC
CAATGATGCAGGGGGGGGGTGTGGGGCCCGCGCGTCGACCGGCCGCGTCACGACTGCTACG
GGGTCATTACCGCCGGGCTGTCGTAGGCCGGCGCTCGCGGGAGGTGGTTCGAATGCCGGG
GCGGGAGGCCCGGATCCGGCCAAGGGGCGGGGCTCGGGGGACGCGAGGCGGAGGGGCGCG
AGGCGGTAGCCGGTGCAAGCGGCGCGTGGCCCCATGAGGAAGAACTCTCTGGTGCCCCAA
TACCACACATTAGCGCCCCCATCTCGGCCGACTTCGAGCACGCTCAGCGAGAACCACCAC
AGGCCTCCAACCCGACGTACCACCCGTCCTCACGAGGCGGGCCGACCCTCGGGTCCAAGC
GAGGGCTCCGGCGCGCACGCCCGGCGTGGTGC
>MAP 058_A_fb1179
CCCCTTAGGCGCCTCGAGCGGGCCCCCGCTTCGCCAGTTCAGCCGTCCCAGTAAGCGCCC
GGGGCACCTCGCCTTCGAGGAGAACCCCCAGAACGCCCCACCGTCTCCCAGCTCGCACAC
GCGCACGCGTCGCACTCTCCCGCGCGCGTACTGTTCCAGCCACCGTCCCCATGCTCCCCG

CGGGGCCGTATGACGAGCCACGCCCCGGCCCCGCCGCCGCCGCTCGCTCACCGAGACCAG
CGCAATGCTATTGGGGCCTGACCACGGCTGGGCGGGGCGCTTCCCACGGAGGGCCCAGTT
TTTGGGAGGAGTTGCCCCTCCAGGGATCGGCCGCGCGGTGCGGCTCCCCGCAGCGGCCGC
GGGCGTCGGGGGGATGCTAGGGAGTGGGGGATGCCCCTTCCGGTCGCAGGCGGAACAGGT
TCGCGGTGGTGCACAGGGGCGTGGTGCTGCAGCTCGAGGCCGCGGAGGGGGGCCGCGGTG
GACTTCGGGTCGGAGGGACCGATATACCAGGGCTCTGGCGCGGGGTAGCCCCAGGACGCT
TTCTCGGGGAGGGGAGGGTACGGGCAGCCAGGTGGGAGTGCAAGAACTGGACGGCCTAAT
CCCTGCTGCCTGCGATGTGGGGTCGGACCCAGCGCACCCTGATGGGGCCTGGCGGCGGGG
GACCCCTGAACGGACGTGGCAGCTACTCTGCCGTTGAGCCGATGCGGACGTGCCCAGTAC
ACGTGGGTGTTGGGGGAACTCCACAGCGGCAGGCGAGGGTTGGGTGTAGCAGGGGGCCCA
CGGGGGCGGCGGGTGGCAGGCGCGAGGGGGGCGCGGGGGAGCGATCCGATGGTCGAGAAC
GATCGGCTAGGAGGCGAGCCCCAGGTGGCGCCGGGCGGGTGCGCCCGTGTCGGCGCGGGG
GGGTTGTGGCGACGAGGCCGCCCTTCAGGCGAGGTGGGAGTGGCCCACGTTGTGGCGCCG
CGGTGGAGGGGGGGGCGGTGAGGTGGGGGGCCAGTCGAGGTTGGTGAGGGCGGGACCAGCC
TCGCTTAGTGCAACGCCAACCCGGATAGTAATTCGTGCGGCTCGTGGGAGCGAGTTCTTC
CAATGATGCAGGGGGGGGGTGTGGGGCCCGCGCGTCGACCGGCCGCGTCACGACTGCTACG
GGGTCATTACCGCCGGGCTGTCGTAGGCCGGCGCTCGCGGGAGGTGGTTCGAATGCCGGG
GCGGGAGGCCCGGATCCGGCCAAGGGGCGGGGCTCGGGGGACGCGAGGCGGAGGGGCGCG
AGGCGGTAGCCGGTGCAAGCGGCGCGTGGCCCCATGAGGAAGAACTCTCTCGTGCCCCAA
CACCACACATTAGCGCCCCCATCTCGGCCGACTTCGAGCACGCTCAGCGAGAACCACCAC
AGGCCTCCAACCCGACGTACCACCCGTCCTCACGAGGCGGGCCGACCCTCGGGTCCAAGC
GAGGGCTCCGGCGCGCACGCCCGGCGTGGTGC
>MAP 150_A_fb1179
CCCCTTAGGCGCCTCGAGCGGGCCCCCGCTTCGCCAGTTCAGCCGTCCCAGTAAGCGCCC
GGGGCACCTCGCCTTCGAGGAGAACCCCCAGAACGCCCCACCGTCTCCCAGCTCGCACAC
GCGCACGCGTCGCACTCTCCCGCGCGCGTACTGTTCCAGCCACCGTCCCCATGCTCCCCG
CGGGGCCGTATGACGAGCCACGCCCCGGCCCCGCCGCCGCCGCTCGCTCACCGAGACCAG
CGCAATGCTATTGGGGCCTGACCACGGCTGGGCGGGGCGCTTCCCACGGAGGGCCCAGTC
TTTGGGAGGAGTTGCCCCTCCAGGGATCGGCCGCGCGGTGCGGCTCCCCGCAGCGGCCGC
GGGCGTCGGGGGGATGCTAGGGAGTGGGGGATGCCCCTTCCGGTCGCAGGCGGAACAGGT
TCGCGGTGGTGCACAGGGGCGTGGTGCTGCAGCTCGAGGCCGCGGAGGGGGGCCGCGGTG
GACTTCGGGTCGGAGGGACCGATATACCAGGGCTCTGGCGCGGGGTAGCCCCAGGACGCT
TTCTCGGGGAGGGGAGGGTACGGGCAGCCAGGTGGGAGTGCAAGAACTGGACGGCCTAAT
CCCTGCTGCCTGCGATGTGGGGTCGGACCCAGCGCACCCTGATGGGGCCTGGCGGCGGGG
GACCCCTGAACGGACGTGGCAGCTACTCTGCCGTTGAGCCGATGCGGACGTGCCCAGTAC
ACGTGGGTGTTGGGGGAACTCCACAGCGGCAGGCGAGGGTTGGGTGTAGCAGGGGGCCCA
CGGGGGCGGCGGGTGGCAGGCGCGAGGGGGGCGCGGGGGAGCGATCCGATGGTCGAGAAC
GATCGGCTAGGAGGCGAGCCCCAGGTGGCGCCGGGCGGGTGCGCCCGTGTCGGCGCGGGG
GGGTTGTGGCGACGAGGCCGCCCTTCAGGCGAGGTGGGAGTGGCCCACGTTGTGGCGCCG
CGGTGGAGGGGGGGGCGGTGAGGTGGGGGGCCAGTCGAGGTTGGTGAGGGCGGGACCAGCC
TCGCTTAGTGCAACGCCAACCCGGATAGTAATTCGTGCGGCTCGTGGGAGCGAGTTCTTC
CAATGATGCAGGGGGGGGGTGTGGGGCCCGCGCGTCGACCGGCCGCGTCACGACTGCTACG
GGGTCATTACCGCCGGGCTGTCGTAGGCCGGCGCTCGCGGGAGGTGGTTCGAATGCCGGG
GCGGGAGGCCCGGATCCGGCCAAGGGGCGGGGCTCGGGGGACGCGAGGCGGAGGGGCGCG
AGGCGGTAGCCGGTGCAAGCGGCGCGTGGCTCCATGAGGAAGAACTCTCTCGTGCCCCAA
CACCACACATTAGCGCCCCCATCTCGGCCGACTTCGAGCACGCTCAGCGAGAACCACCAC
AGGCCTCCAACCCGACGTACCACCCGTCCTCACGAGGCGGGCCGACCCTCGGGTCCAAGC
GAGGGCTCCGGCGCGCACGCCCGGCGTGGTGC
>MAP 152_A_fb1179
CCCCTTAGGCGCCTCGAGCGGGCCCCCGCTTCGCCAGTTCAGCCGTCCCAGTAAGCGCCC
GGGGCACCTCGCCTTCGAGGAGAACCCCCAGAACGCCCCACCGTCTCCCAGCTCGCACAC
GCGCACGCGTCGCACTCTCCCGCGCGCGTACTGTTCCAGCCACCGTCCCCATGCTCCCCG
CGGGGCCGTATGACGAGCCACGCCCCGGCCCCGCCGCCGCCGCTCGCTCACCGAGACCAG
CGCAATGCTATTGGGGCCTGACCACGGCTGGGCGGGGCGCTTCCCACGGAGGGCCCAGTC
TTTGGGAGGAGTTGCCCCTCCAGGGATCGGCAGCGCGGTGCGGCTCCCCGCAGCGGCCGC
GGGCGTCGGGGGGATGCTAGGGAGTGGGGGATGCCCCTTCCGGTCGCAGGCGGAACAGGT
TCGCGGTGGTGCACAGGGGCGTGGTGCTGCAGCTCGAGGCCGCGGAGGGGGGCCGCGGTG
GACTTCGGGTCGGAGGGACCGATATACCAGGGCTCTGGCGCGGGGTAGCCCCAGGACGCT
TTCTCGGGGAGGGGAGGGTACGGGCAGCCAGGTGGGAGTGCAAGAACTGGACGGCCTAAT
CCCTGCTGCCTGCGATGTGGGGTCGGACCCAGCGCACCCTGATGGGGCCTGGCGGCGGGG
GACCCCTGAACGGACGTGGCAGCTACTCTGCCGTTGAGCCGATGCGGACGTGCCCAGTAC
ACGTGGGTGTTGGGGGAACTCCACAGCGGCAGGCGAGGGTTGGGTGTAGCAGGGGGCCCA
CGGGGGCGGCGGGTGGCAGGCGCGAGGGGGGCGCGGGGGAGCGATCCGATGGTCGAGAAC
GATCGGCTAGGAGGCGAGCCCCAGGTGGCGCCGGGCGGGTGCGCCCGTGTCGGCGCGGGG
GGGTTGTGGCGACGAGGCCGCCCTTCAGGCGAGGTGGGAGTGGCCCACGTTGTGGCGCCG
CGGTGGAGGGGGGGGCGGTGAGGTGGGGGGCCAGTCGAGGTTGGTGAGGGCGGGACCAGCC
TCGCTTAGTGCAACGCCAACCCGGATAGTAATTCGTGCGGCTCGTGGGAGCGAGTTCTTC
CAATGATGCAGGGGGGGGGTGTGGGGCCCGCGCGTCGACCGGCCGCGTCACGACTGCTACG

GGGTCATTACCGCCGGGCTGTCGTAGGCCGGCGCTCGCGGGAGGTGGTTCGAATGCCGGG
GCGGGAGGCCCGGATCCGGCCAAGGGGCGGGGCTCGGGGGACGCGAGGCGGAGGGGCGCG
AGGCGGTAGCCGGTGCAAGCGGCGCGTGGCCCCATGAGGAAGAACTCTCTCGTGCCCCAA
CACCACACATTAGCGCCCCCATCTCGGCCGACTTCGAGCACGCTCAGCGAGAACCACCAC
AGGCCTCCAACCCGACGTACCACCCGTCCTCACGAGGCGGGCCGACCCTCGGGTCCAAGC
GAGGGCTCCGGCGCGCACGCCCGGCGTGGTGC
>MAP 146_A_tb1179
CCCCTTAGGCGCCTCGAGCGGGCCCCCGCTTCGCCAGTTCAGCCGTCCCAGTAAGCGCCC
GGGGCACCTCGCCTTCGAGGAGAACCCCCAGAACGCCCCACCGTCTCCCAGCTCGCACAC
GCGCACGCGTCGCACTCTCCCGCGCGCGTACTGTTCCAGCCACCGTCCCCATGCTCCCCG
CGGGGCCGTATGACGAGCCACGCCCCGGCCCCGCCGCCGCCGCTCGCTCACCGAGACCAG
CGCAATGCTATTGGGGCCTGACCACGGCTGGGCGGGGCGCTTCCCACGGAGGGCCCAGTC
TTTGGGAGGAGTTGCCCCTCCAGGGATCGGCCGCGCGGTGCGGCTCCCCGCAGCGGCCGC
GGGCGTCGGGGGGATGCTAGGGAGTGGGGGATGCCCCTTCCGGTCGCAGGCGGAACAGGT
TCGCGGTGGTGCACAGGGGCGTGGTGCTGCAGCTCGAGGCCGCGGAGGGGGGCCGCGGTG
GACTTCGGGTCGGAGGGACCGATATACCAGGGCTCTGGCGCGGGGTAGCCCCAGGACGCT
TTCTCGGGGAGGGGAGGGTACGGGCAGCCAGGTGGGAGTGCAAGAACTGGACGGCCTAAT
CCCTGCTGCCTGCGATGTGGGGTCGGACCCAGCGCACCCTGATGGGGCCTGGCGGCGGGG
GACCCCTGAACGGACGTGGCAGCTACTCTGCCGTTGAGCCGATGCGGACGTGCCCAGTAC
ACGTGGGTGTTGGGGGAACTCCACAGCGGCAGGCGAGGGTTGGGTGTAGCAGGGGGCCCA
CGGGGGCGGCGGGTGGCAGGCGCGAGGGGGGCGCGGGGGAGCGATCCGATGGTCGAGAAC
GATCGGCTAGGAGGCGAGCCCCAGGTGGCGCCGGGCGGGTGCGCCCGTGTCGGCGCGGGG
GGGTTGTGGCGACGAGGCCGCCCTTCAGGCGAGGTGGGAGTGGCCCACGTTGTGGCGCCG
CGGTGGAGGGGGGGCGGTGAGGTGGGGGGCCAGTCGAGGTTGGTGAGGGCGGGACCAGCC
TCGCTTAGTGCAACGCCAACCCGGATAGTAATTCGTGCGGCTCGTGGGAGCGAGTTCTTC
CAATGATGCAGGGGGGGGTGTGGGGCCCGCGCGTCGACCGGCCGCGTCACGACTGCTACG
GGGTCATTACCGCCGGGCTGTCGTAGGCCGGCGCTCGCGGGAGGTGGTTCGAATGCCGGG
GCGGGAGGCCCGGATCCGGCCAAGGGGCGGGGCTCGGGGGACGCGAGGCGGAGGGGCGCG
AGGCGGTAGCCGGTGCAAGCGGCGCGTGGCCCCATGAGGAAGAACTCTCTCGTGCCCCAA
CACCACACATTAGCGCCCCCATCTCGGCCGACTTCGAGCACGCTCAGCGAGAACCACCAC
AGGCCTCCAACCCGACGTACCACCCGTCCTCACGAGGCGGGCCGACCCTCGGGTCCAAGC
GAGGGCTCCGGCGCGCACGCCCGGCGTGGTGC
>MAP 001_A_fb1180
CCCCTTAGGCGCCTCGAGCGGGCCCCCGCTTCGCCAGTTCAGCCGTCCCAGTAAGCGCCC
GGGGCACCTCGCCTTCGAGGAGAACCCCCAGAACGCCCCACCGTCTCCCAGCTCGCACAC
GCGCACGCGTCGCACTCTCCCGCGCGCGTACTGTTCCAGCCACCGTCCCCATGCTCCCCG
CGGGGCCGTATGACGAGCCACGCCCCGGCCCCGCCGCCGCCGCTCGCTCACCGAGACCAG
CGCAATGCTATTGGGGCCTGACCACGGCTGGGCGGGGCGCTTCCCACGGAGGGCCCAGTC
TTTGGGAGGAGTTGCCCCTCCAGGGATCGGCAGCGCGGTGCGGCTCCCCGCAGCGGCCGC
GGGCGTCGGGGGGATGCTAGGGAGTGGGGGATGCCCCTTCCGGTCGCAGGCGGAACAGGC
TCGCGGTGGTGCACAGGGGCGTGGTGCTGCAGCTCGAGGCCGCGGAGGGGGGCCGCGGTG
GACTTTGGGTCGGAGGGACCGATATACCAGGGCTCTGGCGCGGGGTAGCCCCAGGACGCT
TTCTCGGGGAGGGGAGGGTACGGGCAGCCAGGTGGGAGTGCAAGAACTGGACGGCCTAAT
CCCTGCTGCCTGCGATGTGGGGTCGGACCCAGCGCACCCTGATGGGGCCTGGCGGCGGGG
GACCCCTGAACGGACGTGGCAGCTACTCTGCCGTTGAGCCGATGCGGACGTGCCCAGTAC
ACGTGGGTGTTGGGGGAACTCCACAGCGGCAGGCGAGGGTTGGGTGTAGCAGGGGGCCCA
CGGGGGCGGCGGGTGGCAGGCGCGAGGGGGGCGCGGGGGAGCGATCCGATGGTCGAGAAC
GATCGGCTAGGAGGCGAGCCCCAGGTGGCGCCGGGCGGGTGCGCCCGTGTCGGCGCGGGG
GGGTTGTGGCGACGAGACCGCCCTTCAGGCGAGGTGGGAGTGGCCCACGTTGTGGCGCCG
CGGTGGAGGGGGGGCGGTGAGGTGGGGGGCCAGTCGAGGTTGGTGAGGGCGGGACCAGCC
TCGCTTAGTGCAACGCCAACCCGGATAGTAATTCGTGCGGCTCGTGGGAGCGAGTTCTTC
CAATGATGCAGGGGGGGGTGTGGGGCCCGCGCGTCGACCGGCCGCGTCACGACTGCTACG
GGGTCATTACCGCCGGGCTGTCGTAGGCCGGCGCTCGCGGGAGGTGGTTCGAATGCCGGG
GCGGGAGGCCCGGATCCGGCCAAGGGGCGGGGCTCGGGGGACGCGAGGCGGAGGGGCGCG
AGGCGGTAGCCGGTGCAAGCGGCGCGTGGCCCCATGAGGAAGAACTCTCTCGTGCCCCAA
CACCACACATTAGCGCCCCCATCTCGGCCGACTTCGAGCACGCTCAGCGAGAACCACCAC
AGGCCTCCAACCCGACGTACCACCCGTCCTCACGAGGCGGGCCGACCCTCGGGTCCAAGC
GAGGGCTCCGGCGCGCACGCCCGGCGTGGTGC
>MAP 033_A_fb1180
CCCCTTAGGCGCCTCGAGCGGGCCCCCGCTTCGCCAGTTCAGCCGTCCCAGTAAGCGCCC
GGGGCACCTCGCCTTCGAGGAGAACCCCCAGAACGCCCCACCGTCTCCCAGCTCGCACAC
GCGCACGCGTCGCACTCTCCCGCGCGCGTACTGTTCCAGCCACCGTCCCCATGCTCCCCG
CGGGGCCGTATGACGAGCCACGCCCCGGCCCCGCCGCCGCCGCTCGCTCACCGAGACCAG
CGCAATGCTATTGGGGCCTGACCACGGCTGGGCGGGGCGCTTCCCACGGAGGGCCCAGTC
TTTGGGAGGAGTTGCCCCTCCAGGGATCGGCAGCGCGGTGCGGCTCCCCGCAGCGGCCGC
GGGCGTCGGGGGGATGCTAGGGAGTGGGGGATGCCCCTTCCGGTCGCAGGCGGAACAGGT
TCGCGGTGGTGCACAGGGGCGTGGTGCTGCAGCTCGAGGCCGCGGAGGGGGGCCGCGGTG
GACTTCGGGTCGGAGGGACCGATATACCAGGGCTCTGGCGCGGGGTAGCCCCAGGACGCT

TTCTCGGGGAGGGGAGGGTACGGGCAGCCAGGTGGGAGTGCAAGAACTGGACGGCCTAAT
CCCTGCTGCCTGCGATGTGGGGTCGGACCCAGCGCACCCTGATGGGGCCTGGCGGCGGGG
GACCCCTGAACGGACGTGGCAGCTACTCTGCCGTTGAGCCGATGCGGACGTGCCCAGTAC
ACGTGGGTGTTGGGGGAACTCCACAGCGGCAGGCGAGGGTTGGGTGTAGCAGGGGGCCCA
CGGGGGCGGCGGGTGGCAGGCGCGAGGGGGGCGCGGGGGAGCGATCCGATGGTCGAGAAC
GATCGGCTAGGAGGCGAGCCCCAGGTGGCGCCGGGCGGGTGCGCCCGTGTCGGCGCGGGG
GGGTTGTGGCGACGAGGCCGCCCTTCAGGCGAGGTGGGAGTGGCCCACGTTGTGGCGCCG
CGGTGGAGGGGGGGCGGTGAGGTGGGGGGCCAGTCGAGGTTGGTGAGGGCGGGACCAGCC
TCGCTTAGTGCAACGCCAACCCGGATAGTAATTCGTGCGGCTCGTGGGAGCGAGTTCTTC
CAATGATGCAGGGGGGGGTGTGGGGCCCGCGCGTCGACCGGCCGCGTCACGACTGCTACG
GGGTCATTACCGCCGGGCTGTCGTAGGCCGGCGCTCGCGGGAGGTGGTTCGAATGCCGGG
GCGGGAGGCCCGGATCCGGCCAAGGGGCGGGGCTCGGGGGACGCGAGGCGGAGGGGCGCG
AGGCGGTAGCCGGTGCAAGCGGCGCGTGGCCCCATGAGGAAGAACTCTCTCGTGCCCCAA
CACCACACATTAGCGCCCCCATCTCGGCCGACTTCGAGCACGCTCAGCGAGAACCACCAC
AGGCCTCCAACCCGACGTACCACCCGTCCTCACGAGGCGGGCCGACCCTCGGGTCCAAGC
GAGGGCTCCGGCGCGCACGCCCGGCGTGGTGC
>MAP 086_A_fb1181
CCCCTTAGGCGCCTCGAGCGGGCTCCCGCTTCGCCAGTTCAGCCGTCCCAGTAAGCGCCC
GGGGCACCTCGCCTTCGAGGAGAACCCCCAGAACGCCCCACCGTCTCCCAGCTCGCACAC
GCGCACGCGTCGCACTCTCCCGCGCGCGTACTGTTCCAGCCACCGTCCCCATGCTCCCCG
CGGGGCCGTATGACGAGCCACGCCCCGGCCCCGCCGCCGCCGCTCGCTCACCGAGACCAG
CGCAATGCTATTGGGGCCTGACCACGGCTGGGCGGGGCGCTTCCCACGGAGGGCCCAGTC
TTTGGGAGGAGTTGCCCCTCCAGGGATCGGCAGCGCGGTGCGGCTCCCCGCAGCGGCCGC
GGGCGTCGGGGGGATGCTAGGGAGTGGGGGATGCCCCTTCCGGTCGCAGGCGGAACAGGT
TCGCGGTGGTGCACAGGGGCGTGGTGCTGCAGCTCGAGGCCGCGGAGGGGGGCCGCGGTG
GACTTCGGGTCGGAGGGACCGATATACCAGGGCTCTGGCGCGGGGTAGCCCCAGGACGCT
TTCTCGGGGAGGGGAGGGTACGGGCAGCCAGGTGGGAGTGCAAGAACTGGACGGCCTAAT
CCCTGCTGCCTGCGATGTGGGGTCGGACCCAGCGCACCCTGATGGGGCCTGGCGGCGGGG
GACCCCTGAACGGACGTGGCAGCTACTCTGCCGTTGAGCCGATGCGGACGTGCCCAGTAC
ACGTGGGTGTTGGGGGAACTCCACAGCGGCAGGCGAGGGTTGGGTGTAGCAGGGGGCCCA
CGGGGGCGGCGGGTGGCAGGCGCGAGGGGGGCGCGGGGGAGCGATCCGATGGTCGAGAAC
GATCGGCTAGGAGGCGAGCCCCAGGTGGCGCCGGGCGGGTGCGCCCGTGTCGGCGCGGGG
GGGTTGTGGCGACGAGGCCGCCCTTCAGGCGAGGTGGGAGTGGCCCACGTTGTGGCGCCG
CGGTGGAGGGGGGGCGGTGAGGTGGGGGGCCAGTCGAGGTTGGTGAGGGCGGGACCAGCC
TCGCTTAGTGCAACGCCAACCCGGATAGTAATTCGTGCGGCTCGTGGGAGCGAGTTCTTC
CAATGATGCAGGGGGGGGTGTGGGGCCCGCGCGTCGACCGGCCGCGTCACGACTGCTACG
GGGTCATTACCGCCGGGCTGTCGTAGGCCGGCGCTCGCGGGAGGTGGTTCGAATGCCGGG
GCGGGAGGCCCGGATCCGGCCAAGGGGCGGGGCTCGGGGGACGCGAGGCGGAGGGGCGCG
AGGCGGTAGCCGGTGCAAGCGGCGCGTGGCCCCATGAGGAAGAACTCTCTCGTGCCCCAA
CACCACACATTAGCGCCCCCATCTCGGCCGACTTCGAGCACGCTCAGCGAGAACCACCAC
AGGCCTCCAACCCGACGTACCACCCGTCCTCACGAGGCGGGCCGACCCTCGGGTCCAAGC
GAGGGCTCCGGCGCGCACGCCCGGCGTGGTGC
>MAP 151_A_fb1181
CCCCTTAGGCGCCTCGAGCGGGCCCCCGCTTCGCCAGTTCAGCCGTCCCAGTAAGCGCCC
GGGGCACCTCGCCTTCGAGGAGAACCCCCAGAACGCCCCACCGTCTCCCAGCTCGCACAC
GCGCACGCGTCGCACTCTCCCGCGCGCGTACTGTTCCAGCCACCGTCCCCATGCTCCCCG
CGGGGCCGTATGACGAGCCACGCCCCGGCCCCGCCGCCGCCGCTCGCTCACCGAGACCAG
CGCAATGCTATTGGGGCCTGACCACGGCTGGGCGGGGCGCTTCCCACGGAGGGCCCAGTC
TTTGGGAGGAGTTGCCCCTCCAGGGATCGGCAGCGCGGTGCGGCTCCCCGCAGCGGCCGC
GGGCGTCGGGGGGATGCTAGGGAGTGGGGGATGCCCCTTCCGGTCGCAGGCGGAACAGGT
TCGCGGTGGTGCACAGGGGCGTGGTGCTGCAGCTCGAGGCCGCGGAGGGGGGCCGCGGTG
GACTTCGGGTCGGAGGGACCGATATACCAGGGCTCTGGCGCGGGGTAGCCCCAGGACGCT
TTCTCGGGGAGGGGAGGGTACGGGCAGCCAGGTGGGAGTGCAAGAACTGGACGGCCTAAT
CCCTGCTGCCTGCGATGTGGGGTCGGACCCAGCGCACCCTGATGGGGCCTGGCGGCGGGG
GACCCCTGAACGGACGTGGCAGCTACTCTGCCGTTGAGCCGATGCGGACGTGCCCAGTAC
ACGTGGGTGTTGGGGGAACTCCACAGCGGCAGGCGAGGGTTGGGTGTAGCAGGGGGCCCA
CGGGGGCGGCGGGTGGCAGGCGCGAGGGGGGCGCGGGGGAGCGATCCGATGGTCGAGAAC
GATCGGCTAGGAGGCGAGCCCCAGGTGGCGCCGGGCGGGTGCGCCCGTGTCGGCGCGGGG
GGGTTGTGGCGACGAGGCCGCCCTTCAGGCGAGGTGGGAGTGGCCCACGTTGTGGCGCCG
CGGTGGAGGGGGGGCGGTGAGGTGGGGGGCCAGTCGAGGTTGGTGAGGGCGGGACCAGCC
TCGCTTAGTGCAACGCCAACCCGGATAGTAATTCGTGCGGCTCGTGGGAGCGAGTTCTTC
CAATGATGCAGGGGGGGGTGTGGGGCCCGCGCGTCGACCGGCCGCGTCACGACTGCTACG
GGGTCATTACCGCCGGGCTGTCGTAGGCCGGCGCTCGCGGGAGGTGGTTCGAATGCCGGG
GCGGGAGGCCCGGATCCGGCCAAGGGGCGGGGCTCGGGGGACGCGAGGCGGAGGGGCGCG
AGGCGGTAGCCGGTGCAAGCGGCGCGTGGCCCCATGAGGAAGAACTCTCTCGTGCCCCAA
CACCACACATTAGCGCCCCCATCTCGGCCGACTTCGAGCACGCTCAGCGAGAACCACCAC
AGGCCTCCAACCCGACGTACCACCCGTCCTCACGAGGCGGGCCGACCCTCGGGTCCAAGC
GAGGGCTCCGGCGCGCACGCCCGGCGTGGTGC

>MAP 018_A_fb1182
CCCCTTAGGCGCCTCGAGCGGGCCCCCGCTTCGCCAGTTCAGCCGTCCCAGTAAGCGTCC
GGGGCACCTCGCCTTCGAGGAGAACCCCCAGAACGCCCCACCGTCTCCCAGCTCGCACAC
GCGCACGCGTCGCACTCTCCCGCGCGCGTACTGTTCCAGCCACCGTCCCCATGCTCCCCG
CGGGGCCGTATGACGAGCCACGCCCCGGCCCCGCCGCCGCCGCTCGCTCACCGAGACCAG
CGCAATGCTATTGGGGCCTGACCACGGCTGGGCGGGGCGCTTCCCACGGAGGGCCCAGTC
TTTGGGAGGAGTTGCCCCTCCAGGGATCGGCAGCGCGGTGCGGCTCCCCGCAGCGGCCGC
GGGCGTCGGGGGGATGCTAGGGAGTGGGGGATGCCCCTTCCGGTCGCAGGCGGAACAGGT
TCGCGGTGGTGCACAGGGGCGTGGTGCTGCAGCTCGAGGCCGCGGAGGGGGGCCGCGGTG
GACTTCGGGTCGGAGGGACCGATATACCAGGGCTCTGGCGCGGGGTAGCCCCAGGACGCT
TTCTCGGGGAGGGGAGGGTACGGGCAGCCAGGTGGGAGTGCAAGAACTGGACGGCCTAAT
CCCTGCTGCCTGCGATGTGGGGTCGGACCTAGCGCACCCTGATGGGGCCTGGCGGCGGGG
GACCCCTGAACGGACGTGGCAGCTACTCTGCCGTTGAGCCGATGCGGACGTGCCCAGTAC
ACGTGGGTGTTGGGGGAACTCCACAGCGGCAGGCGAGGGTTGGGTGTAGCAGGGGGCCCA
CGGGGGCGGCGGGTGGCAGGCGCGAGGGGGGCGCGGGGGAGCGATCCGATGGTCGAGAAC
GATCGGCTAGGAGGCGAGCCCCAGGTGGCGCCGGGCGGGTGCGCCCGTGTCGGCGCGGGG
GGGTTGTGGCGACGAGGCCGCCCTTCAGGCGAGGTGGGAGTGGCCCACGTTGTGGCGCCG
CGGTGGAGGGGGGGCGGTGAGGTGGGGGGCCAGTCGAGGTTGGTGAGGGCGGGACCAGCC
TCGCTTAGTGCAACGCCAACCCGGATAGTAATTCGTGCGGCTCGTGGGAGCGAGTTCTTC
CAATGATGCAGGGGGGGGTGTGGGGCCCGCGCGTCGACCGGCCGCGTCACGACTGCTACG
GGGTCATTACCGCCGGGCTGTCGTAGGCCGGCGCTCGCGGGAGGTGGTTCGAATGCCGGG
GCGGGAGGCCCGGATCCGGCCAAGGGGCGGGGCTCGGGGGACGCGAGGCGGAGGGGCGCG
AGGCGGTAGCCGGTGCAAGCGGCGCGTGGCCCCATGAGGAAGAACTCTCTCGTGCCCCAA
CACCACACATTAGCGCCCCCATCTCGGCCGACTTCGAGCACGCTCAGCGAGAACCACCAC
AGGCCTCCAACCCGACGTACCACCCGTCCTCACGAGGCGGGCCGACCCTCGGGTCCAAGC
GAGGGCTCCGGCGCGCACGCCCGGCGTGGTGC
>MAP 071_A_tb1189
CCCCTTAGGCGCCTCGAGCGGGCCCCCGCTTCGCCAGTTCACCCGTCCCAGTAAGCGCCC
GGGGCACCTCGCCTTCGAGGAGAACCCCCAGAACGCCCCACCGTCTCCCAGCTCGCACAC
GCGCACGCGTCGCACTCTCCAGCGCGCGTACTGTTCCAGCCACCGTCCCCATGCTCCCCG
CGGGGCCGTATGACGAGCCACGCCCCGGCCCCGCCGCCGTCGCTCGCCCGCCGAGACAAG
CGCAATGCTATTGGGGCCTGACCACGGCTGGGCGGGGCGCTTCCCACGGAGGGCCCAGTC
TTTGGGAGGAGTTGCCCCTCCAGGGATCGGTAGCGCGGTGCGGCTCCCCGCAGCGGCCGC
AGGCGTCGGGGGGATGCTGGGGAGTGGGGGATGCCCCTTCCGGTCGCAGGCGGAACAGGT
TCGCGGTGGTGCACAGGGGCGTGGTGCTGCAGCTCGAGGCCGCGGAGGGGGGCCGCGGTG
GACTTCGGGTCGGAGGGACCGATATACCAGGGCTCTGGCGCGGGGTAGCCCCAGGACGCT
TTCTCGGGGAGGGGAGGGTACGGGCAGCCAGGTGGGAGTGCCAGAACTGGACGGCGTAAT
CCCTGCTGCCTGCGATGTGGGGTCGGACCCAGCGCACCCTGATGGGGCCTGGCGGCGGGG
GACCCCTGAACGGACGTGGCAGCTACTCTGCCGTTGAGCCGATGCGGACGTGCCCAGTAC
ACGTGGGTGTTGGGGGAACTCCACAGCGGCAGGCGAGGGTTGGGTGTAGCAGGGGGCCCA
CGGGGGCGGCGGGTGGCAGGCGCGAGGGGGGCGCGGGGGAGCGATCCGATGGTCGAGAAC
GATCGGCTAGGAGGCGAGCCCCAGGTGGCACCGGGCGGGTGCGCCCGTGTCGGCGCGGGG
GGGTTGTGGCGACGAGGCCGCCCTTCAGGCGAGGTGGGAGTGGCCCAAGTTGTGGCGCCG
CGGTGGAGGGGGGGCGGTGAGGTGGGGGGCCAGTCGAGGTTGGTGAGGGCGGGACCAGCC
TCGTTTAGTGAAACGCCAACCCGGATAGTAATCCGTGCGGCTCGTGGGAGCGAGTTCTTC
CAATGATGCAGGGGGGGGTGTGGGGCCCGCGCGTCGACCGGTCGCGTCACGACTGCTACG
GGGTCATTACCGCCGGGCTGTCGTAGGCCGGCGCTCGCGGGAGGTGGTTCGAATGCCGGG
GCGGGAGGCCCGGATCCGGCGAAGGGGCGGGGCTCGGGGGACGCGAGGCGGAGGGGCGCG
AGGCGGTAGCCGGTGCAAGCGGCGCGTGGCCCCATGAGGAAGAACTCTCTCGTGCCCCAA
CACCACACATTAGCGCCCCCATCTCGGCCGACTTCGAGCACGCTCAACGAGAACCACCAC
AGGCCTCCAACCCGACGTACCACCCGTCCTCACGAGGCGGGCCGACCCTCGGGTCCAAGC
GAGGGCTCCGCCGCGTATGCCCAGCGTGGTGC
>MAP 091_A_fb1194
CCCCTTAGGCGCCTCGAGCGGGCCCCCGCTTCGCCAGTTCACCCGTCCCAGTAAGCGCCC
GGGGCACCTCGCCTTCGAGGAGAACCCCCAGAACGCCCCACCGTCTCCCAGCTCGCACAC
GCGCACGCGTCGCACTCTCCAGCGCGCGTACTGTTCCAGCCACCGTCCCCATGCTCCCCG
CGGGGCCGTATGACGAGCCACGCCCCGGCCCCGCCGCCGCCGCTCGCCCACCGAGACAAG
CGCAATGCTATTGGGGCCTGACCACGGCTGGGCGGGGCGCTTCCCACGGAGGGCCCAGTC
TTTGGGAGGAGTTGCCCCTCCAGGAATCGGTAGCGCGGTGCGGCTCCCCGCAGCGGCCGC
AGGCGTTGGGGGGATGCTGGGGAGTGGGGGATGCCCCTTCCGGTCGCAGGCGGAACAGGT
TCGCGGTGGTGCACAGGGGCGTGGTGCTGCAGCTCGAGGCCGCGGAGGGGGGCCGCGGTG
GACTTCGGGTCGGAGGGACCGATATACCAGGGCTCTGGCGCGGGGTAGCCCCAGGACGCT
TTCTCGGGGAGGGGAGGGTACGGGCAGTCAGGTGGGAGTGCCAGAACTGGACGGCGTAAT
CCCTGCTGCCTGCGATGTGGGGTCGGACCCAGCGCACCCTGATGGGGCCTGGCGGCGGGG
GACCCCTGAACGGACATGGCAGCTACTCTGCCGTTGAGCCGATGCGGACGTGCCCAGTAC
ACGTGGGTGTTGGGGGAACTCCACAGCGGCAGGCGAGGGTTGGGTGTAGCAGGGGGCCCA
CGGGGGCGGCGGGTGGCAGGCGCGAGGGGGGCGCGGGGGAGCGATCCGATGGTCGAGAAC
GACCGGCTAGGAGGCGAGCCCCAGGTGGCACCGGGCGGGTGCGCCCGTGTCGGCGCGGGG

GGGTTGTGGCGACGAGGCCGCCCTTCAGGCGAGGTGGGAGTGGCCCAAGTTGTGGCGCCG
CGGTGGAGGGGGGGTGGTGAGGTGGGGGGCCAGTCGAGGTTGGTGAGGGCGGGACCAGCC
TCGTTTAGTGAAACGCCAACCCGGATAGTAATTCGTGCGGCTCGTGGGAGCGAGTTCTTC
CAATGATGCAGGGGGGGGTGTGGGGCCCGCGCGTCGACCGGTCGCGTCACGACTGCTACG
GGGTCATTACCGCCGGGCTGTCGTAGGCCGGCGCTCGCGGGAGGTGGTTCGAATGCCGGG
GCGGGAGGCCCGGATCCGGCGAAGGGGCGGGGCTCGGGGGACGCGAGGCGGAGGGGCGCG
AGGCGGTAGCCGGTGCAAGCGGCGCGTGGCCCCATGAGGAAGAACTCTCTCGTGCCCCAA
CACCACACATTAGCGCCCCCATCTCGGCCGACTTCGAGCACGCTCAACGAGAACCACCAC
AGGCCTTCAACCCGACGTACCACCTGTCCTCACGAGGCGGGCCGACCCTCGGGTCCAAGC
GAGGGCTCCGCCGCGCACGCCCGGCGTGGTGC
>MAP 184_A_fb1200
CCCCTTAGGCGCCTCGAGCGGGCCCCCGCTTCGCCAGTTCAGCCGTCCCAGTAAGCGCCC
GGGGCACCTCGCCTTCGAGGAGAACCCCCAGAACGCCCCACCGTCTCCCAGCTCGCACAC
GCGCACGCGTCGCACTCTCCCGCGCGCGTACTGTTCCAGCCACCGTCCCCATGCTCCCCG
CGGGGCCGTATGACGAGCCACGCCCCGGCCCCGCCGCCGCCGCTCGCTCACCGAGACCAG
CGCAATGCTATTGGGGCCTGACCACGGCTGGGCGGGGCGCTTCCCACGGAGGGCCCAGTC
TTTGGGAGGAGTTGCCCCTCCAGGGATCGGCAGCGCGGTGCGGCTCCCCGCAGCGGCCGC
GGGCGTCGGGGGGATGCTAGGGAGTGGGGGATGCCCCTTCCGGTCGCAGGCGGAACAGGT
TCGCGGTGGTGCACAGGGGCGTGGTGCTGCAGCTCGAGGCCGCGGAGGGGGGCCGCGGTG
GACTTCGGGTCGGAGGGACCGATATACCAGGGCTCTGGCGCGGGGTAGCCCCAGGACGCT
TTCTCGGGGAGGGGAGGGTACGGGCAGCCAGGTGGGAGTGCAAGAACTGGACGGCCTAAT
CCCTGCTGCCTGCGATGTGGGGTCGGACCCAGCGCACCCTGATGGGGCCTGGCGGCGGGG
GACCCCTGAACGGACGTGGCAGCTACTCTGCCGTTGAGCCGATGCGGACGTGCCCAGTAC
ACGTGGGTGTTGGGGGAACTCCACAGCGGCAGGCGAGGGTTGGGTGTAGCAGGGGGCCCA
CGGGGGCGGCGGGTGGCAGGCGCGAGGGGGGCGCGGGGGAGCGATCCGATGGTCGAGAAC
GATCGGCTAGGAGGCGAGCCCCAGGTGGCGCCGGGCGGGTGCGCCCGTGTCGGCGCGGGG
GGGTTGTGGCGACGAGGCCGCCCTTCAGGCGAGGTGGGAGTGGCCCACGTTGTGGCGCCG
CGGTGGAGGGGGGGCGGTGAGATGGGGGGCCAGTCGAGGTTGGTGAGGGCGGGACCAGCC
TCGCTTAGTGCAACGCCAACCCGGATAGTAATTCGTGCGGCTCGTGGGAGCGAGTTCTTC
CAATGATGCAGGGGGGGGTGTGGGGCCCGCGCGTCGACCGGCCGCGTCACGACTGCTACG
GGGTCATTACCGCCGGGCTGTCGTAGGCCGGCGCTCGCGGGAGGTGGTTCGAATGCCGGG
GCGGGAGGCCCGGATCCGGCCAAGGGGCGGGGCTCGGGGGACGCGAGGCGGAGGGGCGCG
AGGCGGTAGCCGGTGCAAGCGGCGCGTGGCCCCATGAGGAAGAACTCTCTCGTGCCCCAA
CACCACACATTAGCGCCCCCATCTCGGCCGACTTCGAGCACGCTCAGCGAGAACCACCAC
AGGCCTCCAACCCGACGTACCACCCGTCCTCACGAGGCGGGCCGACCCTCGGGTCCAAGC
GAGGGCTCCGGCGCGCACGCCCGGCGTGGTGC
>MAP 220_A_fb1231
CCCCTTAGGCGCCTCGAGCGGGCCCCCGCTTCGCCAGTTCACCCGTCCCAGTAAGCGCCC
GGGGCACCTCGCCTTCGAGGAGAACCCCCAGAACGCCCCACCGTCTCCCAGCTCGCACAC
GCGCACGCGTCGCACTCTCCAGCGCGCGTACTGTTCCAGCCACCGTCCCCATGCTCCCCG
CGGGGCCGTATGACGAGCCACGCCCCGGCCCCGCCGCCGCCGCTCGCCCACCGAGACAAG
CGCAATGCTATTGGGGCCTGACCACGGCTGGGCGGGGCGCTTCCCACGGAGGGCCCAGTC
TTTGGGAGGAGTTACCCCTCCAGGGATCGGTAGCGCGGTGCGGCTCCCCGCAGCGGCCGC
GGGCGTCGGGGGGATGCTGGGGAGTGGGGGATGCCCCTTCCGGTCGCAGGCGGAACAGGT
TCGCGGTGGTGCACAGGGGCGTGGTGCTGTAGCTCGAGGCCGCGGAGGGGGGCCGCGGTG
GACTTCGGGTCGGAGGGACTGATATACCAGGGCTCTGGCGCGGGGTAGCCCCAGGACGCT
TTCTCGGGGAGGGGAGGGTACGGGCAGCCAGGTGGGAGTGCAAGAACTGGACGGCGTAAT
CCCTGCTGCCTGCGATGTGGGGTCGGACCCAGCGCACCCTGATGGGGCCTGGCGGCGGGG
GACCCCTGAGCGGACGTGGCAGCTACTCTGCCGTTGAGCCGATGCGGACGTGCCCAGTAC
ACGTGGGTGTTGGGGGAACTCCACAGCGGCAGGCGAGGGTTGGGTGTAGCAGGGGGCCCA
CGGGGGCGGCGGGTGGCAGGCGCGAGGGGGGCGCGGGGGAGCGATCCGATGGTCGAGAAC
GATCGGCTAGGAGGCGAGCCCCAGGTGGCGCCGGTCGGGTGCGCCCGTGTCGGCGCGGGG
GGGTTGTGGCGACGAGGCCGCCCTTCAGGCGAGGTGGGAGTGGCCCAAGTTGTGGCGCCG
CGGTGGAGGGGGGGCGGTGAGGTGGGGGGCCAGTCGAGGCTGGTGAGAGCGGGACCAGCC
TCGCTTAGTGCAACGCCAACCTGGATAGTAATTCGTGAGGCTCGTGGGAGCGAGTTCTTC
CAACGATGCAGGGGGGGGTGTGGGGCCCGCGCGTCGACCGGCCGCGTCACGACTGCTACG
GGGTCATTACCGCCGGGCTGTCGTAGGCCGGCGCTCGCGGGAGGTGGTTCGAATGCCGGG
GCGGGAGGCCCGGATCCGGCGAAGGGGCGGGGCTCGGGGGACGCGTGGCGGAGGAGCGCG
AGGCGGTAGCCGGTGCAACCGGCGCGTGGCCCCATGAGGAAGAACTCTCTCGTGCCCCAA
CATCACACATTAGCGCCCCCATCTCGGCCGACTTCGAGCGCGCTCAGCGAGAACCACCAC
AGGCCTCCAACCCGACGTACCACCCGTCCTCACGAGGCGGGCCGACCCTCGGATCCAAGC
GAGGGCTCCGGCGCGCACGCCCGGCGTGGTGC
>MAP 206_A_fb1231
CCCCTTAGGCGCCTCGAGCGGGCCCCCGCTTCGCCAGTTCACCCGTCCCAGTAAGCGCCC
GGGGCACCTCGCCTTCGAGGAGAACCCCCAGAACGCCCCACCGTCTCCCAGCTCGCACAC
GCGCACGCGTCGCACTCTCCAGCGCGCGTACTGTTCCAGCCACCGTCCCCATGCTCCCCG
CGGGGCCGTATGACGAGCCACGCCCCGGCCCCGCCGCCGCCGCTCGCCCACCGAGACAAG
CGCAATGCTATTGGGGCCTGACCACGGCTGGGCGGGGCGCTTCCCACGGAGGGCCCAGTC

TTTGGGAGGAGTTGCCCCTCCAGGGATCGGTAGCGCGGTGCGGCTCCCCGCAGCGGCCGC
AGGCGTCGGGGGGATGCTGGGGAGTGGGGGATGCCCCTGCCGGTCGCAGGCGGAACAGGT
TCGCGGTGGTGCACAGGGGCGTGGTGCTGCAGCTCGAGGCCGCGGAGGGGGGCCGCGGTG
SACTTCGGGTCGGAGGGACCGGGATACCAGGGCTCTGGCGCGGGGTAGCCCCAGGACGCT
TTCTCGGGGAGGGGAGGGTACGGGCAGCCAGGTGGGAGTGCCAGAACTGGACGGCGTAAT
CCCTGCTGCCTGCGATGTGGGGTCGGACCCAGCGCACCCTGATGGGGCCTGGCGGCGGGG
GACCCCTGAACGGACGTGGCAGCTACTCTGCCGTTGAGCCGATGCGGACGTGCCCAGTAC
ACGTGGGTGTTGGGGGAACTCCACAGCGGCAGGCGAGGGTTGGGTGTAGCAGGGGGCCCA
CGGGGGCGGCGGGTGGCAGGCGCGAGGGGGGCGCGGGGGAGCGATCCGATGGTCGAGAAC
GATCGGCTAGGAGGCGAGCCCCAGGTGGCACCGGGCGGGTGCGCCCGTGTCGGCGCGGGG
GGGTTGTGGCGACGAGGCCGCCCTTCAGGCGAGGTGGGAGTGGCCCAAGTTGTGGCGCCG
CGGTGGAGGGGGGGCGGTGAGGTGGGGGGCCAGTCGAGGTTGGTGAGGGCGGGACCAGCC
TCGTTTAGTGAAACGCCAACCCGGATAGTAATTCGTGCGGCTCGTGGGAGCGAGTTCTTC
CAATGATGCAGGGGGGGGTGTGGGGCCCGCGCGTCGACCGGTCGCGTCACGACTGCTACG
GGGTCATTACCGCCGGGCTGTCGTAGGCCGGCGCTCGCGGAGGGAGGTTCGAATGCCGGG
GCGGGAGGCCCGGATCCGGCGAAGGGGCGGGGCTCGGGGGACGCGAGGCGGAGGGGCGCG
AGGCGGTAGCCGGTGCAAGCGGCGCGTGGCCCCATGAGGAAGAACTCTCTCGTGCCCCAA
CACCACACATTAGCGCCCCCATCTCGGCCGACTTCGAGCACGCTCAACGAGAACCACCAC
AGGCCTCCAACCCGACGTACCACCCGTCCTCACGAGGCGGGCCGACCCTCGGGTCCAAGC
GAGGGCTCCGCCGCGCACGCCCGGCGTGGTGC
>MAP 230_A_tb1231
CCCCTTAGGCGCCTCGAGCGGGCCCCCGCTTCGCCAGTTCACCCGTCCCAGTAAGCGCCC
GGGGCACCTCGCCTTCGAGGAGAACCCCCAGAACGCCCCACCGTCTCCCAGCTCGCACAC
GCGCACGCGTCGCACTCTCCAGCGCGCGTACTGTTCCAGCCACCGTCCCCATGCTCCCCG
CGGGGCCGTATGACGAGCCACGCCCCGGCCCCGCCGCCGCCGCTCGCCCACCGAGACAAG
CGCAATGCTATTGGGGCCTGACCACGGCTGGGCGGGGCGCTTCCCACGGAGGGCCCAGTC
TTTGGGAGGAGTTGCCCCTCCAGGGATCGGTAGCGCGGTGCGGCTCCCCGCAGCGGCCGC
AGGCGTCGGGGGGATGCTGGGGAGTGGGGGATGCCCCTGCCGGTCGCAGGCGGAACAGGT
TCGCGGTGGTGCACAGGGGCGTGGTGCTGCAGCTCGAGGCCGCGGAGGGGGGCCGCGGTG
GACTTCGGGTCGGAGGGACCGGGATACCAGGGCTCTGGCGCGGGGTAGCCCCAGGACGCT
TTCTCGGGGAGGGGAGGGTACGGGCAGCCAGGTGGGAGTGCCAGAACTGGACGGCGTAAT
CCCTGCTGCCTGCGATGTGGGGTCGGACCCAGCGCACCCTGATGGGGCCTGGCGGCGGGG
GACCCCTGAACGGACGTGGCAGCTACTCTGCCGTTGAGCCGATGCGGACGTGCCCAGTAC
ACGTGGGTGTTGGGGGAACTCCACAGCGGCAGGCGAGGGTTGGGTGTAGCAGGGGGCCCA
CGGGGGCGGCGGGTGGCAGGCGCGAGGGGGGCGCGGGGGAGCGATCCGATGGTCGAGAAC
GATCGGCTAGGAGGCGAGCCCCAGGTGGCACCGGGCGGGTGCGCCCGTGTCGGCGCGGGG
GGGTTGTGGCGACGAGGCCGCCCTTCAGGCGAGGTGGGAGTGGCCCAAGTTGTGGCGTCG
CGGTGGAGGGGGGGCGGTGAGGTGGGGGGCCAGTCGAGGTTGGTGAGGGCGGGACCAGCC
TCGTTTAGTGAAACGCCAACCCGGATAGTAATTCGTGCGGCTCGTGGGAGCGAGTTCTTC
CAATGATGCAGGGGGGGGTGTGGGGCCCGCGCGTCGACCGGTCGCGTCACGACTGCTACG
GGGTCATTACCGCCGGGCTGTCGTAGGCCGGCGCTCGCGGAGGGAGGTTCGAATGCCGGG
GCGGGAGGCCCGGATCCGGCGAAGGGGCGGGGCTCGGGGGACGCGAGGCGGAGGGGCGCG
AGGCGGTAGCCGGTGCAAGCGGCGCGTGGCCCCATGAGGAAGAACTCTCTCGTGCCCCAA
CACCACACATTAGCGCCCCCATCTCGGCCGACTTCGAGCACGCTCAACGAGAACCACCAC
AGGCCTCCAACCCGACGTACCACCCGTCCTCACGAGGCGGGCCGACCCTCGGGTCCAAGC
GAGGGCTCCGCCGCGCACGCCCGGCGTGGTGC
>MAP 224_A_tb1243
CCCCTTAGGCGCCTCGAGCGGGCCCCCGCTTCGCCAGTTCACCCGTCCCAGTAAGCGCCC
GGGGCACCTCGCCTTCGAGGAGAACCCCCAGAACGCCCCACCGTCTCCCAGCTCGCACAC
GCGCACGCGTCGCACTCTCCAGCGCGCGTACTGTTCCAGCCACCGTCCCCATGCTCCCCG
CGGGGCCGTATGACGAGCCACGCCCCGGCCCCGCCGCCGCCGCTCGCCCACCGAGACAAG
CGCAATGCTATTGGGGCCTGACCACGGCTGGGCGGGGCGCTTCCCACGGAGGGCCCAGTC
TTTGGGAGGAGTTGCCCCTCCAGGGATCGGTAGCGCGGTGCGGCTCCCCGCAGCGGCCGC
AGGCGTCGGGGGGATGCTGGGGAGTGGGGGATGCCCCTGCCGGTCGCAGGCGGAACAGGT
TCGCGGTGGTGCACAGGGGCGTGGTGCTGCAGCTCGAGGCCGCGGAGGGGGGCCGCGGTG
GACTTCGGGTCGGAGGGACCGGGATACCAGGGCTCTGGCGCGGGGTAGCCCCAGGACGCT
TTCTCGGGGAGGGGAGGGTACGGGCAGCCAGGTGGGAGTGCCAGAACTGGACGGCGTAAT
CCCTGCTGCCTGCGATGTGGGGTCGGACCCAGCGCACCCTGATGGGGCCTGGCGGCGGGG
GACCCCTGAACGGACGTGGCAGCTACTCTGCCGTTGAGCCGATGCGGACGTGCCCAGTAC
ACGTGGGTGTTGGGGGAACTCCACAGCGGCAGGCGAGGGTTGGGTGTAGCAGGGGGCCCA
CGGGGGCGGCGGGTGGCAGGCGCGAGGGGGGCGCGGGGGAGCGATCCGATGGTCGAGAAC
GATCGGCTAGGAGGCGAGCCCCAGGTGGCACCGGGCGGGTGCGCCCGTGTCGGCGCGGGG
GGGTTGTGGCGACGAGGCCGCCCTTCAGGCGAGGTGGGAGTGGCCCAAGTTGTGGCGCCG
CGGTGGAGGGGGGGCGGTGAGGTGGGGGGCCAGTCGAGGTTGGTGAGGGCGGGACCAGCC
TCGTTTAGTGAAACGCCAACCCGGATAGTAATTCGTGCGGCTCGTGGGAGCGAGTTCTTC
CAATGATGCAGGGGGGGGTGTGGGGCCCGCGCGTCGACCGGTCGCGTCACGACTGCTACG
GGGTCATTACCGCCGGGCTGTCGTAGGCCGGCGCTCGCGGAGGGAGGTTCGAATGCCGGG
GCGGGAGGCCCGGATCCGGCGAAGGGGCGGGGCTCGGGGGACGCGAGGCGGAGCGGCGCG

AGGCGGTAGCCGGTGCAAGCGGCGCGTGGCCCCATGAGGAAGAACTCTCTCGTGCCCCAA
CACCACACATTAGCGCCCCCATCTCGGCCGACTTCGAGCACGCTCAACGAGAACCACCAC
AGGCCTCCAACCCGACGTACCACCCGTCCTCACGAGGCGGGCCGCCCCTCGGGTCCAAGC
GAGGGCTCCGCCGCGCACGCCCGGCGTGGTGC
>MAP 180_A_tb1249
CCCCTTAGGCGCCTCGAGCGGGCCCCCGCTTCGCCAGTTCACCCGTCCCAGTAAGCGCCC
GGGGCACCTCGCCTTCGAGGAGAACCCCCAGAACGCCCCACCGTCTCCCAGCTCGCACAC
GCGCACGCGTCGCACTCTCCAGCGCGCGTACTGTTCCAGCCACCGTCCCCATGCTCCCCG
CGGGGCCGTATGACGAGCCACGCCCCGGCCCCGCCGCCGCCGCTCGCCCACCGAGACAAG
CGCAATGCTATTGGGGCCTGACCACGGCTGGGCGGGGCGCTTCCCACGGAGGGCCCAGTC
TTTGGGAGGAGTTGCCCCTCCAGGGATCGGTAGCGCGGTGCGGCTCCCCGCAGCGGCCGC
AGGCGTCGGGGGGATGCTGGGGAGTGGGGGATGCCCCTGCCGGTCGCAGGCGGAACAGGT
TCGCGGTGGTGCACAGGGGCGTGGTGCTGCAGCTCGAGGCCGCGGAGGGGGGCCGCGGTG
GACTTCGGGTCGGAGGGACCGGGATACCAGGGCTCTGGCGCGGGGTAGCCCCAGGACGCT
TTCTCGGGGAGGGGAGGGTACGGGCAGCCAGGTGGGAGTGCCAGAACTGGACGGCGTAAT
CCCTGCTGCCTGCGATGTGGGGTCGGACCCAGCGCACCCTGATGGGGCCTGGCGGCGGGG
GACCCCTGAACGGACGTGGCAGCTACTCTGCCGTTGAGCCGATGCGGACGTGCCCAGTAC
ACGTGGGTGTTGGGGGAACTCCACAGCGGCAGGCGAGGGTTGGGTGTAGCAGGGGGCCCA
CGGGGGCGGCGGGTGGCAGGCGCGAGGGGGGCGCGGGGGAGCGATCCGATGGTCGAGAAC
GATCGGCTAGGAGGCGAGCCCCAGGTGGCACCGGGCGGGTGCGCCCGTGTCGGCGCGGGG
GGGTTGTGGCGACGAGGCCGCCCTTCAGGCGAGGTGGGAGTGGCCCAAGTTGTGGCGCCG
CGGTGGAGGGGGGGCGGTGAGGTGGGGGGCCAGTCGAGGTTGGTGAGGGCGGGACCAGCC
TCGTTTAGTGAAACGCCAACCCGGATAGTAATTCGTGCGGCTCGTGGGAGCGAGTTCTTC
CAATGATGCAGGGGGGGGTGTGGGGCCCGCGCGTCGACCGGTCGCGTCACGACTGCTACG
GGTTGATTACCGCCGGGCTGTCGTAGGCCGGCGCTCGCGGAGGGAGGTTCGAATGCCGGG
GCGGGAGGCCCGGATCCGGCGAAGGGGCGGGGCTCGGGGGACGCGAGGCGGAGGGGCGCG
AGGCGGTAGCCGGTGCAAGCGGCGCGTGGCCCCATGAGGAAGAACTCTCTCGTGCCCCAA
CACCACACATTAGCGCCCCCATCTCGGCCGACTTCGAGCACGCTCAACGAGAACCACCAC
AGGCCTCCAACCCGACGTACCACCCGTCCTCACGAGGCGGGCCGACCCTCGGGTCCAAGC
GAGGGCTCCGCCGCGCACGCCCGGCGTGGTGC
>MAP 004_A_fb1255
CCCCTTAGGCGCCTCGAGCGGGCCCCCGCTTCGCCAGTTCACCCGTCCCAGTAAGCGCCC
GGGGCACCTCGCCTTCGAGGAGAACCCCCAGAACGCCCCACCGTCTCCCAGCTCGCACAC
GCGCACGCGTCGCACTCTCCAGCGCGCGTACTGTTCCAGCCACCGTCCCCATGCTCCCCG
CGGGGCCGTATGACGAGCCACGCCCCGGCCCCGCCGCCGCCGCTCGCCCACCGAGACAAG
CGCAATGCTATTGGGGCCTGACCACGGCTGGGCGGGGCGCTTCCCACGGAGGGCCCAGTC
TTTGGGAGGAGTTGCCCCTCCAGGGATCGGTAGCGCGGTGCGGCTCCCCGCAGCGGCCGC
AGGCGTCGGGGGGATGCTGGGGAGTGGGGGATGCCCCTGCCGGTCGCAGGCGGAACAGGT
TCGCGGTGGTGCACAGGGGCGTGGTGCTGCAGCTCGAGGCCGCGGAGGGGGGCCGCGGTG
GACTTCGGGTCGGAGGGACCGGGATACCAGGGCTCTGGCGCGGGGTAGCCCCAGGACGCT
TTCTCGGGGAGGGGAGGGTACGGGCAGCCAGGTGGGAGTGCCAGAACTGGACGGCGTAAT
CCCTGCTGCCTGCGATGTGGGGTCGGACCCAGCGCACCCTGATGGGGCCTGGCGGCGGGG
GACCCCTGAACGGACGTGGCAGCTACTCTGCCGTTGAGCCGATGCGGACGTGCCCAGTAC
ACGTGGGTGTTGGGGGAACTCCACAGCGGCAGGCGAGGGTTGGGTGTAGCAGGGGGCCCA
CGGGGGCGGCGGGTGGCAGGCGCGAGGGGGGCGCGGGGGAGCGATCCGATGGTCGAGAAC
GATCGGCTAGGAGGCGAGCCCCAGGTGGCACCGGGCGGGTGCGCCCGTGTCGGCGCGGGG
GGGTTGTGGCGACGAGGCCGCCCTTCAGGCGAGGTGGGAGTGGCCCAAGTTGTGGCGCCG
CGGTGGAGGGGGGGCGGTGAGGTGGGGGGCCAGTCGAGGTTGGTGAGGGCGGGACCAGCC
TCGTTTAGTGAAACGCCAACCCGGATAGTAATTCGTGCGGCTCGTGGGAGCGAGTTCTTC
CAATGATGCAGGGGGGGGTGTGGGGCCCGCGCGTCGACCGGTCGCGTCACGACTGCTACG
GGGTCATTACCGCCGGGCTGTCGTAGGCCGGCGCTCGCGGAGGGAGGTTCGAATGCCGGG
GCGGGAGGCCCGGATCCGGCGAAGGGGCGGGGCTCGGGGGACGCGAGGCGGAGGGGCGCG
AGGCGGTAGCCGGTGCAAGCGGCGCGTGGCCCCATGAGGAAGAACTCTCTCGTGCCCCAA
CACCACACATTAGCGCCCCCATCTCGGCCGACTTCGAGCACGCTCAACGAGAACCACCAC
AGGCCTCCAACCCGACGTACCACCCGTCCTCACGAGGCGGGCCGACCCTCGGGTCCAAGC
GAGGGCTCCGCCGCGCACGCCCGGCGTGGTGC
>MAP 191_A_fb1255
CCCCTTAGGCGCCTCGAGCGGGCCCCCGCTTCGCCAGTTCACCCGTCCCAGTAAGCGCCC
GGGGCACCTCGCCTTCGAGGAGAACCCCCAGAACGCCCCACCGTCTCCCAGCTCGCACAC
GCGCACGCGTCGCACTCTCCAGCGCGCGTACTGTTCCAGCCACCGTCCCCATGCTCCCCG
CGGGGCCGTATGACGAGCCACGCCCCGGCCCCGCCGCCGCCGCTCGCCCACCGAGACAAG
CGCAATGCTATTGGGGCCTGACCACGGCTGGGCGGGGCGCTTCCCACGGAGGGCCCAGTC
TTTGGGAGGAGTTGCCCCTCCAGGGATCGGTAGCGCGGTGCGGCTCCCCGCAGCGGCCGC
AGGCGTCGGGGGGATGCTGGGGAGTGGGGGATGCCCCTGCCGGTCGCAGGCGGAACAGGT
TCGCGGTGGTGCACAGGGGCGTGGTGCTGCAGCTCGAGGCCGCGGAGGGGGGCCGCGGTG
GACTTCGGGTCGGAGGGACCGGGATACCAGGGCTCTGGCGCGGGGTAGCCCCAGGACGCT
TTCTCGGGGAGAGGAGGGTACGGGCAGCCAGGTGGGAGTGCCAGAACCGGACGGCGTAAT
CCCTGCTGCCTGCGATGTGGGGTCGGACCCAGCGCACCCTGATGGGGCCTGGCGGCGGGG

GACCCCTGAACGGACGTGGCAGCTACTCTGCCGTTGAGCCGATGCGGACGTGCCCAGTAC
ACGTGGGTGTTGGGGGAACTCCACAGCGGCAGGCGAGGGTTGGGTGTAGCAGGGGGCCCA
CGGGGGCGGCGGGTGGCAGGCGCGAGGGGGGCGCGGGGGAGCGATCCGATGGTCGAGAAC
GATCGGCTAGGAGGCGAGCCCCAGGTGGCACCGGGCGGGTGCGCCCGTGTCGGCGCGGGG
GGGTTGTGGCGACGAGGCCGCCCTTCAGGCGAGGTGGGAGTGGCCCAAGTTGTGGCGCCG
CGGTGGAGGGGGGGCGGTGAGGTGGGGGGCCAGTCGAGGTTGGTGAGGGCGGGACCAGCC
TCGTTTAGTGAAACGCCAACCCGGATAGTAATTCGTGCGGCTCGTGGGAGCGAGTTCTTC
CAATGATGCAGGGGGGGGTGTGGGGCCCGCGCGTCGACCGGTCGCGTCACGACTGCTACG
GGGTCATTACCGCCGGGCTGTCGTAGGCCGGCGCTCGCGGAGGGAGGTTCGAATGCCGGG
GCGGGAGGCCCGGATCCGGCGAAGGGGCGGGGCTCGGGGGACGCGAGGCGGAGGGGCGCG
AGGCGGTAGCCGGTGCAAGCGGCGCGTGGCCCCATGAGGAAGAACTCTCTCGTGCCCCAA
CACCACACATTAGCGCCCCCATCTCGGCCGACTTCGAGCACGCTCAACGAGAACCACCAC
AGGCCTCCAACCCGACGTACCACCCGTCCTCACGAGGCGGGCCGACCCTCGGGTCCAAGC
GAGGGCTCCGCCGCGCACGCCCGGCGTGGTGC
>MAP 166_A_tb1257
CCCCTTAGGCGCCTCGAGCGGGCCCCCGCTTCGCCAGTTCAGCCGTCCCAGTAAGCGCCC
GGGGCACCTCGCCTTCGAGGAGAACCCCCAGAACGCCCCACCGTCTCCCAGCTCGCACAC
GCGCACGCGTCGCACTCTCCCGCGCGCGTACTGTTCCAGCCACCGTCCCCATGCTCCCCG
CGGGGCCGTATGACGAGCCACGCCCCGGCCCCGCCGCCGCCGCTCGCTCACCGAGACCAG
CGCAATGCTATTGGGGCCTGACCACGGCTGGGCGGGGCGCTTCCCACGGAGGGCCCAGTC
TTTGGGAGGAGTTGCCCCTCCAGGGATCGGCCGCGCGGTGCGGCTCCCCGCAGCGGCCGC
GGGCGTCGGGGGGATGCTAGGGAGTGGGGGATGCCCCTTCCGGTCGCAGGCGGAACAGGT
TCGCGGTGGTGCACAGGGGCGTGGTGCTGCAGCTCGAGGCCGCGGAGGGGGGCCGCGGTG
GACTTCGGGTCGGAGGGACCGATATACCAGGGCTCTGGCGCGGGGTAGCCCCAGGACGCT
TTCTCGGGGAGGGGAGGGTACGGGCAGCCAGGTGGGAGTGCAAGAACTGGACGGCCTAAT
CCCTGCTGCCTGCGATGTGGGGTCGGACCCAGCGCACCCTGATGGGGCCTGGCGGCGGGG
GACCCCTGAACGGACGTGGCAGCTACTCTGCCGTTGAGCCGATGCGGACGTGCCCAGTAC
ACGTGGGTGTTGGGGGAACTCCACAGCGGCAGGCGAGGGTTGGGTGTAGCAGGGGGCCCA
CGGGGGCGGCGGGTGGCAGGCGCGAGGGGGGCGCGGGGGAGCGATCCGATGGTCGAGAAC
GATCGGCTAGGAGGCGAGCCCCAGGTGGCGCCGGGCGGGTGCGCCCGTGTCGGCGCGGGG
GGGTTGTGGCGACGAGGCCGCCCTTCAGGCGAGGTGGGAGTGGCCCACGTTGTGGCGCCG
CGGTGGAGGGGGGGCGGTGAGGTGGGGGGCCAGTCGAGGTTGGTGAGGGCGGGACCAGCC
TCGCTTAGTGCAACGCCAACCCGGATAGTAATTCGTGCGGCTCGTGGGAGCGAGTTCTTC
CAATGATGCAGGGGGGGGTGTGGGGCCCGCGCGTCGACCGGCCGCGTCACGACTGCTACG
GGGTCATTACCGCCGGGCTGTCGTAGGCCGGCGCTCGCGGGAGGTGGTTCGAATGCCGGG
GCGGGAGGCCCGGATCCGGCCAAGGGGCGGGGCTCGGGGGACGCGAGGCGGAGGGGCGCG
AGGCGGTAGCCGGTGCAAGCGGCGCGTGGCCCCATGAGGAAGAACTCTCTCGTGCCCCAA
CACCACACATTAGCGCCCCCATCTCGGCCGACTTCGAGCACGCTCAGCGAGAACCACCAC
AGGCCTCCAACCCGACGTACCACCCGTCCTCACGAGGCGGGCCGACCCTCGGGTCCAAGC
GAGGGCTCCGGCGCGCACGCCCGGCGTGGTGC
>MAP 167_A_tb1257
CCCCTTAGGCGCCTCGAGCGGGCCCCCGCTTCGCCAGTTCAGCCGTCCCAGTAAGCGCCC
GGGGCACCTCGCCTTCGAGGAGAACCCCCAGAACGCCCCACCGTCTCCCAGCTCGCACAC
GCGCACGCGTCGCACTCTCCCGCGCGCGTACTGTTCCAGCCACCGTCCCCATGCTCCCCG
CGGGGCCGTATGACGAGCCACGCCCCGGCCCCGCCGCCGCCGCTCGCTCACCGAGACCAG
CGCAATGCTATTGGGGCCTGACCACGGCTGGGCGGGGCGCTTCCCACGGAGGGCCCAGTC
TTTGGGAGGAGTTGCCCCTCCAGGGATCGGCAGCGCGGTGCGGCTCCCCGCAGCGGCCGC
GGGCGTCGGGGGGATGCTAGGGAGTGGGGGATGCCCCTTCCGGTCGCAGGCGGAACAGGT
TCGCGGTGGTGCACAGGGGCGTGGTGCTGCAGCTCGAGGCCGCGGAGGGGGGCCGCGGTG
GACTTCGGGTCGGAGGGACCGATATACCAGGGCTCTGGCGCGGGGTAGCCCCAGGACGCT
TTCTCGGGGAGGGGAGGGTACGGGCAGCCAGGTGGGAGTGCAAGAACTGGACGGCCTAAT
CCCTGCTGCCTGCGATGTGGGGTCGGACCCAGCGCACCCTGATGGGGCCTGGCGGCGGGG
GACCCCTGAACGGACGTGGCAGCTACTCTGCCGTTGAGCCGATGCGGACGTGCCCAGTAC
ACGTGGGTGTTGGGGGAACTCCACAGCGGCAGGCGAGGGTTGGGTGTAGCAGGGGGCCCA
CGGGGGCGGCGGGTGGCAGGCGCGAGGGGGGCGCGGGGGAGCGATCCGATGGTCGAGAAC
GATCGGCTAGGAGGCGAGCCCCAGGTGGCGCCGGGCGGGTGCGCCCGTGTCGGCGCGGGG
GGGTTGTGGCGACGAGGCCGCCCTTCAGGCGAGGTGGGAGTGGCCCACGTTGTGGCGCCG
CGGTGGAGGGGGGGCGGTGAGGTGGGGGGCCAGTCGAGGTTGGTGAGGGCGGGACCAGCC
TCGCTTAGTGCAACGCCAACCCGGATAGTAATTCGTGCGGCTCGTGGGAGCGAGTTCTTC
CAATGATGCAGGGGGGGGTATGGGGCCCGCGCGTCGACCGGCCGCGTCACGACTGCTACG
GGGTCATTACCGCCGGGCTGTCGTAGGCCGGCGCTCGCGGGAGGTGGTTCGAATGCCGGG
GCGGGAGGCCCGGATCCGGCCAAGGGGCGGGGCTCGGGGGACGCGAGGCGGAGGGGCGCG
AGGCGGTAGCCGGTGCAAGCGGCGCGTGGCCCCATGAGGAAGAACTCTCTCGTGCCCCAA
CACCACACATTAGCGCCCCCATCTCGGCCGACTTCGAGCACGCTCAGCGAGAACCACCAC
AGGCCTCCAACCCGACGTACCACCCGTCCTCACGAGGCGGGCCGACCCTCGGGTCCAAGC
GAGGGCTCCGGCGCGCACGCCCGGCGTGGTGC
>MAP 120_A_tb1261
CCCCTTAGGCGCCTCGAGCGGGTCCCCGCTTCGCCAGTTCACCCGTCCCAGTAAGCGCCC

GCGGCACCTCGCCTTCGCGGAGAACCCCCAGAACGCCCCACCGTCTCCCAGCTCGCACAC
GCGCACGTGTCGCGCTCTCCAGCGCGCGGACTGTTCCAGCCACTGTTCTCATGCTCCCCG
CGGGGCCGTATGACGAGCCACGCCCCAGCCCCGCCGCCGCCGCTCGTCCACCGAGACAAG
CGTAACGCTATTGGGGCCTGACCATGGCTGGGCGGGGCGCTTCCCACGGAGGGCCCAGTC
TTTGGGAGGAGTTGCCCCTCAAGGGATCGGTAGCGCGGTGCGGCTCCCCGCAGCGGCCGC
GAGCGTCGGGGTGATGCTGGAGAGTGGGGCATGCCCCTTCCGGCCGCAGGCGGAACAGGT
TCGCGGTGGTGCACAGGGACGCGGTGCTGCAGCTCGAGACCGCGGAGGGGAGCCGCGGTG
GACTTCGGGTCGGAGGGACCGATATACCAGGGCTCTGACGCGGGGTAACTCTAGGACGCT
CTCTCGAGGAGGGGAAGGTACGGGCAGCCAGGTGGGCGTGCAGGAACTGGACGGCGTAAT
CCCTGGTGACTGCGATGTGGGGTAGGACCCAGCGCACCCTGATGGGGCCTGGCGGCGGGG
GACCCCTGGACGGACGTGGCAGCTACCCTGCCGTTGAGCCGATGCGGACGTGCCTGGTAC
ACGTGGGTGTTGGGGGAACTCCACAGCGGCAGGCGACGGTTGGGTGGGGCCGGGGGCCCA
CGGGGGAGGCGGGTGGCGGTCACGAGGGGGACACGGGGGAGCGATCCGATGGTCGAGAAC
GATCGGCTAGGAGGCGAGCCCCAGGTGGCGCAGGGCGGGTGCGCCCGTGTCGGCGCGGGG
GGGAGGTGGCGACGAGGCGGCCCTTCAGGCGGGGTGAGAATGGCCCAAGTTATGGCGCCG
CGGTGGAGGGGGGGCGGTGAGGTGGGGGGCCAGTCGGGGTTGGTGAGGGCGGGACCAGCC
TCGCTTGATGCAACGCCAACCCGGATAGTAATTCGTGCGGCTCGTGGTCGCGAGTTCTTC
CAATGATTCAGGGGGGGGTGTGGGGCCCGCCCGTCGACCGGCCGCGTCACGACTGCTACG
GGGTCATTACCGCCGGGCTGTCGTAGGCCGGCGATCGCGGGAGGTGGTTCGAATGCCGCG
GCGGGAGGCCCGGATCAAGCGAAGGGGCGGGGCCCGGGGCACGCGAGGCGGGGGGGGGCG
AGACGGTAGCCGGTGCAGGCGACGCGTGGCCCCATGAGGTAGAACTCTCTCGTGCCCCAA
CACCACACATTAGCGCCCCCATCTCGGCCGACTTCGAGCGCGCTCAGCGAGAACCACCAC
AGGCCTCCAACCCGACGTACTACCCGTCCTCACGAGGCGGGCCGACCCCCGGGCCCAAGC
GAAGGCTCCGGCGCGCACACCCGGCGTGCTGC
>MAP 121_A_tb1261
CCCCTTGGGCGCCTCGAGCGGGTCCCCGCTTCGCCAGTTCACCCGTCCCAGTAAGCGCCC
GCAGCACCTCGCCTTCGCGGAGAACCCCCAGAACGCCCCACCGTCTCCCAGCTCGCACAC
GCGCACGTGTCGCACTCTCCAGCGCGCGGACTGTTCCAGCCACCGTTTTCATGCTCCCGG
CGGGGCCGTATGACGAGCCACGCCCCAGCCCCGCCGCCGCCGCTCGCCCACCGAGACAAG
CGTAACGCTATTGGGGCCTGACCATGGCTGGGCGGGGTGCTTCCCACGGAGGGCCCAGTC
TTTGGGAGGAGTTGCCCCTCAAGGGATCGGTAGCGCAGTGCGGCTCCCCGCAGCGGCCGC
GAGCGTCGGGGTGATGCTGGAGAGTGGGGCATGCCCCTTCCGGCCGCAGGCGGAACAGGT
TCGCGGTGGTGCACAGGGACGCGGTGTTGCAGCTCGAGGCCGCGGAGGGGAGCCGCGGTG
GACTTCGGGTCGGAGGGACCAATATACCAGGGCTCTGACGCGGGGTAACTCTAGGACGCC
CTCTCGAGGAGGGGAGGGTACGGGCAGCCAGGTGGGCGTGCAGGAACTGGACGGCGTAAT
CCCTGGTGACTGCGATGTGGGGTAGGACCCAGCGCACCCTGATGGGGCCTGGCGGCGGGG
GACCCCTGGACGGACGTGGCAGCTACCCTGCCGTTGAGCCGATGCGGACGTGCCCAGTAC
ACGTAGGTGTTGGGGGAACTCCACAGCGGCAGGCAACGGTTGGGTGGGGCCGGGGGCCCA
CGGGGGCGGCGGGTGGCGGTCACGAGGGGGGCACGGGGGAGCGATCCGATGGTCGAGAAC
GATCGGCTAGGAGGCGAGCCCCAGGTGGCGCAGGGCGGGTGCGCCCATGTCGGCGCGGGG
GGGAGGTGACGACGAGGCGGCCCTTCAGGCGGGGTGAGAGTGGCCCAAGTTGTGGCGCCG
CGGTGGAGGGGGGGCGGTGAGGTGGGGGGCCAGTCGGAGTTGGTGAGGGCGGGACCAGCC
TCACTTGATGCAACGCCAACCCGGATAGTAATTCGTGCGGCCCGTGGTCGCGAGTTCTTC
CAATGATTCAGGGGGGGGTGTGGGGCCCGCCCGTCGACCGGCCGCGTCACGACTGCTACG
GGGTCATTACCGCCCGGCTGTCGTAGGCCGGCGATCGCGGGAGGTGGTTCGAATGCCGCG
GCGGGAGGCCCGGATCAAGCGAAGGGGCGGGGCCCGGGGCACGCGAGGCGGGGGGGGACG
AGGCGGTAGCCGGTGCAAGCGACGCGTGGCCCCATGAGGTAGAATTCTCTCGTGCCCCAA
CACCACACATTAGCGCCCCCATCTCGGTCGACTTCGAGCGCGCTCAGCGAGAACCACCAC
AGGCCTCCAACCCGACGTACCACCCGTCCTCACGAGGCGGGCCGACCCCCGGGCCCAAGC
GAGGGCTCCGGCGCGCACACCCGGCGTGCTGC
>MAP 122_A_tb1261
CCCCTTAGGCGCCTCGAGCGGGTCCCCRCTTCGCCAGTTCACCCGTCCCAGTAAGCGCCC
GCGGCACCTCGCCTTCGCGGAGAACCCCCAGAACGCCCCACCGTCTCCCAGCTCGCACAC
GCGCACGTGTCGCGCTCTCCAGCGCGCGGACTGTTCCAGCCACTGTTCTCATGCTCCCCG
CGGGGCCGTATGACGAGCCACGCCCCAGCCCCGCCGCCGCCGCTCGCCCACCGAGACAAG
CGTAACGCTATTGGGGCCTGACCATGGCTGGGCGGGGCGCTTCCCACGGAGGGCCCAGTC
TTTGGGAGGAGTTGCCCCTCAAGGGATCGGTAGCGCGGTGCGGCTCCCCGCAGCGGCCGC
GAGCGTCGGGGTGATGCTGGAGAGTGGGGCATGCCCCTTCCGGCCGCAGGCGGAACAGGT
TCGCGGTGGTGCACAGGGACGCGGTGCTGCAGCTCGAGACCGCGGAGGGGAGCCGCGGTG
GACTTCGGGTCGGAGGGACCGATATACCAGGGCTCTGACGCGGGGTAACTCTAGGACGCT
CTCTCGAGGAGGGGAGGGTACGGGCAGCCAGGTGGGCGTGCAGGAACTGGACGGCGTAAT
CCCTGGTGACTGCGATGTGGGGTAGGACCCAGCGCACCCTGATGGGGCCTGGCGGCGGGG
GACCCCTGGACGGACGTGGCAGCTACCCTGCCGTTGAGCCGATGCGGACGTGCCTGGTAC
ACGTGGGTGTTGGGGGAACTCCACAGCGGCAGGCGACGGTTGGGTGGGGCCGGGGGCCCA
CGGGGGAGGCGGGTGGCGGTCACGAGGGGGACACGGGGGAGCGATCCGATGGTCGAGAAC
GATCGGCTAGGAGGCGAGCCCCAGGTGGCGCAGGGCGGGTGCGCCCGTGTCGGCGCGGGG
GGGAGGTGGCGACGAGGCGGCCCTTCAGGCGGGGTGAGAATGGCCCAAGTTATGGCGCCG
CGGTGGAGGGGGGGCGGTGAGGTGGGGGGCCAGTCGGGGTTGGTGAGGGCGGGACCAGCC

TCGCTTGATGCAACGCCAACCCGGATAGTAATTCGTGCGGCTCGTGGTCGCGAGTTCTTC
CAATGATTCAGGGGGGGGTGTGGGGCCCGCCCGTCGACCGGCCGCGTCACGACTGCTACG
GGGTCATTACCGCCGGGCTGTCGTAGGCCGGCGATCGCGGGAGGTGGTTCGAATGCCGCG
GCGGGAGGCCCGGATCAAGCGAAGGGGCGGGGCCCGGGGCACGCGAGGCGGGGGGGGGCG
AGACGGTAGCCGGTGCAGGCGACGCGTGGCCCCATGAGGTAGAACTCTCTCGTGCCCCAA
CACCACACATTAGCGCCCCCATCTCGGCCGACTTCGAGCGCGCTCAGCGAGAACCACCAC
AGGCCTCCAACCCGACGTACTACCCGTCCTCACGAGRCGGGCCGACCCCCGGGCCCAAGC
GAAGGCTCCGGCGCGCACACCCGGCGTGCTGC
>MAP 127_A_tb1283
CCCCTTAGGCGCCTCGAGCGGGTCCCCGCTTCGCCAGTTCACCCGTCCCAGTAAGCGCCC
GCGGCACCTCGCCTTCGCGGAGAACCCCCAGAACGCCCCACCGTCTCCCAGCTCGCACAC
GCGCACGTGTCGCGCTCTCCAGCGCGCGGACTGTTCCAGCCACTGTTCTCATGCTCCCCG
CGGGGCCGTATGACGAGCCACGCCCCAGCCCCGCCGCCGCCGCTCGCCCACCGAGACAAG
CGTAACGCTATTGGGGCCTGACCATGGCTGGGCGGGGCGCTTCCCACGGAGGGCCCAGTC
TTTGGGAGGAGTTGCCCCTCAAGGGATCGGTAGCGCGGTGCGGCTCCCCGCAGCGGCCGC
GAGCGTCGGGGTGATGCTGGAGAGTGGGGCATACCCCTTCCGGCCGCAGGCGGAACAGGT
TCGCGGTGGTGCACAGGGACGCGGTGCTGCAGCTCGAGACCGCGGAGGGGAGCCGCGGTG
GACTTCGGGTCGGAGGGACCGATATACCAGGGCTCTGACGCGGGGTAACTCTAGGACGCT
CTCTCGAGGAGGGGAGGGTACGGGCAGCCAGGTGGGCGTGCAGGAACTGGACGGCGTAAT
CCCTGGTGACTGCGATGTGGGGTAGGACCCAGCGCACCCTGATGGGGCCTGGCGGCGGGG
GACCCCTGGACGGACGTGGCAGCTACCCTGCCGTTGAGCCGATGCGGACGTGCCTGGTAC
ACGTGGGTGTTGGGGGAACTCCACAGCGGCAGGCGACGGTTGGGTGGGGCCGGGGGCCCA
CGGGGGAGGCGGGTGGCGGTCACGAGGGGGACACGGGGGAGCGATCCGATGGTCGAGAAC
GATCGGCTAGGAGGCGAGCCCCAGGTGGCGCAGGGCGGGTGCGCCCGTGTCGGCGCGGGG
GGGAGGTGGCGACGAGGCGGCCCTTCAGGCGGGGTGAGAATGGCCCAAGTTATGGCGCCG
CGGTGGAGGGGGGGCGGTGAGGTGGGGGTCCCGTCGGGGTTGGTGAGGGCGGGACCAGCC
TCGCTTGATGCAACGCCAACCCGGATAGTAATTCGTGCGGCTCGTGGTCGCGAGTTCTTC
CAATGATTCAGGGGGGGGTGTGGGGCCCGCCCGTCGACCGGCCGCGTCACGACTGCTACG
GGGTCATTACCGCCGGGCTGTCGTAGGCCGGCGATCGCGGGAGGTGGTTCGAATGCCGCG
GCGGGAGGCCCGGATCAAGCGAAGGGGCGGGGCCCGGGGCACGCGAGGCGGGGGGGGGCG
AGACGGTAGCCGGTGCAGGCGACGCGTGGCCCCATGAGGTAGAACTCTCTCGTGCCCCAA
CACCACACATTAGCGCCCCCATCTCGGCCGACTTCGAGCGCGCTCAGCGAGAACCACCAC
AGGCCTCCAACCCGACGTACTACCCGTCCTCACGAGGCGGGCCGACCCCCGGGCCCAAGC
GAAGGCTCCGGCGCGCACACCCGGCGTGCTGC
>MAP 128_A_tb1283
CCCCTTAGGCGCCTCGAGCGGGTCCCCGCTTCGCCAGTTCACCCGTCCCAGTAAGCGCCC
GCGGCACCTCGCCTTCGCGGAGAACCCCCAGAACGCCCCACCGTCTCCCAGCTCGCACAC
GCGCACGTGTCGCGCTCTCCAGCGCGCGGACTGTTCCAGCCACTGTTCTCATGCTCCCCG
CGGGGCCGTATGACGAGCCACGCCCCAGCCCCGCCGCCGCCGCTCGCCCACCGAGACAAG
CGTAACGCTATTGGGGCCTGACCATGGCTGAGCGGGGCGCTTCCCACGGAGGGCCCAGTC
TTTGGGAGGAGTTGCCCCTCAAGGGATCGGTAGCGCGGTGCGGCTCCCCGCAGCGGCCGC
GAGCGTCGGGGTGATGCTGGAGAGTGGGGCATGCCCCTTCCGGCCGCAGGCGGAACAGGT
TCGCGGTGGTGCACAGGGACGCGGTGCTGCAGCTCGAGACCGCGGAGGGGAGCCGCGGTG
GACTTCGGGTCGGAGGGACCGATATACCAGGGCTCTGACGCGGGGTAACTCTAGGACGCT
CTCTCGAGGAGGGGAGGGTACGGGCAGCCAGGTGGGCGTGCAGGAACTGGACGGCGTAAT
CCCTGGTGACTGCGATGTGGGGTAGGACCCAGCGCACCCTGATGGGGCCTGGCGGCGGGG
GACCCCTGGACGGACGTGGCAGCTACCCTGCCGTTGAGCCGATGCGGACGTGCCTGGTAC
ACGTGGGTGTTGGGGGAACTCCACAGCGGCAGGCGACGGTTGGGTGGGGCCGGGGGCCCA
CGGGGGAGGCGGGTGGCGGTCACGAGGGGGACACGGGGGAGCGATCCGATGGTCGAGAAC
GATCGGCTAGGAGGCGAGCCCCAGGTGGCGCAGGGCGGGTGCGCCCGTGTCGGCGCGGGG
GGGAGGTGGCGACGAGGCGGCCCTTCAGGCGGGGTGAGAATGGCCCAAGTTATGGCGCCG
CGGTGGAGGGGGGGCGGTGAGGTGGGGGGCCAGTCGGGGTTGGTGAGGGCGGGACCAGCC
TCGCTTGATGCAACGCCAACCCGGATAGTAATTCGTGCGGCTCGTGGTCGCGAGTTCTTC
CAATGATTCAGGGGGGGGTGTGGGGCCCGCCCGTCGACCGGCCGCGTCACGACTGCTACG
GGGTCATTACCGCCGGGCTGTCGTAGGCCGGCGATCGCGGGAGGTGGTTCGAATGCCGCG
GCGGGAGGCCCGGATCAAGCGAAGGGGCGGGGCCCGGGGCACGCGAGGCGGGGGGGGGCG
AGACGGTAGCCGGTGCAGGCGACGCGTGGCCCCATGAGGTAGAACTCTCTCGTGCCCCAA
CACCACACATTAGCGCCCCCATCTCGGCCGACTTCGAGCGCGCTCAGCGAGAACCACCAC
AGGCCTCCAACCCGACGTACTACCCGTCCTCACGAGGCGGGCCGACCCCCGGGCCCAAGC
GAAGGCTCCGGCGCGCACACCCGGCGTGCTGC
>MAP 130_A_tb1283
CCCCTTAGGCGCCTCGAGCGGGTCCCCRCTTCGCCAGTTCACCCGTCCCAGTAAGCGCCC
GCGGCACCTCGCCTTCGCGGAGAACCCCCAGAACGCCCCACCGTCTCCCAGCTCGCACAC
GCGCACGTGTCGCGCTCTCCAGCGCGCGGACTGTTCCAGCCACTGTTCTCATGCTCCCCG
CGGGGCCGTATGACGAGCCACGCCCCAGCCCCGCCGCCGCCGCTCGCCCACCGAGACAAG
CGTAACGCTATTGGGGCCTGACCATGGCTGGGCGGGGCGCTTCCCACGGAGGGCCCAGTC
TTTGGGAGGAGTTGCCCCTCAAGGGATCGGTAGCGCGGTGCGGCTCCCCGCAGCGGCCGC
GAGCGTCGGGGTGATGCTGGAGAGTGGGGCATGCCCCTTCCGGCCGCAGGCGGAACAGGT

TCGCGGTGGTGCACAGGGACGCGGTGCTGCAGCTCGAGACCGCGGAGGGGAGCCGCGGTG
GACTTCGGGTCGGAGGGACCGATATACCAGGGCTCTGACGCGGGGTAACTCTAGGACGCT
CTCTCGAGGAGGGGAGGGTACGGGCAGCCAGGTGGGCGTGCAGGAACTGGACGGCGTAAT
CCCTGGTGACTGCGATGTGGGGTAGGACCCAGCGCACCCTGATGGGGCCTGGCGGCGGGG
GACCCCTGGACGGACGTGGCAGCTACCCTGCCGTTGAGCCGATGCGGACGTGCCTGGTAC
ACGTGGGTGTTGGGGGAACTCCACAGCGGCAGGCGACGGTTGGGTGGGGCCGGGGGCCCA
CGGGGGAGGCGGGTGGCGGTCACGAGGGGGACACGGGGGAGCGATCCGATGGTCGAGAAC
GATCGGCTAGGAGGCGAGCCCCAGGTGGCGCAGGGCGGGTGCGCCCGTGTCGGCGCGGGG
GGGAGGTGGCGACGAGGCGGCCCTTCAGGCGGGGTGAGAATGGCCCAAGTTATGGCGCCG
CGGTGGAGGGGGGGCGGTGAGGTGGGGGGCCAGTCGGGGTTGGTGAGGGCGGGACCAGCC
TCGCTTGATGCAACGCCAACCCGGATAGTAATTCGTGCGGCTCGTGGTCGCGAGTTCTTC
CAATGATTCAGGGGGGGGTGTGGGGCCCGCCCGTCGACCGGCCGCGTCACGACTGCTACG
GGGTCATTACCGCCGGGCTGTCGTAGGCCGGCGATCGCGGGAGGTGGTTCGAATGCCGCG
GCGGGAGGCCCGGATCAAGCGAAGGGGCGGGGCCCGGGGCACGCGAGGCGGGGGGGGGCG
AGACGGTAGCCGGTGCAGGCGACGCGTGGCCCCATGAGGTAGAACTCTCTCGTGCCCCAA
CACCACACATTAGCGCCCCCATCTCGGCCGACTTCGAGCGCGCTCAGCGAGAACCACCAC
AGGCCTCCAACCCGACGTACTACCCGTCCTCACGAGGCGGGCCGACCCCCGGGCCCAAGC
GAAGGCTCCGGCGCGCACACCCGGCGTGCTGC
>MAP 161_A_tb1286
CCCCTTAGGCGCCTCGAGCGGGCCCCCGCTTCGCCAGTTCAGCCGTCCCAGTAAGCGCCC
GGGGCACCTCGCCTTCGAGGAGAACCCCCAGAACGCCCCACCGTCTCCCAGCTCGCACAC
GCGCACGCGTCGCACTCTCCCGCGCGCGTACTGTTCCAGCCACCGTCCCCATGCTCCCCG
CGGGGCCGTATGACGAGCCACGCCCCGGCCCCGCCGCCGCCGCTCGCTCACCGAGACCAG
CGCAATGCTATTGGGGCCTGACCACGGCTGGGCGGGGCGCTTCCCACGGAGGGCCCAGTC
TTTGGGAGGAGTTGCCCCTCCAGGGATCGGCAGCGCGGTGCGGCTCCCCGCAGCGGCCGC
GGGCGTCGGGGGGATGCTAGGGAGTGGGGGATGCCCCTTCCGGTCGCAGGCGGAACAGGT
TCGCGGTGGTGCACAGGGGCGTGGTGCTGCAGCTCGAGGCCGCGGAGGGGGGCCGCGGTG
GACTTCGGGTCGGAGGGACCGATATACCAGGGCTCTGGCGCGGGGTAGCCCCAGGACGCT
TTCTCGGGGAGGGGAGGGTACGGGCAGCCAGGTGGGAGTACAAGAACTGGACGGCCTAAT
CCCTGCTGCCTGCGATGTGGGGTCGGACCCAGCGCACCCTGATGGGGCCTGGCGGCGGGG
GACCCCTGAACGGACGTGGCAGCTACTCTGCCGTTGAGCCGATGCGGACGTGCCCAGTAC
ACGTGGGTGTTGGGGGAACTCCACAGCGGCAGGCGAGGGTTGGGTGTAGCAGGGGGCCCA
CGGGGGCGGCGGGTGGCAGGCGCGAGGGGGGCGCGGGGGAGCGATCCGATGGTCGAGAAC
GATCGGCTAGGAGGCGAGCCCCAAGTGGCGCCGGGCGGGTGCGCCCGTGTCGGCGCGGGG
GGGTTGTGGCGACGAGGCCGCCCTTCAGGCGAGGTGGGAGTGGCCCACGTTGTGGCGCCG
CGGTGGAGGGGGGGCGGTGAGGTGGGGGGCCAGTCGAGGTTGGTGAGGGCGGGACCAGCC
TCGCTTAGTGCAACGCCAACCCGGATAGTAATTCGTGCGGCTCGTGGGAGCGAGTTCTTC
CAATGATGCAGGGGGGGGTGTGGGGCCCGCGCGTCGACCGGCCGCGTCACGACTGCTACG
GGGTCATTACCGCCGGGCTGTCGTAGGCCGGCGCTCGCGGGAGGTGGTTCGAATGCCGGG
GCGGGAGGCCCGGATCCGGCCAAGGGGCGGGGCTCGGGGGACGCGAGGCGGAGGGGCGCG
AGGCGGTAGCCGGTGCAAGCGGCGCGTGGCCCCATGAGGAAGAACTCTCTCGTGCCCCAA
CACCACACATTAGCGCCCCCATCTCGGCCGACTTCGAGCACGCTCAGCGAGAACCACCAC
AGGCCTCCAACCCGACGTACCACCCGTCCTCACGAGGCGGGCCGACCCTCGGGTCCAAGC
GAGGGCTCCGGCGCGCACGCCCGGCGTGGTGC
>MAP 162_A_tb1286
CCCCTTAGGCGCCTCGGGCGGGCCCCCGCCTCGCCAGTTCACCCGTCCCAGTAAGCGCCT
GCGGCCCCTCGCCTTCGCGGAGAACCCCCAGAACGCCCCACCGTATCCCTGCTCGCACAC
GCGCACGCGTCGCACTCTCCAGCGCGCGGACTGTTCCAGCCACCGTCCCCATGTTCCCCG
CGAGGCCGTATGACGAGTCACGCCCCGGCCCCGCCGCCGCCGCTCGCCCACCGAGACAAG
CGCAACGCTATTGGGGCCTGACCACGGCTGGGCGGGGCGCTTCCCACGGAGAACCCAGCC
TTTGGGAGGAGTTGCCCCTCAGGGGACCGGTAGCGCGGTGCGGCTCCCCGCAGCGGCCGC
GGGCGTCGGGGTGATGCTGGGGAGTGGGGGATGTCCCTTCCGGCCGCAGGCGGAACAGGT
TCGCGGTGGTGCACAGGGGCATGGTGCTGCAGCTCAAGGCCGCAGAGGGAGGCCGCGGTG
GACTTCGGGTCGGAGGGACCGATATACCAGGGCTCTGGCGCGGGGTAGCCCCAGGACGCT
CTCCCGGGGAGGGGAGAGTACGGGCAGCCAGGTGGGAGTGCAGGAACTGGACGGCGTAAT
CCCTGGTGACTGCGATGTTGGGGAGGACCCAGCGCACCCTGATGGGGCCTGACGGCGGGG
GATCCCTGGACCGACGTGGCAGCTACCCCGCCGTTGGGCCGATGCGGACGTGTCCAGTAC
ACGTGGGTGTTGGGGGAACTCCACAGCGGCGGGCGAGTGTTGGGTGTGGCCGGGGGCCCA
CAGGGGCGAGGGGTGACGGTCGCGAGGGGGGCGTGGGGGAGCGATCCGATGGTCGAGAAC
GATTAGCTGGGAGGCGAGACCCAGGCGGCGCAGGGCGGGTGCGCCCGTGTCGGCGCGGGG
GGGTTGTGGCGACGAGGCCGCCCTTCAGGCGGGGTGGGAGTGACCCAAGTTGTGGCGCCG
CGGCGGAGGGGGGGCGGTGAGGTGGGGGGCCAGTCGGGGTTGGTGAGGACGGGATCAGCC
TTGCTTGGTGCAACGCCAACCCGGATAGTGATTCGTGCGGCTCGTGGGCGCGGGTTGCTC
CAATGATTCAGGGGGGGGTGTGGGGCCCGCGCGTCGACCGGCCGCGTCACGACTGCTACG
GGGTCATTACCGCCGGGCTGTCGTAGGCCAGCGATCGCGGGAGGTGGTTCGAATGCCGGG
GCCGGAGGCCCGAATCAGGCGAAGGGGCGGGGTCCGCGGCACGCGAGGCGGGGGGGCGCG
AGGCGGTAGCCGGTGCAAGCGGCGCCTGGCCCCATGAGGTAGAACTCTCTCGTCCCCCAA
CACCACACACTAGCGCCTCCATCTCGGCCAACTTCGAGCGCGCTCAGGGAGAACCACCAC

AGGCCTCCAACCCGACGTACCACCCGTCCTCACGAGGCGGGCCGAGCCCCGGGCCCAAGC
GAGGGCTCCGGCGAGCACGCCCGGCGTGGTAC
>MAP 008_A_fb1288
CCCCTTAGGCGCCTCGAGCGGGCCCCCGCTTCGCCAGTTCACCCGTCCCAGTAAGCGCCC
GGGGCACCTCGCCTTCGAGGAGAACCCCCAGAACGCCCCACCGTCTCCCAGCTCGCACAC
GCGCACGCGTCGCACTCTCCAGCGCGCGTACTGTTCCAGCCACCGTCCCCATGCTCCCCG
CGGGGCCGTATGACGAGCCACGCCCCGGCCCCGCCGCCGTCGCTCGCCCGCCGAGACAAG
CGCAATGCTATTGGGGCCTGACCACGGCTGGGCGGGGCGCTTCCCACGGAGGGCCCAGTC
TTTGGGAGGAGTTGCCCCTCCAGGGATCGGTAGCGCGGTGCGGCTCCCCGCAGCGGCCGC
AGGCGTCGGGGGGATGCTGGGGAGTGGGGGATGCCCCTTCCGGTCGCAGGCGGAACAGGT
TCGCGGTGGTGCACAGGGGCGTGGTGCTGCAGCTCGAGGCCGCGGAGGGGGGGCCGCGGTG
GACTTCGGGTCGGAGGGACCGATATACCAGGGCTCTGGCGCGGGGTAGCCCCAGGACGCT
TTCTCGGGGAGGGGAGGGTACGGGCAGCCAGGTGGGAGTGCCAGAACTGGACGGCGTAAT
CCCTGCTGCCTGCGATGTGGGGTCGGACCCAGCGCACCCTGATGGGGCCTGGCGGCGGGG
GACCCCTGAACGGACGTGGCAGCTACTCTGCCGTTGAGCCGATGCGGACGTGCCCAGTAC
ACGTGGGTGTTGGGGGAACTCCACAGCGGCAGGCGAGGGTTGGGTGTAGCAGGGGGCCCA
CGGGGGCGGCGGGTGGCAGGCGCGAGGGGGGCGCGGGGGAGCGATCCGATGGTCGAGAAC
GATCGGCTAGGAGGCGAGCCCCAGGTGGCACCGGGCGGGTGCGCCCGTGTCGGCGCGGGG
GGGTTGTGGCGACGAGGCCGCCCTTCAGGCGAGGTGGGAGTGGCCCAAGTTGTGGCGCCG
CGGTGGAGGGGGGGCGGTGAGGTGGGGGGCCAGTCGAGGTTGGTGAGGGCGGGACCAGCC
TCGTTTAGTGAAACGCCAACCCGGATAGTAATTCGTGCGGCTCGTGGGAGCGAGTTCTTC
CAATGATGCAGGGGGGGGTGTGGGGCCCGCGCGTCGACCGGTCGCGTCACGACTGCTACG
GGGTCATTACCGCCGGGCTGTCGTAGGCCGGCGCTCGCGGGAGGTGGTTCGAATGCCGGG
GCGGGAGGCCCGGATCCGGCGAAGGGGCGGGGCTCGGGGGACGCGAGGCGGAGGGGCGCG
AGGCGGTAGCCGGTGCAAGCGGCGCGTGGCCCCATGAGGAAGAACTCTCTCGTGCCCCAA
CACCACACATTAGCGCCCCCATCTCGGCCGACTTCGAGCACGCTCAACGAGAACCACCAC
AGGCCTCCAACCCGACGTACCACCCGTCCTCACGAGGCGGGCCGACCCTCGGGTCCAAGC
GAGGGCTCCGCCGCGTATGCCCGGCGTGGTGC
>MAP 205_A_fb1291
CCCCTTAGGCGCCTCGAGCGGGCCCCCGCTTCGCCAGTTCACCCGTCCCAGTAAGCGCCC
GGGGCACCTCGCCTTCGAGGAGAACCCCCAGAACGCCCCACCGTCTCCCAGCTCGCACAC
GCGCACGCGTCGCACTCTCCAGCGCGCGTACTGTTCCAGCCACCGTCCCCATGCTCCCCG
CGGGGCCGTATGACGAGCCACGCCCCGGCCCCGCCGCCGTCGCTCGCCCGCCGAGACAAG
CGCAATGCTATTGGGGCCTGACCACGGCTGGGCGGGGCGCTTCCCACGGAGGGCCCAGTC
TTTGGGAGGAGTTGCCCCTCCAGGGATCGGTAGCGCGGTGCGCCTCCCCGCAGCGGCCGC
AGGCGTCGGGGGGATGCTGGGGAGTGGGGGATGCCCCTTCCGGTCGCAAGCGGAACAGGT
TCGCGGTGGTGCACAGAGGCGTGGTGCTGCAGCTCGAGGCCGCGGAGGGGGGGCCGCGGTG
GACTTCGGGTCGGAGGGACCGATATACCAGGGCTCTGGCGCGGGGTAGCCCCAGGACGCT
TTCTCGGGGAGGGGAGGGTACGGGCAGCCAGGTGGGAGTGCCAGAACTGGACGGCGTAAT
CCCTGCTGCCTGCGATGTGGGGTCGGACCCAGCGCACCCTGATGGGGCCTGGCGGCGGGG
GACCCCTGAACGGACGTGGCAGCTACTCTGCCGTTGAGCCGATGCGGACGTGCCCAGTAC
ACGTGGGTGTTGGGGGGACTCCACAGCGGCAGGCGAGGGTTGGGTGTAGCAGGGGGCCCA
CGGGGGCGGCGGGTGGCAGGCGCGAGGGGGGCGCGGGGGAGCGATCCGATGGTCGAGAAC
GATCGGCTAGGAGGCGAGCCCCAGGTGGCACCGGGCGGGTGCGCCCGTGTCGGCGCGGGG
GGGTTGTGGCGACGAGGCCGCCCTTCAGGCGAGGTGGGAGTGGCCCAAGTTGTGGCGCCG
CGGTGGAGGGGGGGCGGTGAGGTGGGGGGCCAGTCGAGGTTGGTGAGGGCGGGACCAGCC
TCGTTTAGTGAAACGCCAACCCGGATAGTAATTTGTGCGGCTCGTGGGAGCGAGTTCTTC
CAATGATGCAGGGGGGGGTGTGGGGCCCGCGCGTCGACCGGTCGCGTCACGACTGCTACG
GGGTCATTACCGCCGGGCTGTCGTAGGCCGGCGCTCGCGGGAGGTGGTTCGAATGCCGGG
GCGGGAGGCCCGGATCCGGCGAAGGGGCGGGGCTCGGGGGACGCGAGGCGGAGGGGCGCG
AGGCGGTAGCCGGTGCAAGCGGCGCGTGGCCCCATGAGGAAGAACTCTCTCGTGCCCCAA
CACCACACATTAGCGCCCCCATCTCGGCCGACTTCGAGCACGCTCAACGAGAACCACCAC
AGGCCTCCAACCCGACGTACCACCCGTCCTCACGAGGCGGGCCGACCCTCGGGTCCAAGC
GAGGGCTCCGCCGCGTATGCCCGGCGTGGTGC
>MAP 204_A_fb1296
CCCCTTAGGCGCCTCGAGCGGGCCCCCGCTTCGCCAGTTCACCCGTCCCAGTAAGCGCCC
GGGGCACCTCGCCTTCGAGGAGAACCCCCAGAACGCCCCACCGTCTCCCAGCTCGCACAC
GCGCACGCGTCGCACTCTCCAGCGCGCGTACTGTTCCAGCCACCGTCCCCATGCTCCCCG
CGGGGCCGTATGACGAGCCACGCCCCGGCCCCGCCGCCGCCGCTCGCCCACCGAGACAAG
CGCAATGCTATTGGGGCCTGACCACGGCTGGGCGGGGCGCTTCCCACGGAGGGCCCAGTC
TTTGGGAGGAGTTGCCCCTCCAGGGATCGGTAGCGCGGTGCGGCTCCCCGCAGCGGCCGC
AGGCGTCGGGGGGATGCTGGGGAGTGGGGGATGCCCCTTCCGGTCGCAGGCGGAACAGGT
TCGCGGTGGTGCACAGGGGCGTGGTGCTGCAGCTCGAGGCCGCGGAGGGGGGGCCGCGGTG
GACTTCGGGTCGGAGGGACCGATATACCAGGGCTCTGGCGCGGGGTAGCCCCAGGACGCT
TTCTCGGGGAGGGGAGGGTACGGGCAGCCAGGTGGGAGTGCCAGAACTGGACGGCGTAAT
CCCTGCTGCCTGCGATGTGGGGTCGGACCCAGCGCACCCTGATGGGGCCTGGCGGCGGGG
GACCCCTGAACGGACGTGGCAGCTACTCTGCCGTTGAGCCGATGCGGACGTGCCCAGTAC
ACGTGGGTGTTGGGGGAACTCCACAGCGGCAGGCGAGGGTTGGGTGTAGCAGGGGGCCCA

CGGGGGCGGCGGGTGGCAGGCGCGAGGGGGGCGCGGGGGAGCGATCCGATGGTCGAGAAC
GATCGGCTAGGAAGCGAGCCCCAGGTGGCACCGGGCGGGTGCGCCCGTGTCGGCGCGGGG
GGGTTGTGGCGACGAGGCCGCCCTTCAGGCGAGGTGGGAGTGGCCCAAGTTGTGGCGCCG
CGGTGGAGGGGGGGCGGTGAGGTGGGGGGCCAGTCGAGGTTGGTGAGGGCGGGACCAGCC
TCGTTTAGTGAAACGCCAACCCGGATAGTAATTCGTGCGGCTCGTGGGAGCGAGTTCTTC
CAATGATGCAGGGGGGGGTGTGGGGCCCGCGCGTCGACCGGTCGCGTCACGACTGCTACG
GGGTCATTACCGCCGGGCTGTCGTAGGCCGGCGCTCGCGGGAGGTGGTTCGAATGCCGGG
GCGGGAGGCCCGGATCCGGCGAAGGGGCGGGGCTCGGGGGACGCGAGGCGGAGGGGCGCG
AGGCGGTAGCCGGTGCAAGCGGCGCGTGGCCCCATGAGGAAGAACTCTCTCGTGCCCCAA
CACCACACATTAGCGCCCCCATCTCGGCCGACTTCGAGCACGCTCAACGAGAACCACCAC
AGGCCTCCAACCCGACGTACCACCCGTCCTCACGAGGCGGGCCGACCGTCGGGTCCAAGC
GAGGGCTCCGCCGCGCACGCCCGGCGTGGTGC
>MAP 187_A_fb1329
CCCCTTAGGCGCCTCGAGCGGGCCCCCGCTTCGCCAGTTCACCCGTCCCAGTAAGCGCCC
GGGGCACCTCGCCTTCCAGGAGAACCCCCAGAACGCCCCACCGTCTCCCAGCTCGCACAC
GCGCACGCGTCGCACTCTCCAGCGCGCGTACTGTTCCAGCCACCGTCCCCATGCTCCCCG
CGGGGCCGTATGACGAGCCACGCCCCGGCCCCGCCGCCGTCGCTCGCCCGCCGAGACAAG
CGCAATGCTATTGGGGCCTGACCACGGCTGGGCGGGGCGCTTCCCACGGAGGGCCCAGTC
TTTGGGAGGAGTTGCCCCTCCAGGGATCGGTAGCGCGGTGCGGCTCCCCGCAGCGGCCGC
AGGCGTCGGGGGGATGCTGGGGAGTGGGGGATGCCCCTTCCGGTCGCAGGCGGAACAGGT
TCGCGGTGGTGCACAGGGGCGTGGTGCTGCAGCTCGAGGCCGCGGAGGGGGGCCGCGGTG
GACTTCGGGTCGGAGGGACCGATATACCAGGGCTCTGGCGCGGGGTAGCCCCAGGACGCT
TTCTCGGGGAGGGGAGGGTACGGGCAGCCAGGTGGGAGTGCCAGAACTGGACGGCGTAAT
CCCTGCTGCCTGCGATGTGGGGTCGGACCCAGCGCACCCTGATGGGGCCTGGCGGCGGGG
GACCCCTGAACGGACGTGGCAGCTACTCTGCCGTTGAGCCGATGCGGACGTGCCCAGTAC
ACGTGGGTGTTGGGGGAACTCCACAGCGGCAGGCGAGGGTTGGGTGTAGCAGGGGGCCCA
CGGGGGCGGCGGGTGGCAGGCGCGAGGGGGGCGCGGGGGAGCGATCCGATGGTCGAGAAC
GATCGGCTAGGAGGCGAGCCCCAGGTGGCACCGGGCGGGTGCGCCCGTGTCGGCGCGGGG
GGGTTGTGGCGACGAGGCCGCCCTTCAGGCGAGGTGGGAGTGGCCCAAGTTGTGGCGCCG
CGGTGAAGGGGGGGCGGTGAGGTGGGGGGCCAGTCGAGGTTGGTGAGGGCGGGACCAGCC
TCGTTTAGTGAAACGCCAACCCGGATAGTAATTCGTGCGGCTCGTGGGAGCGAGTTCTTC
CAATGATGCAGGGGGGGGTGTGGGGCCCGCGCGTCGACCGGTCGCGTCACGACTGCTACG
GGGTCATTACCGCCGGGCTGTCGTAGGCCGGCGCTCGCGGGAGGTGGTTCGAATGCCGGG
GCGGGAGGCCCGGATCCGGCGAAGGGGCGGGGCTCGGGGGACGCGAGGCGGAGGGGCGCG
AGGCGGTAGCCGGTGCAAGCGGCGCGTGGCCCCATGAGGAAGAACTCTCTCGTGCCCCAA
CACCACACATTAGCGCCCCCATCTCGGCCGACTTCGAGCACGCTCAACGAGAACCACCAC
AGGCCTCCAACCCGACGTACCACCCGTCCTCACGAGGCGGGCCGACCCTCGGGTCCAAGC
GAGGGCTCCGCCGCGTATGCCCGGCGTGGTGC
>MAP 113_A_fasb1342
CCCCTTAGGCGCCTCGAGCGGGCCCCCGCTTCGCCAGTTCAGCCGTCCCAGTAAGCGCCC
GGGGCACCTCGCCTTCGAGGAGAACCCCCAGAACGCCCCACCGTCTCCCAGCTCGCACAC
GCGCACGCGTCGCACTCTCCCGCGCGCGTACTGTTCCAGCCACCGTCCCCATGCTCCCCG
CGGGGCCGTATGACGAGCCACGCCCCGGCCCCGCCGCCGCCGCTCGCTCACCGAGACCAG
CGCAATGCTATTGGGGCCTGACCACGGCTGGGCGGGGCGCTTCCCACGGAGGGCCCAGTC
TTTGGGAGGAGTTGCCCCTCCAGGGATCGGCAGCGCGGTGCGGCTCCCCGCAGCGGCCGC
GGGCGTCGGGGGGATGCTAGGGAGTGGGGGATGCCCCTTCCGGTCGCAGGCGGAACAGGT
TCGCGGTGGTGCACAGGGGCGTGGTGCTGCAGCTCGAGGCCGCGGAGGGGGGCCGCGGTG
GACTTCGGGTCGGAGGGACCGATATACCAGGGCTCTGGCGCGGGGTAGCCCCAGGACGCT
TTCTCGGGGAGGGGAGGGTACGGGCAGCCAGGTGGGAGTGCAAGAACTGGACGGCCTAAT
CCCTGCTGCCTGCGATGTGGGGTCGGACCCAGCGCACCCTGATGGGGCCTGGCGGCGGGG
GACCCCTGAACGGACGTGGCAGCTACTCTGCCGTTGAGCCGATGCGGACGTGCCCAGTAC
ACGTGGGTGTTGGGGGAACTCCACAGCGGCAGGCGAGGGTTGGGTGTAGCAGGGGGCCCA
CGGGGGCGGCGGGTGGCAGGCGCGAGGGGGGCGCGGGGGAGCGATCCGATGGTCGAGAAC
GATCGGCTAGGAGGCGAGCCCCAGGTGGCGCCGGGCGGGTGCGCCCGTGTCGGCGCGGGG
GGGTTGTGGCGACGAGGCCGCCCTTCAGGCGAGGTGGGAGTGGCCCACGTTGTGGCGCCG
CGGTGGAGGGGGGGCGGTGAGGTGGGGGGCCAGTCGAGGTTGGTGAGGGCGGGACCAGCC
TCGCTTAGTGCAACGCCAACCCGGATAGTAATTCGTGCGGCTCGTGGGAGCGAGTTCTTC
CAATGATGCAGGGGGGGGTGTGGGGCCCGCGCGTCGACCGGCCGCGTCACGACTGCTACG
GGGTCATTACCGCCGGGCTGTCGTAGGCCGGCGCTCGCGGGAGGTGGTTCGAATGCCGGG
GCGGGAGGCCCGGATCCGGCCAAGGGGCGGGGCTCGGGGGACGCGAGGCGGAGGGGCGCG
AGGCGGTAGCCGGTGCAAGCGGCGCGTGGCCCCATGAGGAAGAACTCTCTCGTGCCCCAA
CACCACACATTAGCGCCCCCATCTCGGCCGACTTCGAGCACGCTCAGCGAGAACCACCAC
AGGCCTCCAACCCGACGTACCACCCGTCCTCACGAGGCGGGCCGACCCTCGGGTCCAAGC
GAGGGCTCCGGCGCGCACGCCCGGCGTGGTGC
>MAP 159_A_tb1355
CCCCTTAGGCGCCTCGAGCGGGCCCCCGCTTCGCCAGTTCACCCGTCCCAGTAAGCGCCC
GGGGCACCTCGCCTTCGAGGAGAACCCCCAGAACGCCCCACCGTCTCCCAGCTCGCACAC
GCGCACGCGTCGCACTCTCCAGCGCGCGTACTGTTCCAGCCACCGTCCCCATGCTCCCCG

CGGGGCCGTATGACGAGCCACGCCCCGGCCCCGCCGCCGTCGCTCGCCCGCCGAGACAAG
CGCAATGCTATTGGGGCCTGACCACGGCTGGGCGGGGCGCTTCCCACGGAGGGCCCAGTC
TTTGGGAGGAGTTGCCCCTCCAGGGATCGGTAGCGCGGTGCGGCTCCCCGCAGCGGCCGC
AGGCGTCGGGGGGATGCTGGGGAGTGGGGGATGCCCCTTCCGGTCGCAGGCGGAACAGGT
TCGCGGTGGTGCACAGGGGCGTGGTGCTGCAGCTCGAGGCCGCGGAGGGGGGCCGCGGTG
GACTTCGGGTCGGAGGGACCGATATACCAGGGCTCTGGCGCGGGGTAGCCCCAGGACGCT
TTCTCGGGGAGGGGAGGGTACGGGCAGCCAGGTGGGAGTGCCAGAACTGGACGGCGTAAT
GCCTGCTGCCTGCGATGTGGGGTCGGACCCAGCGCACCCTGATGGGGCCTGGCGGCGGGG
GACCCCTGAACGGACGTGGCAGCTACTCTGCCGTTGAGCCGATGCGGACGTGCCCAGTAC
ACGTGGGTGTTGGGGGAACTCCACAGCGGCAGGCGAGGGTTGGGTGTAGCAGGGGGCCCA
CGGGGGCGGCGGGTGGCAGGCGCGAGGGGGGCGCGGGGGAGCGATCCGATGGTCGAGAAC
GATCGGCTAGGAGGCGAGCCCCAGGTGGCACCGGGCGGGTGCGCCCGTGTCGGCGCGGGG
GGGTTGTGGCGACGAGGCCGCCCTTCAGGCGAGGTGGGAGTGGCCCAAGTTGTGGCGCCG
CGGTGGAGGGGGGGCGGTGAGGTGGGGGGCCAGTCGAGGTTGGTGAGGGCGGGACCAGCC
TCGTTTAGTGAAACGCCAACCCGGATAGTAATTCGTGCGGCTCGTGGGAGCGAGTTCTTC
CAATGATGCAGGGGCGGGTGTGGGGCCCGCGCGTCGACCGGTCGCGTCACGACTGCTACG
GGGTCATTACCGCCGGGCTGTCGTAGGCCGGCGCTCGCGGGAGGTGGTTCGAATGCCGGG
GCGGGAGGCCCGGATCCGGCGAAGGGGCGGGGCTCGGGGGACGCGAGGCGGAGGGGCGCG
AGGCGGTAGCCGGTGCAAGCGGCGCGTGGCCCCATGAGGAAGAACTCTCTCGTGCCCCAA
CACCACACATTAGCGCCCCCATCTCGGCCGACTTCGAGCACGCTCAACGAGAACTACCAC
AGGCCTCCAACCCGACGTACCACCCGTCCTCACGAGGCGGGCCGACCCTCGGGTCCAAGC
GAGGGCTCCGCCGCGTATGCCCGGCGTGGTGC
>MAP 160_A_tb1355
CCCCTTAGGCGCCTCGAGCGGGCCCCCGCTTCGCCAGTTCACCCGTCCCAGTAAGCGCCC
GGGGCACCTCGCCTTCGAGGAGAACCCCCAGAACGCCCCACCGTCTCCCAGCTCGCACAC
GCGCACGCGTCGCACTCTCCAGCGCGCGTACTGTTCCAGCCACCGTCCCCATGCTCCCCG
CGGGGCCGTATGACGAGCCACGCCCCGGCCCCGCCGCCGCCGCTCGCCCACCGAGACAAG
CGCAATGCTATTGGGGCCTGACCACGGCTGGGCGGGGCGCTTCCCACGGAGGGCCCAGTC
TTTGGGAGGAGTTGCCCCTCCAGGGATCGGTAGCGCGGTGCGGCTCCCCGCAGCGGCCGC
AGGCGTCGGGGGGATGCTGGGGAGTGGGGGATGCCCCTTCCGGTCGCAGGCGGAACAGGT
TCGCGGTGGTGCACAGGGGCGTGGTGCTGCAGCTCGAGGCCGCGGAGGGGGGCCGCGGTG
GACTTCGGGTCGGAGGGACCGATATACCAGGGCTCTGGCGCGGGGTAGCCCCAGGACGCT
TTCTCGGGGAGGGGAGGGTACGGGCAGCCAGGTGGGAGTGCCAGAACTGGACGGCGTAAT
CCCTGCTGCCTGCGATGTGGGGTCGGACCCAGCGCACCCTGATGGGGCCTGGCGGCGGGG
GACCCCTGAACGGACGTGGCAGCTACTCTGCCGTTGAGCCGATGCGGACGTGCCCAGTAC
ACGTGGGTGTTGGGGGAACTCCACAGCGGCAGGCGAGGGTTGGGTGTAGCAGGGGGCCCA
CGGGGGCGGCGGGTGGCAGGCGCGAGGGGGGCGCGGGGGAGCGATCCGATGGTCGAGAAC
GATCGGCTAGGAAGCGAGCCCCAGGTGGCACCGGGCGGGTGCGCCCGTGTCGGCGCGGGG
GGGTTGTGGCGACGAGGCCGCCCTTCAGGCGARGTGGGAGTGGCCCAAGTTGTGGCGCCG
CGGTGGAGGGGGGGCGGTGAGGTGGGGGGCCAGTCGAGGTTGGTGAGGGCGGGACCAGCC
TCGTTTAGTGAAACGCCAACCCGGATAGTAATTCGTGCGGCTCGTGGGAGCGAGYTCTTC
CAATGATGCAGGGGGGGGTGTGGGGCCCGCGCGTCGACCGGTCGCGTCACGACTGCTACG
GGGTCATTACCGCCGGGCTGTCGTAGGCCGGCGCTCGCGGGAGGTGGTTCGAATGCCGGG
GCGGGAGGCCCGGATCCGGCGAAGGGGCGGGGCTCGGGGGACGCGAGGCGGAGGGGCGCG
AGGCGGTAGCCGGTGCAAGCGGCGCGTGGCCCCATGAGGAAGAACTCTCTCGKGCCCCAA
CACCACACATTAGCGCCCCCATCTCGGCCGACTTCGAGCACGCTCAACGAGAACCACCAC
AGGCCTCCAACCCGACGTACCACCCGTCCTCACGAGGCGGGCCGACCGTCGGGTCCAAGC
GAGGGCTCCGCCGCGCACGCCCGGCGTGGTGC
>MAP 186_A_fb1372
CCCCTTAGGCGCCTCGAGCGGGCCCCCGCTTCGCCAGTTCACCCGTCCCAGTAAGCGCCC
GGGGCACCTCGCCTTCGAGGAGAACCCCCAGAACGCCCCACCGTCTCCCAGCTCGCACAC
GCGCACGCGTCGCACTCTCCAGCGCGCGTACTGTTCCAGCCACCGTCCCCATGCTCCCCG
CGGGGCCGTATGACGAGCCACGCCCCGGCCCCGCCGCCGTCGCTCGCCCGCCGAGACAAG
CGCAATGCTATTGGGGCCTGACCACGGCTGGGCGGGGCGCTTCCCACGGAGGGCCCAGTC
TTTGGGAGGAGTTGCCCCTCCAGGGATCGGTAGCGCGGTGCGGCTCCCCGCAGCGGCCGC
AGGCGTCGGGGGGATGCTGGGGAGTGGGGGATGCCCCTTCCGGTCGCAGGCGGAACAGGT
TCGCGGTGGTGCACAGGGGCGTGGTGCTGCAGCTCGAGGCCGCGGAGGGGGGCCGCGGTG
GACTTCGGGTCGGAGGGACCGATATACCAGGGCTCTGGCGCGGGGTAGCCCCAGGACGCT
TTCTCGGGGAGGGGAGGGTACGGGCAGCCAGGTGGGAGTGCCAGAACTGGACGGCGTAAT
CCCTGCTGCCTGCGATGTGGGGTCGGACCCAGCGCACCCTGATGGGGCCTGGCGGCGGGG
GACCCCTGAACGGACGTGGCAGATACTCTGCCGTTGAGCCGATGCGGACGTGCCCAGTAC
ACGTGGGTGTTGGGGGAACTCCACAGCGGCAGGCGAGGGTTGGGTGTAGCAGGGGGCCCA
CGGGGGCGGCGGGTGGCAGGCGCGAGGGGGGCGCGGGGGAGCGATCCGATGGTCGAGAAC
GATCGGCTAGGAGGCGAGCCCCAGGTGGCACCGGGCGGGTGCGCCCGTGTCGGCGCGGGG
GGGTTGTGGCGACGAGGCCGCCCTTCAGGCGAGGTGGGAGTGGCCCAAGTTGTGGCGCCG
CGGTGGAGGGGGGGCGGTGAGGTGGGGGGCCAGTCGAGGTTGGTGAGGGCGGGACCAGCC
TCGTTTAGTGAAACGCCAACCCGGATAGTAATTCGTGCGGCTCGTGGGAGCGAGTTCTTC
CAATGATGCAGGGGGGGGTGTGGGGCCCGCGCGTCGACCGGTCGCGTCACGACTGCTACG

GGGTCATTACCGCCGGGCTGTCCTAGGCCGGCGCTCGCGGGAGGTGGTTCGAATGCCGGG
GCGGGAGGCCCGGATCCGGCGAAGGGGCGGGGCTCGGGGGACGCGAGGCGGAGGGGCGCG
AGGCGGTAGCCGGTGCAAGCGGCGCGTGGCCCCATGAGGAAGAACTCTCTCGTGCCCCAA
CACCACACATTAGCGCCCCCATCTCGGCCGACTTCGAGCACGCTCAACGAGAACCACCAC
AGGCCTCCAACCCGACGTACCACCCGTCCTCACGAGGCGGGCCGACCCTCGGGTCCAAGC
GAGGGCTCCGCCGCGTATGCCCGGCGTGGTGC
>MAP 228_A_tb1377
CCCCTTAGGCGCCTCGAGCGGGCCCCCGCTTCGCCAGTTCAGCCGTCCCAGTAAGCGCCC
GGGGCACCTCGCCTTCGAGGAGAACCCCCAGAACGCCCCACCGTCTCCCAGCTCGCACAC
GCGCACGCGTCGCACTCTCCCGCGCGCGTACTGTTCCAGCCACCGTCCCCATGCTCCCCG
CGGGGCCGTATGACGAGCCACGCCCCGGCCCCGCCGCCGCCGCTCGCTCACCGAGACCAG
CGCAATGCTATTGGGGCCTGACCACGGCTGGGCGGGGCGCTTCCCACGGAGGGCCCAGTC
TTTGGGAGGAGTTGCCCCTCCAGGGATCGGCCGCGCGGTGCGGCTCCCCGCAGCGGCCGC
GGGCGTCGGGGGGATGCTAGGGAGTGGGGGATGCCCCTTCCGGTCGCAGGCGGAACAGGT
TCGCGGTGGTGCACAGGGGCGTGGTGCTGCAGCTCGAGGCCGCGGAGGGGGGCCGCGGTG
GACTTCGGGTCGGAGGGACCGATATACCAGGGCTCTGGCGCGGGGTAGCCCCAGGACGCT
TTCTCGGGGAGGGGAGGGTACGGGCAGCCAGGTGGGAGTGCAAGAACTGGACGGCCTAAT
CCCTGCTGCCTGCGATGTGGGGTCGGACCCAGCGCACCCTGATGGGGCCTGGCGGCGGGG
GACCCCTGAACGGACGTGGCAGCTACTCTGCCGTTGAGCCGATGCGGACGTGCCCAGTAC
ACGTGGGTGTTGGGGGAACTCCACAGCGGCAGGCGAGGGTTGGGTGTAGCAGGGGGCCCA
CGGGGGCGGCGGGTGGCAGGCGCGAGGGGGGCGCGGGGGAGCGATCCGATGGTCGAGAAC
GATCGGCTAGGAGGCGAGCCCCAGGTGGCGCCGGGCGGGTGCGCCCGTGTCGGCGCGGGG
GGGTTGTGGCGACGAGGCCGCCCTTCAGGCGAGGTGGGAGTGGCCCACGTTGTGGCGCCG
CGGTGGAGGGGGGGCGGTGAGGTGGGGGGCCAGTCGAGGTTGGTGAGGGCGGGACCAGCC
TCGCTTAGTGCAACGCCAACCCGGATAGTAATTCGTGCGGCTCGTGGGAGCGAGTTCTTC
CAATGATGCAGGGGGGGGTGTGGGGCCCGCGCGTCGACCGGCCGCGTCACGACTGCTACG
GGGTCATTACCGCCGGGCTGTCGTAGGCCGGCGCTCGCGGGAGGTGGTTCGAATGCCGGG
GCGGGAGGCCCCGATCCGGCCAAGGGGCGGGGCTCGGGGGACGCGAGGCGGAGGGGCGCG
AGGCGGTAGCCGGTGCAAGCGGCGCGTGGCCCCATGAGGAAGAACTCTCTCGTGCCCCAA
CACCACACATTAGCGCCCCCATCTCGGCCGACTTCGAGCACGCTCAGCGAGAACCACCAC
AGGCCTCCAACCCGACGTACCACCCGTCCTCACGAGGCGGGCCGACCCTCGGGTCCAAGC
GAGGGCTCCGGCGCGCACGCCCGGCGTGGTGC
>MAP 226_A_tb1387
CCCCTTAGGCGCCTCGAGCGGGCCCCCGCTTCGCCAGTTCACCCGTCCCAGTAAGCGCCC
GGGGCACCTCGCCTTCGAGGAGAACCCCCAGAACGCCCCACCGTCTCCCAGCTCGCACAC
GCGCACGCGTCGCACTCTCCAGCGCGCGTACTGTTCCAGCCACCGTCCCCATGCTCCCCG
CGGGGCCGTATGACGAGCCACGCCCCGGCCCCGCCGCCGCCGCTCGCCCACCGAGACAAG
CGCAATGCTATTGGGGCCTGACCACGGCTGGGCGGGGCGCTTCCCACGGAGGGCCCAGTC
TTTGGGAGGAGTTGCCCCTCCAGGGATCGGTAGCGCGGTGCGGCTCCCCGCAGCGGCCGC
AGGCGTCGGGGGGATGCTGGGGAGTGGGGGATGCCCCTGCCGGTCGCAGGCGGAACAGGT
TCGCGGTGGTGCACAGGGGCGTAGTGCTGCAGCTCGAGGCCGCGGAGGGGGGCCGCGGTG
GACTTCGGGTCGGAGGGACCGGGATACCAGGGCTCTGGCGCGGGGTAGCCCCAGGACGCT
TTCTCGGGGAGGGGAGGGTACGGGCAGCCAGGTGGGAGTGCCAGAACTGGACGGCGTAAT
CCCTGCTGCCTGCGATGTGGGGTCGGACCCAGCGCACCCTGATGGGGCCTGGCGGCGGGG
GACCCCTGAACGGACGTGGCAGCTACTCTGCCGTTGAGCCGATGCGGACGTGCCCAGTAC
ACGTGGGTGTTGGGGGAACTCCACAGCGGCAGGCGAGGGTTGGGTGTAGCAGGGGGCCCA
CGGGGGCGGCGGGTGGCAGGCGCGAGGGGGGCGCGGGGGAGCGATCCGATGGTCGAGAAC
GATCGGCTAGGAGGCGAGCCCCAGGTGGCACCGGGCGGGTGCGCCCGTGTCGGCGCGGGG
GGGTTGTGGCGACGAGGCCGCCCTTCAGGCGAGGTGGGAGTGGCCCAAGTTGTGGCGCCG
CGGTGGAGGGGGGGCGGTGAGGTGGGGGGCCAGTCGAGGTTGGTGAGGGCGGGACCAGCC
TCGTTTAGTGAAACGCCAACCCGGATAGTAATTCGTGCGGCTCGTGGGAGCGAGTTCTTC
CAATGATGCAGGGGGGGGTGTGGGGCCCGCGCGTCGACCGGTCGCGTCACGACTGCTACG
GGGTCATTACCGCCGGGCTGTCGTAGGCCGGCGCTCGCGGAGGGAGGTTCGAATGCCGGG
GCGGGAGGCCCGGATCCGGCGAAGGGGCGGGGCTCGGGGGACGCGAGGCGGAGGGGCGCG
AGGCGGTAGCCGGTGCAAGCGGCGCGTGGCCCCATGAGGAAGAACTCTCTCGTGCCCCAA
CACCACACATTAGCGCCCCCATCTCGGCCGACTTCGAGCACGCTCAACGAGAACCACCAC
AGGCCTCCAACCCGACGTACCACCCGTCCTCACGAGGCGGGCCGACCCTCGGGTCCAAGC
GAGGGCTCCGCCGCGCACGCCCGGCGTGGTGC
>MAP 227_A_tb1387
CCCCTTAGGCGCCTCGAGCGGGTCCCCGCTTCGCCAGTTCACCCGTCCCAGTAAGCGCCC
GCGGCACCTCGCCTTTGCGGAGAACCCCCAGAACGCCCCACCGTCTCCCAGCTCGCACAC
GCGCACGTGTCGCACTCTCCAGCACGCGGACTGTTCCAGCCACCGTCCTCATGCTCCCCG
CGGGGCCGTATGACGAGCCACGCCCCGGCCCCGCCGCCGCCGCTCGCCCACCGAGACAAG
CGTAACGCTATTGGGGCCTGACTATGGCTGGGCGGGGCGCTTCCCACGGAGGGCCCAGTC
TTTGGGAGGAGTTGCCCCTCAAGGGATCGGTAGCGCGGTGCGGCTCCCCGCAGCGGCCGC
GAGCGTCGGGGTGATGCTGGAGAGTGGGGCATGCTTGTTCCGGCCGCAGGCGGAACAGGT
TCGCGGTGGTGCACAGGGACGCGGTGCTGCAGCTCGAGGCCGCGGAGGGGAGCCGCGGTG
GACCTCGGGTCGGAGGGACCGATATACCAGGGCTCTGACGCGGGGTAACTCTAGGACGCT

CTCTCGAGGAGGGGAGGGTACGAGCAGCCAGGTGGGCGTGCAGGAACTGGACGGCGTAAT
CTCTGGTGACTGCGATGTGGGGTAGGACCCAGCGCACCCTGATGGGGCTTGGCGGCAGGG
GACCCCTGGACGGACGTGGCAGCTACCCTGCCGTTGAGCCGATGCGGACGTGCCCAGTAC
ACGTGGGTGTTGGGGGAACTCCACAGCGGCAGGCGACGGTTGGGTGGGGCCGGGGGCCCA
CGGGGGCGGCGGGTGGCGGTCACGAGGGGGGCACGGGGGAGCGATCCGATGGTTGAGAAC
GATCGGCTAGGAGGCGAGCCCCAGGTGGTGCAGGGCGGGTGCGCTCGTGTCGGCGCGGGG
GGGAGGTGGCGACGAGGCGGCCCTTCAGGCGGGGTGAGAGTGGCCCAAGTTGTGGCGCCG
CGGTGGAGGGGGGGCGGTGAGGTGGGGGGCCAGTCGGGGTTGGTGAGGGCGGGACCAGCC
TCGCTTGATGCAACGCCAACCCGGATAGTAATTCGTGCGGCTCGTGGTCGCGAGTTCTTC
CAATGATTCAGGGGGGGGTGTGGGGCCCGCCCGTCGACCGGCCGCGTCACGACTGCTACG
GGGTCATTACCGCCGGGTTGTCGTAGGCCGGCGATCGCGGGAGGTGGTTCGAATGCCGCG
GCGGGAGGCCCGGATCAAGCGAAGGGGCGGGGCCCGGGGCACGCGAGGCGGGGGGGGGCA
AGGCGGTAGCCGGTGCAAGCGACGCGTGGCCCCATGAGGTAGAACTCTCTCGTGCCCCAA
CACCACACATTAGCGTCCCCATCTCGGCTGACTTCGAGCGCGCTCAGCGAGAACCACCAT
AGGCCTCCAACCCGACGTACCACCCGCCCTCACGAGGCGGGCCGACCCCCGGGCCCAAGC
GAGGGCTCCGGCGCGCACGCCCGGCGTGCTGC
>MAP 137_A_tb1405
CCCCTTAGGCGCCTCGAGCGGGCCCCCGCTTCGCCAGTTCAGCCGTCCCAGTAAGCGCCC
GGGGCACCTCGCCTTCGAGGAGAACCCCCAGAACGCCCCACCGTCTCCCAGCTCGCACAC
GCGCACGCGTCGCACTCTCCCGCGCGCGTACTGTTCCAGCCACCGTCCCCATGCTCCCCG
CGGGGCCGTATGACGAGCCACGCCCCGGCCCCGCCGCCGCCGCTCGCTCACCGAGACCAG
CGCAATGCTATTGGGGCCTGACCACGGCTGGGCGGGGCGCTTCCCACGGAGGGCCCAGTC
TTTGGGAGGAGTTGCCCCTCCAGGGATCGGCAGCGCGGTGCGGCTCCCCGCAGCGGCCGC
GGGCGTCGGGGGGATGCTAGGGAGTGGGGGATGCCCCTTCCGGTCGCAGGCGGAACAGGT
TCGCGGTGGTGCACAGGGGCGTGGTGCTGCAGCTCGAGGCCGCGGAGAGGGGCCGCGGTG
GACTTCGGGTCGGAGGGACCGATATACCAGGGCTCTGGCGCGGGGTAGCCCCAGGACGCT
TTCTCGGGGAGGGGAGGGTACGGGCAGCCAGGTGGGAGTGCAAGAACTGGACGGCCTAAT
CCCTGCTGCCTGCGATGTGGGGTCGGACCCAGCGCACCCTGATGGGGCCTGGCGGCGGGG
GACCCCTGAACGGACGTGGCAGCTACTCTGCCGTTGAGCCGATGCGGACGTGCCCAGTAC
ACGTGGGTGTTGGGGGAACTCCACAGCGGCAGGCGAGGGTTGGGTGTAGCAGGGGGCCCA
CGGGGGCGGCGGGTGGCAGGCGCGAGGGGGGCGCGGGGGAGCGATCCGATGGTCGAGAAC
GATCGGCTAGGAGGCGAGCCCCAGGTGGCGCCGGGCGGGTGCGTCCGTGTCGGCGCGGGG
GGGTTGTGGCGACGAGGCCGCCCTTCAGGCGAGGTGGGAGTGGCCCACGTTGTGGCGCCG
CGGTGGAGGGGGGGCGGTGAGGTGGGGGGCCAGTCGAGGTTGGTGAGGGCGGGACCAGCC
TCGCTTAGTGCAACGCCAACCCGGATAGTAATTCGTGCGGCTCGTGGGAGCGAGTTCTTC
CAATGATGCAGGGGGGGGTGTGGGGCCCGCGCGTCGACCGGCCGCGTCACGACTGCTACG
GGGTCATTACCGCCGGGCTGTCGTAGGCCGGCGCTCGCGGGAGGTGGTTCGAATGCCGGG
GCGGGAGGCCCGGATCCGGCCAAGGGGCGGGGCTCGGGGGACGCGAGGCGGAGGGGCGCG
AGGCGGTAGCCGGTGCAAGCGGCGCGTGGCCCCATGAGGAAGAACTCTCTCGTGCCCCAA
CACCACACATTAGCGCCCCCATCTCGGCCGACTTCGAGCACGCTCAGCGAGAACCACCAC
AGGCCTCCAACCCGACGTACCACCCGTCCTCACGAGGCGGGCCGACCCTCGGGTCCAAGC
GAGGGCTCCGGCGCGCACGCCCGGCGTGGTGC
>MAP 225_A_tb1415
CCCCTTAGGCGTCTCGAGCGGATCCCCGTTTCGCCAGTTCACCCGTCCCAGTAAGCGCCC
GCGGCACCTCGCCTTCGCGGAGAACCCCCAGAACGCCCCACCGCCTCCCAGCTCGCACAC
GCGCACGTGTCGCACTCTCCAGGGCGCGGACTGTTCCAGCCACCGTTCTCATGCTCCCCG
CGGGGCCGTATGACGAGCCACGCCCCAGCCCCGCCGCCGCCGTTCGCCCACCGAGACAAG
CGTAACGCTATTGGGGCCTGACCATGGCTGGGCGGGGCGCTTCCCACGGAGGGCCCAGTC
TTTGGGAGGAGTTGCCCCTCAAGGGATCGGTAGCGCGGTGCGGCTCCCCGCAGCGGCCGC
GAGCGTCGGGGTGATGCTGGAGAGTGGGGCATGCCCCTTCCGGCCGCAGGCGGAACAGGT
TCGCGGTGGTGCACAGGGACGCGGTGCTGCAGCTCGAGGCCGCGGAGGGGAGCCGCGGTG
GACTTCGGGTCGGAGGGACCGATATACCAGGGCTCTAACGCGGGGTAACTCTAGGACGCT
CTCTCGAGGAGGGGAGGGTACGGGCAGCCAGGTGGGCGTGCAGGAACTGGACGGCGTAAT
CCCTGGTGACTACGATGTGGGGTAGGACCCAGCGCACCTTGATGGGGCCTGGCGACGGGG
AACCCCTGGACGGACGTGGCAGCTACCCTGCCATTGAGCCGATGCGGACGTGCCCAGTAC
ACGTGGGTGCTGGGGGAACTCCACAGCGGCAGGCGACGGTTGGGTGGGGCCGGGAGCCCA
CGGGGGCGGCGGGTGGCGGTCACGAGAGGGGCACGGGGGAGCGATCCGATGGTCGAGAAC
GATCGGCTAGGAGGCGAGCCCCAGGTGGCGCAGGGCGGGTGCGCCCGTGTCGGCGCGGGG
GGGAGGTGGCGACGAGGCGGCCCTTCAGGCGGGGTGAGAGTGGCCCAAGTTGTGGCGCCG
CGGTGGAGGGGGGGCGGTGAGGTGGGGGGCCAGTCGGGGTTGGTGAGGGCGGGACCAGCC
TCGCTTGATGCAACGCCAACCCGGATAGTAATTCGTGCGGCTCGTGGTCGCGAGTTCTTC
CAATGATTCAGGGGGGGGTGTGGGACCCGCCCGTCGACCGGCCGCGTCACGACTGCTACG
GGGTCATTACCGCCGGGCTGTCGTAGGCCGGCGATCGCGGGAGGTGGTTCAAATGCCGCG
GCGGGAGGCCCGGATCAAGCGAAGGGGCGGGGCCCGGGGCACGCGAGGCGGGGGGGGGCG
AGGCGGTAGCCGGTGCAAGCGACGCGTGGCCCCATGAGGTAGAACTCTCTCGTGCCTCAA
CACCACACATTAGCGCCCCCATCTCGGCCGACTTCGAGCGCGCTCAGCGAGAACCACCAC
AGGCCTCCAACCCGACGTACCACCCGTCCTCACGAGGCGGGCCGACCCCCGGGCCCAAGC
GAGGGCTCCGGCGCGCACACCCGGCGTGCTGC

>MAP 094_A_tb1427
CCCCTTAGGCGCCTCGAGCGGGCCCCCGCCTCGCCAGTTCACCCGTCCCAGTAAGCGCCT
GCGGCCCCTCGCCTTCGCGGAGAACCCCCAGAACGCCCCACCGTATCCCTGCTCGCACAC
GCGCACGCGTCGCACTCTCCAGCGCGCGGACTGTCCCGCCCACCGTCCCCATCCTCCCCG
CGGGGCCGTATGACGAGCCACGCCCCGGCCCCGCCGCCGCCGCTCGCCCACCGAGACAAG
CGCAACGCTATTAGGGCCTGACCGCGGCTGGGCGGGGCGCTTCCCACGGAGAACCCAGCC
TTTGGAAGGAATTGCCCCTCAGGGGGTCGGTAGCGCGGTGCGGCTCCCCGCGGCGGCCGC
GGGCGTCGGGGTGATGCTGGGGAGTGGGGGATGCCCCTTCCGGCCGCAGGCGGAACACGT
TCGCGGTGGTGCACAGGGGCGTGGTGCTACAGCTCGAGGCCGCAGAGGGGGGCCGCGGTG
GACTTCGGGTCCGAGGGACCGATATACCAGGGCTCTGGCGCGGGGTAGCCCCAGGACGCT
CTCCCGGGGAGGGGAGGGTACGGGCAGCCAGGTGGGAGTGCAGGAACTGGACGGCGTAAT
CCCTGGTGACTGCGATGTTGAGGAGGACCCAGCGCACCCTGATGGGGCCTGACGGCGGGG
GATCCCTGGACCGACGTGGCAGCTACCCTACCGTTGGGCCGATGCGGACGCGCCCAGTAC
ACGTGGGTGTTGGGGGAACGCCACAGCGGCGGGCGAGTGTTGGGTGTGGCCGGGGGCCCA
CAGGGGCGAGGGGTGACGGTCGCGAGGGGGGCGTGGGGGAGCGATCCGATGGTCGAGAAC
GATCGGCTAGGAGGCGAGACCCAGGCGGCGCAGGGCGGGTGCGCCCGTGTCGGCGCGGGG
GGGTTGTGGCGACGAGGCCGCCCTTCAGGCGGGGTGGGAGTGGCCCAAGCTGCGGAGCCG
CGGCGGAGGGGGGGCGGTGAGGTGGGGGGCCAGTCGGGGTTGGTGAGGACGGGACCAGCC
TTGCTCGGTGCAACGCCAGCCCGGATAGTAATTCGTGCGGCTCGTGGGCGCGGGTTCTTC
CAATGATTCAGGGGGGCGTGTGGGGCCCGCGCGTCGACCGGCCGCGTCACGACTGCTACG
GGGTCATTACCGACGGGCTGTCGTAGGCCGGCGATCGCGGGAGGTGGTTCGAATGCCGGG
GCCGGAGGCCCGAATCAGGCGAAGGGGCGGGGTCCGGGGCACGCGAGGCGGGGGGGCGCG
AGGCGGTGGCCGGTGCAAGCGGCGCCTGGCCCCATGAGGTAGAACTCTCTCGTCCCCCAA
CACCACACACTAGCGCCTCCATCTCGGCCGACTTCGAGCGCGCTCAGGGAGAATCACCAC
AGGCCTCCAACCCGACGTACCACCCGTCCCCACGAGGCGGGCCGACCCCCGGGCCCAAGC
GAGGGCTCCGGCGAGCACGCCCGGCGAGGTAC
>MAP 119_A_tb1429
CCCCTTAGGCGCCTCGAGCGGGCCCCCGCTTCGCCAGTTCACCCGTCCCAGTAAGCGCCC
GGGGCACCTCGCCTTCGAGGAGAACCCCCAGAACGCCCCACCGTCTCCCAGCTCGCACAC
GCGCACGCGTCGCACTCTCCAGCGCGCGTACTGTTCCAGCCACCGTCCCCATGCTCCCCG
CGGGGCCGTATGACGAGCCACGCCCCGGCCCCGCCGCCGCCGCTCGCCCACCGAGACAAG
CGCAATGCTATTGGGGCCTGACCACGGCTGGGCGGGGCGCTTCCCACGGAGGGCCCAGTC
TTTGGGAGGAGTTGCCCCTCCAGGGATCGGTAGCGCGGTGCGGCTCCCCGCAGCGGCCGC
AGGCGTCGGGGGGATGCTGGGGAGTGGGGGATGCCCCTGCCGGTCGCAGGCGGAACAGGT
TCGCGGTGGTGCACAGGGGCGTGGTGCTGCAGCTCGAGGCCGCGGAGGGGGGCCGCGGTG
GACTTCGGGTCGGAGGGACCGGGATACCAGGGCTCTGGCGCGGGGTAGCCCCAGGACGCT
TTCTCGGGGAGGGGAGGGTACGGGCAGCCAGGTGGGAGTGCCAGAACTGGACGGCGTAAT
CCCTGCTGCCTGCGATGTGGGGTCGGACCCAGCGCACCCTGATGGGGCCTGGCGGCGGGG
GACCCCTGAACGGACGTGGCAGCCACTCTGCCGTTGAGCCGATGCGGACGTGCCCAGTAC
ACGGGGGTGTTGGGGGAACTCCACAGCGGCAGGCGAGGGTTGGGTGTAGCAGGGGGCCCA
CGGGGGCGGCGGGTGGCAGGCGCGAGGGGGGCGCGGGGGAGCGATCCGATGGTCGAGAAC
GATCGGCTAGGAGGCGAGCCCCAGGTGGCACCGGGCGGGTGCGCCCGTGTCGGCGCGGGG
GGGTTGTGGCGACGAGGCCGCCCTTCAGGCGAGGTGGGAGTGGCCCAAGTTGTGGCGCCG
CGGTGGAGGGGGGGCGGTGAGGTGGGGGGCCAGTCGAGGTTGGTGAGGGCGGGACCAGCC
TCGTTTAGTGAAACGCCAACCCGGATAGTAATTCGTGCGGCTCGTGGGAGCGAGTTCTTC
CAATGATGCAGGGGGGGGTGTGGGGCCCGCGCGTCGACCGGTCGCGTCACGACTGCTACG
GGGTCATTACCGCCGGGCTGTCGTAGGCCGGCGCTCGCGGAGGGAGGTTCGAATGCCGGG
GCGGGAGGCCCGGATCCGGCGAAGGGGCGGGGCTCGGGGGACACGAGGCGGAGGGGCGCG
AGGCGGTAGCCGGTGCAAGCGGCGCGTGGCCCCATGAGGAAGAACTCTCTCGTGCCCCAA
CACCACACATTAGCGCCCCCATCTCGGCCGACTTCGAGCACGCTCAACGAGAACCACCAC
AGGCCTCCAACCCGACGTACCACCCGTCCTCACGAGGCGGGCCGACCCTCGGGTCCAAGC
GAGGGCTCCGCCGCGCACGCCCGGCGTGGTGC
>MAP 214_A_tb1445
CCCCTTAGGCGCCTCGAGCGGGCCCCCGCTTCGCCAGTTCAGCCGTCCCAGTAAGCGYCC
GGGGCACCTCGCCTTCGAGGAGAACCCCCAGAACGCCCCACCGTCTCCCAGCTCGCACAC
GCGCACGCGTCGCACTCTCCCGCGCGCGTACTGTTCCAGCCACCGTCCCCATGCTCCCCG
CGGGGCCGTATGACGAGCCACGCCCCGGCCCCGCCGCCGCCGCTCGCTCACCGAGACCAG
CGCAATGCTATTGGGGCCTGACCACGGCTGGGCGGGGCGCTTCCCACGGAGGGCCCAGTC
TTTGGGAGGAGTTGCCCCTCCAGGGATCGGCAGCGCGGTGCGGCTCCCCGCAGCGGCCGC
GGGCGTCGGGGGGATGCTAGGGAGTGGGGGATGCCCCTTCCGGTCGCAGGCGGAACAGGT
TCGCGGTGGTGCACAGGGGCGTGGTGCTGCAGCTCGAGGCCGCGGAGGGGGGCCGCGGTG
GACTTCGGGTCGGAGGGACCGATATACCAGGGCTCTGGCGCGGGGTAGCCCCAGGACGCT
TTCTCGGGGAGGGGAGGGTACGGGCAGCCAGGTGGGAGTGCAAGAACTGGACGGCCTAAT
CCCTGCTGCCTGCGATGTGGGGTCGGACCYAGCGCACCCTGATGGGGCCTGGCGGCGGGG
GACCCCTGAACGGACGTGGCAGCTACTCTGCCGTTGAGCCGATGCGGACGTGCCCAGTAC
ACGTGGGTGTTGGGGGAACTCCACAGCGGCAGGCGAGGGTTGGGTGTAGCAGGGGGCCCA
CGGGGGCGGCGGGTGGCAGGCGCGAGGGGGGCGCGGGGGAGCGATCCGATGGTCGAGAAC
GATCGGCTAGGAGGCGAGCCCCAGGTGGCGCCGGGCGGGTGCGCCCGTGTCGGCGCGGGG

GGGTTGTGGCGACGAGGCCGCCCTTCAGGCGAGGTGGGAGTGGCCCACGTTGTGGCGCCG
CGGTGGAGGGGGGGCGGTGAGGTGGGGGGCCAGTCGAGGTTGGTGAGGGCGGGACCAGCC
TCGCTTAGTGCAACGCCAACCCGGATAGTAATTCGTGCGGCTCGTGGGAGCGAGTTCTTC
CAATGATGCAGGGGGGGGTGTGGGGCCCGCGCGTCGACCGGCCGCGTCACGACTGCTACG
GGGTCATTACCGCCGGGCTGTCGTAGGCCGGCGCTCGCGGGAGGTGGTTCGAATGCCGGG
GCGGGAGGCCCGGATCCGGCCAAGGGGCGGGGCTCGGGGGACGCGAGGCGGAGGGGCGCG
AGGCGGTAGCCGGTGCAAGCGGCGCGTGGCCCCATGAGGAAGAACTCTCTCGTGCCCCAA
CACCACACATTAGCGCCCCCATCTCGGCCGACTTCGAGCACGCTCAGCGAGAACCACCAC
AGGCCTCCAACCCGACGTACCACCCGTCCTCACGAGGCGGGCCGACCCTCGGGTCCAAGC
GAGGGCTCCGGCGCGCACGCCCGGCGTGGTGC
>MAP 171_A_fb1491
CCCCTTAGGCGCCTCGAGCGGGCCCCCGCTTCGCCAGTTCAGCCGTCCCAGTAAGCGCCC
GGGGCACCTCGCCTTCGAGGAGAACCCCCAGAACGCCCCACCGTCTCCCAGCTCGCACAC
GCGCACGCGTCGCACTCTCCCGCGCGCGTACTGTTCCAGCCACCGTCCCCATGCTCCCCG
CGGGGCCGTATGACGAGCCACGCCCCGGCCCCGCCGCCGCCGCTCGCTCACCGAGACCAG
CGCAATGCTATTGGGGCCTGACCACGGCTGGGCGGGGCGCTTCCCACGGAGGGCCCAGTC
TTTGGGAGGAGTTGCCCCTCCAGGGATCGGCAGCGCGGTGCGGCTCCCCGCAGCGGCCGC
GGGCGTCGGGGGGATGCTAGGGAGTGGGGGATGCCCCTTCCGGTCGCAGGCGGAACAGGT
TCGCGGTGGTGCACAGGGGCGTGGTGCTGCAGCTCGAGGCCGCGGAGGGGGGCCGCGGTG
GACTTCGGGTCGGAGGGACCGATATACCAGGGCTCTGGCGCGGGGTAGCCCCAGGACGCT
TTCTCGGGGAGGGGAGGGTACGGGCAGCCAGGTGGGAGTGCAAGAACTGGACGGCCTAAT
CCCTGCTGCCTGCGATGTGGGGTCGGACCCAGCGCACCCTGATGGGGCCTGGCGGCGGGG
GACCCCTGAACGGACGTGGCAGCTACTCTGCCGTTGAGCCGATGCGGACGTGCCCAGTAC
ACGTGGGTGTTGGGGGAACTCCACAGCGGCAGGCGAGGGTTGGGTGTAGCAGGGGGCCCA
CGGGGGCGGCGGGTGGCAGGCGCGAGGGGGGCGCGGGGGAGCGATCCGATGGTCGAGAAC
GATCGGCTAGGAGGCGAGCCCCAGGTGGCGCCGGGCGGGTGCGCCCGTGTCGGCGCGGGG
GGGTTGTGGCGACGAGGCCGCCCTTCAGGCGAGGTGGGAGTGGCCCACGTTGTGGCGCCG
CGGTGGAGGGGGGGCGGTGAGGTGGGGGGCCAGTCGAGGTTGGTGAGGGCGGGACCAGCC
TCGCTTAGTGCAACGCCAACCCGGATAGTAATTCGTGCGGCTCGTGGGAGCGAGTTCTTC
CAATGATGCAGGGGGGGGTGTGGGGCCCGCGCGTCGACCGGCCGCGTCACGACTGCTACG
GGGTCATTACCGCCGGGCTGTCGTAGGCCGGCGCTCGCGGGAGGTGGTTCGAATGCCGGG
GCGGGAGGCCCGGATCCGGCCAAGGGGCGGGGCTCGGGGGACGCGAGGCGGAGGGGCGCG
AGGCGGTAGCCGGTGCAAGCGGCGCGTGGCCCCATGAGGAAGAACTCTCTCGTGCCCCAA
CACCACACATTAGCGCCCCCATCTCGGCCGACTTCGAGCACGCTCAGCGAGAACCACCAC
AGGCCTCCAACCCGACGTACCACCCGTCCTCACGAGGCGGGCCGACCCTCGGGTCCAAGC
GAGGGCTCCGGCGCGCACGCCCGGCGTGGTGC
>MAP 194_A_fb1494
CCCCTTAGGCGCCTCGAGCGGGCCCCCGCTTCGCCAGTTCAGCCGTCCCAGTAAGCGCCC
GGGGCACCTCGCCTTCGAGGAGAACCCCCAGAACGCCCCACCGTCTCCCAGCTCGCACAC
GCGCACGCGTCGCACTCTCCCGCGCGCGTACTGTTCCAGCCACCGTCCCCATGCTCCCCG
CGGGGCCGTATGACGAGCCACGCCCCGGCCCCGCCGCCGCCGCTCGCTCACCGAGACCAG
CGCAATGCTATTGGGGCCTGACCACGGCTGGGCGGGGCGCTTCCCACGGAGGGCCCAGTC
TTTGGGAGGAGTTGCCCCTCCAGGGATCGGCAGCGCGGTGCGGCTCCCCGCAGCGGCCGC
GGGCGTCGGGGGGATGCTAGGGAGTGGGGGATGCCCCTTCCGGTCGCAGGCGGAACAGGT
TCGCGGTGGTGCACAGGGGCGTGGTGCTGCAGCTCGAGGCCGCGGAGGGGGGCCGCGGTG
GACTTCGGGTCGGAGGGACCGATATACCAGGGCTCTGGCGCGGGGTAGCCCCAGGACGCT
TTCTCGGGGAGGGGAGGGTACGGGCAGCCAGGTGGGAGTGCAAGAACTGGACGGCCTAAT
CCCTGCTGCCTGCGATGTGGGGTCGGACCCAGCGCACCCTGATGGGGCCTGGCGGCGGGG
GACCCCTGAACGGACGTGGCAGCTACTCTGCCGTTGAGCCGATGCGGACGTGCCCAGTAC
ACGTGGGTGTTGGGGGAACTCCACAGCGGCAGGCGAGGGTTGGGTGTAGCAGGGGGCCCA
CGGGGGCGGCGGGTGGCAGGCGCGAGGGGGGCGCGGGGGAGCGATCCGATGGTCGAGAAC
GATCGGCTAGGAGGCGAGCCCCAGGTGGCGCCGGGCGGGTGCGCCCGTGTCGGCGCGGGG
GGGTTGTGGCGACGAGGCCGCCCTTCAGGCGAGGTGGGAGTGGCCCACGTTGTGGCGCCG
CGGTGGAGGGGGGGCGGTGAGGTGGGGGGCCAGTCGAGGTTGGTGAGGGCGGGACCAGCC
TCGCTTAGTGCAACGCCAACCCGGATAGTAATTCGTGCGGCTCGTGGGAGCGAGTTCTTC
CAATGATGCAGGGGGGGGTGTGGGGCCCGCGCGTCGACCGGCCGCGTCACGACTGCTACG
GGGTCATTACCGCCGGGCTGTCGTAGGCCGGCGCTCGCGGGAGGTGGTTCGAATGCCGGG
GCGGGAGGCCCGGATCCGGCCAAGGGGCGGGGCTCGGGGGACGCGAGGCGGAGGGGCGCG
AGGCGGTAGCCGGTGCAAGCGGCGCGTGGCCCCATGAGGAAGAACTCTCTCGTGCCCCAA
CACCACACATTAGCGCCCCCATCTCGGCCGACTTCGAGCACGCTCAGCGAGAACCACCAC
AGGCCTCCAACCCGGCGTACCACCCGTCCTCACGAGGCGGGCCGACCCTCGGGTCCAAGC
GAGGGCTCCGGCGCGCACGCCCGGCGTGGTGC
>MAP 237_A_fasb1494
CCCCTTAGGCGCCTCGAGCGGGCCCCCGCTTCGCCAGTTCAGCCGTCCCAGTAAGCGCCC
GGGGCACCTCGCCTTCGAGGAGAACCCCCAGAACGCCCCACCGTCTCCCAGCTCGCACAC
GCGCACGCGTCGCACTCTCCCGCGCGCGTACTGTTCCAGCCACCGTCCCCATGCTCCCCG
CGGGGCCGTATGACGAGCCACGCCCCGGCCCCGCCGCCGCCGCTCGCTCACCGAGACCAG
CGCAATGCTATTGGGGCCTGACCACGGCTGGGCGGGGCGCTTCCCACGGAGGGCCCAGTC

```
TTTGGGAGGAGTTGCCCCTCCAGGGATCGGCAGCGCGGTGCGGCTCCCCGCAGCGGCCGC
GGGCGTCGGGGGGATGCTAGGGAGTGGGGGATGCCCCTTCCGGTCGCAGGCGGAACAGGT
TCGCGGTGGTGCACAGGGGCGTGGTGCTGCAGCTCGAGGCCGCGGAGGGGGGCCGCGGTG
GACTTCGGGTCGGAGGGACCGATATACCAGGGCTCTGGCGCGGGGTAGCCCCAGGACGCT
TTCTCGGGGAGGGGAGGGTACGGGCAGCCAGGTGGGAGTGCAAGAACTGGACGGCCTAAT
CCCTGCTGCCTGCGATGTGGGGTCGGACCCAGCGCACCCTGATGGGGCCTGGCGGCGGGG
GACCCCTGAACGGACGTGGCAGCTACTCTGCCGTTGAGCCGATGCGGACGTGCCCAGTAC
ACGTGGGTGTTGGGGGAACTCCACAGCGGCAGGCGAGGGTTGGGTGTAGCAGGGGGCCCA
CGGGGGCGGCGGGTGGCAGGCGCGAGGGGGGCGCGGGGGAGCGATCCGATGGTCGAGAAC
GATCGGCTAGGAGGCGAGCCCCAGGTGGCGCCGGGCGGGTGCGCCCGTGTCGGCGCGGGG
GGGTTGTGGCGACGAGGCCGCCCTTCAGGCGAGGTGGGAGTGGCCCACGTTGTGGCGCCG
CGGTGGAGGGGGGGGCGGTGAGGTGGGGGGCCAGTCGAGGTTGGTGAGGGCGGGACCAGCC
TCGCTTAGTGCAACGCCAACCCGGATAGTAATTCGTGCGGCTCGTGGGAGCGAGTTCTTC
CAATGATGCAGGGGGGGGGTGTGGGGCCCGCGCGTCGACCGGCCGCGTCACGACTGCTACG
GGGTCATTACCGCCGGGCTGTCGTAGGCCGGCGCTCGCGGGAGGTGGTTCGAATGCCGGG
GCGGGAGGCCCGGATCCGGCCAAGGGGCGGGGCTCGGGGGACGCGAGGCGGAGGGGCGCG
AGGCGGTAGCCGGTGCAAGCGGCGCGTGGCCCCATGAGGAAGAACTCTCTCGTGCCCCAA
CACCACACATTAGCGCCCCCATCTCGGCCGACTTCGAGCACGCTCAGCGAGAACCACCAC
AGGCCTCCAACCCGACGTACCACCCGTCCTCACGAGGCGGGCCGACCCTCGGGTCCAAGC
GAGGGCTCCGGCGCGCACGCCCGGCGTGGTGC
>MAP 131_A_tb1504
CCCCTTAGGCGCCTCGAGCGGGTCCCCGCTTCGCCAGTTCACCCGTCCCAGTAAGCGCCC
GCGGCACCTCGCCTTCGCGGAGAACCCCCAGAACGCCCCACCGTCTCCCAGCTCGCACAC
GCGCACGTGTCGCGCTCTCCAGCGCGCGGACTGTTCCAGCCACTGTTCTCATGCTCCCCG
CGGGGCCGTATGACGAGCCACGCCCCAGCCCCGCCGCCGCCGCTCGCCCACCGAGACAAG
CGTAACGCTATTGGGGCCTGACCATGGCTGGGCGGGGCGCTTCCCACGGAGGGCCCAGTC
TTTGGGAGGAGTTGCCCCTCAAGGGATCGGTAGCGCGGTGCGGCTCCCCGCAGCGGCCGC
GAGCGTCGGGGTGATGCTGGAGAGTGGGGCATGCCCCTTCCGGCCGCAGGCGGAACAGGT
TCGCGGTGGTGCACAGGGACGCGGTGCTGCAGCTCGAGACCGCGGAGGGGAGCCGCGGTG
GACTTCGGGTCGGAGGGACCGATATACCAGGGCTCTGACGCGGGGTAACTCTAGGACGCT
CTCTCGAGGAGGGGAGGGTACGGGCAGCCAGGTGGGCGTGCAGGAACTGGACGGCGTAAT
CCCTGGTGACTGCGATGTGGGGTAGGACCCAGCGCACCCTGATGGGGCCTGGCGGCGGGG
GACCCCTGGACGGACGTGGCAGCTACCCTGCCGTTGAGCCGATGCGGACGTGCCTGGTAC
ACGTGGGTGTTGGGGGAACTCCACAGCGGCAGGCGACGGTTGGGTGGGGCCGGGGGCCCA
CGGGGGAGGCGGGTGGCGGTCACGAGGGGGACACGGGGGAGCGATCCGATGGTCGAGAAC
GATCGGCTAGGAGGCGAGCCCCAGGTGGCGCAGGGCGGGTGCGCCCGTGTCGGCGCGGGG
GGGAGGTGGCGACGAGGCGGCCCTTCAGGCGGGGTGAGAATGGCCCAAGTTATGGCGCCG
CGGTGGAGGGGGGGGCGGTGAGGTGGGGGGCCAGTCGGGGTTGGTGAGGGCGGGACCAGCC
TCGCTTGATGCAACGCCAACCCGGATAGTAATTCGTGCGGCTCGTGGTCGCGAGTTCTTC
CAATGATTCAGGGGGGGGGTGTGGGGCCCGCCCGTCGACCGGCCGCGTCACGACTGCTACG
GGGTCATTACCGCCGGGCTGTCGTAGGCCGGCGATCGCGGGAGGTGGTTCGAATGCCGCG
GCGGGAGGCCCGGATCAAGCGAAGGGGCGGGGCCCGGGGCACGCGAGGCGGGGGGGGGCG
AGACGGTAGCCGGTGCAGGCGACGCGTGGCCCCATGAGGTAGAACTCTCTCGTGCCCCAA
CACCACACATTAGCGCCCCCATCTCGGCCGACTTCGAGCGCGCTCAGCGAGAACCACCAC
AGGCCTCCAACCCGACGTACTACCCGTCCTCACGAGGCGGGCCGACCCCCGGGCCCAAGC
GAAGGCTCCGGCGCGCACACCCGGCGTGCTGC
>MAP 109_A_tb1536
CCCCTTAGGCGCCTCGAGCGGGCCCCCGCTTCGCCAGTTCAGCCGTCCCAGTAAGCGCCC
GGGGCACCTCGCCTTCGAGGAGAACCCCCAGAACGCCCCACCGTCTCCCAGCTCGCACAC
GCGCACGCGTCGCACTCTCCCGCGCGCGTACTGTTCCAGCCACCGTCCCCATGCTCCCCG
CGGGGCCGTATGACGAGCCACGCCCCGGCCCCGCCGCCGCCGCTCGCTCACCGAGACCAG
CGCAATGCTATTGGGGCCTGACCACGGCTGGGCGGGGCGCTTCCCACGGAGGGCCCAGTC
TTTGGGAGGAGTTGCCCCTCCAGGGATCGGCAGCGCGGTGCGGCTCCCCGCAGCGGCCGC
GGGCGTCGGGGGGATGCTAGGGAGTGGGGGATGCCCCTTCCGGTCGCAGGCGGAACAGGT
TCGCGGTGGTGCACAGGGGCGTGGTGCTGCAGCTCGAGGCCGCGGAGGGGGGCCGCGGTG
GACTTCGGGTCGGAGGGACCGATATACCAGGGCTCTGGCGCGGGGTAGCCCCAGGACGCT
TTCTCGGGGAGGGGAGGGTACGGGCAGCCAGGTGGGAGTGCAAGAACTGGACGGCCTAAT
CCCTGCTGCCTGCGATGTGGGGTCGGACCCAGCGCACCCTGATGGGGCCTGGCGGCGGGG
GACCCCTGAACGGACGTGGCAGCTACTCTGCCGTTGAGCCGATGCGGACGTGCCCAGTAC
ACGTGGGTGTTGGGGGAACTCCACAGCGGCAGGCGAGGGTTGGGTGTAGCAGGGGGCCCA
CGGGGGCGGCGGGTGGCAGGCGCGAGGGGGGCGCGGGGGAGCGATCCGATGGTCGAGAAC
GATCGGCTAGGAGGCGAGCCCCAGGTGGCGCCGGGCGGGTGCGCCCGTGTCGGCGCGGGG
GGGTTGTGGCGACGAGGCCGCCCTTCAGGCGAGGTGGGAGTGGCCCACGTTGTGGCGCCG
CGGTGGAGGGGGGGGCGGTGAGGTGGGGGGCCAGTCGAGGTTGGTGAGGGCGGGACCAGCC
TCGCTTAGTGCAACGCCAACCCGGATAGTAATTCGTGCGGCTCGTGGGAGCGAGTTCTTC
CAATGATGCAGGGGGGGGGTGTGGGGCCCGCGCGTCGACCGGCCGCGTCACGACTGCTACG
GGGTCATTACCGCCGGGCTGTCGTAGGCCGGCGCTCGCGGGAGGTGGTTCGAATGCCGGG
GCGGGAGGCCCGGATCCGGCCAAGGGGCGGGGCTCGGGGGACGCGAGGCGGAGGGGCGCG
```

AGGCGGTAGCCGGTGCAAGCGGCGCGTGGCCCCATGAGGAAGAACTCTCTCGTGCCCCAA
CACCACACATTAGCGCCCCCATCTCGGCCGACTTCGAGCACGCTCAGCGAGAACCACCAC
AGGCCTCCAACCCGACGTACCACCCGTCCTCACGAGGCGGGCCGACCCTCGGGTCCAAGC
GAGGGCTCCGGCGCGCACGCCCGGCGTGGTGC
>MAP_111_A_tb1536
CCCCTTAGGCGCCTCGAGCGGGCCCCCGCTTCGCCAGTTCACCCGTCCCAGTAAGCGCCC
GGGGCACCTCGCCTTCGAGGAGAACCCCCAGAACGCCCCACCGTCTCCCAGCTCGCACAC
GCGCACGCGTCGCACTCTCCAGCGCGCGTACTGTTCCAGCCACCGTCCCCATGCTCCCCG
CGGGGCCGTATGACGAGCCACGCCCCGGCCCCGCCGCCGCCGCTCGCCCACCGAGACAAG
CGCAATGCTATTGGGGCCTGACCACGGCTGGGCGGGGCGCTTCCCACGGAGGGCCCAGTC
TTTGGGAGGAGTTGCCCCTCCAGGGATCGGTAGCGCGGTGCGGCTCCCCGCAGCGGCCGC
AGGCGTCGGGGGGATGCTGGGGAGTGGGGGATGCCCCTGCCGGTCGCAGGCGGAACAGGT
TCGCGGTGGTGCACAGGGGCGTGGTGCTGCAGCTCGAGGCCGCGGAGGGGGGCCGCGGTG
GACTTCGGGTCGGCGGGACCGGGATACCAGGGCTCTGGCGCGGGGTAGCCCCAGGACGCT
TTCTCGGGGAGGGGAGGGTACGGGCAGCCAGGTGGGAGTGCCAGAACTGGACGGCGTAAT
CCCTGCTGCCTGCGATGTGGGGTCGGACCCAGCGCACCCTGATGGGGCCTGGCGGCGGGG
GACCCCTGAACGGACGTGGCAGCTACTCTGCCGTTGAGCCGATGCGGACGTGCCCAGTAC
ACGGGGGTGTTGGGGGAACTCCACAGCGGCAGGCGAGGGTTGGGTGTAGCAGGGGGCCCA
CGGGGGCGGCGGGTGGCAGGCGCGAGGGGGGCGCGGGGGAGCGATCCGATGGTCGAGAAC
GATCGGCTAGGAGGCGGGCCCCAGGTGGCACCGGGCGGGTGCGCCCGTGTCGGCGCGGGG
GGGTTGTGGCGACGAGGCCGCCCTTCAGGCGAGGTGGGAGTGGCCCAAGTTGTGGCGCCG
CGGTGGAGGGGGGGCGGTGAGGTGGGGGGCCAGTCGAGGTTGGTGAGGGCGGGACCAGCC
TCGTTTAGTGAAACGCCAACCCGGATAGTAATTCGTGCGGCTCGTGGGAGCGAGTTCTTC
CAATGATGCAGGGGGGGGTGTGGGGCCCGCGCGTCGACCGGTCGCGTCACGACTGCTACG
GGGTCATTACCGCCGGGCTGTCGTAGGCCGGCGCTCGCGGAGGGAGGTTCGAATGCCGGG
GCGGGAGGCCCGGATCCGGCGAAGGGGCGGGGCTCGGGGGACGCGAGGCGGAGGGGCGCG
AGGCGGTAGCCGGTGCAAGCGGCGCGTGGCCCCATGAGGAAGAACTCTCTCGTGCCCCAA
CACCACACATTAGCGCCCCCATCTCGGCCGACTTCGAGCACGCTCAACGAGAACCACCAC
AGGCCTCCAACCCGACGTACCACCCGTCCTCACGAGGCGGGCCGACCCTCGGGTCCAAGC
GAGGGCTCCGCCGCGCACGCCCGGCGTGGTGC
>MAP_112_A_tb1536
CCCCTTAGGCGCCTCGAGCGGGCCCCCGCTTCGCCAGTTCAGCCGTCCCAGTAAGCGCCC
GGGGCACCTCGCCTTCGAGGAGAACCCCCAGAACGCCCCACCGTCTCCCAGCTCGCACAC
GCGCACGCGTCGCACTCTCCCGCGCGCGTACTGTTCCAGCCACCGTCCCCATGCTCCCCG
CGGGGCCGTATGACGAGCCACGCCCCGGCCCCGCCGCCGCCGCTCGCTCACCGAGACCAG
CGCAATGCTATTGGGGCCTGACCACGGCTGGGCGGGGCGCTTCCCACGGAGGGCCCAGTC
TTTGGGAGGAGTTGCCCCTCCAGGGATCGGCAGCGCGGTGCGGCTCCCCGCAGCGGCCGC
GGGCGTCGGGGGGATGCTAGGGAGTGGGGGATGCCCCTTCCGGTCGCAGGCGGAACAGGT
TCGCGGTGGTGCACAGGGGCGTGGTGCTGCAGCTCGAGGCCGCGGAGGGGGGCCGCGGTG
GACTTCGGGTCGGAGGGACCGATATACCAGGGCTCTGGCGCGGGGTAGCCCCAGGACGCT
TTCTCGGGGAGGGGAGGGTACGGGCAGCCAGGTGGGAGTGCAAGAACTGGACGGCCTAAT
CCCTGCTGCCTGCGATGTGGGGTCGGACCCAGCGCACCCTGATGGGGCCTGGCGGCGGGG
GACCCCTGAACGGACGTGGCAGCTACTCTGCCGTTGAGCCGATGCGGACGTGCCCAGTAC
ACGTGGGTGTTGGGGGAACTCCACAGCGGCAGGCGAGGGTTGGGTGTAGCAGGGGGCCCA
CGGGGGCGGCGGGTGGCAGGCGCGAGGGGGGCGCGGGGGAGCGATCCGATGGTCGAGAAC
GATCGGCTAGGAGGCGAGCCCCAGGTGGCGCCGGGCGGGTGCGCCCGTGTCGGCGCGGGG
GGGTTGTGGCGACGAGGCCGCCCTTCAGGCGAGGTGGGAGTGGCCCACGTTGTGGCGCCG
CGGTGGAGGGGGGGCGGTGAGGTGGGGGGCCAGTCGAGGTTGGTGAGGGCGGGACCAGCC
TCGCTTAGTGCAACGCCAACCCGGATAGTAATTCGTGCGGCTCGTGGGAGCGAGTTCTTC
CAATGATGCAGGGGGGGGTGTGGGGCCCGCGCGTCGACCGGCCGCGTCACGACTGCTACG
GGGTCATTACCGCCGGGCTGTCGTAGGCCGGCGCTCGCGGGAGGTGGTTCGAATGCCGGG
GCGGGAGGCCCGGATCCGGCCAAGGGGCGGGGCTCGGGGGACGCGAGGCGGAGGGGCGCG
AGGCGGTAGCCGGTGCAAGCGGCGCGTGGCCCCATGAGGAAGAACTCTCTCGTGCCCCAA
CACCACACATTAGCGCCCCCATCTCGGCCGACTTCGAGCACGCTCAGCGAGAACCACCAC
AGGCCTCCAACCCGACGTACCACCCGTCCTCACGAGGCGGGCCGACTCTCGGGTCCAAGC
GAGGGCTCCGGCGCGCACGCCCGGCGTGGTGC
>MAP_108_A_tb1567
CCCCTTAGGCGCCTCGAGCGGGCCCCCGCTTCGCCAGTTCAGCCGTCCCAGTAAGCGCCC
GGGGCACCTCGCCTTCGAGGAGAACCCCCAGAACGCCCCACCGTCTCCCAGCTCGCACAC
GCGCACGCGTCGCACTCTCCCGCGCGCGTACTGTTCCAGCCACCGTCCCCATGCTCCCCG
CGGGGCCGTATGACGAGCCACGCCCCGGCCCCGCCGCCGCCGCTCGCTCACCGAGACCAG
CGCAATGCTATTGGGGCCTGACCACGGCTGGGCGGGGCGCTTCCCACGGAGGGCCCAGTC
TTTGGGAGGAGTTGCCCCTCCAGGGATCGGCAGCGCGGTGCGGCTCCCCGCAGCGGCCGC
GGGCGTCGGGGGGATGCTAGGGAGTGGGGGATGCCCCTTCCGGTCGCAGGCGGAACAGGT
TCGCGGTGGTGCACAGGGGCGTGGTGCTGCAGCTCGAGGCCGCGGAGGGGGGCCGCGGTG
GACTTCGGGTCGGAGGGACCGATATACCAGGGCTCTGGCGCGGGGTAGCCCCAGGACGCT
TTCTCGGGGAGGGGAGGGTACGGGCAGCCAGGTGGGAGTGCAAGAACTGGACGGCCTAAT
CCCTGCTGCCTGCGATGTGGGGTCGGACCCAGCGCACCCTGATGGGGCCTGGCGGCGGGG

GACCCCTGAACGGACGTGGCAGCTACTCTGCCGTTGAGCCGATGCGGACGTGCCCAGTAC
ACGTGGGTGTTGGGGGAACTCCACAGCGGCAGGCGGGGGTTGGGTGTAGCAGGGGGCCCA
CGGGGGCGGCGGGTGGCAGGCGCGAGGGGGGCGCGGGGGAGCGATCCGATGGTCGAGAAC
GATCGGCTAGGAGGCGAGCCCCAGGTGGCGCCGGGCGGGTGCGCCCGTGTCGGCGCGGGG
GGGTTGTGGCGACGAGGCCGCCCTTCAGGCGAGGTGGGAGTGGCCCACGTTGTGGCGCCG
CGGTGGAGGGGGGGGCGGTGAGGTGGGGGGCCAGTCGAGGTTGGTGAGGGCGGGACCAGCC
TCGCTTAGTGCAACGCCAACCCGGATAGTAATTCGTGCGGCTCGTGGGAGCGAGTTCTTC
CAATGATGCAGGGGGGGGGTGTGGGGCCCGCGCGTCGACCGGCCGCGTCACGACTGCTACG
GGGTCATTACCGCCGGGCTGTCGTAGGCCGGCGCTCGCGGGAGGTGGTTCGAATGCCGGG
GCGGGAGGCCCGGATCCGGCCAAGGGGCGGGGCTCGGGGGACGCGAGGCGGAGGGGCGCG
AGGCGGTAGCCGGTGCAAGCGGCGCGTGGCCCCATGAGGAAGAACTCTCTCGTGCCCCAA
CACCACACATTAGCGCCCCCATCTCGGCCGACTTCGAGCACGCTCAGCGAGAACCACCAC
AGGCCTCCAACCCGACGTACCACCCGTCCTCACGAGGCGGGCCGACCCTCGGGTCCAAGC
GAGGGCTCCGGCGCGCACGCCCGGCGTGGTGC
>MAP 110_A_tb1567
CCCCTTAGGCGCCTCGAGCGGGCCCCCGCTTCGCCAGTTCAGCCGTCCCAGTAAGCGCCC
GGGGCACCTCGCCTTCGAGGAGAACCCCCAGAACGCCCCACCGTCTCCCAGCTCGCACAC
GCGCACGCGTCGCACTCTCCCGCGCGCGTACTGTTCCAGCCACCGTCCCCATGCTCCCCG
CGGGGCCGTATGACGAGCCACGCCCCGGCCCCGCCGCCGCCGCTCGCTCACCGAGACCAG
CGCAATGCTATTGGGGCCTGACCACGGCTGGGCGGGGCGCTTCCCACGGAGGGCCCAGTC
TTTGGGAGGAGTTGCCCCTCCAGGGATCGGCAGCGCGGTGCGGCTCCCCGCAGCGGCCGC
GGGCGTCGGGGGGATGCTAGGGAGTGGGGGATGCCCCTTCCGGTCGCAGGCGGAACAGGT
TCGCGGTGGTGCACAGGGGCGTGGTGCTGCAGCTCGAGGCCGCGGAGGGGGGGCCGCGGTG
GACTTCGGGTCGGAGGGACCGATATACCAGGGCTCTGGCGCGGGGTAGCCCCAGGACGCT
TTCTCGGGGAGGGGAGGGTACGGGCAGCCAGGTGGGAGTGCAAGAACTGGACGGCCTAAT
CCCTGCTGCCTGCGATGTGGGGTCGGACCCAGCGCACCCTGATGGGGCCTGGCGGCGGGG
GACCCCTGAACGGACGTGGCAGCTACTCTGCCGTTGAGCCGATGCGGACGTGCCCAGTAC
ACGTGGGTGTTGGGGGAACTCCACAGCGGCAGGCGAGGGTTGGGTGTAGCAGGGGGCCCA
CGGGGGCGGCGGGTGGCAGGCGCGAGGGGGGCGCGGGGGAGCGATCCGATGGTCGAGAAC
GATCGGCTAGGAGGCGAGCCCCAGGTGGCGCCGGGCGGGTGCGCCCGTGTCGGCGCGGGG
GGGTTGTGGCGACGAGGCCGCCCTTCAGGCGAGGTGGGAGTGGCCCACGTTGTGGCGCCG
CGGTGGAGGGGGGGGCGGTGAGGTGGGGGGCCAGTCGAGGTTGGTGAGGGCGGGACCAGCC
TCGCTTAGTGCAACGCCAACCCGGATAGTAATTCGTGCGGCTCGTGGGAGCGAGTTCTCC
CAATGATGCAGGGGGGGGGTGTGGGGCCCGCGCGTCGACCGGCCGCGTCACGACTGCTACG
GGGTCATTACCGCCGGGCTGTCGTAGGCCGGCGCTCGCGGGAGGTGGTTCGAATGCCGGG
GCGGGAGGCCCGGATCCGGCCAAGGGGCGGGGCTCGGGGGACGCGAGGCGGAGGGGCGCG
AGGCGGTAGCCGGTGCAAGCGGCGCGTGGCCCCATGAGGAAGAACTCTCTCGTGCCCCAA
CACCACACATTAGCGCCCCCATCTCGGCCGACTTCGAGCACGCTCAGCGAGAACCACCAC
AGGCCTCCAACCCGACGTACCACCCGTCCTCACGAGGCGGGCCGACCCTCGGGTCCAAGC
GAGGGCTCCGGCGCGCACGCCCGGCGTGGTGC
>MAP 189_A_fb1580
CCCCTTAGGCGCCTCGAGCGGGCCCCCGCTTCGCCAGTTCACCCGTCCCAGTAAGCGCCC
GGGGCACCTCGCCTTCGAGGAGAACCCCCAGAACGCCCCACCGTCTCCCAGCTCGCACAC
GCGCACGCGTCGCACTCTCCAGCGCGCGTACTGTTCCAGCCACCGTCCCCATGCTCCCCG
CGGGGCCGTATGACGAGCCACGCCCCGGCCCCGCCGCCGCCGCTCGCCCACCGAGACAAG
CGCAATGCTATTGGGGCCTGACCACGGCTGGGCGGGGCGCTTCCCACGGAGGGCCCAGTC
TTTGGGAGGAGTTGCCCCTCCAGGGATCGGTAGCGCGGTGCGGCTCCCCGCAGCGGCCGC
AGGCGTCGGGGGGATGCTGGGGAGTGGGGGATGCCCCTGCCGGTCGCAGGCGGAACAGGT
TCGCGGTGGTGCACAGGGGCGTGGTGCTGCAGCTCGAGGCCGCGGAGGGGGGGCCGCGGTG
GACTTCGGGTCGGAGGGACCGGGATACCAGGGCTCTGGCGCGGGGTAGCCCCAGGACGCT
TTCTCGGGGAGGGGAGGGTACGGGCAGCCAGGTGGGAGTGCCAGAACTGGACGGCGTAAT
CCCTGCTGCCTGCGATGTGGGGTCGGACCCAGCGCACCCTGATGGGGCCTGGCGGCGGGG
GACCCCTGAACGGACGTGGCAGCTACTCTGCCGTTGAGCCGATGTGGACGTGCCCAGTAC
ACGTGGGTGTTGGGGGAACTCCACAGCGGCAGGCGAGGGTTGGGTGTAGCAGGGGGCCCA
CGGGGGCGGCGGGTGGCAGGCGCGAGGGGGGCGCGGGGGAGCGATCCGATGGTCGAGAAC
GATCGGCTAGGAGGCGAGCCCCAGGTGGCACCGGGCGGGTGCGCCCGTGTCGGCGCGGGG
GGGTTGTGGCGACGAGGCCGCCCTTCAGGCGAGGTGGGAGTGGCCCAAGTTGTGGCGCCG
CGGTGGAGGGGGGGGCGGTGAGGTGGGGGGCCAGTCGAGGTTGGTGAGGGCGGGACCAGCC
TCGTTTAGTGAAACGCCAACCCGGATAGTAATTCGTGCGGCTCGTGGGAGCGAGTTCTTC
CAATGATGCAGGGGGGGGGTGTGGGGCCCGCGCGTCGACCGGTCGCGTCACGACTGCTACG
GGGTCATTACCGCCGGGCTGTCGTAGGCCGGCGCTCGCGGAGGGAGGTTCGAATGCCGGG
GCGGGAGGCCCGGATCCGGCGAAGGGGCGGGGCTCGGGGGACGCGAGGCGGAGGGGCGCG
AGGCGGTAGCCGGTGCAAGCGGCGCGTGGCCCCATGAGGAAGAACTCTCTCGTGCCCCAA
CACCACACATTAGCGCCCCCATCTCGGCCGACTTCGAGCACGCTCAACGAGAACCACCAC
AGGCCTCCAACCCGACGTACCACCCGTCCTCACGAGGCGGGCCGACCCTCGGGTCCAAGC
GAGGGCTCCGCCGCGCACGCCCGGCGTGGTGC
>MAP 215_A_tb1594
CCCCTTAGGCGCCTCGAGCGGGCCCCCGCTTCGCCAGTTCAGCCGTCCCAGTAAGCGCCC

GGGGCACCTCGCCTTCGAGGAGAACCCCCAGAACGCCCCACCGTCTCCCAGCTCGCACAC
GCGCACGCGTCGCACTCTCCCGCGCGCGTACTGTTCCAGCCACCGTCCCCATGCTCCCCG
CGGGGCCGTATGACGAGCCACGCCCCGGCCCCGCCGCCGCCGCTCGCTCACCGAGACCAG
CGCAATGCTATTGGGGCCTGACCACGGCTGGGCGGGGCGCTTCCCACGGAGGGCCCAGTC
TTTGGGAGGAGTTGCCCCTCCAGGGATCGGCAGCGCGGTGCGGCTCCCCGCAGCGGCCGC
GGGCGTCGGGGGGATGCTAGGGAGTGGGGGATGCCCCTTCCGGTCGCAGGCGGAACAGGT
TCGCGGTGGTGCACAGGGGCGTGGTGCTGCAGCTCGAGGCCGCGGAGGGGGGCCGCGGTG
GACTTCGGGTCGGAGGGACCGATATACCAGGGCTCTGGCGCGGGGTAGCCCCAGGACGCT
TTCTCGGGGAGGGGAGGGTACGGGCAGCCAGGTGGGAGTGCAAGAACTGGACGGCCTAAT
CCCTGCTGCCTGCGATGTGGGGTCGGACCCAGCGCACCCTGATGGGGCCTGGCGGCGGGG
GACCCCTGAACGGACGTGGCAGCTACTCTGCCGTTGAGCCGATGCGGACGTGCCCAGTAC
ACGTGGGTGTTGGGGGAACTCCACAGCGGCAGGCGAGGGTTGGGTGTAGCAGGGGGCCCA
CGGGGGCGGCGGGTGGCAGGCGCGAGGGGGGCGCGGGGGAGCGATCCGATGGTCGAGAAC
GATCGGCTAGGAGGCGAGCCCCAGGTGGCGCCGGGCGGGTGCGCCCGTGTCGGCGCGGGG
GGGTTGTGGCGACGAGGCCGCCCTTCAGGCGAGGTGGGAGTGGCCCACGTTGTGGCGCCG
CGGTGGAGGGGGGGCGGTGAGGTGGGGGGCCAGTCGAGGTTGGTGAGGGCGGGACCAGCC
TCGCTTAGTGCAACGCCAACCCGGATAGTAATTCGTGCGGCTCGTGGGAGCGAGTTCTTC
CAATGATGCAGGGGGGGGGTGTGGGGCCCGCGCGTCGACCGGCCGCGTCACGACTGTTACG
GGGTCATTACCGCCGGGCTGTCGTAGGCCGGCGCTCGCGGGAGGTGGTTCGAATGCCGGG
GCGGGAGGCCCGGATCCGGCCAAGGGGCGGGGCTCGGGGGACGCGAGGCGGAGGGGCGCG
AGGCGGTAGCCGGTGCAAGCGGCGCGTGGCCCCATGAGGAAGAACTCTCTCGTGCCCCAA
CACCACACATTAGCGCCCCCATCTCGGCCGACTTCGAGCACGCTCAGCGAGAACCACCAC
AGGCCTCCAACCCGACGTACCACCCGTCCTCACGAGGCGGGCCGACCCTCGGGTCCAAGC
GAGGGCTCCGGCGCGCACGCCCGGCGTGGTGC
>MAP 181_A_tb1597
CCCCTTAGACGCCTCGAGCGGGTCCCCGCTTCGCCAGTTCACCCGTCCCAGTAAGCGCTC
GCGGCACCTCGCCTTCGCGGAGAACCCCCAGAACGCCCCACCGTCTCCCAGCTCGCACAC
GCGCACGTGTCACACTCTCCAGCGCGCGGACTGTTCCAGCCACCGTCCCCATGCTCCCCG
CGGGGCCGTATGACGAGCCACGCCCCGGCCCCGCCGCCGCCGCTCGCCCACCGAGACAAG
CGTAACGCTATTGGGGCCTGACCATAGCTGGGCGGGGCGCTTCCCACGGAGGGCCCAGTC
TTTGGGAGGAGTTGCCCCTCAAGGGATCGGTAGCGCGGTGCGGCTCCCCGCAGCGGCCGC
GAGCGTCGGGGTGATGCTGGAGAGTGGGGCATGCCCCTTCCGGCCGCAGGCGGAACAGGT
TCGTGGTGGTGCACAGGGACGCGGTGCTGCAGCTCGAGGCCGCGGAGGGGAGCCGCGGTG
GACTTCGGGCCGGAGGGACCGATATACCAGGGCTCTGACGCGGGGTAACTCTAGGACGCT
CTCTCGGGGAGGGGAGGGTACGGGCAGCCAGGTGGGCGTGCAGGAACTGGACGGCGTAAT
CCCTGGTGACTGCGATGTGGGGTAGGACCCAGCGCACCCTGATAGGGCCTGGCGGCGGGG
GACCCCTGGACGGACGTGGCAGCTACCCTGCCGTTGAGCCGATGCGGACGTGCCCAGTAC
ATGTGGGTGTTGGGGGAACTCCACAGCGGCAGGCGACGGGTGGGTGGGGCCGGGGGCCCA
CGGGGGCGGCGGGTGGCGGTCACGAGGGGGGCACGGGGGAGCGATCCGATGGTCGAGAAC
GATCGGCTAGGAGGCGAGCCCCCGGTGGCGCAGGGCGGGTGCGCCCGTGTCGGCGCGGGG
GGGAGGTGGCGACGAGGCGGCCCTTACGGCGGGGTGGGAGTGGCCCAAGTTGTGGCGCCT
CGGTGGAGGGGGGGCGGTGAGGTGGGGGGCCAAGCGGGATTGGTGAGGGCGGGACCAGCC
TCGCTTGATGCAACGCCAACCCGGATAGTAACTCGTGCGGCTTGTGGTCGCGAGTTCTTC
CAATGATTCAGGGGGGGGGTGTGGGGCCCGCCCCTGGACCGGCCGCGTCACGACTGCTACG
GGGTCATTACCGCCGGGCTGTCGTAGGCCGATGATCGCGGGAGGTGGTTCGAATGCCGCG
GCGGGAGGCCCGGATCAAGAGAAGGGGCGGGGCCCGGGGCACGCGAGGTGGGGGGGGGCG
AGGCGGTAGCCGATGCAAGCGGCGCGTGGCCCCATGAGGTAGAACTTTCTCGTGCCCCAA
CACCACACATCAGCGCCCCCATCTCGGCCGACTTCGAGCGCGCTCAGCGAGAACCACCAC
AGGCCTCCAACCCGACGTACCACCCGTCCTCACGAGGCGGGCCGACCCCCGGGCCCAAGC
GAGGGCTCCGGCGCGCACGCCCGGCGTGCTGC
>MAP 182_A_tb1620
CCCCTTAGGCGCCTCGAGCGGGCCCCCGCTTCGCCAGTTCAGCCGTCCCAGTAAGCGCCC
GGGGCACCTCGCCTTCGAGGAGAACCCCCAGAACGCCCCACCGTCTCCCAGCTCGCACAC
GCGCACGCGTCGCACTCTCCCGCGCGCGTACTGTTCCAGCCACCGTCCCCATGCTCCCCG
CGGGGCCGTATGACGAGCCACGCCCCGGCCCCGCCGTCGCCGCTCGCTCACCGAGACCAG
CGCAATGCTATTGGGGCCTGACCACGGCTGGGCGGGGCGCTTCCCACGGAGGGCCCAGTC
TTTGGGAGGAGTTGCCCCTCCAGGGATCGGCAGCGCGGTGCGGCTCCCCGCAGCGGCCGC
GGGCGTCGGGGGGATGCTAGGGAGTGGGGGATGCCCCTTCCGGTCGCAGGCGGAACAGGT
TCGCGGTGGTGCACAGGGGCGTGGTGCTGCAGCTCGAGGCCGCGGAGGGGGGCCGCGGTG
GACTTCGGGTCGGAGGGACCGATATACCAGGGCTCTGGCGCGGGGTAGCCCCAGGACGCT
TTCTCGGGGAGGGGAGGGTACGGGCAGCCAGGTGGGAGTGCAAGAACTGGACGGCCTAAT
CCCTGCTGCCTGCGATGTGGGGTCGGACCCAGCGCACCCTGATGGGGCCTGGCGGCGGGG
GACCCCTGAACGGACGTGGCAGCTACTCTGCCGTTGAGCCGATGCGGACGTGCCCAGTAC
ACGTGGGTGTTGGGGGAACTCCACAGCGGCAGGCGAGGGTTGGGTGTAGCAGGGGGCCCA
CGGGGGCGGCGGGTGGCAGGCGCGAGGGGGGCGCGGGGGAGCGATCCGATGGTCGAGAAC
GATCGGCTAGGAGGCGAGCCCCAGGTGGCGCCGGGCGGGTGCGCCCGTGTCGGCGCGGGG
GGGTTGTGGCGACGAGGCCGCCCTTCAGGCGAGGTGGGAGTGGCCCACGTTGTGGCGCCG
CGGTAGAGGGGGGGCGGTGAGGTGGGGGGCCAGTCGAGGTTGGTGAGGGCGGGACCAGCC

TCGCTTAGTGCAACGCCAACCCGGATAGTAATTCGTGCGGCTCGTGGGAGCGAGTTCTTC
CAATGATGCAGGGGGGGGTGTGGGGCCCGCGCGTCGACCGGCCGCGTCACGACTGCTACG
GGGTCATTACCGCCGGGCTGTCGTAGGCCGGCGCTCGCGGGAGGTGGTTCGAATGCCGGG
GCGGGAGGCCCGGATCCGGCCAAGGGGCGGGGCTCGGGGGACGCGAGGCGGAGGGGCGCG
AGGCGGTAGCCGGTGCAAGCGGCGCGTGGCCCCATGAGGAAGAACTCTCTCGTGCCCCAA
CACCACACATTAGCGCCCCCATCTCGGCCGACTTCGAGCACGCTCAGCGAGAACCACCAC
AGGCCTCCAACCCGACGTACCACCCGTCCTCACGAGGCGGGCCGACCCTCGGGTCCAAGC
GAGGGCTCCGGCGCGCACGCCCGGCGTGGTGC
>MAP 185_A_fb1625
CCCCTTAGGCGCCTCGAGCGGGCCCCCGCTTCGCCAGTTCAGCCGTCCCAGTAAGCGCCC
GGGGCACCTCGCCTTCGAGGAGAACCCCCAGAACGCCCCACCGTCTCCCAGCTCGCACAC
GCGCACGCGTCGCACTCTCCCGCGCGCGTACTGTTCCAGCCACCGTCCCCATGCTCCCCG
CGGGGCCGTATGACGAGCCACGCCCCGGCCCCGCCGCCGCCGCTCGCTCACCGAGACCAG
CGCAATGCTATTGGGGCCTGACCACGGCTGGGCGGGGCGCTTCCCACGGAGGGCCCAGTC
TTTGGGAGGAGTTGCCCCTCCAGGGATCGGCAGCGCGGTGCGGCTCCCCGCAGCGGCCGC
GGGCGTCGGGGGGATGCTAGGGAGTGGGGGATGCCCCTTCCGGTCGCAGGCGGAACAGGT
TCGCGGTGGTGCACGGGGGCGTGGTGCTGCAGCTCGAGGCCGCGGAGGGGGGCCGCGGTG
GACTTCGGGTCGGAGGGACCGATATACCAGGGCTCTGGCGCGGGGTAGCCCCAGGACGCT
TTCTCGGGGAGGGGAGGGTACGGGCAGCCAGGTGGGAGTGCAAGAACTGGACGGCCTAAT
CCCTGCTGCCTGCGATGTGGGGTCGGACCCAGCGCACCCTGATGGGGCCTGGCGGCGGGG
GACCCCTGAACGGACGTGGCAGCTACTCTGCCGTTGAGCCGATGCGGACGTGCCCAGTAC
ACGTGGGTGTTGGGGGAACTCCACAGCGGCAGGCGAGGGTTGGGTGTAGCAGGGGGCCCA
CGGGGGCGGCGGGTGGCAGGCGCGAGGGGGGCGCGGGGGAGCGATCCGATGGTCGAGAAC
GATCGGCTAGGAGGCGAGCCCCAGGTGGCGCCGGGCGGGTGCGCCCGTGTCGGCGCGGGG
GGGTTGTGGCGACGAGGCCGCCCTTCAGGCGAGGTGGGAGTGGCCCACGTTGTGGCGCCG
CGGTGGAGGGGGGGCGGTGAGGTGGGGGGCCAGTCGAGGTTGGTGAGGGCGGGACCAGCC
TCGCTTAGTGCAACGCCAACCCGGATAGTAATTCGTGCGGCTCGTGGGAGCGAGTTCTTC
CAATGATGCAGGGGGGGGTGTGGGGCCCGCGCGTCGACCGGCCGCGTCACGACTGCTACG
GGGTCATTACCGCCGGGCTGTCGTAGGCCGGCGCTCGCGGGAGGTGGTTCGAATGCCGGG
GCGGGAGGCCCGGATCCGGCCAAGGGGCGGGGCTCGGGGGACGCGAGGCGGAGGGGCGCG
AGGCGGTAGCCGGTGCAAGCGGCGCGTGGCCCCATGAGGAAGAACTCTCTCGTGCCCCAA
CACCACACATTAGCGCCCCCATCTCGGCCGACTTCGAGCACGCTCAGCGAGAACCACCAC
AGGCCTCCAACCCGACGTACCACCCGTCCTCACGAGGCGGGCCGACCCTCGGGTCCAAGC
GAGGGCTCCGGCGCGCACGCCCGGCGTGGTGC
>MAP 217_A_fb1625
CCCCTTAGGCGCCTCGAGCGGGCCCCCGCTTCGCCAGTTCACCCGTCCCAGTAAGCGCCC
GGGGCACCTCGCCTTCGAGGAGAACCCCCAGAACGCCCCACCGTCTCCCAGCTCGCACAC
GCGCACGCGTCGCACTCTCCAGCGCGCGTACTGTTCCAGCCACCGTCCCCATGCTCCCCG
CGGGGCCGTATGACGAGCCACGCCCCGGCCCCGCCGCCGCCGCTCGCCCACCGAGACAAG
CGCAATGCTATTGGGGCCTGACCACGGCTGGGCGGGGCGCTTCCCACGGAGGGCCCAGTC
TTTGGGAGGAGTTGCCCCTCCAGGGATCGGTAGCGCGGTGCGGCTCCCCGCAGCGGCCGC
AGGCGTCGGGGGGATGCTGGGGAGTGGGGGATGCCCCTTCCGGTCGCAGGCGGAACAGGT
TCGCGGTGGTGCACAGGGGCGTGGTGCTGCAGCTCGAGGCCGCGGAGGGGGGCCGCGGTG
GACTTCGGGTCGGAGGGACCGATATACCAGGGCTCTGGCGCGGGGTAGCCCCAGGACGCT
TTCTCGGGGAGGGGAGGGTACGGGCAGCCAGGTGGGAGTGCCAGAACTGGACGGCGTAAT
CCCTGCTGCCTGCGATGTGGGGTCGGACCCAGCGCACCCTGATGGGGCCTGGCGGCGGGG
GACCCCTGAACGGACGTGGCAGCTACTCTGCCGTTGAGCCGATGCGGACGTGCCCAGTAC
ACGTGGGTGTTGGGGGAACTCCACAGCGGCAGGCGAGGGTTGGGTGTAGCAGGGGGCCCA
CGGGGGCGGCGGGTGGCAGGCGCGAGGGGGGCGCGGGGGAGCGATCCGATGGTCGAGAAC
GATCGGCTAGGAAGCGAGCCCCAGGTGGCACCGGGCGGGTGCGCCCGTGTCGGCGCGGGG
GGGTTGTGGCGACGAGGCCGCCCTTCAGGCGAAGTGGGAGTGGCCCAAGTTGTGGCGCCG
CGGTGGAGGGGGGGCGGTGAGGTGGGGGGCCAGTCGAGGTTGGTGAGGGCGGGACCAGCC
TCGTTTAGTGAAACGCCAACCCGGATAGTAATTCGTGCGGCTCGTGGGAGCGAGTTCTTC
CAATGATGCAGGGGGGGGTGTGGGGCCCGCGCGTCGACCGGYCGCGTCACGACTGCTACG
GGGTCATTACCGCCGGGCTGTCGTAGGCCGGCGCTCGCGGGAGGTGGTTCGAATGCCGGG
GCGGGAGGCCCGGATCCGGCGAAGGGGCGGGGCTCGGGGGACGCGAGGCGGAGGGGCGCG
AGGCGGTAGCCGGTGCAAGCGGCGCGTGGCCCCATGAGGAAGAACTCTCTCGGGCCCCAA
CACCACACATTAGCGCCCCCATCTCGGCCGACTTCGAGCACGCTCAACGAGAACCACCAC
AGGCCTCCAACCCGACGTACCACCCGTCCTCACGAGGCGGGCCGACCGTCGGGTCCAAGC
GAGGGCTCCGCCGCGCACGCCCGGCGTGGTGC
>MAP 168_A_fb1645
CCCCTTAGGCGCCTCGAGCGGGCCCCCGCTTCGCCAGTTCAGCCGTCCCAGTAAGCGCCC
GGGGCACCTCGCCTTCGAGGAGAACCCCCAGAACGCCCCACCGTCTCCCAGCTCGCACAC
GCGCACGCGTCGCACTCTCCCGCGCGCGTACTGTTCCAGCCACCGTCCCCATGCTCCCCG
CGGGGCCGTATGACGAGCCACGCCCCGGCCCCGCCGCCGCCGCTCGCTCACCGAGACCAG
CGCAATGCTATTGGGGCCTGACCACGGCTGGGCGGGGCGCTTCCCACGGAGGGCCCAGTC
TTTGGGAGGAGTTGCCCCTCCAGGGATCGGCAGCGCGGTGCGGCTCCCCGCAGCGGCCGC
GGGCGTCGGGGGGATGCTAGGGAGTGGGGGATGCCCCTTCCGGTCGCAGGCGGAACAGGT

TCGCGGTGGTGCACAGGGGCGTGGTGCTGCAGCTCGAGGCCGCGGAGGGGGGCCGCGGTG
GACTTCGGGTCGGAGGGACCGATATACCAGGGCTCTGGCGCGGGGTAGCCCCAGGACGCT
TTCTCGGGGAGGGGAGGGTACGGGCAGCCAGGTGGGAGTGCAAGAACTGGACGGCCTAAT
CCCTGCTGCCTGCGATGTGGGGTCGGACCCAGCGCACCCTGATGGGGCCTGGCGGCGGGG
GACCCCTGAACGGACGTGGCAGCTACTCTGCCGTTGAGCCGATGCGGACGTGCCCAGTAC
ACGTGGGTGTTGGGGGAACTCCACAGCGGCAGGCGAGGGTTGGGTGTAGCAGGGGGCCCA
CGGGGGCGGCGGGTGGCAGGCGCGAGGGGGGCGCGGGGGAGCGATCCGATGGTCGAGAAC
GATCGGCTAGGAGGCGAGCCCCAGGTGGCGCCGGGCGGGTGCGCCCGTGTCGGCGCGGGG
GGGTTGTGGCGACGAGGCCGCCCTTCAGGCGAGGTGGGAGTGGCCCACGTTGTGGCGCCG
CGGTGGAGGGGGGGCGGTGAGGTGGGGGGCCAGTCGAGGTTGGTGAGGGCGGGACCAGCC
TCGCTTAGTGCAACGCCAACCCGGATAGTAATTCGTGCGGCTCGTGGGAGCGAGTTCTTC
CAATGATGCAGGGGGGGGTGTGGGGCCCGCGCGTCGACCGGCCGCGTCACGACTGCTACG
GGGTCATTACCGCCGGGCTGTCGTAGGCCGGCGCTCGCGGGAGGTGGTTCGAATGCCGGG
GCGGGAGGCCCGGATCCGGCCAAGGGGCGGGGCTCGGGGGACGCGAGGCGGAGGGGCGCG
AGGCGGTAGCCGGTGCAAGCGGCGCGTGGCCCCATGAGGAAGAACTCTCTCGTGCCCCAA
CACCACACATTAGCGCCCCCATCTCGGCCGACTTCGAGCACGCTCAGCGAGAACCACCAC
AGGCCTCCAACCCGACGTACCACCCGTCCTCACGAGGCGGGCCGACCCTCGGGTCCAAGC
GAGGGCTCCGGCGCGCACGCCCGGCGTGGTGC
>MAP 241_A_tb1683
TCCCTCAGGTGCCTTGAGTGGGCCCCCGCTTCGCCAGTCCACCTGTCCCAGTAAGCGCCC
GCGGCCCCTCGCCTTCGCATGGAACCCTTAGAACGCCTCACCGTCTCCCAGCTTGCACGC
GCGCACGCGTCGCACTTTCCAGCGCGCGGACTTCTCCAGCCGCCGTCCCCAGGCTCCCCG
CGGGGCCGCACGACAAGCTACGTCCCGGTCCCGCCGCCGCCGCCCGCCCACCGAGACAAC
TTCAACGCTATTGGGGCTTGACCACGGCTGGGCATGGCACTTCCCACGGAGGGCCTAGTC
TTCCTGAGGAGCTGCCCCTCAAGGGATCGGTAGTGCGGTGCGGTTCCCCGCAGCGGCCGT
GGGCGTCGGGGTGATCCTGGGGAGTGGGGGATGCCCCTTTCTGCCGCGGGTAGCGCGGGT
TCGCGGTGGTGCACAGGCGCGTGGTGCCGCGGTTCGAGGTGGCGGAGGGGGGTTGCGGTG
GACTTCGGGTCGGATGGACCGATAGACCAGGGCTTTGGCGCGGCGCGGCCGCAGGACACT
CCCTCGGGGAGGGGAGGGTACGGGCAGCCAGGTGGGAGTGCAGGCGCTGGACGGCGTACT
CCCTGGTGACTGGAATGTGGGGTAGGGCCCAGCGCACTCTGGCGAGGCCTGGCGGCGGGG
GCCGCCCGGACGCACGCGGCGGCTACCTTGGCGTTGGGCCGATGCGGACGTGCCCAGTAC
ACGTGGGTGTTAGGGGAACTCCACAGCGGCAGGCGAGGGTTGGGCGTGGCCGAGGGCCCA
CGGGGGCGGCGGGGGGCGGTCGCGAGGGGGGTGCGGGAGAACGATCTGATGATCGAGAAC
GATCGGCCAGAAGGCGAGCCCCAGGTGGCGCAGAGCGTGTGCGCCCGCGTCGGCGCGGGC
GGGTTGTGGCGACGCGGCCACCCTTCAGGCGGGGCGGGAGTGGCCCAAGTTGTGGCGCCG
CGGTGGAGGGGGCCCGGTGAGGTGGGGGGCCAGTCGGGGTTGGTGAGGGCGGAACCAGCC
ACGCTTGGCGCATGGCCAAGGCAGCTAGTAATTCGTGCAGCTCGCGGGCGCAAGTCCTTC
CAATGATTCAGGGGGGGGTGGGGGGCCCGCGCGTCGGCCGGCCGCGTCACGAGTGCTGCG
GGGCCATTACCGCCGGGCCGCCGTAGGCCGGCGATCGCGGGAGGTGGTTCGTGTGCCGGG
GCGGGAAGGCCGGATCAGGCGAAGGGGCGGGGCCCGGGGCGAGCGAGGCGGGGGGACGCG
AGGCGGTAGTCGGCGCAAGCGGCGCGTGGCCTCATGAGGTAGAACTCTCTCATGCCCCAA
CACCATACATTAGCGCCCCCGCTTCGGCCGGGCTCGAGTGCGCTCAGCGAGAACCACCAC
AGGCCTCCAACCCGATGTTCCACTCGTCCTCACGGGGCGGGCCGACCCCGGGGCCCAAGC
GAGGCCTCCGGCGAGCACGCCCGGTGTGGTGC
>MAP 210_A_fb1695
CCCCTTAGGCGCCTCGAGCGGGCCCCCGCTTCGCCAGTTCACCCGTCCCAGTAAGCGCCC
GGGGCACCTCGCCTTCGAGGAGAACCCCCAGAACGCCCCACCGTCTCCCAGCTCGCACAC
GCGCACGCGTCGCACTCTCCAGCGCGCGTACTGTTCCAGCCACCGTCCCCATGCTCCCCG
CGGGGCCGTATGACGAGCCACGCCCCGGCCCCGCCGCCGCCGCTCGCCCACCGAGACAAG
CGCAATGCTATTGGGGCCTGACCACGGCTGGGCGGGGCGCTTCCCACGGAGGGCCCAGTC
TTTGGGAGGAGTTGCCCCTCCAGGGATCGGTAGCGCGGTGCGGCTCCCCGCAGCGGCCGC
AGGCGTCGGGGGGATGCTGGGGAGTGGGGGATGCCCCTGCCGGTCGCAGGCGGAACAGGT
TCGCGGTGGTGCACAGGGGCGTGGTGCTGCAGCTCGAGGCCGCGGAGGGGGGCCGCGGTG
GACTTCGGGTCGGAGGGACCGGGATACCAGGGCTCTGGCGCGGGGTAGCCCCAGGACGCT
TTCTCGGGGAGGGGAGGGTACGGGCAGCCAGGTGGGAGTGCCAGAACTGGACGGCGTAAT
CCCTGCTGCCTGCGATGTGGGGTCGGACCCAGCGCACCCTGATGGGGCCTGGCGGCGGGG
GACCCCTGAACGGACGTGGCAGCTACTCTGCCGTTGAGCCGATGCGGACGTGCCCAGTAC
ACGTGGGTGTTGGGGGAACTCCACAGCGGCAGGCGAGGGTTGGGTGTAGCAGGGGGCCCA
CGGGGGCGGCGGGTGGCAGGCGCGAGGGGGGCGCGGGGGAGCGATCCGATGGTCGAGAAC
GATCGGCTAGGAGGCGAGCCCCAGGTGGCACCGGGCGGGTGCGCCCGTGTCGGCGCGGGG
GGGTTGTGGCGACGAGGCCGCCCTTCAGGCGAGGTGGGAGTGGCCCAAGTTGTGGCGCCG
CGGTGGAGGGGGGGCGGTGAGGTGGGGGGCCAGTCGAGGTTGGTGAGGGCGGGACCAGCC
TCGTTTAGTGAAACGCCAACCCGGATAGTAATTCGTGCGGCTCGTGGGAGCGAGTTCTTC
CAATGATGCAGGGGGGGGTGTGGGGCCCGCGCGTCGACCGGTCGCGTCACGACTGCTACG
GGGTCATTACCGCCGGGCTGTCGTAGGCCGGCGCTCGCGGAGGGAGGTTCGAATGCCGGG
GCGGGAGGCCCGGATCCGGCGAAGGGGCGGGGCTCGGGGGACGCGAGGCGGAGGGGCGCG
AGGCGGTAGCCGGTGCAAGCGGCGCGTGGCCCCATGAGGAAGAACTCTCTCGTGCCCCAA
CACCACACATTAGCGCCCCCATCTCGGCCGACTTCGAGCACGCTCAACGAGAACCACCAC

AGGCCTCCAACCCGACGTACCACCCGTCCTCACGAGGCGGGCCGACCCTCGGGTCCAAGC
GAGGGCTCCGCCGCGCACGCCCGGCGTGGTGC
>MAP 135_A_tb1719
CCCCTTAGGCGCCTCGAGCGGGTCCCCGCTTCGCCAGTTCACCCGTCCCAGTAAGCGCCC
GCGGCACCTCGCCTTCGCGGAGAACCCCCAGAACGCCCCACCGTCTCCCAGCTCGCACAC
GCGCACGTGTCGCGCTCTCCAGCGCGCGGACTGTTCCAGCCACTGTTCTCATGCTCCCCG
CGGGGCCGTATGACGAGCCACGCCCCAGCCCCACCGCCGCCGCTCGCCCACCGAGACAAG
CGTAACGCTATTGGGGCCTGACCATGGCTGGGCGGGGCGCTTCCCACGGAGGGCCCAGTC
TTTGGGAGGAGTTGCCCCTCAAGGGATCGGTAGCGCGGTGCGGCTCCCCGCAGCGGCCGC
GAGCGTCGGGGTGATGCTGGAGAGTGGGGCATGCCCCTTCCGGCCGCAGGCGGAACAGGT
TCGCGGTGGTGCACAGGGACGCGGTGCTGCAGCTCGAGACCGCGGAGGGGAGCCGCGGTG
GACTTCGGGTCGGAGGGACCGATATACCAGGGCTCTGACGCGGGGTAACTCTAGGACGCT
CTCTCGAGGAGGGGAGGGTACGGGCAGCCAGGTGGGCGTGCAGGAACTGGACGGCGTAAT
CCCTGGTGACTGCGATGTGGGGTAGGACCCAGCGCACCCTGATGGGGCCTGGCGGCGGGG
GACCCCTGGACGGACGTGGCAGCTACCCTGCCGTTGAGCCGATGCGGACGTGCCTGGTAC
ACGTGGGTGTTGGGGGAACTCCACAGCGGCAGGCGACGGTTGGGTGGGGCCGGGGGCCCA
CGGGGGAGGCGGGTGGCGGTCACGAGGGGGACACGGGGGAGCGATCCGATGGTCGAGAAC
GATCGGCTAGGAGGCGAGCCCCAGGTGGCGCAGGGCGGGTGCGCCCGTGTCGGCGCGGGG
GGGAGGTGGCGACGAGGCGGCCCTTCAGGCGGGGTGAGAATGGCCCAAGTTATGGCGCCG
CGGTGGAGGGGGGGCGGTGAGGTGGGGGGCCAGTCGGGGTTGGTGAGGGCGGGACCAGCC
TCGCTTGATGCAACGCCAACCCGGATAGTAATTCGTGCGGCTCGTGGTCGCGAGTTCTTC
CAATGATTCAGGGGGGGGTGTGGGGCCCGCCCGTCGACCGGCCGCGTCACGACTGCTACG
GGGTCATTACCGCCGGGCTGTCGTAGGCCGGCGATCGCGGGAGGTGGTTCGAATGCCGCG
GCGGGAGGCCCGGATCAAGCGAAGGGGCGGGGCCCGGGGCACGCGAGGCGGGGGGGGGCG
AGACGGTAGCCGGTGCAGGCGACGCGTGGCCCCATGAGGTAGAACTCTCTCGTGCCCCAA
CACCACACATTAGCGCCCCCATCTCGGCCGACTTCGAGCGCGCTCAGCGAGAACCACCAC
AGGCCTCCAACCCGACGTACTACCCGTCCTCACGAGGCGGGCCGACCCCCGGGCCCAAGC
GAAGGCTCCGGCGCGCACACCCGGCGTGCTGC
>MAP 136_A_tb1719_06
TCCCTCAGGCGCCTCGAGTGGGCCCCCGCTTCGCCAGTCCACCTGTCCCAGTAAGCGCCC
GCGGCCCCTCGCCTTCGCATGGAACCCTTAGAACGGCTCACCGTCTCCCAGCTTGCACGC
GCGCACGCGTCGCACTTTCCAGCGCGCGGACTTCTCCAGCCGCCGTCCCCAGGCTCCCCG
TGGGGCCGCACGACAAGCTACGTCCCGGTCCCGCCGCCGCCGCCCGCCCACCGAGACAAC
TTCAACGCTATTGGGGCTTGACCACGGCTGGGCATGGCACTTCCCACGGAGGGCCCAGTC
TTCCTGAGGAGCTGCCCCTCAAGGGATCGGTAGTGCGGTGCGGTTCCCCGCAGCGGCCGT
GGGCGTCGGGGTGATCCTGGGGAGTGGGGGATGCCCCTTTCTGCCGCGGGTAGCGCGGGT
TCGCGGTGGTGCACAGGCGCGTGGTGCCGCGGTTCGAGGTGGCGGAGGGGGGTTGCGGTG
GACTTCGGGTCGGATGGACCGATAGACCAGGGCTTTGGCGCGGCGCGGCCGCAGGACACT
CTCTCGGGGAGGGGAGGGTACGGGCAGCCAGGTGGGAGTGCAGGCGCTGGACGGTGTACT
CCCTGGTGACTGGGATGTGGGGTAGGACCCAGCGCACTCTGGCGAGGCCTGGCGGCGGGG
GCCCCCCGGACGCACGCGGCGGCTACCTTGGCGTTGGGCCGATGCGGACGTGCCCAGTAC
ACGTGGGTGTTAGGGGAACTCCACAGCGGCAGGCGAGGGTTGGGCGTGGCCGAGGGCCCA
CGGGGGCGGCGGTGGGCGGTCGCGAGGGGGGCGCGGGAGAGCGATCCGATGATCGAGAGC
GATCGGCCAGAAGGCGAGCCCCAGGTGGCGCAGAGCGTGTGCGCCCGCGTCGGCGCGGGG
GGGTTGTGGCGACGCGGCCACCCTTCAGGCGGGGCGGGAGTGGCCCAAGTTGTGGCGCCG
CGGTGGAGGGGGCCCGGTGAGGTGGGGGGCCAGTCGGGGTTGGTGAGGGCGGAACCAGCC
ACGCTTGGCGCATGGCCAAGGCAGATAGTAATTCGTGCAGCTCGCGGGCGCAAGTCCTTC
CAATGATTCAGGGGGGGGTGGGGGGCCCGCGCGTCGGCCGGCCGCGTCACGAGTGCTGCG
GGGCCATTACCGCCGGGCCGCCGTAGGCCGGCGATCGCGGGAGATGGTTCGTGTGCCGGG
GCGGGAAGGCCGGATCAGGCGAAGGGACGGGGCCCGGGGCACGCGAGGCGGGGGGACGCG
AGGCGGTAGTCGGCGCAAGCGGCGCGTGGCCTCATGAGGTAGAACTCTCTCATGCCCCAA
CACCATACATTAGCGCCCCCGCTTCGGCCGGGCTCGAGTGCGCTCAGCGAGAACCACCAC
AGGCCTCCAACCCGACGTTCCACTCGTCCTCACGGGGCGGGCCGACCCCGGGGCCCAAGC
GAGGCCTCCGGCGAGCACGCCCGGTGTGGTGC
>MAP 208_A_fb1773
CCCCTTAGGCGCCTCGAGCGGGCCCCCGCTTCGCCAGTTCACCCGTCCCAGTAAGCGCCC
GGGGCACCTCGCCTTCGAGGAGAACCCCCAGAACGCCCCACCGTCTCCCAGCTCGCACAC
GCGCACGCGTCGCACTCTCCAGCGCGCGTACTGTTCCAGCCACCGTCCCCATGCTCCCCG
CGGGGCCGTATGACGAGCCACGCCCCGGCCCCGCCGCCGCCGCTCGCCCACCGAGACAAG
CGCAATGCTATTGGGGCCTGACCACGGCTGGGCGGGGCGCTTCCCACGGAGGGCCCAGTC
TTTGGGAGGAGTTGCCCCTCCAGGGATCGGTAGCGCGGTGCGGCTCCCCGCAGCGGCCGC
AGGCGTCGGGGGGATGCTGGGGAGTGGGGGATGCCCCTGCCGGTCGCAGGCGGAACAGGT
TCGCGGTAGTGCACAGGGGCGTGGTGCTGCAGCTCGAGGCCGCGGAGGGGGGCCGCGGTG
GACTTCGGGTCGGAGGGACCGGGATACCAGGGCTCTGGCGCGGGGTAGCCCCAGGACGCT
TTCTCGGGGAGGGGAGGGTACGGGCAGCCAGGTGGGAGTGCCAGAACTGGACGGCGTAAT
CCCTGCTGCCTGCGATGTGGGGTCGGACCCAGCGCACCCTGATGGGGCCTGGCGGCGGGG
GACCCCTGAACGGACGTGGCAGCTACTCTGCCGTTGAGCCGATGCGGACGTGCCCAGTAC
ACGTGGGTGTTGGGGGAACTCCACAGCGGCAGGCGAGGGTTGGGTGTAGCAGGGGGGCCCA

CGGGGGCGGCGGGTGGCAGGCGCGAGGGGGGCGCGGGGGAGCGATCCGATGGTCGAGAAC
GATCGGCTAGGAGGCGAGCCCCAGGTGGCACCGGGCGGGTGCGCCCGTGTCGGCGCGGGG
GGGTTGTGGCGACGAGGCCGCCCTTCAGGCGAGGTGGGAGTGGCCCAAGTTGTGGCGCCG
CGGTGGAGGGGGGGCGGTGAGGTGGGGGGCCAGTCGAGGTTGGTGAGGGCGGGACCAGCC
TCGTTTAGTGAAACGCCAACCCGGATAGTAATTCGTGCGGCTCGTGGGAGCGAGTTCTTC
CAATGATGCAGGGGGGGGTGTGGGGCCCGCGCGTCGACCGGTCGCGTCACGACTGCTACG
GGGTCATTACCGCCGGGCTGTCGTAGGCCGGCGCTCGCGGAGGGAGGTTCGAATGCCGGG
GCGGGAGGCCCGGATCCGGCGAAGGGGCGGGGCTCGGGGGACGCGAGGCGGAGGGGCGCG
AGGCGGTAGCCGGTGCAAGCGGCGCGTGGCCCCATGAGGAAGAACTCTCTCGTGCCCCAA
CACCACACATTAGCGCCCCCATCTCGGCCGACTTCGAGCACGCTCAACGAGAACCACCAC
AGGCCTCCAACCCGACGTACCACCCGTCCTCACGAGGCGGGCCGACCCTCGGGTCCAAGC
GAGGGCTCCGCCGCGCACGCCCGGCGTGGTGC
>MAP 139_A_tb2158
CCCCTTAGGCGCCTCGAGCGGGCCCCCGCTTCGCCAGTTCACCCGTCCCAGTAAGCGCCC
GGGGCACCTCGCCTTCGAGGAGAACCCCCAGAACGCCCCACCGTCTCCCAGCTCGCACAC
GCGCACGCGTCGCACTCTCCAGCGCGCGTACTGTTCCAGCCACCGTCCCCATGCTCCCCG
CGGGGCCGTATGACGAGCCACGCCCCGGCCCCGCCGCCGTCGCTCGCCCGCCGAGACAAG
CGCAATGCTATTGGGGCCTGACCACGGCTGGGCGGGGCGCTTCCCACGGAGGGCCCAGTC
TTTGGGAGGAGTTGCCCCTCCAGGGATCGGTAGCGCGGTGCGGCTCCCCGCAGCGGCCGC
AGGCGTCGGGGGGATGCTGGGGAGTGGGGGATGCCCCTTCCGGTCGCAGGCGGAACAGGT
TCGCGGTGGTGCACAGGGGCGTGGTGCTGCAGCTCGAGGCCGCGGAGGGGGGCCGCGGTG
GACTTCGGGTCGGAGGGACCGATATACCAGGGCTCTGGCGCGGGGTAGCCCCAGGACGCT
TTCTCGGGGAGGGGAGGGTACGGGCAGCCAGGTGGGAGTGCCAGAACTGGACGGCGTAAT
CCCTGCTGCCTGCGATGTGGGGTCGGACCCAGCGCACCCTGATGGGGCCTGGCGGCGGGG
GACCCCTGAACGGACGTGGCAGMTACTCTGCCGTTGAGCCGATGCGGACGTGCCCAGTAC
ACGTGGGTGTTGGGGGAACTCCACAGCGGCAGGCGAGGGTTGGGTGTAGCAGGGGGCCCA
CGGGGGCGGCGGGTGGCAGGCGCGAGGGGGGCGCGGGGGAGCGATCCGATGGTCGAGAAC
GATCGGCTAGGAGGCGAGCCCCAGGTGGCACCGGGCGGGTGCGCCCGTGTCGGCGCGGGG
GGGTTGTGGCGACGAGGCCGCCCTTCAGGCGAGGTGGGAGTGGCCCAAGTTGTGGCGCCG
CGGTGGAGGGGGGGCGGTGAGGTGGGGGGCCAGTCGAGGTTGGTGAGGGCGGGACCAGCC
TCGTTTAGTGAAACGCCAACCCGGATAGTAATTCGTGCGGCTCGTGGGAGCGAGTTCTTC
CAATGATGCAGGGGGGGGTGTGGGGCCCGCGCGTCGACCGGTCGCGTCACGACTGCTACG
GGGTCATTACCGCCGGGCTGTCSTAGGCCGGCGCTCGCGGGAGGTGGTTCGAATGCCGGG
GCGGGAGGCCCGGATCCGGCGAAGGGGCGGGGCTCGGGGGACGCGAGGCGGAGGGGCGCG
AGGCGGTAGCCGGTGCAAGCGGCGCGTGGCCCCATGAGGAAGAACTCTCTCGTGCCCCAA
CACCACACATTAGCGCCCCCATCTCGGCCGACTTCGAGCACGCTCAACGAGAACYACCAC
AGGCCTCCAACCCGACGTACCACCCGTCCTCACGAGGCGGGCCGACCCTCGGGTCCAAGC
GAGGGCTCCGCCGCGTATGCCCGGCGTGGTGC
>MAP 142_A_tb2158
CCCCTTAGGCGCCTCGAGCGGGCCCCCGCTTCGCCAGTTCACCCGTCCCAGTAAGCGCCC
GGGGCACCTCGCCTTCGAGGAGAGCCCCCAGAACGCCCCACCGTCTCCCAGCTCGCACAC
GCGCACGCGTCGCACTCTCCAGCGCGCGTACTGTTCCAGCCACCGTCCCCATGCTCCCCG
CGGGGCCGTATGACGAGCCACGCCCCGGCCCCGCTGCCGCCGCTCGCCCACCGAGACAAG
CGCAATGCTATTGGGGCCTGACCACGGCTGGGCGGGGCGCTTCCCACGGAGGGCCCAGTC
TTTGGGAGGAGTTGCCCCTCCAGGGATCGGTAGCGCGGTGCGGCTCCCCGCAGCGGCCGC
AGGCGTCGGGGGGATGCTGGGGAGTGGGGGATGCCCCTGCCGGTCGCAGGCGGAAAAGGT
TCGCGGTGGTGCACAGGGGCGTGGTGCTGCAGCTCGAGGCCGCGGAGGGGGGCCGCGGTG
CACTTCGGGTCGGAGTGACCGGGATACCAGGGCTCTGGCGCGGGGTAGCCCCAGGACGCT
TTCTCGGGGAGGGGAGGGTACGGGCAGCCAGGTGGGAGTGCCAGAACTGGACGGCGTAAT
CCCTGCTGCCTGCGATGTGGGGTCGGACCCAGCGCACCCTGATGGGGCCTGGCGGCGGGG
GACCCCTGAACGGACGTGGCAGCTACTCTGCCGTTGAGCCGATGCGGACGTGCCCAGTAC
ACGTGGGTGTTGGGGGAACTCCACAGCGGCAGGCGAGGGTTGGGTGTAGCAGGGGGCCCA
CGGGGGCGGCGGGTGGCAGGCGCGCGGGGGGCGCGGGGGAGCGATCCGATGGTCGAGAAC
GATCGGCTAGGAGGCGAGCCCCAGGTGGCACCGGGCGGGTGCGCCCGTGTCGGCGCGGGG
GGGTTGTGGCGAGGAGGCCGCCCTTCAGGCGAGGTGGGAGTGGCCCAAGTTGTGGCGCCG
CGGTGGAGGGGGGGCGGTGAGGTGGGGGGCCAGTCGAGGTTGGTGAGGGCGGGACCAGCC
TCGTTTAGTGAAACGCCAACCCGGATAGTAATTCGTGCGGCTCGTGGGAGCGAGTTCTTC
CAATGATGCAGGGGGGGGTGTGGGGCCCGCGCGTCGACCGGTCGCGTCACGACTGCTACG
GGGTCATTACCGCCGGGCTGTCGTAGGCCGGCGCTCGCGGAGGGAGGTTCGAATGCCGGG
GCGGGAGGCCCGGATCCGGCGAAGGGGCGGGGCTCGGGGGACGCGAGGCGGAGGGGCGCG
AGGCGGTAGCCGGTGCAAGCGGCGCGTGGCCCCATGAGGAAGAACTCTCTCGTGCCCCAA
CACCACACATTAGCGCCCCCATCTCGGCCGACTTCGAGCACGCTCAACGAGAACCACCAC
AGGCCTCCAACCCGACGTACCACCCGTCCTCACGAGGCGGGCCGACCCTCGGGTCCAAGC
GAGGGCTCCGCCGCGCACGCCCGGCGTGGTGC
>MAP 143_A_fasb2158
CCCCTTAGGCGCCTCGAGCGGGCCCCCGCTTCGCCAGTTCACCCGTCCCAGTAAGCGCCC
GGGGCACCTCGCCTTCGAGGAGAACCCCCAGAACGCCCCACCGTCTCCCAGCTCGCACAC
GCGCACGCGTCGCACTCTCCAGCGCGCGTACTGTTCCAGCCACCGTCCCCATGCTCCCCG

CGGGGCCGTATGACGAGCCACGCCCCGGCCCCGCCGCCGCCGCTCGCCCACCGAGACAAG
CGCAATGCTACTGGGGCCTGACCACGGCTGGGCGGGGCGCTCCCCACGGAGGGCCCAGTC
TTTGGGAGGAGTTGCCCCTCCAGGGATCGGTAGCGCGGTGCGGCTCCCCGCAGCGGCCGC
AGGCGTCGGGGGGATGCTGGGGAGTGGGGGATGCCCCTGCCGGTCGCAGGCGGAACAGGT
TCGCGGTGGTGCACAGGGGCGTGGTGCTGCAGCTCGAGGCCGCGGAGGGGGGCCGCGGTG
GACTTCGGGTCGGAGGGACCGGGATACCAGGGCTCTGGCGCGGGGTAGCCCCAGGACGCT
TTCTCGGGGAGGGGAGGGTACGGGCAGCCAGGTGGGAATGCCAGAACTGGACGGCGTAAT
CCCTGCTGCCTGCGATGTGGGCTCGGACCCAGCGCACCCTGATGGGGCCTGGCGGCGGGG
GACCCCTGAACGGACGTGGCAGCTACTCTGCCGTTGAGCCGATGCGGACGTGCCCAGTAC
ACGTGGGTGTTGGGGGAACTCCACAGCGGCAGGCGAGGGTTGGGTGTAGCAGGGGGCCCA
CGGGGGCGGCGGGTGGCAGGCGCGAGGGGGGCGCGGGGGAGCGATCCGATGGTCGAGAAC
GATCGGCTAGGAGGCGAGCCCCAGGTGGCACCGGGCGGGTGCGCCCGTGTCGGCGCGGGG
GGGTTGTGGCGACGAGGCCGCCCTTCAGGCGAGGTGGGAGTGGCCCAAGTTGTGGCGCCG
CGGTGGAGGGGGGGCGGTGAGGTGGGGGGCCAGTCGAGGTTGGTGAGGGCGGGACCAGCC
TCGTTTAGTGAAACGCCAACCCGGATAGTAATTCGTGCGGCTCGTGGGAGCGAGTTCTTC
CAATGATGCAGGGGGGGGGGTGGGGCCGGCGCGTCGACCGGTCGCGTCACGACTGCTACG
GGGTCATTACCGCCGGGCTGTCGTAGGCCGGCGCTCGCGGGAGGAGGTTCGAATGCCGGG
GCGGGAGGCCCGGATCCGGCGAAGGGGCGGGGCTCGGGGGACGCGAGGCGGAGGGGCGCG
AGGCGGTAGCCGGTGCAAGCGGCGCGTGGCCCCATGAGGAAGAACTCTCTCGTGCCCCAA
CACCACACATTAGCGCCCCCATCTCGGCCGACTTCGAGCACGCTCAACGAGAACCACCAC
AGGCCTCCAACCCGACGTACCACCCGTCCTCACGAGGCGGGCCGACCCTCGGGTCCAAGC
GAGGGCTCCGCCGCGCACGCCCGGCGTGGTGC
>MAP_066_A_fb2653_98
CCCCTTAGGCGCCTCGAGCGGGCCCCCGCTTCGCCAGTTCACCCGTCCCAGTAAGCGCCC
GGGGCACCTCGCCTTCGAGGAGAACCCCCAGAACGCCCCACCGTCTCCCAGCTCGCACAC
GCGCACGCGTCGCACTCTCCAGCGCGCGTACTGTTCCAGCCACCGTCCCCATGCTCCCCG
CGGGGCCGTATGACGAGCCACGCCCCGGCCCCGCCGCCGCCGCTCGCCCACCGAGACAAG
CGCAATGCTATTGGGGCCTGACCACGGCTGGGCGGGGCGCTTCCCACGGAGGGCCCAGTC
TTTGGGAGGAGTTGCCCCTCCAGGGATCGGTAGCGCGGTGCGGCTCCCCGCAGCGGCCGC
AGGCGTCGGGGGGATGCTGGGGAGTGGGGGATGCCCCTGCCGGTCGCAGGCGGAACAGGT
TCGCGGTGGTGCACAGGGGCGTGGTGCTGCAGCTCGAGGCCGCGGAGGGGGGCCGCGGTG
GACTTCGGGTCGGAGGGACCGGGATACCAGGGCTCTGGCGCGGGGTAGCCCCAGGACGCT
TTCTCGGGGAGGGGAGGGTACGGGCAGCCAGGTGGGAGTGCCAGAACTGGACGGCGTAAT
CCCTGCTGCCTGCGATGTGGGGTCGGACCCAGCGCACCCTGATGGGGCCTGGCGGCGGGG
GACCCCTGAACGGACGTGGCAGCTACTCTGCCGTTGAGCCGATGCGGACGTGCCCAGTAC
ACGTGGGTGTTGGGGGAACTCCACAGCGGCAGGCGAGGGTTGGGTGTAGCAGGGGGCCCA
CGGGGGCGGCGGGTGGCAGGCGCGAGGGGGGCGCGGGGGAGCGATCCGATGGTCGAGAAC
GATCGGCTAGGAGGCGAGCCCCAGGTGGCACCGGGCGGGTGCGCCCGTGTCGGCGCGGGG
GGGTTGTGGCGACGAGGCCGCCCTTCAGGCGAGGTGGGAGTGGCCCAAGTTGTGGCGCCG
CGGTGGAGGGGGGGCGGTGAGGTGGGGGGCCAGTCGAGGTTGGTGAGGGCGGGACCAGCC
TCGTTTAGTGAAACGCCAACCCGGATAGTAATTCGTGCGGCTCGTGGGAGCGAGTTCTTC
CAATGATGCAGGGGGGGGGTGTGGGGCCCGCGCGTCGACCGGTCGCGTCACGACTGCTACG
GGGTCATTACCGCCGGGCTGTCGTAGGCCGGCGCTCGCGGAGGGAGGTTCGAATGCCGGG
GCGGGAGGCCCGGATCCGGCGAAGGGGCGGGGCTCGGGGGACGCGAGGCGGAGGGGCGCG
AGGCGGTAGCCGGTGCAAGCGGCGCGTGGCCCCATGAGGAAGAACTCTCTCGTGCCCCAA
CACCACACATTAGCGCCCCCATCTCGGCCGACTTCGAGCACGCTCAACGAGAACCACCAC
AGGCCTCCAACCCGACGTACCACCCGTCCTCACGAGGCGGGCCGACCCTCGGGTCCAAGC
GAGGGCTCCGCCGCGCACGCCCGGCGTGGTGC
>MAP_023_A_fb2669_99
CCCCTTAGGCGCCTCGAGCGGGCCCCCGCTTCGCCAGTTCACCCGTCCCAGTAAGCGCCC
GGGGCACCTCGCCTTCGAGGAGAACCCCCAGAACGCCCCACCGTCTCCCAGCTCGCACAC
GCGCACGCGTCGCACTCTCCAGCGCGCGTACTGTTCCAGCCACCGTCCCCATGCTCCCCG
CGGGGCCGTATGACGAGCCACGCCCCGGCCCCGCCGCCGCCGCTCGCCCACCGAGACAAG
CGCAATGCTATTGGGGCCTGACCACGGCTGGGCGGGGCGCTTCCCACGGAGGGCCCAGTC
TTTGGGAGGAGTTGCCCCTCCAGGGATCGGTAGCGCGGTGCGGCTCCCCGCAGCGGCCGC
AGGCGTCGGGGGGATGCTGGGGAGTGGGGGATGCCCCTGCCGGTCGCAGGCGGAACAGGT
TCGCGGTGGTGCACAGGGGCGTGGTGCTGCAGCTCGAGGCCGCGGAGGGGGGCCGCGGTG
GACTTCGGGTCGGAGGGACCGGGCTACCAGGGCTCTGGCGCGGGGTAGCCCCAGGACGCT
TTCTCGGGGAGGGGAGGGTACGGGCAGCCAGGTGGGAGTGCCAGAACTGGACGGCGTAAT
CCCTGCTGCCTGCGATGTGGGGTCGGACCCAGCGCACCCTGATGGGGCCTGGCGGCGGGG
GACCCCTGAACGGACGTGGCAGCTACTCTGCCGTTGAGCCGATGCGGACGTGCCCAGTAC
ACGTGGGTGTTGGGGGAACTCCACAGCGGCAGGCGAGGGTTGGGTGTAGCAGGGGGCCCA
CGGGGGCGGCGGGTAGCAGGCGCGAGGGGGGCGCGGGGGAGCGATCCGATGGTCGAGAAC
GATCGGCTAGGAGGCGAGCCCCAGGTGGCACCGGGCGGGTGCGCCCGTGTCGGCGCGGGG
GGGTTGTGGCGACGAGGCCGCCCTTCAGGCGAGGTGGGAGTGGCCCAAGTTGTGGCGCCG
CGGTGGAGGGGGGGCGGTGAGGTGGGGGGCCAGTCGAGGTTGGTGAGGGCGGGACCAGCC
TCGTTTAGTGAAACGCCAACCCGGATAGTAATTCGTGCGGCTCGTGGGAGCGAGTTCTTC
CAATGATGCAGGGGGGGGGTGTGGGGCCCGCGCGTCGACCGGTCGCGTCACGACTGCTACG

GGGTCATTACCGCCGGGCTGTCGTAGGCCGGCGCTCGCGGAGGGAGGTTCGAATGCCGGG
GCGGGAGGCCCGGATCCGGCGAAGGGGCGGGGCTCGGGGGACGCGAGGCGGAGGGGCGCG
AGGCGGTAGCCGGTGCAAGCGGCGCGTGGCCCCATGAGGAAGAACTCTCTCGTGCCCCAA
CACCACACATTAGCGCCCCCATCTCGGCCGACTTCGAGCACGCTCAACGAGAACCACCAC
AGGCCTCCAACCCGACGTACCACCCGTCCTCACGAGGCGGGCCGACCCTCGGGTCCAAGC
GAGGGCTCCGCCGCGCACGCCCGGCGTGGTGC
>MAP 169_A_fb727
TCCCTCAGGCGCCTCGAGTGGGCCCCCGCTTCGCCAGTCCACCTGTCCCAGTAAGCGCCC
GCGGCCCCTCGCCTTCGCATGGAACCCTTAGAACGCCTCACCGTCTCCCAGCTTGCACGC
GCGCACGCGTCGCACTTTCCAGCGCGCGGACTTCTCCAGCCGCCGTCCCCAGGCTCTCCG
CGGGGCCGCACGACAAGCTACGTCCCGGTCCCGCCGCCGCCGCCCGCCCACCGAGACAAC
TTCAACGCTATTGGGGCTTGACCACGGCTGGGCATGGCACTTCCCACGGAGGGCCCAGTC
TTCCTGAGGAGCTGCCCCTCAAGGGATCGGTAGTGCGGTGCGGTTCCCCGCAGCGGCCGT
GGGCGTCGGGGTGATCCTGGGGAGTGGGGGATGCCCCTTTCTGCCGCGGGTAGCGCGGGT
TCGCGGTGGTGCACAGGCGCGTGGTGCCGCGGTTCGAGGTGGCGGAGGGGGGTTGCGGTG
GACTTCGGGTCGGATGGACCGATAGACCAGGGCTTTGGCGCGGCGCGGCCGCAGGACACT
CTCTCGGGGAGGGGAGGGTACGGGCAGCCAGGTGGGAGTGCAGGCGCTGGACGGCGTACT
CCCTGGTGACTGGGATGTGGGGTAGGACCCAGCGTACTCTGGCGAGGCCTGGCGGCGGGG
GCCCCCCGGACGCACGCGGCGGCTACCTTGGCGTTGGGCCGATGCGGACGTGCGCAGTAC
ACGTGGGTGTTAGGGGAACTACACAGCGGCAGGCGAGGGTTGGGCGTGGCCGAGGGCCCA
CGGGGGCGGCGGGGGGCGGTCGCGAGGGGGGCGCGGGAGAGCGATCCGATGATCGAGAAC
GATCGGCCAGAAGGCGAGCCCCAGGTGGCGCAGAGCGTGTGCGCCCGCGTCGGCGCGGGG
GGGTTGTGGCGACGCGGCCACCCTTCAGGCGGGGCGGGAGTGGCCCAAGTTGTGGCGCCG
CGGTGGAGGGGGCCCGGTGAGGTGGGGGGCCAGTCGGGGTTGGTGAGGGCGGAACCAGCC
ACGCTTGGCGCATGGCCAAGGCAGATAGTAATTCGTGCAGCTCGCGGGCGCAAGTCCTTC
CAATGATTCAGGGGGGGGTGGGGGGCCCGCGCGTCGGCCGGCCGCGTCACGAGTGCTGCG
GGGCCATTACCGCCGGGCCGCCGTAGGCCGGCGATCGCGGGAGGTGGTTCGTGTGCCGGG
GCGGGAAGGCCGGATCAGGCGAAGGGGCGGGGCCCGGGGCACGCGAGGCGGGGGGACGCG
AGGCGGTAATCGGCGCAAGCGGCGCGTGGCCTCATGAGGTAGAACTCTCTCATGCCCCAA
CACCATACATTAGCGCCCCCGCTTCGGCCGGGCTCGAGTGCGCTCAGCGAGAACCACCAC
AGGCCTCCAACCCGACGTTCCACTCGTCCTCACGGGGCGGGCCGACCCCGGGGCCCAAGC
GAGGCCTCCGGCGAGCACGCCCGGTGTGGTGC
>MAP 012_A_e
CCCCTTAGGCGCCTCGAGCGGGCCCCCGCTTCGCCAGTTCACCCGTCCCAGTAAGCGCCC
GGGGCACCTCGCCTTCGAGGAGAACCCCCAGAACGCCCCACCGTCTCCCAGCTCGCACAC
GCGCACGCGTCGCACTCTCCAGCGCGCGTACTGTTCCAGCCACCGTCCCCATGCTCCCCG
CGGGGCCGTATGACGAGCCACGCCCCGGCCCCGCCGCCGCCGCTCGCCCACCGAGACAAG
CGCAATGCTATTGGGGCCTGACCACGGCTGGGCGGGGCGCTTCCCACGGAGGGCCCAGTC
TTTGGGAGGAGTTGCCCCTCCAGGGATCGGTAGCGCGGTGCGGCTCCCCGCAGCGGCCGC
AGGCGTCGGGGGGATGCTGGGGAGTGGGGGATGCCCCTGCCGGTCGCAGGCGGAACAGGT
TCGCGGTGGTGCACAGGGGCGTGGTGCTGCAGCTCGAGGCCGCGGAGGGGGGCCGCGGTG
GACTTCGGGTCGGAGGGACCGGGATACGAGGGCTCTGGCGCGGGGTAGCCCCAGGACGCT
TTCTCGGGGAGGGGAGGGTACGGGCAGCCAGGTGGGAGTGCCAGAACTGGACGGCGTAAT
CCCTGCTGCCTGCGATGTGGGGTCGGACCCAGCGCACCCTGATGGGGCCTGGCGGCGGGG
GACCCCTGAACGGACGTGGCAGCTACTCTGCCGTTGAGCCGATGCGGACGTGCCCAGTAC
ACGTGGGTGTTGGGGGAACTCCACAGCGGCAGGCGAGGGTTGGGTGTAGCAGGGGGCCCA
CGGGGGCGGCGGGTGGCAGGCGCGAGGGGGGCGCGGGGGAGCGATCCGATGGTCGAGAAC
GATCGGCTAGGAGGTGAGCCCCAGGTGGCACCGGGCGGGTGCGCCCGTGTCGGCGCGGGG
GGGTTGTGGCGACGAGGCCGCCCTTCAGGCGAGGTGGGAGTGGCCCAAGTTGTGGCGCCG
CGGTGGAGGGGGGGCGGTGAGGTGGGGGGCCAGTCGAGGTTGGTGAGGGCGGGACCAGCC
TCGTTTAGTGAAACGCCAACCCGGATAGTAATTCGTGCGGCTCGTGGGAGCGAGTTCTTC
CAATGATGCAGGGGGGGGTGTGGGGCCCGCGCGTCGACCGGTCGCGTCACGACTGCTACG
GGGTCATTACCGCCGGGCTGTCGTAGGCCGGCGCTCGCGGAGGGAGGTTCGAATGCCGGG
GCGGGAGGCCCGGATCCGGCGAAGGGGCGGGGCTCGGGGGACGCGAGGCGGAGGGGCGCG
AGGCGGTAGCCGGTGCAAGCGGCGCGTGGCCCCATGAGGAAGAACTCTCTCGTGCCCCAA
CACCACATATTAGCGCCCCCATCTCGGCCGACTTCGAGCACGCTCAACGAGAACCACCAC
AGGCCTCCAACCCGACGTACCACCCGTCCTCACGAGGCGGGCCGACCCTCGGGTCCAAGC
GAGGGCTCCGCCGCGCACGCCCGGCGTGGTGC
>MAP 014_A_e
CCCCTTAGGCGCCTCGAGCGGGCCCCCGCTTCGCCAGTTCAGCCGTCCCAGTAAGCGCCC
GGGGCACCTCGCCTTCGAGGAGAACCCCCAGAACGCCCCACCGTCTCCCAGCTCGCACAC
GCGCACGCGTCGCACTCTCCCGCGCGCGTACTGTTCCAGCCACCGTCCCCATGCTCCCCG
CGGGGCCGTATGACGAGCCACGCCCCGGCCCCGCCGCCGCCGCTCGCTCACCGAGACCAG
CGCAATGCTATTGGGGCCTGACCACGGCTGGGCGGGGCGCTTCCCACGGAGGGCCCAGTC
TTTGGGAGGAGTTGCCCCTCCAGGGATCGGCAGCGCGGTGCGGCTCCCCGCAGCGGCCGC
GGGCGTCGGGGGGATGCTAGGGAGTGGGGGATGCCCCTTCCGGTCGCAGGCGGAACAGGT
TCGCGGTGGTGCACAGGGGCGTGGTGCTGCAGCTCGAGGCCGCGGAGGGGGGCCGCGGTG
GACTTCGGGTCGGAGGGACCGATATACCAGGGCTCTGGCGCGGGGTAGCCCCAGGACGCT

TTCTCGGGGAGGGGAGGGTACGGGCAGCCAGGTGGGAGTGCAAGAACTGGACGGCCTAAT
CCCTGCTGCCTGCGATGTGGGGTCGGACCCAGCGCACCCTGATGGGGCCTGGCGGCGGGG
GACCCCTGAACGGACGTGGCAGCTACTCTGCCGTTGAGCCGATGCGGACGTGCCCAGTAC
ACGTGGGTGTTGGGGGAACTCCACAGCGGCAGGCGAGGGTTGGGTGTAGCAGGGGGCCCA
CGGGGGCGGCGGGTGGCAGGCGCGAGGGGGGCGCGGGGGAGCGATCCGATGGTCGAGAAC
GATCGGCTAGGAGGCGAGCCCCAGGTGGCGCCGGGCGGGTGCGCCCGTGTCGGCGCGGGG
GGGTTGTGGCGACGAGGCCGCCCTTCAGGCGAGGTGGGAGTGGCCCACGTTGTGGCGCCG
CGGTGGAGGGGGGGCGGTGAGGTGGGGGGCCAGTCGAGGTTGGTGAGGGCGGGACCAGCC
TCGCTTAGTGCAACGCCAACCCGGATAGTAATTCGTGCGGCTCGTGGGAGCGAGTTCTTC
CAATGATGCAGGGGGGGGTGTGGGGCCCGCGCGTCGACCGGCCGCGTCACGACTGCTACG
GGGTCATTACCGCCGGGCTGTCGTAGGCCGGCGCTCGCGGGAGGTGGTTCGAATGCCGGG
GCGGGAGGCCCGGATCCGGCCAAGGGGCGGGGCTCGGGGGACGCGAGGCGGAGGGGCGCG
AGGCGGTAGCCGGTGCAAGCGGCGCGTGGCCCCATGAGGAAGAACTCTCTCGTGCCCCAA
CACCACACATTAGCGCCCCCATCTCGGCCGACTTCGAGCACGCTCAGCGAGAACCACCAC
AGGCCTCCAACCCGACGTACCACCCGTCCTCACGAGGCGGGCCGACCCTCGGGTCCAAGC
GAGGGCTCCGGCGCGCACGCCCGGCGTGGTGC
>MAP 015_A_e
CCCCTTAGGCGCCTCGAGCGGGCCCCCGCTTCGCCAGTTCACCCGTCCCAGTAAGCGCCC
GGGGCACCTCGCCTTCGAGGAGAACCCCCAGAACGCCCCACCGTCTCCCAGCTCGCACAC
GCGCACGCGTCGCACTCTCCAGCGCGCGTACTGTTCCAGCCACCGTCCCCATGCTCCCCG
CGGGGCCGTATGACGAGCCACGCCCCGGCCCCGCCGCCGCCGCTCGCCCACCGAGACAAG
CGCAATGCTATTGGGGCCTGACCACGGCTGGGCGGGGCGCTTCCCACGGAGGGCCCAGTC
TCTGGGAGGAGTTACCCCTCCAGGGATCGGTAGCGCGGTGCGGCTCCCCGCAGCGGCCGC
GGGCGTCGGAGGGATGCTGGGGAGTGGGGGATGCCCCTTCCGGTCGCAGGCGGAACAGGT
TCGCGGTGGTGCACAGGGGCGTGGTGCTGCAGCTCGAGGCCGCGGAGGGGGGCCGCGATG
GACTTCGGGTCGCAGGAACCGATATACCAGGGCTCTGGCGCGGGGTAGCCCCAGGACGCT
TTCTCGGGGAGGGGAGGGTACGGGCAGCCAGGTGGGAGTGCAAGAACTGGACGGCGTAAT
CCCTGCTGCCTGCGATGTGGGGTCGGACCCAGCGCACCCTGATGGGGCCTGGCGGCGGGG
GACCCCTGAGCGGACGTGGCAGCTACTCTGCCGTTGAGCCGATGCGGACGTGCCCAGTAC
ACGTGGGTGTTGGGGGAACTCCACAGCGGCAGGCGAGGGTTGGGTGTAGCAGGGGGCCCA
CGGGGGCGGCGGGTGGCAGGCGCGATGGGGGCGCGGGGGAGCGATCCGATGGTCGAGAAC
GATCGGCTAGGAGGCGAGCCCCAGGTGGCGCCGGTCGGGTGCGCCCGTGTCGGCGCGGGG
GGGTTGTGGCGACGAGGCCGCCCTTCAGGCGAGGTGGGAGCGGCCCAAGTTGTGGCGCCG
CGGTGGAGGGGGGGCGGTGAGGTGGGGGGCCAGTCGAGGCTGGTGAGGGCGGGACCAACC
TCGCTTAGTGCAACGCCAACCTGGATAGTAATTCGTGCGGCTCGTGGGAGCGAGTTCTTC
CAACGATGCAGGGGGGGGTGTGGGGCCCGCGCGTCGACCGGCCGCGTCACGACTGCTACG
GGGTCATTACCGCCGGGCTGTCGTAGGCCGGCGCTCGCGGGAGGTGGTTCGAATGCCGGG
GCGGGAGGCCCGGATCCGGCGAAGGGGCGGGGCTCGGGGGACGCGAGGCGGAGGAGCGCG
AGGCGGTAGCCGGTGCAAGAGGCGCGTGGCCCCATGAGGAAGAACTCTCTCGTGCCCCAA
CATCACACATTAGCGCCCCCATCTCGGCCGACTTCGAGCGCGCTCAGCGAGAACCACCAC
AGGCCTCCAACCCGACGTACCACCCATCCTCACGAGGCGGGCCGACCCTCGGGTCCAAGC
GAGGGCTCCGGCGCGCACGCCCGGCGTGGTGC
>MAP 016_A_e
CCCCTTAGGCGCCTCGAGCGGGCCCCCGCTTCGCCAGTTCAGCCGTCCCAGTAAGCGCCC
GGGGCACCTCGCCTTCGAGGAGAACCCCCAGAACGCCCCACCGTCTCCCAGCTCGCACAC
GCGCACGCGTCGCACTCTCCCGCGCGCGTACTGTTCCAGCCACCGTCCCCATGCTCCCCG
CGGGGCCGTATGACGAGCCACGCCCCGGCCCCGCCGCCGCCGCTCGCTCACCGAGACCAG
CGCAATGCTATTGGGGCCTGACCACGGCTGGGCGGGGCGCTTCCCACGGAGGGCCCAGTC
TTTGGGAGGAGTTGCCCCTCCAGGGATCGGCAGCGCGGTGCGGCTCCCCGCAGCGGCCGC
GGGCGTCGGGGGGATGCTAGGGAGTGGGGGATGCCCCTTCCGGTCGCAGGCGGAACAGGT
TCGCGGTGGTGCACAGGGGCGTGGTGCTGCAGCTCGAGGCCGCGGAGGGGGGCCGCGGTG
GACTTCGGGTCGGAGGGACCGATATACCAGGGCTCTGGCGCGGGGTAGCCCCAGGACGCT
TTCTCGGGGAGGGGAGGGTACGGGCAGCCAGGTGGGAGTGCAAGAACTGGACGGCCTAAT
CCCTGCTGCCTGCGATGTGGGGTCGGACCCAGCGCACCCTGATGGGGCCTGGCGGCGGGG
GACCCCTGAACGGACGTGGCAGCTACTCTGCCGTTGAGCCGATGCGGACGTGCCCAGTAC
ACGTGGGTGTTGGGGGAACTCCACAGCGGCAGGCGAGGGTTGGGTGTAGCAGGGGGCCCA
CGGGGGCGGCGGGTGGCAGGCGCGAGGGGGGCGCGGGGGAGCGATCCGATGGTCGAGAAC
GATCGGCTAGGAGGCGAGCCCCAGGTGGCGCCGGGCGGGTGCGCCCGTGTCGGCGCGGGG
GGGTTGTGGCGACGAGGCCGCCCTTCAGGCGAGGTGGGAGTGGCCCACGTTGTGGCGCCG
CGGTGGAGGGGGGGCGGTGAGGTGGGGGGCCAGTCGAGGTTGGTGAGGGCGGGACCAGCC
TCGCTTAGTGCAACGCCAACCCGGATAGTAATTCGTGCGGCTCGTGGGAGCGAGTTCTTC
CAATGATGCAGGGGGGGGTGTGGGGCCCGCGCGTCGACCGGCCGCGTCACGACTGCTACG
GGGTCATTACCGCCGGGCTGTCGTAGGCCGGCGCTCGCGGGAGGTGGTTCGAATGCCGGG
GCGGGAGGCCCGGATCCGGCCAAGGGGCGGGGCTCGGGGGACGCGAGGCGGAGGGGCGCG
AGGCGGTAGCCGGTGCAAGCGGCGCGTGGCCCCATGAGGAAGAACTCTCTCGTGCCCCAA
CACCACACATTAGCGCCCCCATCTCGGCCGACTTCGAGCACGCTCAGCGAGAACCACCAC
AGGCCTCCAACCCGACGTACCACCCGTCCTCACGAGGCGGGCCGACCCTCGGGTCCAAGC
GAGGGCTCCGGCGCGCACGCCCGGCGTGGTGC

>MAP 019_A_e
CCCCTTAGGCGCCTCGAGCGGGCCCCCGCTTCGCCAGTTCACCCGTCCCAGTAAGCGCCC
GGGGCACCTCGCCTTCGAGGAGAACCCCCAGAACGCCCCACCGTCTCCCAGCTCGCACAC
GCGCACGCGTCGCACTCTCCAGCGCGCGTACTGTTCCAGCCACCGTCCCCATGCTCCCCG
CGGGGCCGTATGACGAGCCACGCCCCGGCCCCGCCGCCGTCGCTCGCCCGCCGAGACAAG
CGCAATGCTATTGGGGCCTGACCACGGCTGGGCGGGGCGCTTCCCACGGAGGGCCCAGTC
TTTGGGAGGAGTTGCCCCTCCAGGGATCGGTAGCGCGGTGCGGCTCCCCGCAGCGGCCGC
AGGCGTCGGGGGGATGCTGGGGAGTGGGGGATGCCCCTTCCGGTCGCAGGCGGAACAGGT
TCGCGGTGGCGCACAGGGGCGTGGTGCTGCAGCTCGAGGCCGCGGAGGGGGGCCGCGGTG
GACTTCGGGTCGGAGGGACCGATATACCAGGGCTCTGGCGCGGGGTAGCCCCAGGACGCT
TTCTCGGGGAGGGGAGGGTACGGGCAGCCAGGTGGGAGTGCCAGAACTGGACGGCGTAAT
CCCTGCTGCCTGCGATGTGGGGTCGGACCCAGCGCACCCTGATGGGGCCTGGCGGCGGGG
GACCCCTGAACGGACGTGGCAGCTACTCTGCCGTTGAGCCGATGCGGACGTGCCCAGTAC
ACGTGGGTGTTGGGGGAACTCCACAGCGGCAGGCGAGGGTTGGGTGTAGCAGGGGGCCCA
CGGGGGCGGCGGGTGGCAGGCGCGAGGGGGGCGCGGGGGAGCGATCCGATGGTCGAGAAC
GATCGGCTAGGAGGCGAGCCCCAGGTGGCACCGGGCGGGTGCGCCCGTGTCGGCGCGGGG
GGGTTGTGGCGACGAGGCCGCCCTTCAGGCGAGGTGGGAGTGGCCCAAGTTGTGGCGCCG
CGGTGGAGGGGGGGCGGTGAGGTGGGGGGCCAGTCGAGGTTGGTGAGGGCGGGACCAGCC
TCGTTTAGTGAAACGCCAACCCGGATAGTAATTCGTGCGGCTCGTGGGAGCGAGTTCTTC
CAATGATGCAGGGGGGGGGTGTGGGGCCCGCGCGTCGACCGGTCGCGTCACGACTGCTACG
GGGTCATTACCGCCGGGCTGTCGTAGGCCGGCGCTCGCGGGAGGTGGTTCGAATGCCGGG
GCGGGAGGCCCGGATCCGGCGAAGGGGCGGGGCTCGGGGGACGCGAGGCGGAGGGGCGCG
AGGCGGTAGCCGGTGCAAGCGGCGCGTGGCCCCATGAGGAAGAACTCTCTCGTGCCCCAA
CACCACACATTAGCGCCCCCATCTCGGCCGACTTCGAGCACGCTCAACGAGAACTACCAC
AGGCCTCCAACCCGACGTACCAGCCGTCCTCACGAGGCGGGCCGACCCTCGGGTCCAAGC
GAGGGCTCCGCCGCGTATGCCCGGCGTGGTGC
>MAP 030_A_e
CCCCTTAGGCGCCTCGAGCGGGCCCCCGCTTCGCCAGTTCAGCCGTCCCAGTAAGCGCCC
GGGGCACCTCGCCTTCGAGGAGAACCCCCAGAACGCCCCACCGTCTCCCAGCTCGCACAC
GCGCACGCGTCGCACTCTCCCGCGCGCGTACTGTTCCAGCCACCGTCCCCATGCTCCCCG
CGGGGCCGTATGACGAGCCACGCCCCGGCCCCGCCGCCGCCGCTCGCTCACCGAGACCAG
CGCAATGCTATTGGGGCCTGACCACGGCTGGGCGGGGCGCTTCCCACGGAGGGCCCAGTC
TTTGGGAGGAGTTGCCCCTCCAGGGATCGGCAGCGCGGTGCGGCTCCCCGCAGCGGCCGC
GGGCGTCGGGGGGATGCTAGGGAGTGGGGGATGCCCCTTCCGGTCGCAGGCGGAACAGGT
TCGCGGTGGTGCACAGGGGCGTGGTGCTGCAGCTCGAGGCCGCGGAGAGGGGCCGCGGTG
GACTTCGGGTCGGAGGGACCGATATACCAGGGCTCTGGCGCGGGGTAGCCCCAGGACGCT
TTCTCGGGGAGGGGAGGGTACGGGCAGCCAGGTGGGAGTGCAAGAACTGGACGGCCTAAT
CCCTGCTGCCTGCGATGTGGGGTCGGACCCAGCGCACCCTGATGGGGCCTGGCGGCGGGG
GACCCCTGAACGGACGTGGCAGCTACTCTGCCGTTGAGCCGATGCGGACGTGCCCAGTAC
ACGTGGGTGTTGGGGGAACTCCACAGCGGCAGGCGAGGGTTGGGTGTAGCAGGGGGCCCA
CGGGGGCGGCGGGTGGCAGGCGCGAGGGGGGCGCGGGGGAGCGATCCGATGGTCGAGAAC
GATCGGCTAGGAGGCGAGCCCCAGGTGGCGCCGGGCGGGTGCGCCCGTGTCGGCGCGGGG
GGGTTGTGGCGACGAGGCCGCCCTTCAGGCGAGGTGGGAGTGGCCCACGTTGTGGCGCCG
CGGTGGAGGGGGGGCGGTGAGGTGGGGGGCCAGTCGAGGTTGGTGAGGGCGGGACCAGCC
TCGCTTAGTGCAACGCCAACCCGGATAGTAATTCGTGCGGCTCGTGGGAGCGAGTTCTTC
CAATGATGCAGGGGGGGGGTGTGGGGCCCGCGCGTCGACCGGCCGCGTCACGACTGCTACG
GGGTCATTACCGCCGGGCTGTCGTAGGCCGGCGCTCGCGGGAGGTGGTTCGAATGCCGGG
GCGGGAGGCCCGGATCCGGCCAAGGGGCGGGGCTCGGGGGACGCGAGGCGGAGGGGCGCG
AGGCGGTAGCCGGTGCAAGCGGCGCGTGGCCCCATGAGGAAGAACTCTCTCGTGCCCCAA
CACCACACATTAGCGCCCCCATCTCGGCCGACTTCGAGCACGCTCAGCGAGAACCACCAC
AGGCCTCCAACCCGACGTACCACCCGTCCTCACGAGGCGGGCCGACCCTCGGGTCCAAGC
GAGGGCTCCGGCGCGCACGCCCGGCGTGGTGC
>MAP 050_A_e
CCCCTTAGGCGCCTCGAGCGGGCCCCCGCTTCGCCAGTTCACCCGTCCCAGTAAGCGCCC
GGGGCACCTCGCCTTCGAGGAGAACCCCCAGAACGCCCCACCGTCTCCCAGCTCGCACAC
GCGCACGCGTCGCACTCTCCAGCGCGCGTACTGTTCCAGCCACCGTCCCCATGCTCCCCG
CGGGGCCGTATGACGAGCCACGCCCCGGCCCCGCCGCCGCCGCTCGCCCACCGAGACAAG
CGCAATGCTATTGGGGCCTGACCACGGCTGGGCGGGGCGCTTCCCACGGAGGGCCCAGTC
TTTGGGAGGAGTTGCCCCTCCAGGGATCGGTAGCGCGGTGCGGCTCCCCGCAGCGGCCGC
AGGCGTCGGGGGGATGCTGGGGAGTGGGGGATGCCCCTGCCGGTCGCAGGCGGAACAGGT
CCGCGATGGTGCACAGGGGCGTGGTGCTGCAGCTCGAGGCCGCGGAGGGGGGCCGCGGTG
GACTTCGGGTCGGAGGGACCGGGATACCAGGGCTCTGGCGCGGGGTAGCCCCAGGACGCT
TTCTCGGGGAGGGGAGGGTACGGGCAGCCAGGTGGGAGTGCCAGAACTGGACGGCGTAAT
CCCTACTGCCTGCGATGTGGGGTCGGACCCAGCGCACCCTGATGGGGCCTGGCGGCGGGG
GACCCCTGAACGGACGTGGCAGCTACTCTGCCGTTGAGCCGATGCGGACGTGCCCAGTAC
ACGTGGGTGTTGGGGGAACTCCACAGCGGCAGGCGAGGGTTGGGTGTAGCAGGGGGCCCA
CGGGGGCGGCGGGTGGCAGGCGCGAGGGGGGCGCGGGGGAGCGATCCGATGGTCGAGAAC
GATCGGCTAGGAGGCGAGCCCCAGGTGGCACCGGGCGGGTGCGCCCGTGTCGGCGCGGGG

GGGTTGTGGCGACGAGGCCGCCCTTCAGGCGAGGTGGGAGTGGCCCAAGTTGTGGCGCCG
CGGTGGAGGGGGGGCGGTGAGGTGGGGGGCCAGTCGAGGTTGGTGAGGGCGGGACCAGCC
TCGTTTAGTGAAACGCCAACCCGGATAGTAATTCGTGCGGCTCGTGGGAGCGAGTTCTTC
CAATGATGCAGGGGGGGGTGTGGGGCCCGCGCGTCGACCGGTCGCGTCACGACTGCTACG
GGGTCATTACCGCCGGGCTGTCGTAGGCCGGCGCTCGCGGAGGGAGGTTCGAATGCCGGG
GCGGGAGGCCCGGATCCGGCGAAGGGGCGGGGCTCGGGGGACGCGAGGCGGAGGGGCGCG
AGGCGGTAGCCGGTGCAAGCGGCGCGTGGCCCCATGAGGAAGAACTCTCTCGTGCCCCAA
CACCACACATTAGCGCCCCTATCTCGGCCGACTTCGAGCACGCTCAACGAGAACCACCAC
AGGCCTCCAACCCGACGTACCACCCGTCCTCACGAGGCGGGCCGACCCTCGGGTCCAAGC
GAGGGCTCCGCCGCGCACGCCCGGCGTGGTGC
>MAP 060_A_e
CCCCTTAGGCGCCTCGAGCGGGCCCCCGCTTCGCCAGTTCAGCCGTCCCAGTAAGCGCCC
GGGGCACCTCGCCTTCGAGGAGAACCCCCAGAACGCCCCACCGTCTCCCAGCTCGCACAC
GCGCACGCGTCGCACTCTCCCGCGCGCGTACTGTTCCAGCCACCGTCCCCATGCTCCCCG
CGGGGCCGTATGACGAGCCACGCCCCGGCCCCGCCGCCGCCGCTCGCTCACCGAGACCAG
CGCAATGCTATTGGGGCCTGACCACGGCTGGGCGGGGCGCTTCCCACGGAGGGCCCAGTC
TTTGGGAGGAGTTGCCCCTCCAGGGATCGGCAGCGCGGTGCGGCTCCCCGCAGTGGCCGC
GGGCGTCGGGGGGATGCTAGGGAGTGGGGGATGCCCCTTCCGGTCGCAGGCGGAACAGGT
TCGCGGTGGTGCACAGGGGAGTGGTGCTGCAGCTCGAGGCCGCGGAGGGGGGCCGCGGTG
GACTTCGGGTCGGAGGGACCGATATACCAGGGCTCTGGCGCGGGGTAGCCCCAGGACGCT
TTCTCGGGGAGGGGAGGGTACGGGCAGCCAGGTGGGAGTGCAAGAACTGGACGGCCTAAT
CCCTGCTGCCTGCGATGTGGGGTCGGACCCAGCGCACCCTGATGGGGCCTGGCGGCGGGG
GACCCCTGAACGGACGTGGCAGCTACTCTGCCGTTGAGCCGATGCGGACGTGCCCAGTAC
ACGTGGGTGTTGGGGGAACTCCACAGCGGCAGGCGAGGGTTGGGTGTAGCAGGGGGCCCA
CGGGGGCGGCGGGTGGCAGGCGCGAGGGGGGCGCGGGGGAGCGATCCGATGGTCGAGAAC
GATCGGCTAGGAGGCGAGCCCCAGGTGGCGCCGGGCGGGTGCGCCCGTGTCGGCGCGGGG
GGGTTGTGGCGACGAGGCCGCCCTTCAGGCGAGGTGGGAGTGGCCCACGTTGTGGCGCCG
CGGTGGAGGGGGGGCGGTGAGGTGGGGGGCCAGTCGAGGTTGGTGAGGGCGGGACCAGCC
TCGCTTAGTGCAACGCCAACCCGGATAGTAATTCGTGCGGCTCGTGGGAGCGAGTTCTTC
CAATGATGCAGGGGGGGGTGTGGGGCCCGCGCGTCGACCGGCCGCGTCACGACTGCTACG
GGGTCATTACCGCCGGGCTGTCGTAGGCCGGCGCTCGCGGGAGGTGGTTCGAATGCCGGG
GCGGGAGGCCCGGATCCGGCCAAGGGGCGGGGCTCGGGGGACGCGAGGCGGAGGGGCGCG
AGGCGGTAGCCGGTGCAAGCGGCGCGTGGCCCCATGAGGAAGAACTCTCTCGTGCCCCAA
CACCACACATTAGCGCCCCCATCTCGGCCGACTTCGAGCACGCTCAGCGAGAACCACCAC
AGGCCTCCAACCCGACGTACCACCCGTCCTCACGAGGCGGGCCGACCCTCGGGTCCAAGC
GAGGGCTCCGGCGCGCACGCCCGGCGTGGTGC
>MAP 107_A_e
CCCCTTAGGCGCCTCGAGCGGGCCCCCGCTTCGCCAGTTCAGCCGTCCCAGTAAGCGCCC
GGGGCACCTCGCCTTCGAGGAGAACCCCCAGAACGCCCCACCGTCTCCCAGCTCGCACAC
GCGCACGCGTCGCACTCTCCCGCGCGCGTACTGTTCCAGCCACCGTCCCCATGCTCCCCG
CGGGGCCGTATGACGAGCCACGCCCCGGCCCCGCCGCCGCCGCTCGCTCACCGAGACCAG
CGCAATGCTATTGGGGCCTGACCACGGCTGGGCGGGGCGCTTCCCACGGAGGGCCCAGTC
TTTGGGAGGAGTTGCCCCTCCAGGGATCGGCAGCGCGGTGCGGCTCCCCGCAGCGGCCGC
GGGCGTCGGGGGGATGCTAGGGAGTGGGGGATGCCCCTTCCGGTCGCAGGCGGAACAGGT
TCGCGGTGGTGCACAGGGGCGTGGTGCTGCAGCTCGAGGCCGCGGAGGGGGGCCGCGGTG
GACTTCGGGTCGGAGGGACCGATATACCAGGGCTCTGGCGCGGGGTAGCCCCAGGACGCT
TTCTCGGGGAGGGGAGGGTACGGGCAGCCAGGTGGGAGTGCAAGAACTGGACGGCCTAAT
CCCTGCTGCCTGCGATGTGGGGTCGGACCCAGCGCACCCTGATGGGGCCTGGCGGCGGGG
GACCCCTGAACGGACGTGGCAGCTACTCTGCCGTTGAGCCGATGCGGACGTGCCCAGTAC
ACGTGGGTGTTGGGGGAACTCCACAGCGGCAGGCGAGGGTTGGGTGTAGCAGGGGGCCCA
CGGGGGCGGCGGGTGGCAGGCGCGAGGGGGGCGCGGGGGAGCGATCCGATGGTCGAGAAC
GATCGGCTAGGAGGCGAGCCCCAGGTGGCGCCGGGCGGGTGCGCCCGTGTCGGCGCGGGG
GGGTTGTGGCGACGAGGCCGCCCTTCAGGCGAGGTGGGAGTGGCCCACGTTGTGGCGCCG
CGGTGGAGGGGGGGCGGTGAGGTGGGGGGCCAGTCGAGGTTGGTGAGGGCGGGACCAGCC
TCGCTTAGTGCAACGCCAACCCGGATAGTAATTCGTGCGGCTCGTGGGAGCGAGTTCTCC
CAATGATGCAGGGGGGGGTGTGGGGCCCGCGCGTCGACCGGCCGCGTCACGACTGCTACG
GGGTCATTACCGCCGGGCTGTCGTAGGCCGGCGCTCGCGGGAGGTGGTTCGAATGCCGGG
GCGGGAGGCCCGGATCCGGCCAAGGGGCGGGGCTCGGGGGACGCGAGGCGGAGGGGCGCG
AGGCGGTAGCCGGTGCAAGCGGCGCGTGGCCCCATGAGGAAGAACTCTCTCGTGCCCCAA
CACCACACATTAGCGCCCCCATCTCGGCCGACTTCGAGCACGCTCAGCGAGAACCACCAC
AGGCCTCCAACCCGACGTACCACCCGTCCTCACGAGGCGGGCCGACCCTCGGGTCCAAGC
GAGGGCTCCGGCGCGCACGCCCGGCGTGGTGC
>MAP 126_A_e
CCCCTTAGGCGCCTCGAGCGGGCCCCCGCTTCGCCAGTTCAGCCGTCCCAGTAAGCGCCC
GGGGCACCTCGCCTTCGAGGAGAACCCCCAGAACGCCCCACCGTCTCCCAGCTCGCACAC
GCGCACGCGTCGCACTCTCCCGCGCGCGTACTGTTCCAGCCACCGTCCCCATGCTCCCCG
CGGGGCCGTATGACGAGCCACGCCCCGGCCCCGCCGCCGCCGCTCGCTCACCGAGACCAG
CGCAATGCTATTGGGGCCTGACCACGGCTGGGCGGGGCGCTTCCCACGGAGGGCCCAGTC

TTTGGGAGGAGTTGCCCCTCCAGGGATCGGCCGCGCGGTGCGGCTCCCCGCAGCGGCCGC
GGGCGTCGGGGGGATGCTAGGGAGTGGGGGATGCCCCTTCCGGTCGCAGGCGGAACAGGT
TCGCGGTGGTGCACAGGGGCGTGGTGCTGCAGCTCGAGGCCGCGGAGGGGGGCCGCGGTG
GACTTCGGGTCGGAGGGACCGATATACCAGGGCTCTGGCGCGGGGTAGCCCCAGGACGCT
TTCTCGGGGAGGGGAGGGTACGGGCAGCCAGGTGGGAGTGCAAGAACTGGACGGCCTAAT
CCCTGCTGCCTGCGATGTGGGGTCGGACCCAGCGCACCCTGATGGGGCCTGGCGGCGGGG
GACCCCTGAACGGACGTGGCAGCTACTCTGCCGTTGAGCCGATGCGGACGTGCCCAGTAC
ACGTGGGTGTTGGGGGAACTCCACAGCGGCAGGCGAGGGTTGGGTGTAGCAGGGGGCCCA
CGGGGGCGGCGGGTGGCAGGCGCGAGGGGGGCGCGGGGGAGCGATCCGATGGTCGAGAAC
GATCGGCTAGGAGGCGAGCCCCAGGTGGCGCCGGGCGGGTGCGCCCGTGTCGGCGCGGGG
GGGTTGTGGCGACGAGGCCGCCCTTCAGGCGAGGTGGGAGTGGCCCACGTTGTGGCGCCG
CGGTGGAGGGGGGGCGGTGAGGTGGGGGGCCAGTCGAGGTTGGTGAGGGCGGGACCAGCC
TCGCTTAGTGCAACGCCAACCCGGATAGTAATTCGTGCGGCTCGTGGGAGCGAGTTCTTC
CAATGATGCAGGGGGGGGTGTGGGGCCCGCGCGTCGACCGGCCGCGTCACGACTGCTACG
GGGTCATTACCGCCGGGCTGTCGTAGGCCGGCGCTCGCGGGAGGTGGTTCGAATGCCGGG
GCGGGAGGCCCGGATCCGGCCAAGGGGCGGGGCTCGGGGGACGCGAGGCGGAGGGGCGCG
AGGCGGTAGCCGGTGCAAGCGGCGCGTGGCCCCATGAGGAAGAACTCTCTCGTGCCCCAA
CACCACACATTAGCGCCCCCATCTCGGCCGACTTCGAGCACGCTCAGCGAGAACCACCAC
AGGCCTCCAACCCGACGTACCACCCGTCCTCACGAGGCGGGCCGACCCTCGGGTCCAAGC
GAGGGCTCCGGCGCGCACGCCCGGCGTGGTGC
>MAP 144_A_e
CCCCTTAGGCGCCTCGAGCGGGCCCCCGCTTCGCCAGTTCACCCGTCCCAGTAAGCGCCC
GGGGCACCTCGCCTTCGAGGAGAACCCCCAGAACGCCCCACCGTCTCCCAGCTCGCACAC
GCGCACGCGTCGCACTCTCCAGCGCGCGTACTGTTCCAGCCACCGTCCCCATGCTCCCCG
CGGGGCCGTATGACGAGCCACGCCCCGGCCCCGCCGCCGCCGCTCGCCCACCGAGACAAG
CGCAATGCTATTGGGGCCTGACCACGGCTGGGCGGGGCGCTTCCCACGGAGGGCCCAGTC
TTTGGGAGGAGTTACCCCTCCAGGGATCGGTAGCGCGGTGCGGCTCCCCGCAGCGGCCGC
GGGCGTCGGGGGGATGCTGGGGAGTGGGGGATGCCCCTTCCGGTCGCAGGCGGAACAGGT
TCGCGGTGGTGCACAGGGGCGTGGTGCTGTAGCTCGAGGCCGCGGAGGGGGGCCGCGGTG
GACTTCGGGTCGGAGGGACTGATATACCAGGGCTCTGGCGCGGGGTAGCCCCAGGACGCT
TTCTCGGGGAGGGGAGGGTACGGGCAGCCAGGTGGGAGTGCAAGAACTGGACGGCGTAAT
CCCTGCTGCCTGCGATGTGGGGTCGGACCCAGCGCACCCTGATGGGGCCTGGCGGCGGGG
GACCCCTGAGCGGACGTGGCAGCTACTCTGCCGTTGAGCCGATGCGGACGTGCCCAGTAC
ACGTGGGTGTTGGGGGAACTCCACAGCGGCAGGCGAGGGTTGGGTGTAGCAGGGGGCCCA
CGGGGGCGGCGGGTGGCAGGCGCGAGGGGGGCGCGGGGGAGCGATCCGATGGTCGAGAAC
GATCGGCTAGGAGGCGAGCCCCAGGTGGCGCCGGTCGGGTGCGCCCGTGTCGGCGCGGGG
GGGTTGTGGCGACGAGGCCGCCCTTCAGGCGAGGTGGGAGTGGCCCAAGTTGTGGCGCCG
CGGTGGAGGGGGGGCGGTGAGGTGGGGGGCCAGTCGAGGCTGGTGAGAGCGGGACCAGCC
TCGCTTAGTGCAACGCCAACCTGGATAGTAATTCGTGAGGCTCGTGGGAGCGAGTTCTTC
CAACGATGCAGGGGGGGGTGTGGGGCCCGCGCGTCGACCGGCCGCGTCACGACTGCTACG
GGGTCATTACCGCCGGGCTGTCGTAGGCCGGCGCTCGCGGGAGGTGGTTCGAATGCCGGG
GCGGGAGGCCCGGATCCGGCGAAGGGGCGGGGCTCGGGGGACGCGTGGCGGAGGAGCGCG
AGGCGGTAGCCGGTGCAACCGGCGCGTGGCCCCATGAGGAAGAACTCTCTCGTGCCCCAA
CATCACACATTAGCGCCCCCATCTCGGCCGACTTCGAGCGCGCTCAGCGAGAACCACCAC
AGGCCTCCAACCCGACGTACCACCCGTCCTCACGAGGCGGGCCGACCCTCGGGTCCAAGC
GAGGGCTCCGGCGCGCACGCCCGGCGTGGTGC
>MAP 195_A_e
CCCCTTAGGCGCCTCGAGCGGGCCCCCGCTTCGCCAGTTCACCCGTCCCAGTAAGCGCCC
GGGGCACCTCGCCTTCGAGGAGAACCCCCAGAACGCCCCACCGTCTCCCAGCTCGCCCAC
GCGCACGCGTCGCACTCTCCAGCGCGCGTACTGTTCCAGCCACCGTCCCCATGCTCCCCG
CGGGGCCGTATGACGAGCCACGCCCCGGCCCCGCCGCCGCCGCTCGCCCACCGAGACAAG
CGCAATGCTATTGGGGCCTGACCACGGCTGGGCGGGGCGCTTCCCACGGAGGGCCCAGTC
TTTGGGAGGAGTTGCCCCTCCAGGGATCGGTAGCGCGGTGCGGCTCCCCGCAGCGGCCGC
AGGCGTCGGGGGGATGCTGGGGAGTGGGGGATGCCCCTGCCGGTCGCAGGCGGAACAGGT
TCGCGGTGGTGCACAGGGGCGTGGTGCTGCAGCTCGAGGCCGCGGAGGGGGGCCGCGGTG
GACTTCGGGTCGGAGGGACCGGGATACCAGGGCTCTGGCGCGGGGTAGCCCCAGGACGCT
TTCTCGGGGAGGGGAGGGTACGGGCAGCCAGGTGGGAGTGCCAGAACTGGACGGCGTAAT
CCCTGCTGCCTGCGATGTGGGGTCGGACCCAGCGCACCCTGATGGGGCCTGGCGGCGGGG
GACCCCTGAACGGACGTGGCAGCTACTCTGCCGTTGAGCCGATGCGGACGTGCCCAGTAC
ACGTGGGTGTTGGGGGAACTCCACAGCGGCAGGCGAGGGTTGGGTGTAGCAGGGGGCCCA
CGGGGGCGGCGGGTGGCAGGCGCGAGGGGGGCGCGGGGGAGCGATCCGATGGTCGAGAAC
GATCGGCTAGGAGGCGAGCCCCAGGTGGCACCGGGCGGGTGCGCCCGTGTCGGCGCGGGG
GGGTTGTGGCGACGAGGCCGCCCTTCAGGCGAGGTGGGAGTGGCCCAAGTTGTGGCGCCG
CGGTGGAGGGGGGGCGGTGAGGTGGGGGGCCAGTCGAGGTTGGTGAGGGCGGGACCAGCC
TCGTTTAGTGAAACGCCAACCCGGATAGTAATTCGTGCGGCTCGTGGGAGCGAGTTCTTC
CAATGATGCAGGGGGGGGTGTGGGGCCCGCGCGTCGACCGGTCGCGTCACGACTGCTACG
GGGTCATTACCGCCGGGCTGTCGTAGGCCGGCGCTCGCGGAGGGAGGTTCGAATGCCGGG
GCGGGAGGCCCGGATCCGGCGAAGGGGCGGGGCTCGGGGGACGCGAGGCGGAGGGGCGCG

AGGCGGTAGCCGGTGCAAGCGGCGCGTGGCCCCATGAGGAAGAACTCTCTCGTGCCCCAA
CACCACACATTAGCGCCCCCATCTCGGCCGACTTCGAGCACGCTCAACGAGAACCACCAC
AGGCCTCCAACCCGACGTACCACCCGTCCTCACGAGGCGGGCCGACCCTCGGGTCCAAGC
GAGGGCTCCGCCGCGCACGCCCGGCGTGGTGC
>MAP 196_A_e
CCCCTTAGGCGCCTCGAGCGGGCCCCCGCTTCGCCAGTTCACCCGTCCCAGTAAGCGCCC
GGGGCACCTCGCCTTCGAGGAGAACCCCCAGAACGCCCCACCGTCTCCCAGCTCGCACAC
GCGCACGCGTCGCACTCTCCAGCGCGCGTACTGTTCCAGCCACCGTCCCCATGCTCCCCG
CGGGGCCGTATGACGAGCCACGCCCCGGCCCCGCCGCCGCCGCTCGCCCACCGAGACAAG
CGCAATGCTATTGGGGCCTGACCACGGCTGGGCGGGGCGCTTCCCACGGAGGGCCCAGTC
TTTGGGAGGAGTTGCCCCTCCAGGGATCGGTAGCGCGGTGCGGCTCCCCGCAGCGGCCGC
AGGCGTCGGGGGGATGCTGGGGAGTGGGGGATGCCCCTGCCGGTCGCAGGCGGAACAGGT
TCGCGGTGGTGCACAGGGGCGTGGTGCTGCAGCTCGAGGCCGCGGAGGGGGGGCCGCGGTG
GACTTCGGGTCGGAGGGACCGGGATACCAGGGCTCTGGCGCGGGGTAGCCCCAGGACGCT
TTCTCGGGGAGGGGAGGGTACGGGCAGCCAGGTGGGAGTGCCAGAACTGGACGGCGTAAT
CCCTGCTGCCTGCGATGTGGGGTCGGACCCAGCGCACCCTGATGGGGCCTGGCGGCGGGG
GACCCCTGAACGGACGTGGCAGCTACTCTGCCGTTGAGCCGATGCGGACGTGCCCAGTAC
ACGTGGGTGTTGGGGGAACTCCACAGCGGCAGGCGAGGGTTGGGTGTAGCAGGGGGCCCA
CGGGGGCGGCGGGTGGCAGGCGCGAGGGGGGCGCGGGGGAGCGATCCGATGGTCGAGAAC
GATCGGCTAGGAGGCGAGCCCCAGGTGGCACCGGGCGGGTGCGCCCGTGTCGGCGCGGGG
GGGTTGTGGCGACGAGGCCGCCCTTCAGGCGAGGTGGGAGTGGCCCAAGTTGTGGCGCCG
CGGTGGAGGGGGGGCGGTGAGGTGGGGGGCCAGTCGAGGTTGGTAAGGGCGGGACCAGCC
TCGTTTAGTGAAACGCCAACCCGGATAGTAATTCGTGCGGCTCGTGGGAGCGAGTTCTTC
CAATGATGCAGGGGGGGGTGTGGGGCCCGCGCGTCGACCGGTCGCGTCACGACTGCTACG
GGGTCATTACCGCCGGGCTGTCGTAGGCCGGCGCTCGCGGAGGGAGGTTCGAATGCCGGG
GCGGGAGGCCCGGATCCGGCGAAGGGGCGGGGCTCGGGGGACGCGAGGCGGAGGGGCGCG
AGGCGGTAGCCGGTGCAAGCGGCGCGTGGCCCCATGAGGAAGAACTCTCTCGTGCCCCAA
CACCACACATTAGCGCCCCCATCTCGGCCGACTTCGAGCACGCTCAACGAGAACCACCAC
AGGCCTCCAACCCGACGTACCACCCGTCCTCACGAGGCGGGCCGACCCTCGGGTCCAAGC
GAGGGCTCCGCCGCGCACGCCCGGCGTGGTGC
>MAP 197_A_e
CCCCTTAGGCGCCTCGAGCGGGCCCCCGCTTCGCCAGTTCACCCGTCCCAGTAAGCGCCC
GGGGCACCTCGCCTTCGAGGAGAACCCCCAGAACGCCCCACCGTCTCCCAGCTCGCACAC
GCGCACGCGTCGCACTCTCCAGCGCGCGTACTGTTCCAGCCACCGTCCCCATGCTCCCCG
CGGGGCCGTATGACGAGCCACGCCCCGGCCCCGCCGCCGCCGCTCGCCCACCGAGACAAG
CGCAATGCTATTGGGGCCTGACCACGGCTGGGCGGGGCGCTTCCCACGGAGGGCCCAGTC
TTTGGGAGGAGTTGCCCCTCCAGGGATCGGTAGCGCGGTGCGGCTCCCCGCAGCGGCCGC
AGGCGTCGGGGGGATGCTGGGGAGTGGGGGATGCCCCTGCCGGTCGCAGGCGGAACAGGT
TCGCGGTGGTGCACAGGGGCGTGGTGCTGCAGCTCGAGGCCGCGGAGGGGGGGCCGCGGTG
GACTTCGGGTCGGAGGGACCGGGATACCAGGGCTCTGGCGCGGGGTAGCCCCAGGACGCT
TTCTCGGGGAGGGGAGGGTACGGGCAGCCAGGTGGGAGTGCCAGAACTGGACGGCGTAAT
CCCTGCTGCCTGCGATGTGGGGTCGGACCCAGCGCACCCTGATGGGGCCTGGCGGCGGGG
GACCCCTGAACGGACGTGGCAGCTACTCTGCCGTTGAGCCGATGCGGACGTGCCCAGTAC
ACGTGGGTGTTGGGGGAACTCCACAGCGGCAGGCGAGGGTTGGGTGTAGCAGGGGGCCCA
CGGGGGCGGCGGGTGGCAGGCGCGAGGGGGGCGCGGGGGAGCGATCCGATGGTCGAGAAC
GATCGGCTAGGAGGCGAGCCCCAGGTGGCACCGGGCGGGTGCGCCCGTGTCGGCGCGGGG
GGGTTGTGGCGACGAGGCCGCCCTTCAGGCGAGGTGGGAGTGGCCCAAGTTGTGGCGCCG
CGGTGGAGGGGGGGCGGTGAGGTGGGGGGCCAGTCGAGGTTGGTGAGGGCGGGACCAGCC
TCGTTTAGTGAAACGCCAACCCGGATAGTAATTCGTGCGGCTCGTGGGAGCGAGTTCTTC
CAATGATGCAGGGGGGGGTGTGGGGCCCGCGCGTCGACCGGTCGCGTCACGACTGCTACG
GGGTCATTACCGCCGGGCTGTCGTAGGCCGGCGCTCGCGGAGGGAGGTTCGAATGCCGGG
GCGGGAGGCCCGGATCCGGCGAAGGGGCGGGGCTCGGGGGACGCGAGGCGGAGGGGCGCG
AGGCGGTAGCCGGTGCAAGCGGCGCGTGGCCCCATGAGGAAGAACTCTCTCGTGCCCCAA
CACCACACATTAGCGCCCCCATCTCGGCCGACTTCGAGCACGCTCAACGAGAACCACCAC
AGGCCTCCAACCCGACGTACCACCCGTCCTCACGAGGCGGGCCGACCCTCGGGTCCAAGC
GAGGGCTCCGCCGCGCACGCCCGGCGTGGTGC
>MAP 198_A_e
CCCCTCAGGCGCCCCAAGCGGGCCCCCGCTTCGCCAGCTCACCCGTTCCAGTAACTGCCC
ACGGCCCCTCGCCTTCGCGGGGGACCCCCAGAACGCCCCACCGTCTCCCAGCTCGCACGC
GCACACGCGTCGCACCCTCCAGCGCGCGGACTGCTCCAGCAGTCATCCCCATGCTCCCCG
CGGGGCTGCACGACGAGCCACGCCTCGGCACCGCCGCTACCGCCCACCCATCGAGACAAG
CGCAACGCTATTGGGGCCTAACCACGGCTGGGTGGGACGCTTCCCACGGAGGGCTCAGTC
CTCGGGAGGAGCTGCCCCTCAAGGGATCGGTAGCGCGGTGCGGTTCCCCGCAGCGGCCGT
GGGCGTCGGGGTGATCCCGAGGAGTGGGGGATGCCCCTTCCGGCAGAGGGCGGAGCGGGT
TCGCAGTGGTGCACAGGGGCGTGGTGCCGCGGCCTGGGGCGGCGGCCGGGGGCCGCGGTG
GACTTCGGGTCGGATGGACCGATATAGCACGGCTCTGGCGCGGCGTGGCCCCAGGACGCT
CTCTCGGGGACGGGAGGGTACGGTGAACCAGGTGGGAGCGCAGGAGCTGGACGGCGTAAT
CCCTGGTGACTGCGACGTGGGGTAGGACCCAGGGCACCCTGGCGATGCCTGGCGGCGGGG

GCCCCCCGGACGGACGCGCCGGCTACCCTGCTGTTGGGACGGTGCGGACGTACCCAGTAG
ACGTGGGTATTGGGGGAACTCTGGAGCGGCAGGCGAGGGTTAGGTGTGAGCGGGGGCCCC
CGGGGGCGGCGGGGGGCGGTCGCGAGGGGGGCGCGAGGAAGTGATGCGATGGTCGAGAAC
AATCGGCCAGGAGGCGAGCCCCAGCTGCCGCAGGGCGGGTGCGCCCGTGTCGGGGCGGGG
GGGTTGTGGCGACGAGGCCGCCCCTCAGGCGGGTCGGGAGTGGCCCAAGTTGTGGCGCTG
CGGTGGAGGGGGGGCAGTGAGGCGGGGGGTGAGTCGGGGTTCGTGCAGGCGGGACCAGCC
TCGCTTGGCGCATGGCCAAGCCGGATAGTAATTCGTGCGACTCGCAGGCACGAGTTCTTC
CAATGATTCAGGGGGGGGTGTGAGGCCCCCGCGTCGGTGGGCCGTGTCACGAGTGCCGCA
GGGTCAGTACCGCCGGACCACCGTAGGCCGGCAATTGCGGGAGGTGGTTCGAATGCCGGG
CCGGGAGGCCCGGATTAGGCGGAGGGGCGGGGCCCGGGGCACGCGAGGCACGGGGGCGCG
AAGCGGTAGCCGGTGCAAGCGGCCCGTGGCCCCATGAGGTAGAACTCTCTCGTGCCCCAA
CACCCCACATTGGCGCCCACGCCTCGTCCGACCTCAAGCGTACTCAGCTGGAACCGCCAC
AGGCCTCTAACCCTACGTTCCACCCGTCCTCACGAGGCGGTCCGACCCCCGGGCCCAAGG
GAGGGCTTCGGCGAGCACGCCCGGCATGGCGC
>MAP 199_A_e
CCCCTTAGGCGCCTCGAGCGGGCCCCCGCTTCGCCAGTTCACCCGTCCCAGTAAGCGCCC
GGGGCACCTCGCCTTCGAGGAGAACCCCCAGAACGCCCCACCGTCTCCCAGCTCGCACAC
GCGCACGCGTCGCACTCTCCAGCGCGCGTACTGTTCCAGCCACCGTCCCCATGCTCCCCG
CGGGGCCGTATGACGAGCCACGCCCCGGCCCCGCCGCCGCCGCTCGCCCACCGAGACAAG
CGCAATGCTATTGGGGCCTGACCACGGCTGGGCGGGGCGCTTCCCACGGAGGGCCCAGTC
TTTGGGAGGAGTTGCCCCTCCAGGGATCGGTAGCGCGGTGCGGCTCCCCGCAGCGGCCGC
AGGCGTCGGGGGGATGCTGGGGAGTGGGGGATGCCCCTGCCGGTCGCAGGCGGAACAGGT
TCGCGGTGGTGCACAGGGGCGTGGTGCTGCAGCTCGAGGCCGCGGAGGGGGGCCGCGGTG
GACTTCGGGTCGGAGGGACCGGGATACCAGGGCTCTGGCGCGGGGTAGCCCCAGGACGCT
TTCTCGGGGAGGGGAGGGTACGGGCAGCCAGGTGGGAGTGCCAGAACTGGACGGCGTAAT
CCCTGCTGCCTGCGATGTGGGGTCGGACCCAGCGCACCCTGATGGGGCCTGGCGGCGGGG
GACCCCTGAACGGACGTGGCAGCTACTCTGCCGTTGAGCCGATGCGGACGTGCCCAGTAC
ACGTGGGTGTTGGGGGAACTCCACAGCGGCAGGCGAGGGTTGGGTGTAGCAGGGGGCCCA
CGGGGGCGGCGGGTGGCAGGCGCGAGGGGGGCGCGGGGGAGCGATCCGATGGTCGAGAAC
GATCGGCTAGGAGGCGAGCCCCAGGTGGCACCGGGCGGGTGCGCCCGTGTCGGCGCGGGG
GGGTTGTGGCGACGAGGCCGCCCTTCAGGCGAGGTGGGAGTGGCCCAAGTTGTGGCGCCG
CGGTGGAGGGGGGGCGGTGAGGTGGGGGGCCAGTCGAGGTTGGTGAGGGCGGGACCAGCC
TCGTTTAGTGAAACGCCAACCCGGATAGTAATTCGTGCGGCTCGTGGGAGCGAGTTCTTC
CAATGATGCAGGGGGGGGTGTGGGGCCCGCGCGTCGACCGGTCGCGTCACGACTGCTACG
GGGTCATTACCGCCGGGCTGTCGTAGGCCGGCGCTCGCGGAGGGAGGTTCGAATGCCGGG
GCGGGAGGCCCGGATCCGGCGAAGGGGCGGGGCTCGGGGGACGCGAGGCGGAGGGGCGCG
AGGCGGTAGCCGGTGCAAGCGGCGCGTGGCCCCATGAGGAAGAACTCTCTCGTGCCCCAA
CACCACACATTAGCGCCCCCATCTCGGCCGACTTCGAGCACGCTCAACGAGAACCACCAC
AGGCCTCCAACCCGACGTACCACCCGTCCTCACGAGGCGGGCCGACCCTCGGGTCCAAGC
GAGGGCTCCGCCGCGCACGCCCGGCGTGGTGC
>MAP 200_A_e
CCCCTTAGGCGCCTCGAGCGGGCCCCCGCTTCGCCAGTTCACCCGTCCCAGTAAGCGCCC
GGGGCACCTCGCCTTCGAGGAGAACCCCCAGAACGCCCCACCGTCTCCCAGCTCGCACAC
GCGCACGCGTCGCACTCTCCAGCGCGCGTACTGTTCCAGCCACCGTCCCCATGCTCCCCG
CGGGGCCGTATGACGAGCCACGCCCCGGCCCCGCCGCCGCCGCTCGCCCACCGAGACAAG
CGCAATGCTATTGGGGCCTGACCACGGCTGGGCGGGGCGCTTCCCACGGAGGGCCCAGTC
TTTGGGAGGAGTTGCCCCTCCAGGGATCGGTAGCGCGGTGCGGCTCCCCGCAGCGGCCGC
AGGCGTCGGGGGGATGCTGGGGAGTGGGGGATGCCCCTGCCGGTCGCAGGCGGAACAGGT
TCGCGGTGGTGCACAGGGGCGTGGTGCTGCAGCTCGAGGCCGCGGAGGGGGGCCGCGGTG
GACTTCGGGTCGGAGGGACCGGGATACCAGGGCTCTGGCGCGGGGTAGCCCCAGGACGCT
TTCTCGGGGAGGGGAGGGTACGGGCAGCCAGGTGGGAGTGCCAGAACTGGACGGCGTAAT
CCCTGCTGCCTGCGATGTGGGGTCGGACCCAGCGCACCCTGATGGGGCCTGGCGGCGGGG
GACCCCTGAACGGACGTGGCAGCTACTCTGCCGTTGAGCCGATGCGGACGTGCCCAGTAC
ACGTGGGTGTTGGGGGAACTCCACAGCGGCAGGCGAGGGTTGGGTGTAGCAGGGGGCCCA
CGGGGGCGGCGGGTGGCAGGCGCGAGGGGGGCGCGGGGGAGCGATCCGATGGTCGAGAAC
GATCGGCTAGGAGGCGAGCCCCAGGTGGCACCGGGCGGGTGCGCCCGTGTCGGCGCGGGG
GGGTTGTGGCGACGAGGCCGCCCTTCAGGCGAGGTGGGAGTGGCCCAAGTTGTGGCGCCG
CGGTGGAGGGGGGGCGGTGAGGTGGGGGGCCAGTCGAGGTTGGTAAGGGCGGGACCAGCC
TCGTTTAGTGAAACGCCAACCCGGATAGTAATTCGTGCGGCTCGTGGGAGCGAGTTCTTC
CAATGATGCAGGGGGGGGTGTGGGGCCCGCGCGTCGACCGGTCGCGTCACGACTGCTACG
GGGTCATTACCGCCGGGCTGTCGTAGGCCGGCGCTCGCGGAGGGAGGTTCGAATGCCGGG
GCGGGAGGCCCGGATCCGGCGAAGGGGCGGGGCTCGGGGGACGCGAGGCGGAGGGGCGCG
AGGCGGTAGCCGGTGCAAGCGGCGCGTGGCCCCATGAGGAAGAACTCTCTCGTGCCCCAA
CACTACACATTAGCGCCCCCATCTCGGCCGACTTCGAGCACGCTCAACGAGAACCACCAC
AGGCCTCCAACCCGACGTACCACCCGTCCTCACGAGGCGGGCCGACCCTCGGGTCCAAGC
GAGGGCTCCGCCGCGCACGCCCGGCGTGGTGC
>MAP 201_A_e
CCCCTTAGGCGCCTCGAGCGGGCCCCCGCTTCGCCAGTTCACCCGTCCCAGTAAGCGCCC

GGGGCACCTCGCCTTCGAGGAGAACCCCCAGAACGCCCCACCGTCTCCCAGCTCGCACAC
GCGCACGCGTCGCACTCTCCAGCGCGCGTACTGTTCCAGCCACCGTCCCCATGCTCCCCG
CGGGGCCGTATGACGAGCCACGCCCCGGCCCCGCCGCCGCCGCTCGCCCACCGAGACAAG
CGCAATGCTATTGGGGCCTGACCACGGCTGGGCGGGGCGCTTCCCACGGAGGGCCCAGTC
TTTGGGAGGAGTTGCCCCTCCAGGGATCGGTAGCGCGGTGCGGCTCCCCGCAGCGGCCGC
AGGCGTCGGGGGGATGCTGGGGAGTGGGGGATGCCCCTGCCGGTCGCAGGCGGAACAGGT
TCGCGGTGGTGCACAGGGGCGTGGTGCTGCAGCTCGAGGCCGCGGAGGGGGGCCGCGGTG
GACTTCGGGTCGGAGGGACCGGGATACCAGGGCTCTGGCGCGGGGTAGCCCCAGGACGCT
TTCTCGGGGAGGGGAGGGTACGGGCAGCCAGGTGGGAGTGCCAGAACTGGACGGCGTAAT
CCCTGCTGCCTGCGATGTGGGGTCGGACCCAGCGCACCCTGATGGGGCCTGGCGGCGGGG
GACCCCTGAACGGACGTGGCAGCTACTCTGCCGTTGAGCCGATGCGGACGTGCCCAGTAC
ACGTGGGTGTTGGGGGAACTCCACAGCGGCAGGCGAGGGTTGGGTGTAGCAGGGGGCCCA
CGGGGGCGGCGGGTGGCAGGCGCGAGGGGGGCGCGGGGGAGCGATCCGATGGTCGAGAAC
GATCGGCTAGGAGGCGAGCCCCAGGTGGCACCGGGCGGGTGCGCCCGTGTCGGCGCGGGG
GGGTTGTGGCGACGAGGCCGCCCTTCAGGCGAGGTGGGAGTGGCCCAAGTTGTGGCGCCG
CGGTGGAGGGGGGGCGGTGAGGTGGGGGGCCAGTCGAGGTTGGTGAGGGCGGGACCAGCC
TCGTTTAGTGAAACGCCAACCCGGATAGTAATTCGTGCGGCTCGTGGGAGCGAGTTCTTC
CAATGATGCAGGGGGGGGTGTGGGGCCCGCGCGTCGACCGGTCGCGTCACGACTGCTACG
GGGTCATTACCGCCGGGCTGTCGTAGGCCGGCGCTCGCGGAGGGAGGTTCGAATGCCGGG
GCGGGAGGCCCGGATCCGGCGAAGGGGCGGGGCTCGGGGGACGCGAGGCGGAGGGGCGCG
AGGCGGTAGCCGGTGCAAGCGGCGCGTGGCCCCATGAGGAAGAACTCTCTCGTGCCCCAA
CACCACACATTAGCGCCCCCATCTCGGCCGACTTCGAGCACGCTCAACGAGAACCACCAC
AGGCCTCCAACCCGACGTACCACCCGTCCTCACGAGGCGGGCCGACCCTCGGGTCCAAGC
GAGGGCTCCGCCGCGCACGCCCGGCGTGGTGC
>MAP 213_A_e
CCCCTTAGGCGCCTCGAGCGGGCCCCCGCTTCGCCAGTTCAGCCGTCCCAGTAAGCGCCC
GGGGCACCTCGCCTTCGAGGAGAACCCCCAGAACGCCCCACCGTCTCCCAGCTCGCACAC
GCGCACGCGTCGCACTCTCCCGCGCGCGTACTGTTCCAGCCACCGTCCCCATGCTCCCCG
CGGGGCCGTATGACGAGCCACGCCCCGGCCCCGCCGCCGCCGCTCGCTCACCGAGACCAG
CGCAATGCTATTGGGGCCTGACCACGGCTGGGCGGGGCGCTTCCCACGGAGGGCCCAGTC
TTTGGGAGGAGTTGCCCCTCCAGGGATCGGCAGCGCGGTGCGGCTCCCCGCAGCGGCCGC
GGGCGTCGGGGGGATGCTAGGGAGTGGGGGATGCCCCTTCCGGTCGCAGGCGGAACAGGT
TCGCGGTGGTGCACAGGGGCGTGGTGCTGCAGCTCGAGGCCGCGGAGGGGGGCCGCGGTG
GACTTCGGGTCGGAGGGACCGATATACCAGGGCTCTGGCGCGGGGTAGCCCCAGGACGCT
TTCTCGGGGAGGGGAGGGTACGGGCAGCCAGGTGGGAGTGCAAGAACTGGACGGCCTAAT
CCCTGCTGCCTGCGATGTGGGGTCGGACCCAGCGCACCCTGATGGGGCCTGGCGGCGGGG
GACCCCTGAACGGACGTGGCAGCTACTCTGCCGTTGAGCCGATGCGGACGTGCCCAGTAC
ACGTGGGTGTTGGGGGAACTCCACAGCGGCAGGCGAGGGTTGGGTGTAGCAGGGGGCCCA
CGGGGGCGGCGGGTGGCAGGCGCGAGGGGGGCGCGGGGGAGCGATCCGATGGTCGAGAAC
GATCGGCTAGGAGGCGAGCCCCAGGTGGCGCCGGGCGGGTGCGCCCGTGTCGGCGCGGGG
GGGTTGTGGCGACGAGGCCGCCCTTCAGGCGAGGTGGGAGTGGCCCACGTTGTGGCGCCG
CGGTGGAGGGGGGGCGGTGAGGTGGGGGGCCAGTCGAGGTTGGTGAGGGCGGGACCAGCC
TCGCTTAGTGCAACGCCAACCCGGATAGTAATTCGTGCGGCTCGTGGGAGCGAGTTCTTC
CAATGATGCAGGGGGGGGTGTGGGGCCCGCGCGTCGACCGGCCGCGTCACGACTGCTACG
GGGTCATTACCGCCGGGCTGTCGTAGGCCGGCGCTCGCGGGAGGTGGTTCGAATGCCGGG
GCGGGAGGCCCGGATCCGGCCAAGGGGCGGGGCTCGGGGGACGCGAGGCGGAGGGGCGCG
AGGCGGTAGCCGGTGCAAGCGGCGCGTGGCCCCATGAGGAAGAACTCTCTCGTGCCCCAA
CACCACACATTAGCGCCCCCATCTCGGCCGACTTCGAGCACGCTCAGCGAGAACCACCAC
AGGCCTCCAACCCGACGTACCACCCGTCCTCACGAGGCGGGCCGACCCTCGGGTCCAAGC
GAGGGCTCCGGCGCGCACGCCCGGCGTGGTGC
>MAP 221_A_e
CCCCTTAGGCGCCTCGAGCGGGCCCCCGCTTCGCCAGTTCACCCGTCCCAGTAAGCGCCC
GGGGCACCTCGCCTTCGAGGAGAACCCCCAGAACGCCCCACCGTCTCCCAGCTCGCACAC
GCGCACGCGTCGCACTCTCCAGCGCGCGTACTGTTCCAGCCACCGTCCCCATGCTCCCCG
CGGGGCCGTATGACGAGCCACGCCCCGGCCCCGCCGCCGCCGCTCGCCCACCGAGACAAG
CGCAATGCTATTGGGGCCTGACCACGGCTGGGCGGGGCGCTTCCCACGGAGGGCCCAGTC
TTTGGGAGGAGTTGCCCCTCCAGGGATCGGTAGCGCGGTGCGGCTCCCCGCAGCGGCCGC
AGGCGTCGGGGGGATGCTGGGGAGTGGGGGATGCCCCTGCCGGTCGCAGGCGGAACAGGT
TCGCGGTGGTGCACAGGGGCGTGGTGCTGCAGCTCGAGGCCGCGGAGGGGGGCCGCGGTG
GACTTCGGGTCGGAGGGACCGGGATACCAGGGCTCTGGCGCGGGGTAGCCCCAGGACGCT
TTCTCGGGGAGGGGAGGGTACGGGCAGCCAGGTGGGAGTGCCAGAACTGGACGGCGTAAT
CCCTGCTGCCTGCGATGTGGGGTCGGACCCAGCGCACCCTGATGGGGCCTGGCGGCGGGG
GACCCCTGAACGGACGTGGCAGCTACTCTGCCGTTGAGCCGATGCGGACGTGCCCAGTAC
ACGTGGGTGTTGGGGGAACTCCACAGCGGCAGGCGAGGGTTGGGTGTAGCAGGGGGCCCA
CGGGGGCGGCGGGTGGCAGGCGCGAGGGGGGCGCGGGGGAGCGATCCGATGGTCGAGAAC
GATCGGCTAGGAGGCGAGCCCCAGGTGGCACCGGGCGGGTGCGCCCGTGTCGGCGCGGGG
GGGTTGTGGCGACGAGGCCGCCCTTCAGGCGAGGTGGGAGTGGCCCAAGTTGTGGCGCCG
CGGTGGAGGGGGGGCGGTGAGGTGGGGGGCCAGTCGAGGTTGGTGAGGGCGGGACCAGCC

TCGTTTAGTGAAACGCCAACCCGGATAGTAATTCGTGCGGCTCGTGGGAGCGAGTTCTTC
CAATGATGCAGGGGGGGGTGTGGGGCCCGCGCGTCGACCGGTCGCGTCACGACTGCTACG
GGGTCATTACCGCCGGGCTGTCGTAGGCCGGCGCTCGCGGAGGGAGGTTCGAATGCCGGG
GCGGGAGGCCCGGATCCGGCGAAGGGGCGGGGCTCGGGGGACGCGAGGCGGAGGGGCGCG
AGGCGGTAGCCGGTGCAAGCGGCGCGTGGCCCCATGAGGAAGAACTCTCTCGTGCCCCAA
CACCACACATTAGCGCCCCCATCTCGGCCGACTTCGAGCACGCTCAACGAGAACCACCAC
AGGCCTCCAACCCGACGTACCACCCGTCCTCACGAGGCGGGCCGACCCTCGGGTCCAAGC
GAGGGCTCCGCCGCGCACGCCCGGCGTGGTGC
>MAP 222_A_e
CCCCTTAGGCGCCTCGAGCAGGCCCCCGCCTCGCCAGTTCACCCGTCCCAGTAAGCGCCT
GCGGCCCCTCGCCTTCGCGGAGAACCCCCAGAACGCCCCACCGTATCCCTGCTCGCACAC
GCGCACGCGTCGCACTCTCCAGCGCGCGGACTGTTCCAGCCACCGTCCCCATGCTCCCCG
CGGGGCCGTATGACGAGCCACGCCCCGGCCCCGCCGCCGCCGCTCGCCCACCGAGGCAAG
CGCAACGCTATTGGGGCCTGACCACGGCGGGGCGGGGCGCTTCCCACGGAGAACCCAGCC
TTTGGGAGGAGTTGCCCCTCAGGGGATCGGTAGCGCGGTGCGGCTCCCCGCAGCGGCCGC
GGGCGTCGGGGTGATGCTGGGGAGTGGGGGATGCCCCTTCCGGCCGCAGGCGGAACAGGT
TCGCGGTGGTGCACAGGGGCATGGTGCTGCAGCTCGAGGCCGCAGAGGGGGGCCGCGGTG
GACTTCGGGTCGGAGGGACCGATATACCAGGGCTCTGGCGCGAGGTAGCCCCAGGACGCT
CTCCCGGGGAGGGGAGAGTACGGGCAGCCAGGTGGGAGTGCAGAAACTGGACGGCGTAAT
CCCTGGCGACTGCGATGTTGGGGAGGACCCAGCGCACCCTGATGGGGCCTGACGGCGGGG
GATCCCTGGACCGACGTGGCAGCTACCCTGCCGTTGGGCCGATGCGGACGTGCCCAGTAC
ACGTGGGTGTTGGGGGAACTCCACAGCGGCGGGCGAGTGTTGGGTGTGGCCGGGGGCCCA
CAGGGGCGAGGGGTGACGGTCGCGAGGGGGGCGTGGGGGAGCGATCCGATGGTCGAGAAC
GATCGGCTAGGAGGCGAGACCCAGGCGGCGCAGGGCGGGTGCGCCCGTGTCGGCGCGGGG
GGGTTGTGGCGACGAGGCCGCCCTTCAGGCGGGGTGGGAGTGGCCCAAGTTGTGGCGCCG
CGGCGGAGGGGGGGCGGTGAGGTGGGGGGCCAGTCGGGGTTGGTGAGGATGGGACCAGCC
TTGCTTGGTGCAACGCCAACCCGGATAGTAATTCGTGCGGCTCGTGGGCGCGGGTTGCTC
CAATGGTTCAGGGGGGGGTGTGGGGCCCGCGCGTCGACCGGCCGCGTCACGACTGCTACG
AGGTCATTACCGCCGGGCTGTCGTAGGCCGGCGATCGCGGGAAGTGGTTCGAATGCCGGG
GCCGGAGGCCCGAATCAGGCGAAGGGGCGGGGTCCGGGGCACGCGAGGCGGGGGGGCGCG
AGGCGGTAGCCGGTGCAAGCGGCGCCTAGCCCCATGAGATAGAACTCTCTCGTCCCCCAA
CACCACACACTAGCGCCTCCATCTCGGCCGACTTCGAGCGCGCTCAGGGAGAACCACCAC
AGGCTCCCGACCCGACGTACCACCCGTCCTCACGAGGCGGGCCGACCCCCGGGCCCAAGC
GAGAGCTCCGGCGAGCACGCCCGGCGTGGTAC
>MAP 223_A_e
CCCCTTAGGCGCCTCGAGCGGGCCCCCGCTTCGCCAGTTCAGCCGTCCCAGTAAGCGCCC
GGGGCACCTCGCCTTCGAGGAGAACCCCCAGAACGCCCCACCGTCTCCCAGCTCGCACAC
GCGCACGCGTCGCACTCTCCCGCGCGCGTACTGTTCCAGCCACCGTCCCCATGCTCCCCG
CGGGGCCGTATGACGAGCCACGCCCCGGCCCCGCCGCCGCCGCTCGCTCACCGAGACCAG
CGCAATGCTATTGGGGCCTGACCACGGCTGGGCGGGGCGCTTCCCACGGAGGGCCCAGTC
TTTGGGAGGAGTTGCCCCTCCAGGGATCGGCAGCGCGGTGCGGCTCCCCGCAGCGGCCGC
GGGCGTCGGGGGGATGCTAGGGAGTGGGGGATGCCCCTTCCGGTCGCAGGCGGAACAGGT
TCGCGGTGGTGCACAGGGGCGTGGTGCTGCAGCTCGAGGCCGCGGAGGGGGGCCGCGGTG
GACTTCGGGTCGGAGGGACCGATATACCAGGGCTCTGGCGCGGGGTAGCCCCAGGACGCT
TTCTCGGGGAGGGGAGGGTACGGGCAGCCAGGTGGGAGTGCAAGAACTGGACGGCCTAAT
CCCTGCTGCCTGCGATGTGGGGTCGGACCCAGCGCACCCTGATGGGGCCTGGCGGCGGGG
GACCCCTGAACGGACGTGGCAGCTACTCTGCCGTTGAGCCGATGCGGACGTGCCCAGTAC
ACGTGGGTGTTGGGGGAACTCCACAGCGGCAGGCGAGGGTTGGGTGTAGCAGGGGGCCCA
CGGGGGCGGCGGGTGGCAGGCGCGAGGGGGGCGCGGGGGAGCGATCCGATGGTCGAGAAC
GATCGGCTAGGAGGCGAGCCCCAGGTGGCGCCGGGCGGGTGCGCCCGTGTCGGCGCGGGG
GGGTTGTGGCGACGAGGCCGCCCTTCAGGCGAGGTGGGAGTGGCCCACGTTGTGGCGCCG
CGGTGGAGGGGGGGCGGTGAGGTGGGGGGCCAGTCGAGGTTGGTGAGGGCGGGACCAGCC
TCGCTTAGTGCAACGCCAACCCGGATAGTAATTCGTGCGGCTCGTGGGAGCGAGTTCTTC
CAATGATGCAGGGGGGGGTGTGGGGCCCGCGCGTCGACCGGCCGCGTCACGACTGCTACG
GGGTCATTACCGCCGGGCTGTCGTAGGCCGGCGCTCGCGGGAGGTGGTTCGAATGCCGGG
GCGGGAGGCCCGGATCCGGCCAAGGGGCGGGGCTCGGGGGACGCGAGGCGGAGGGGCGCG
AGGCGGTAGCCGGTGCAAGCGGCGCGTGGCCCCATGAGGAAGAACTCTCTCGTGCCCCAA
CACCACACATTAGCGCCCCCATCTCGGCCGACTTCGAGCACGCTCAGCGAGAACCACCAC
AGGCCTCCAACCCGACGTACCACCCGTCCTCACGAGGCGGGCCGACCCTCGGGTCCAAGC
GAGGGCTCCGGCGCGCACGCCCGGCGTGGTGC
>MAP 172_A_tb1171
CCCCTTAGGCGCCTCGAGCGGGCCTCCGCTTCGCCAGTTCACCCGTCCCAGTAAGCGCCC
GGGGCACCTCGCCTTCGAGGAGAACCCCCAGAACGCCCCACCGTCTCCCAGCTCGCACAC
GCGCACGCGTCGCACTCTCCAGCGCGCGTACTGTTCCAGCCACCGTCCCCATGCTCCCCG
CGGGGCCGTATGACGAGCCACGCCCCGGCCCCGCCGCCGCCGCTCGCCCACCGAGACAAG
CGCAATGCTATTGGGGCCTGACCACGGCTGGGCGGGGCGCTTCCCACGGAGGGCCCAGTC
TTTGGGGGGAGTTGCCCTTCCAGGGATCGGTAGCGCGGTGCGGCTCCCCGCAGCGGCCGC
GGGCGTCGGGGGGATGCTGGGGAGTGGGGGATGCCCCTTCCGGTCGCAGGCGGAACAGGT

TCGCGGTGGTGCACAGGGGCGTGGTGCTGCAGCTCGAGGCCGCGGAGGGGGGCCGTGGTG
GACTTCGGGTCGGAGGGACCGATATACCAGGGCTCTGGCGCGGGGTAGCCCCAGGACGCT
TTCTCGGGGAGGGGAGGGTACGGGCAGCCAGGTGGGAGTGCAAGAACTGGACGGCGTAAT
CCCTGCTGCCCGCGATGTGGGGTCGGACCCAGCGCACCCCGATGGGGCCTGGCGGCGGGG
GACCCCTGAACGGACGTGGCAGCTACTCTGCCGTTGAGCCGATGCGGACGTGCCCAGTAC
ACGTGGGTGTTGGGGGAACTCCACAGCGGCAGGCGAGGGTTGGGTGTAGCAGGGGGCCCA
CGGGGGCGGCGGGTGGCAGGCGCGAGGGGGGCGCGGGGGAGCGATCCGATGGTCGAGAAC
GATCGGCTAGGAGGCGAGCCCCAGGTGGCGCCGGGCGGGTGCGCCCGTGTCGGCGCGGGG
GGGTTGTGGCGACGAGGCCGCCCTTCAGGCGAGGTGGGAGTGGCCCAAGTTGTGGCGCCG
CGGTGGAGGGGGGGCGGTGAGGTGGGGGGCCAGTCGAGGTTGGTGAGGGCGGGACCGGCC
TCGCTTAGTGCAACACCAACCCGGATAGTAATTCGTGCGGCTCGTGGGAGCGAGTTCTTC
CAATGATGCAGGGGGGGGTGTAGGGCCCGCGCGTCGACCGGCCGCGTCACGACTGCTACG
GGGTCATTACCGCCGGGCTGTCGTAGGCCGGCGCTCGCGGGAGGTGGTTCGAATGCCGGG
GCGGGAGGCCCGGATCCGGCGAAGGGGCGGGGCTCGGGGGACGCGAGGCGGAGGGGCGCG
AGGCGGTAGCCGGTGCAAGCGGCGCGTGGCCCCATGAGGAAGAACTCTCTCGTGCCCCAA
CACCACACATTAGCGCCCCCATCTCGGCCGACTTCGAGCACGCTCAGCGAGAACCACCAC
AGGCCTCCAACCCGACGTACCACCCGTTCTCACGAGGCGGGCCGACCCTCGGGTCCAAGC
GAGGGCTCCGGCGCGCACGCCCGGCGTGGTGC
>MAP 173_A_tb1171
CCCCTTAGGCGCCTCGAGCGGGCCTCCGCTTCGCCAGTTCACCCGTCCCAGTAAGCGCCC
GGGGCACCTCGCCTTCGAGGAGAACCCCCAGAACGCCCCACCGTCTCCCAGCTCGCACAC
GCGCACGCGTCGCACTCTCCAGCGCGCGTACTGTTCCAGCCACCGTCCCCATGCTCCCCG
CGGGGCCGTATGACGAGCCACGCCCCGGCCCCGCCGCCGCCGCTCGCCCACCGAGACAAG
CGCAATGCTATTGGGGCCTGACCACGGCTGGGCGGGGCGCTTCCCACGGAGGGCCCAGTC
TTTGGGGGGAGTTGCCCTTCCAGGGATCGGTAGCGCGGTGCGGCTCCCCGCAGCGGCCGC
GGGCGTCGGGGGGATGCTGGGGAGTGGGGGATGCCCCTTCCGGTCGCAGGCGGAACAGGT
TCGCGGTGGTGCACAGGGGCGTGGTGCTGCAGCTCGAGGCCGCGGAGGGGGGCCGTGGTG
GACTTCGGGTCGGAGGGACCGATATACCAGGGCTCTGGCGCGGGGTAGCCCCAGGACGCT
TTCTCGGGGAGGGGAGGGTACGGGCAGCCAGGTGGGAGTGCAAGAACTGGACGGCGTAAT
CCCTGCTGCCCGCGATGTGGGGTCGGACCCAGCGCACCCCGATGGGGCCTGGCGGCGGGG
GACCCCTGAACGGAGGTGGCAGCTACTCTGCCGTTGAGCCGATGCGGACGTGCCCAGTAC
ACGTGGGTGTTGGGGGAACTCCACAGCGGCAGGCGAGGGTTGGGTGTAGCAGGGGGCCCA
CGGGGGCGGCGGGTGGCAGGCGCGAGGGGGGCGCGGGGGAGCGATCCGATGGTCGAGAAC
GATCGGCTAGGAGGCGAGCCCCAGGTGGCGCCGGGCGGGTGCGCCCGTGTCGGCGCGGGG
GGGTTGTGGCGACGAGGCCGCCCTTCAGGCGAGGTGGGAGTGGCCCAAGTTGTGGCGCCG
CGGTGGAGGGGGGGCGGTGAGGTGGGGGGCCAGTCGAGGTTGGTGAGGGCGGGACCGGCC
TCGCTTAGTGCAACACCAACCCGGATAGTAATTCGTGCGGCTCGTGGGAGCGAGTTCTTC
CAATGATGCAGGGGGGGGTGTAGGGCCCGCGCGTCGACCGGCCGCGTCACGACTGCTACG
GGGTCATTACCGCCGGGCTGTCGTAGGCCGGCGCTCGCGGGAGGTGGTTCGAATGCCGGG
GCGGGAGGCCCGGATCCGGCGAAGGGGCGGGGCTCGGGGGACGCGAGGCGGAGGGGCGCG
AGGCGGTAGCCGGTGCAAGCGGCGCGTGGCCCCATGAGGAAGAACTCTCTCGTGCCCCAA
CACCACACATTAGCGCCCCCATCTCGGCCGACTTCGAGCACGCTCAGCGAGAACCACCAC
AGGCCTCCAACCCGACGTACCACCCGTTCTCACGAGGCGGGCCGACCCTCGGGTCCAAGC
GAGGGCTCCGGCGCGCACGCCCGGCGTGGTGC
>MAP 175_A_tb1171
CCCCTTAGGCGCCTCGAGCGGGCCTCCGCTTCGCCAGTTCACCCGTCCCAGTAAGCGCCC
GGGGCACCTCGCCTTCGAGGAGAACCCCCAGAACGCTCCACCGTCTCCCAGCTCGCACAC
GCGCACGCGTCGCACTCTCCAGCGCGCGTACTGTTCCAGCCACCGTCCCCATGCTCCCCG
CGGGGCCGTATGACGAGCCACGCCCCGGCCCCGCCGCCGCCGCTCGCCCACCGAGACAAG
CGCAATGCTATTGGGGCCTGACCACGGCTGGGCGGGGCGCTTCCCACGGAGGGCCCAGTC
TTTGGGGGGAGTTGCCCTTCCAGGGATCGGTAGCGCGGTGCGGCTCCCCGCAGCGGCCGC
GGGCGTCGGGGGGATGCTGGGGAGTGGGGGATGCCCCTTCCGGTCGCAGGCGGAACAGGT
TCGCGGTGGTGCACAGGGGCGTGGTGCTGCAGCTCGAGGCCGCGGAGGGGGGCCGTGGTG
GACTTCGGGTCGGAGGGACCGATATACCAGGGCTCTGGCGCGGGGTAGCCCCAGGACGCT
TTCTCGGGGAGGGGAGGGTACGGGCAGCCAGGTGGGAGTGCAAGAACTGGACGGCGTAAT
CCCTGCTGCCCGCGATGTGGGGTCGGACCCAGCGCACCCCGATGGGGCCTGGCGGCGGGG
GACCCCTGAACGGACGTGGCAGCTACTCTGCCGTTGAGCCGATGCGGACGTGCCCAGTAC
ACGTGGGTGTTGGGGGAACTCCACAGCGGCAGGCGAGGGTTGGGTGTAGCAGGGGGCCCA
CGGGGGCGGCGGGTGGCAGGCGCGAGGGGGGCGCGGGGGAGCGATCCGATGGTCGAGAAC
GATCGGCTAGGAGGCGAGCCCCAGGTGGCGCCGGGCGGGTGCGCCCGTGTCGGCGCGGGG
GGGTTGTGGCGACGAGGCCGCCCTTCAGGCGAGGTGGGAGTGGCCCAAGTTGTGGCGCCG
CGGTGGAGGGGGGGCGGTGAGGTGGGGGGCCAGTCGAGGTTGGTGAGGGCGGGACCGGCC
TCGCTTAGTGCAACACCAACCCGGATAGTAATTCGTGCGGCTCGTGGGAGCGAGTTCTTC
CAATGATGCAGGGGGGGGTGTAGGGCCCGCGCGTCGACCGGCCGCGTCACGACTGCTACG
GGGTCATTACCGCCGGGCTGTCGTAGGCCGGCGCTCGCGGGAGGTGGTTCGAATGCCGGG
GCGGGAGGCCCGGATCCGGCGAAGGGGCGGGGCTCGGGGGACGCGAGGCGGAGGGGCGCG
AGGCGGTAGCCGGTGCAAGCGGCGCGTGGCCCCATGAGGAAGAACTCTCTCGTGCCCCAA
CACCACACATTAGCGCCCCCATCTCGGCCGACTTCGAGCACGCTCAGCGAGAACCACCAC

AGGCCTCCAACCCGACGTACCACCCGTTCTCACGAGGCGGGCCGACCCTCGGGTCCAAGC
GAGGGCTCCGGCGCGCACGCCCGGCGTGGTGC
>MAP 133_A_tb1504
CCCCTTAAGCGCCTCGAGCGGGCCCCCGCCCCTCCAGTTCACCCGTCCCAGTAAGCGCCT
GCGGCCTCACGCCTTCGCGGAGAACACCCAGAACGCCCCACCCTATCCTAGGGCGCACAC
GCGCACGCGTCGCATTCTCCAGCGCGCGGGCTGTTCTAGCCACCGTCCCCATGCTCCCCG
CGGGGCCGTATGACGAGCCACGCCCCGGCCCCGCCGCCGCCGCTGGCCCACCGAGACAAG
CGCAACGCTGTTGGGGCCTGACCACGGCTGGGCGGGGCGCTTCCCACGGCGGGCCCAGTC
TTTGGGAGGAGTTGTCCCTCAAGGGATCGGTAGCGCGGTGCGGCTCCCCGCAGCGGCCGC
GGGCGTCGGGGTGATGCTGGGGAGTGGGGGATGCCCCTTCCGGCCGCAGGCGGAACAGGT
TCGCGGTGGTGCACAGGGGCGTGGTGCTGCAGCTCGAGGCCGCAGAGGGGGGCCGCGGTG
GACTTCGGGTCGGAGGGACCGATATACCAGGGCTCTGGCGCGGGGTAGCCCCAGGACGCT
CTCTCGGGGAGGGGAGGGTACTGGCAGCCAGGTGGGAGTGCAGGAACTGGACGGCGTAAT
CCCTGGTGACTGCGATGTTGGGTAGGACCCAGCGCGCCCTGATGGGGCCTGGCGGCGGGG
GACCCCTGGATCGACGTCGCAGCTACCCTGCCGTTGGGCCGACGCGGTCGTGCCCAGTAC
ACGTGGGTGTTGGGGGAACTCCACAGGGGTAGGCGAGTGTTGGGTGTGGCCGGGGACCCA
CGCGGGCGGGGGGTGGCGGTCGCGAGGGGGGCGTGGGGGAGCGATCCGATGGTCGAGAAC
GATCGGATAGGAGGCAAGACCCAGGCGGCGCAGGGCGGGTGCGCCCGTGTCGGCGCGGGG
GGGTTGTGGCAACAAGGCCGCCCTTCAGGCGGGGTGGGAGTGGCCCAAGTTGTGGCGCCG
CGGTGGAGGGGGGGCGGTGAGGTGGGGGGCCAGTCGGGGTTGGTGAGGGCGGGACCAGCC
TCGCTTGGTGCAACGCCAACCCGGATAGTAATTCGTGCGGCTCGTGGGCGCGAATTCTTC
CAATGACTCAGAGGGGGGTGTGGGGCCCGAGCGTCGACCGGCCGCGTCACGGCTGCTACG
GGGTCATTACCGCCGGGCTGTCGTAGGCGGGCGATCGCGGGAGGTGGTTCGAATGCCGGG
GCCGGAGGCCCGGATCAGGCGAAGGGGCGGGGCCCGGGGCACGCAAGGCGGGGGGGCGCG
AGGCGGTAGCCGGTGCAAGCGGCGCGTGTCCCCATGAGGTAAAACTCTCTCGTCCCCCAC
CACCACACATTAGCGCCTCCATCTCGGCCGACTTCGAGCGCGTTCAGGGAGAACCACCAC
AGGCCTCCAACCCGACGTACCACCCGTCCTCACGAGGCGGGCCGACCCCCGGGCCCAAGC
GAGGGCTCCGGCGAGCACGCGCGGCGTGGTGC
>MAP 134_A_tb1504
CCCCTTAAGCGCCTCGAGCGGGCCCCCGCCCCTCCAGTTCACCCGTCCCAGTAAGCGCCT
GCGGCCTCACGCCTTCGCGGAGAACACCCAGAACGCCCCACCCTATCCTAGGGCGCACAC
GCGCACGCGTCGCATTCTCCAGCGCGCGGGCTGTTCTAGCCACCGTCCCCATGCTCCCCG
CGGGGCCGTATGACGAGCCACGCCCCGGCCCCGCCGCCGCCGCTGGCCCACCGAGACAAG
CGCAACGCTGTTGGGGCCTGACCACGGCTGGGCGGGGCGCTTCCCACGGCGGGCCCAGTC
TTTGGGAGGAGTTGTCCCTCAAGGGATCGGTAGCGCGGTGCGGCTCCCCGCAGCGGCCGC
GGGCGTCGGGGTGATGCTGGGGAGTGGGGGATGCCCCTTCCGGCCGCAGGCGGAACAGGT
TCGCGGTGGTGCACAGGGGCGTGGTGCTGCAGCTCGAGGCCGCAGAGGGGGGCCGCGGTG
GACTTCGGGTCGGAGGGACCGATATACCAGGGCTCTGGCGCGGGGTAGCCCCAGGACGCT
CTCTCGGGGAGGGGAGGGTACTGGCAGCCAGGTGGGAGTGCAGGAACTGGACGGCGTAAT
CCCTGGTGACTGCGATGTTGGGTAGCACCCAGCGCGCCCTGATGGGGCCTGGCGGCGGGG
GACCCCTGGATCGACGTCGCAGCTACCCTGCCGTTGGGCCGACGCGGTCGTGCCCAGTAC
ACGTGGGTGTTGGGGGAACTCCACAGGGGTAGGCGAGTGTTGGGTGTGGCCGGGGACCCA
CGCGGGCGGGGGGTGGCGGTCGCGAGGGGGGCGTGGGGGAGCGATCCGATGGTCGAGAAC
GATCGGATAGGAGGCAAGACCCAGGCGGCGCAGGGCGGGTGCGCCCGTGTCGGCGCGGGG
GGGTTGTGGCAACAAGGCCGCCCTTCAGGCGGGGTGGGAGTGGCCCAAGTTGTGGCGCCG
CGGTGGAGGGGGGGCGGTGAGGTGGGGGGCCAGTCGGGGTTGGTGAGGGCGGGACCAGCC
TCGCTTGGTGCAACGCCAACCCGGATAGTAATTCGTGCGGCTCGTGGGCGCGAATTCTTC
CAATGACTCAGAGGGGGGTGTGGGGCCCGAGCGTCGACCGGCCGCGTCACGGCTGCTACG
GGGTCATTACCGCCGGGCTGTCGTAGGCGGGCGATCGCGGGAGGTGGTTCGAATGCCGGG
GCCGGAGGCCCGGATCAGGCGAAGGGGCGGGGCCCGGGGCACGCAAGGCGGGGGGGCGCG
AGGCGGTAGCCGGTGCAAGCGGCGCGTGTCCCCATGAGGTAAAACTCTCTCGTCCCCCAC
CACCACACATTAGCGCCTCCATCTCGGCCGACTTCGAGCGCGTTCAGGGAGAACCACCAC
AGGCCTCCAACCCGACGTACCACCCGTCCTCACGAGGCGGGCCGACCCCCGGGCCCAAGC
GAGGGCTCCGGCGAGCACGCGCGGCGTGGTGC
>MAP 176_A_tb1532
CCCCTTAGGCGCCTCGAGCGGGCCTCCGCTTCGCCAGTTCACCCGTCCCAGTAAGCGCCC
GGGGCACCTCGCCTTCGAGGAGAACCCCCAGAACGCCCCACCGTCTCCCAGCTCGCACAC
GCGCACGCGTCGCACTCTCCAGCGCGCGTACTGTTCCAGCCACCGTCCCCATGCTCCCCG
CGGGGCCGTATGACGAGCCACGCCCCGGCCCCGCCGCCGCCGCTCGCCCACCGAGACAAG
CGCAATGCTATTGGGGCCTGACCACGGCTGGGCGGGGCGCTTCCCACGGAGGGCCCAGTC
TTTGGGGGGAGTTGCCCTTCCAGGGATCGGTAGCGCGGTGCGGCTCCCCGCAGCGGCCGC
GGGCGTCGGGGGGATGCTGGGGAGTGGGGGATGCCCCTTCCGGTCGCAGGCGGAACAGGT
TCGCGGTGGTGCACAGGGGCGTGGTGCTGCAGCTCGAGGCCGCGGAGGGGGGCCGTGGTG
GACTTCGGGTCGGAGGGACCGATATACCAGGGCTCTGGCGCGGGGTAGCCCCAGGACGCT
TTCTCGGGGAGGGGAGGGTACGGGCAGCCAGGTGGGAGTGCAAGAACTGGACGGCGTAAT
CCCTGCTGCCCGCGATGTGGGGTCGGACCCAGCGCACCCCGATGGGGCCTGGCGGCGGGG
GACCCCTGAACGGACGTGGCAGCTACTCTGCCGTTGAGCCGATGCGGACGTGCCCAGTAC
ACGTGGGTGTTGGGGGAACTCCACAGCGGCAGGCGAGGGTTGGGTGTAGCAGGGGGCCCA

CGGGGGCGGCGGGTGGCAGGCGCGAGGGGGGCGCGGGGGAGCGATCCGATGGTCGAGAAC
GATCGGCTAGGAGGCGAGCCCCAGGTGGCGCCGGGCGGGTGCGCCCGTGTCGGCGCGGGG
GGGTTGTGGCGACGAGGCCGCCCTTCAGGCGAGGTGGGAGTGGCCCAAGTTGTGGCGCCG
CGGTGGAGGGGGGGCGGTGAGGTGGGGGGCCAGTCGAGGTTGGTGAGGGCGGGACCGGCC
TCGCTTAGTGCAACACCAACCCGGATAGTAATTCGTGCGGCTCGTGGGAGCGAGTTCTTC
CAATGATGCAGGGGGGGGTGTAGGGCCCGCGCGTCGACCGGCCGCGTCACGACTGCTACG
GGGTCATTGCCGCCGGGCTGTCGTAGGCCGGCGCTCGCGGGAGGTGGTTCGAATGCCGGG
GCGGGAGGCCCGGATCCGGCGAAGGGGCGGGGCTCGGGGGACGCGAGGCGGAGGGGCGCG
AGGCGGTAGCCGGTGCAAGCGGCGCGTGGCCCCATGAGGAAGAACTCTCTCGTGCCCCAA
CACCACACATTAGCGCCCCCATCTCGGCCGACTTCGAGCACGCTCAGCGAGAACCACCAC
AGGCCTCCAACCCGACGTACCACCCGTTCTCACGAGGCGGGCCGACCCTCGGGTCCAAGC
GAGGGCTCCGGCGCGCACGCCCGGCGTGGTGC
>MAP 177_A_tb1532
CCCCTTAGGCGCCTCGAGCGGGCCTCCGCTTCGCCAGTTCACCCGTCCCAGTAAGCGCCC
GGGGCACCTCGCCTTCGAGGAGAACCCCCAGAACGCCCCACCGTCTCCCAGCTCGCACAC
GCGCACGCGTCGCACTCTCCAGCGCGCGTACTGTTCCAGCCACCGTCCCCATGCTCCCCG
CGGGGCCGTATGACGAGCCACGCCCCGGCCCCGCCGCCGCCGCTCGCCCACCGAGACAAG
CGCAATGCTATTGGGGCCTGACCACGGCTGGGCGGGGCGCTTCCCACGGAGGGCCCAGTC
TTTGGGGGGAGTTGCCCTTCCAGGGATCGGTAGCGCGGTGCGGCTCCCCGCAGCGGCCGC
GGGCGTCGGGGGGATGCTGGGGAGTGGGGGATGCCCCTTCCGGTCGCAGGCGGAACAGGT
TCGCGGTGGTGCACAGGGGCGTGGTGCTGCAGCTCGAGGCCGCGGAGGGGGGCCGTGGTG
GACTTCGGGTCGGAGGGACCGATATACCAGGGCTCTGGCGCGGGGTAGCCCCAGGACGCT
TTCTCGGGGAGGGGAGGGTACGGGCAGCCAGGTGGGAGTGCAAGAACTGGACGGCGTAAT
CCCTGCTGCCCGCGATGTGGGGTCGGACCCAGCGCACCCCGATGGGGCCTGGCGGCGGGG
GACCCCTGAACGGACGTGGCAGCTACTCTGCCGTTGAGCCGATGCGGACGTGCCCAGTAC
ACGTGGGTGTTGGGGGAACTCCACAGCGGCAGGCGAGGGTTGGGTGTAGCAGGGGGCCCA
CGGGGGCGGCGGGTGGCAGGCGCGAGGGGGGCGCGGGGGAGCGATCCGATGGTCGAGAAC
GATCGGCTAGGAGGCGAGCCCCAGGTGGCGCCGGGCGGGTGCGCCCGTGTCGGCGCGGGG
GGGTTGTGGCGACGAGGCCGCCCTTCAGGCGAGGTGGGAGTGGCCCAAGTTGTGGCGCCG
CGGTGGAGGGGGGGCGGTGAGGTGGGGGGCCAGTCGAGGTTGGTGAGGGCGGGACCGGCC
TCGCTTAGTGCAACACCAACCCGGATAGTAATTCGTGCGGCTCGTGGGAGCGAGTTCTTC
CAATGATGCAGGGGGGGGTGTAGGGCCCGCGCGTCGACCGGCCGCGTCACGACTGCTACG
GGGTCATTACCGCCGGGCTGTCGTAGGCCGGCGCTCGCGGGAGGTGGTTCGAATGCCGGG
GCGGGAGGCCCGGATCCGGCGAAGGGGCGGGGCTCGGGGGACGCGAGGCGGAGGGGCGCG
AGGCGGTAGCCGGTGCAAGCGGCGCGTGGCCCCATGAGGAAGAACTCTCTCGTGCCCCAA
CACCACACATTAGCGCCCCCATCTCGGCCGACTTCGAGCACGCTCAGCGAGAACCACCAC
AGGCCTCCAACCCGACGTACCACCCGTTCTCACGAGGCGGGCCGACCCTCGGGTCCAAGC
GAGGGCTCCGGCGCGCACGCCCGGCGTGGTGC
>MAP 178_A_tb1532
CCCCTTAGGCGCCTCGAGCGGGCCTCCGCTTCGCCAGTTCACCCGTCCCAGTAAGCGCCC
GGGGCACCTCGCCTTCGAGGAGAACCCCCAGAACGCCCCACCGTCTCCCAGCTCGCACAC
GCGCACGCGTCGCACTCTCCAGCGCGCGTACTGTTCCAGCCACCGTCCCCATGCTCCCCG
CGGGGCCGTATGACGAGCCACGCCCCGGCCCCGCCGCCGCCGCTCGCCCACCGAGACAAG
CGCAATGCTATTGGGGCCTGACCACGGCTGGGCGGGGCGCTTCCCACGGAGGGCCCAGTC
TTTGGGGGGAGTTGCCCTTCCAGGGATCGGTAGCGCGGTGCGGCTCCCCGCAGCGGCCGC
GGGCGTCGGGGGGATGCTGGGGAGTGGGGGATGCCCCTTCCGGTCGCAGGCGGAACAGGT
TCGCGGTGGTGCACAGGGGCGTGGTGCTGCAGCTCGAGGCCGCGGAGGGGGGCCGTGGTG
GACTTCGGGTCGGAGGGACCGATATACCAGGGCTCTGGCGCGGGGTAGCCCCAGGACGCT
TTCTCGGGGAGGGGAGGGTACGGGCAGCCAGGGGGGGAGTGCAAGAACTGGACGGCGTAAT
CCCTGCTGCCCGCGATGTGGGGTCGGACCCAGCGCACCCCGATGGGGCCTGGCGGCGGGG
GACCCCTGAACGGACGTGGCAGCTACTCTGCCGTTGAGCCGATGCGGACGTGCCCAGTAC
ACGTGGGTGTTGGGGGAACTCCACAGCGGCAGGCGAGGGTTGGGTGTAGCAGGGGGCCCA
CGGGGGCGGCGGGTGGCAGGCGCGAGGGGGGCGCGGGGGAGCGATCCGATGGTCGAGAAC
GATCGGCTAGGAGGCGAGCCCCAGGTGGCGCCGGGCGGGTGCGCCCGTGTCGGCGCGGGG
GGGTTGTGGCGACGAGGCCGCCCTTCAGGCGAGGTGGGAGTGGCCCAAGTTGTGGCGCCG
CGGTGGAGGGGGGGCGGTGAGGTGGGGGGCCAGTCGAGGTTGGTGAGGGCGGGACCGGCC
TCGCTTAGTGCAACACCAACCCGGATAGTAATTCGTGCGGCTCGTGGGAGCGAGTTCTTC
CAATGATGCAGGGGGGGGTGTAGGGCCCGCGCGTCGACCGGCCGCGTCACGACTGCTACG
GGGTCATTACCGCCGGGCTGTCGTAGGCCGGCGCTCGCGGGAGGTGGTTCGAATGCCGGG
GCGGGAGGCCCGGATCCGGCGAAGGGGCGGGGCTCGGGGGACGCGAGGCGGAGGGGCGCG
AGGCGGTAGCCGGTGCAAGCGGCGCGTGGCCCCATGAGGAAGAACTCTCTCGTGCCCCAA
CACCACACATTAGCGCCCCCATCTCGGCCGACTTCGAGCACGCTCAGCGAGAACCACCAC
AGGCCTCCAACCCGACGTACCACCCGTTCTCACGAGGCGGGCCGACCCTCGGGTCCAAGC
GAGGGCTCCGGCGCGCACGCCCGGCGTGGTGC
