## Supplementary material for "Who infects Whom? - Reconstructing infection chains of *Mycobacterium avium* ssp. *paratuberculosis* in an endemically infected dairy herd by use of genomic data": S2 File.R

```
## Supporting information S2 File

## This is the R code for the analyis of following research article:
## Title: "Who infects Whom": Reconstructing infection chains of
Mycobacterium avium subsp. paratuberculosis in an
##        endemically infected dairy herd by use of genomic data
## by: Annette Nigsch, Suelee Robbe-Austerman, Tod Stuber, Paulina D.
Pavinski Bitar, Yrjö Gröhn and Ynte H. Schukken



## 1.) PREPARATIONS

## for details on SeqTrack, see SeqTrack helpfile and Jombart et al.
(2011), Heredity 106, 383-390.

## load packages
library("ape")
library("adegenet")
library("igraph")
library("ggplot2")
library("plyr")

## set working directory on your computer
## setwd("C:/Users/.../MAP")

## load the SNP sequence data
snp <-read.dna("MAP.fasta", format="fa")


## load case file
# columns:  id = UID, label = sequence ID, cow = cow ID, birthd = birth
date of cow,
#           shedd = potential start of the (genotype-specific) infectious
period,
cases <- read.csv("MAP_cases.CSV")


## define the columms with dates as GMT dates in POSIXct format.
birth <- as.POSIXct(cases$birthd, tz = "GMT")
shed <- as.POSIXct(cases$shedd, tz = "GMT")  # time zone (tz) in Universal
Time, Coordinated (UTC), which is GMT


## create distance matrix
distmat <- dist.dna(snp, model="N", pairwise.deletion = TRUE, as.matrix =
TRUE)
write.csv(distmat, "DistMat_pairwDeletion.csv")


## define length of nucleotide sequence and mutation rate mu
nbNucl <- ncol(as.matrix(snp))
mu_MAP <- 2.91e-08*2/365    ## 0.25 substitutions per genome per year


## load weighting matrices for Exposure [E] and Susceptibility [S]
scenarios
# MAP_E.csv: cells A1:DX1 = isolate ID, cells A2:DX129 = number of exposure
days between pairs of isolates [E]
# MAP_S.csv: cells A1:DX1 = isolate ID, cells A2:DX129 = weights from 6 to
0 for seven social network pattern [S]
Matrix_E <- read.csv("MAP_E.csv")
Matrix_S <- read.csv("MAP_S.csv")


## convert data into matrices
M_E <- as.matrix(Matrix_E)
M_S <- as.matrix(Matrix_S)
```

```
## 2.) RECONSTRUCTION OF MAP TRANSMISSION TREES WITH SEQTRACK

## select time stamp for the scenario to calculate (shed or birth), start
with "birth".
# then repeat lines 67 - 91 with "shed" and continue in line 99.
Date <- birth # shed


## run SeqTrack analysis for [Basic], [E] and [S] scenarios
res_Basic <- seqTrack(distmat, x.names=cases$label, x.dates=Date,
mu=mu_MAP, haplo.le=nbNucl)
res_E <- seqTrack(distmat, x.names=cases$label, x.dates=Date, prox.mat=M_E,
mu=mu_MAP, haplo.le=nbNucl)
res_S <- seqTrack(distmat, x.names=cases$label, x.dates=Date, prox.mat=M_S,
mu=mu_MAP, haplo.le=nbNucl)


## calculate statistical support for inferred ancestries
p_Basic <- get.likelihood(res_Basic, mu=mu_MAP, haplo.length=nbNucl)
p_E <- get.likelihood(res_E, mu=mu_MAP, haplo.length=nbNucl)
p_S <- get.likelihood(res_S, mu=mu_MAP, haplo.length=nbNucl)


# replace all ancestors with weight > "x" with NA. In the article we used x
= 6.
x <- 6
res_Basic$ances[res_Basic$weight > x] <- NA
res_E$ances[res_E$weight > x] <- NA
res_S$ances[res_S$weight > x] <- NA


## create summary files: "ares" contains ancestries, "ap" contains p-values
of statistical support for ancestries
ares <- data.frame(res_Basic$ances, res_E$ances, res_S$ances)
ap <- data.frame(p_Basic, p_E, p_S)


## save time stamp-specific summary files, begin with "birth"
ares_birth <- ares
ap_birth <- ap

# then repeat lines 67 - 91 with "shed" and continue in line 99, save
"shed" summary files.
ares_shed <- ares
ap_shed <- ap


## graphs
## plot pairwise genomic distance of all snp sequences (Fig. 2 in article)
hist <- hist(distmat,  col="lightgrey", nclass=250, xlim = c(0,250), ylim =
c(0,1200),
             main="Distribution of pairwise genomic distances",
             xlab="Number of differing SNPs")

## plot SeqTrack transmission trees once with "birth" and once with "shed"
as time stamp (Fig. 3. Fig. 4, Fig. S.1 in article)
res_graph <- as.igraph(res_Basic)
tkplot(res_graph, vertex.color="white") # layout transmission tree in newly
opened window

res_graph <- as.igraph(res_E)
tkplot(res_graph, vertex.color="white")

res_graph <- as.igraph(res_S)
```

```
tkplot(res_graph, vertex.color="white")


## 3.) ANALYSIS OF RECONSTRUCTED MAP TRANSMISSION TREES

## create table with number of descending isolates and number of offspring
# select time stamp for the scenario to calculate (shed or birth), start
with "ares_birth".
# repeat lines 125 - 138 with "ares_shed" and continue in line 146.
a <- ares_birth  # ares_shed
ancesfreq_Basic <- as.data.frame(table(a$res_Basic))
ancesfreq_E <- as.data.frame(table(a$res_E))
ancesfreq_S <- as.data.frame(table(a$res_S))

cas <- data.frame(cow=cases$cow, MAP_id=cases$id)
cas$freq_Basic = ancesfreq_Basic$Freq[match(cases$id, ancesfreq_Basic
$Var1)]  # gives number of descending isolates at isolate level
cas$freq_E = ancesfreq_E$Freq[match(cases$id, ancesfreq_E$Var1)]
cas$freq_S = ancesfreq_S$Freq[match(cases$id, ancesfreq_S$Var1)]
cas[is.na(cas)] <- 0
# replaces all "NA" by 0

cow_freq_Basic <- count(cas, "cow", "freq_Basic")
# gives number of descending isolates at cow level
cow_freq_E <- count(cas, "cow", "freq_E")
cow_freq_S <- count(cas, "cow", "freq_S")


cow_freqb <- data.frame(cow=cow_freq_Basic$cow, b_Basic=cow_freq_Basic
$freq, b_E= cow_freq_E$freq,
                        b_S= cow_freq_S$freq)
cas_birth <- cas

# repeat lines 125 - 138 with "ares_shed" and continue in line 146, save
"shed" results.
cow_freqs <- data.frame(cow=cow_freq_Basic$cow, s_Basic=cow_freq_Basic
$freq, s_E= cow_freq_E$freq,
                        s_S= cow_freq_S$freq)
cas_shed <- cas

## save results for data handling and further analysis in R or in another
software
# data handling outside R: accout for potential right and left censoring of
data by using
# a subsample of all isolates to investigate the role of an individual cow
in MAP spread (see 4.)
cow_freqbs <- data.frame(cow_freqb, cow_freqs)
cas_bs <- data.frame(b=cas_birth, e=cas_shed)

write.csv(cas_bs, "cas_bs.csv")
write.csv(cow_freqbs, "cow_freqbs.csv")


## 4.) CORRELATION BETWEEN RECONSTRUCTED NUMBER OF OFFSPRING AND DISEASE
PHENOTYPES

## load cow disease phenotypes and concatenate them to final (adapted)
table with number of offspring per cow.
# format of table with cow disease phenotypes: "cow_id", "disease
phenotypes" in binary or categorical coding.
# make sure that both tables are sorted in the same order of cows.
pheno <- read.csv("MAP_disease_phenotypes.CSV")
cow_freqbsx <- read.csv("Cow_freqbs_adapt.CSV")
phenodf <- data.frame(pheno, cow_freqbsx)
write.csv(phenodf, "Cow_freqbsx_phenodf.csv")

## multifactorial analysis: correlate number of offspring (by scenario)
with disease phenotype
```

```
# categorical coded phenotype - example: "shedding level"
cor.test(phenodf$b_Basic,phenodf$Sheddinglevel, method="spearman")
cor.test(phenodf$b_E,phenodf$Sheddinglevel, method="spearman")
cor.test(phenodf$b_S,phenodf$Sheddinglevel, method="spearman")
cor.test(phenodf$s_Basic,phenodf$Sheddinglevel, method="spearman")
cor.test(phenodf$s_E,phenodf$Sheddinglevel, method="spearman")
cor.test(phenodf$s_S,phenodf$Sheddinglevel, method="spearman")

# boxplots with numbers of offspring produced by individual cows, by
disease phenotype and scenario (Fig. 5 in article)
par(mfrow = c(1,6))
boxplot(phenodf$b_Basic~phenodf$Sheddinglevel,
        main="Shedding Level",
        ylab="Offspring (n)")
boxplot(phenodf$b_E~phenodf$Sheddinglevel)
boxplot(phenodf$b_S~phenodf$Sheddinglevel)
boxplot(phenodf$s_Basic~phenodf$Sheddinglevel)
boxplot(phenodf$s_E~phenodf$Sheddinglevel)
boxplot(phenodf$s_S~phenodf$Sheddinglevel)

# binary coded phenotype - example: "SeroStatus"
t.test(phenodf$b_Basic~phenodf$SeroStatus)
t.test(phenodf$b_E~phenodf$SeroStatus)
t.test(phenodf$b_S~phenodf$SeroStatus)
t.test(phenodf$s_Basic~phenodf$SeroStatus)
t.test(phenodf$s_E~phenodf$SeroStatus)
t.test(phenodf$s_S~phenodf$SeroStatus)

boxplot(phenodf$b_Basic~phenodf$SeroStatus,
        main="Serostatus",
        ylab="Offspring (n)")
boxplot(phenodf$b_E~phenodf$SeroStatus)
boxplot(phenodf$b_S~phenodf$SeroStatus)
boxplot(phenodf$s_Basic~phenodf$SeroStatus)
boxplot(phenodf$s_E~phenodf$SeroStatus)
boxplot(phenodf$s_S~phenodf$SeroStatus)


# end
```
